# OzFAD: Ozone-enabled fatty acid discovery reveals unexpected diversity in the human lipidome

**DOI:** 10.1101/2022.10.24.513604

**Authors:** Jan Philipp Menzel, Reuben S.E. Young, Aurélie H. Benfield, Julia Scott, Lisa M. Butler, Sónia Troeira Henriques, Berwyck L.J. Poad, Stephen J. Blanksby

## Abstract

Fatty acid isomers are responsible for an under-reported lipidome diversity across all kingdoms of life. Isomers of unsaturated fatty acid are often masked in contemporary analysis by incomplete separation and the absence of sufficiently diagnostic methods for structure elucidation. Here, we introduce a comprehensive workflow to discover new unsaturated fatty acids through coupling liquid chromatography and mass spectrometry with gas-phase ozonolysis of double bonds. The workflow encompasses semi-automated data analysis and enables *de novo* identification in complex media including human plasma, cancer cell lines and human sebaceous wax (i.e., vernix caseosa). The targeted analysis including ozonolysis enables structural assignment over a dynamic range of five orders of magnitude, even in instances of incomplete chromatographic separation. Thereby we expand the number of identified plasma fatty acids two-fold, including non-methylene interrupted fatty acids. Detection, without prior knowledge, allows discovery of non-canonical double bond positions. Changes in relative isomer abundances reflect underlying perturbations in lipid metabolism.

## INTRODUCTION

Advances in the molecular-level description of the genome, transcriptome, proteome, lipidome or metabolome over recent decades,^1^ has opened new frontiers in medicine, biology, and nutrition.^2^ The contemporary challenge of uncovering biomolecular diversity increases in complexity from genetics and proteomics to lipidomics due to the expanding array of molecular building blocks. Conversely, lipidomics thus presents a greater scope for biomolecular discovery due to the vast structural diversity of fatty acid building blocks.^3^ Moreover, many fatty acids remain entirely undescribed or widely underreported, due to limitations in chromatographic and mass spectrometric sensitivity and selectivity. Notably, unsaturated fatty acid isomers with varied double bond position and configuration make up a major part of the diversity across the categories of lipids, where they are responsible for potentially thousands of structurally distinct molecular species; many of which are isomers of each other and have not been reported with full structural assignment.^4^ Full lipidome coverage is a frontier challenge in lipidomics that can only be met by approaches that include a comprehensive survey of regio- and stereoisomeric fatty acid building blocks.^5^

Contemporary methods for fatty acid analysis are based on the hyphenation of chromatographic separation with mass spectrometric detection. Despite the high peak capacity of modern gas chromatography columns and ultra-performance reversed-phase liquid chromatography, many fatty acids remain incompletely resolved.^6^ In hyphenated chromatography-mass spectrometry methods, incomplete chromatographic resolution can be easily overcome when ionized lipids have different mass-to-charge ratios (*m/z*). For lipid isomers however, mass spectral discrimination relies on distinctive fragmentation patterns arising from ionization or ion activation. Unfortunately, the archetype ion activation approaches for fatty acids, namely electron ionization (for GC-MS) and collision-induced dissociation (CID, for LC-MS), most often yield indistinguishable mass spectra for isomers of unsaturated fatty acids.^7^

Efforts toward mass spectral discrimination of isomeric fatty acids can broadly be grouped into two categories. The first group of ion activation strategies have been optimized to promote wide-ranging fragmentation of carbon-carbon bonds resulting in a rich fragmentation pattern that is interrupted at, or adjacent to, the site(s) of unsaturation. Examples of these approaches include; (i) GC-MS strategies exploiting electron ionization of picolinyl esters (pyridyl carbinol derivatives) and dimethyloxazoline derivatives of fatty acids,^8^ and (ii) LC-MS approaches such as electron impact excitation of ions from organics,^9^ fixed-charge derivatives that promote charge-remote fragmentation^10^ and laser-based radical-directed dissociation methods.^11^ These methods generate complex mass spectra and thus typically require full chromatographic separation of each isomer or are reliant on spectral libraries from analytical standards, because the overlay of complex spectra containing common product ions prevents unambiguous identification of individual isomers within such mixtures.

The second group of methods takes advantage of selective fragmentation of ionized fatty acids at, or adjacent to, carbon-carbon double bonds. Examples of this approach in the context of GC-MS include off-line dimethyl disulfide derivatization,^12^ and online gas-phase derivatization at the double bond *via* covalent-adduct chemical ionization mass spectrometry.^13,14^ Analogous LC-MS approaches include derivatization at the site(s) of unsaturation *via* Paternò-Büchi reactions that are often conducted between the chromatograph and the mass spectrometer.^15,16^ Once chemically activated by derivatization, carbon-carbon double bonds are cleaved by CID leading to two characteristic product ions demarking each double bond position. Two further methods in this category are direct UV-photodissociation of ionized lipids^17^ and the gas-phase ion-molecule reaction with ozone; so-called ozone-induced dissociation (OzID).^18-22^ The latter is particularly suited for *de novo* identification of previously unknown regio- and stereoisomers at a high dynamic range, because mass selection of the fatty acid species can be performed prior to the ion activation and mass spectral analysis. In particular, the diagnostic OzID ions of any possible regioisomer are predictable and distinct, resulting in tandem mass spectra that allow *de novo* structural assignment across a wide dynamic range. Prior applications of OzID to a diverse range of lipids has led to numerous discoveries of novel lipids, confirming that current understanding of the natural diversity in fatty acid building blocks is incomplete.^23^

Systematically surveying all fatty acids in complex lipidomes critically relies on robust data acquisition and automation in data analysis that does not introduce assumptions on possible structural diversity. Several software packages have been developed for lipidomics and metabolomics applications, such as the LC-MS based computational lipid library software LipidBlast,^24^ the intact lipid analysis software MS-Dial 4,^25^ the lipid class separation based quantification workflow LipidQuant,^26^ the shotgun lipidomics based tools ALEX^27^ and LipidXplorer^28^ among other tools.^29,30^ These tools predominantly focus on the analysis of intact lipids, often only assigning fatty acyl chains at the level of numbers of carbons and the degree of unsaturation. Consequently, in addition to the analytical strategies above there is a need for specific software tools to empower the discovery of fatty acid regio- and stereoisomers to fully map the lipidome.

Herein we introduce the combination of charge-tagged fatty acid analysis *via* LC-OzID and tailor-made source code for highly automated data analysis as an end-to-end workflow for the *de novo*, library independent, discovery of fatty acid isomers for complete fatty acid profiling. Termed ozone-enabled fatty acid discovery, or OzFAD, the analytical pipeline presented here is specifically created to interrogate the structural diversity of fatty acids, including double bond positional and configurational information. The data analysis workflow is engineered around the Skyline Mass Spectrometry Environment^31-34^ to allow intermittent user input and validation. The OzFAD methods are here evaluated against selected aspects of the human lipidome, namely pooled human plasma, sebaceous lipids (vernix caseosa) and human derived cancer cells (cell culture).

## RESULTS

### Ozone-enabled Fatty Acid Discovery enables *de novo* identification of fatty acid double bond isomers

Here we introduce a comprehensive workflow as a discovery platform for fatty acids in biological samples. OzFAD is a sophisticated approach to semi-automated data analysis including target list creation for data dependent acquisition of LC-OzID-MS/MS spectra that offers facile access to assigned spectra and relative quantitation without prior knowledge (Fig. 1). The workflow is based on the extraction of lipids from a biological matrix following the protocol of Matyash *et al*.,^35^ subsequent hydrolysis and derivatization of fatty acids with a positive fixed charge as described by Bollinger *et al*.^10^ (Fig. 1a and b). Samples are loaded onto reversed-phase LC and analyzed by a travelling-wave ion mobility mass spectrometer^21^ modified to deliver ozone to the ion-mobility region to promote efficient gas-phase ion-molecule reactions. In this data-independent acquisition (DIA) there is no mass selection of ionized lipids, but all ions are subjected to ozonolysis and the resulting aldehyde and Criegee product ions arising from unsaturated fatty acids are assigned to precursor ions by retention time alignment (Fig. 1c and d). Conducting identification and relative quantification based only on the DIA approach (*vide infra*) is possible but challenging. In the absence of mass selection of the precursors, OzID product ions could be assigned to more than one coeluting fatty acid. Thus, to enhance confidence in structural assignments and to lower the limits of detection, a data-dependent acquisition(DDA), an LC-OzID-MS/MS experiment, is introduced. To accommodate this, the workflow branches into analysis of the data-independent acquisition (Fig. 1d), and generation of a target list for data-dependent acquisition (Fig. 1e).

**Figure 1:**
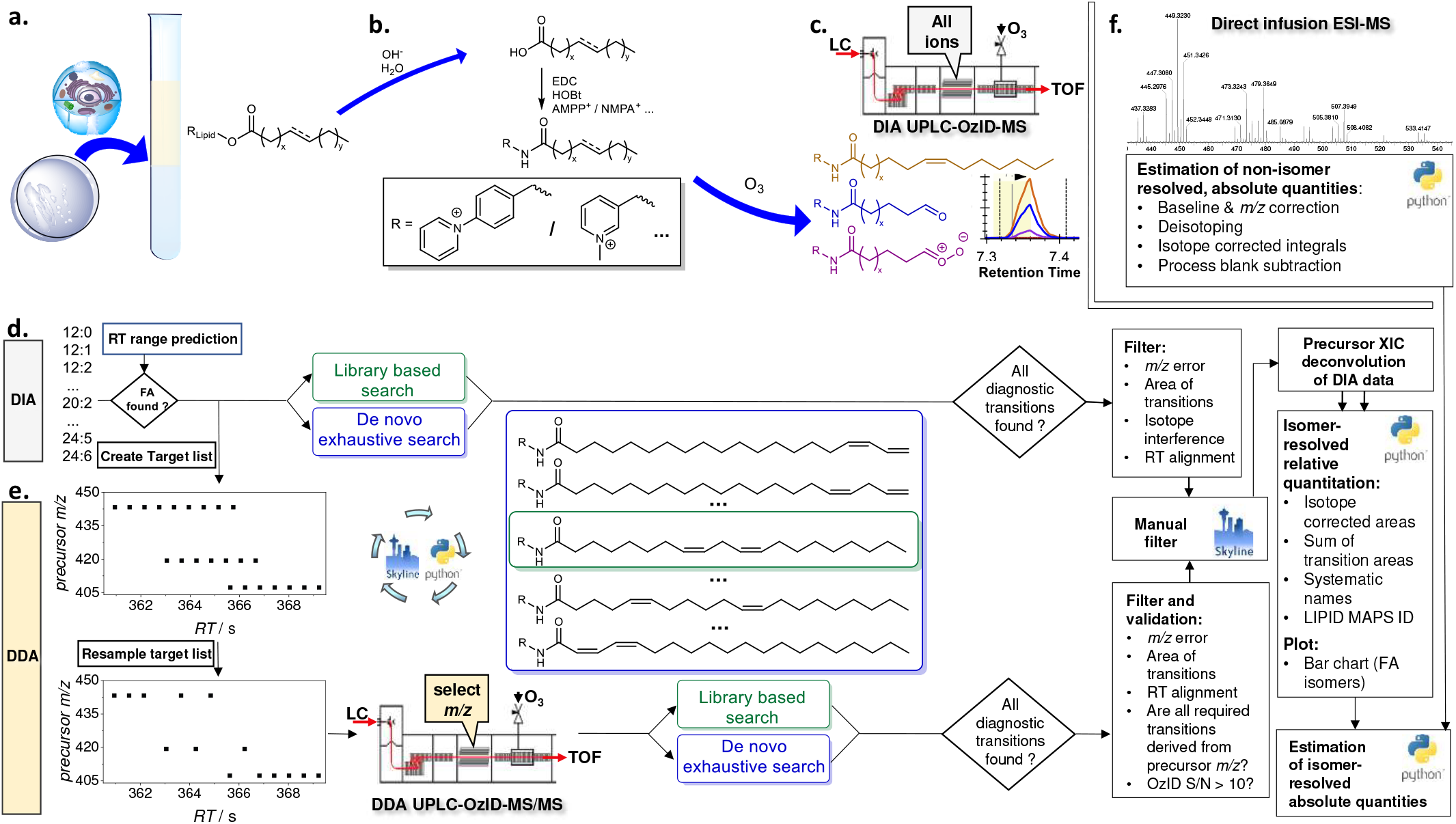
*De-novo* workflow for semi-automated fatty acid analysis with isomer resolution. **a**. Lipids are extracted from cell cultures, sebaceous waxes or human blood plasma. **b**. After hydrolysis of lipids and addition of internal standards, fatty acids are derivatized with a fixed charge. **c**. Liquid chromatography separates derivatized fatty acids, that are then ionized by electrospray ionization (ESI) and subjected to ozone-induced dissociation (OzID) with subsequent mass analysis (data-independent acquisition: DIA LC-OzID-MS). **d**. Analysis of the DIA LC-OzID-MS dataset is initiated by a windows batch file that controls python scripts and instances of Skyline Runner. First, the retention times of precursors (unfragmented fatty acids) that can be identified in the dataset are saved. **e**. Second, a target list (for a separate DDA LC-OzID-MS/MS run) is built based on precursor *m/z* values and retention times. After acquisition, an exhaustive search for all possible double bond positions (*i*.*e*., OzID product ions) and an automated filtering of the data is carried out. Manual inspection in Skyline allows the deletion of remaining false positives. Relative quantification is based on the DIA data including a manual correction of the deconvolution of extracted ion chromatograms. Finally, a python script formats the data, generates systematic names, retrieves LIPID MAPS IDs and common names from the online database (Lipidomics Gateway)^36^ where available and generates a bar chart. The latter visualizes the relative abundance of fatty acid isomers and their double bond positions. **f**. Direct infusion ESI-MS allows an estimation of abundance of the fatty acids without isomer resolution and combined with the relative abundance, an estimation of the absolute quantities of each fatty acid species.

Applying an initial retention time range prediction for each precursor (the retention time of palmitic and stearic acid are used to predict retention time ranges of all other fatty acids at the sum composition level, Supplementary Information, section 3.1.2; Fig. S10), those *m/z*-values with a precursor ion abundance above a user defined threshold and an associated *m/z*-error within another user-defined threshold (e.g., between 10 to 60 ppm, depending on the derivatization agent and instrument parameters), are selected within the respective retention time ranges (precursor analysis step). For each such precursor signal, targets are added to a raw target list (Fig. 1e). The raw target list is resampled, leading to a final target list, which contains one single precursor target at each defined retention time increment, with typically a total of 1000 to 3000 individual targets (*m/z* value for mass selection with associated retention time). The increments are defined such that the mass spectrometer can carry out for each increment at least one tandem MS measurement including mass selection, OzID of the mass selected precursor and analysis of the precursor and the respective OzID product ions in the time-of-flight analyzer. The resampling algorithm is designed to retain sufficient targets for each precursor that is detected, irrespective of abundance. This allows detection of low abundant fatty acids (e.g., mead acid, FA 20:3*n*-9,12,15) that are not isomers of coeluting, highly abundant fatty acids (e.g., oleic acid, FA 18:1*n*-9) in human plasma (Fig. 2). Acquisition of tandem mass-spectra enables explicit assignment of OzID product ions to precursors, thus minimizing assignment errors while the enhanced signal-to-noise ratio (S/N) reduces the limits of detection.

**Figure 2:**
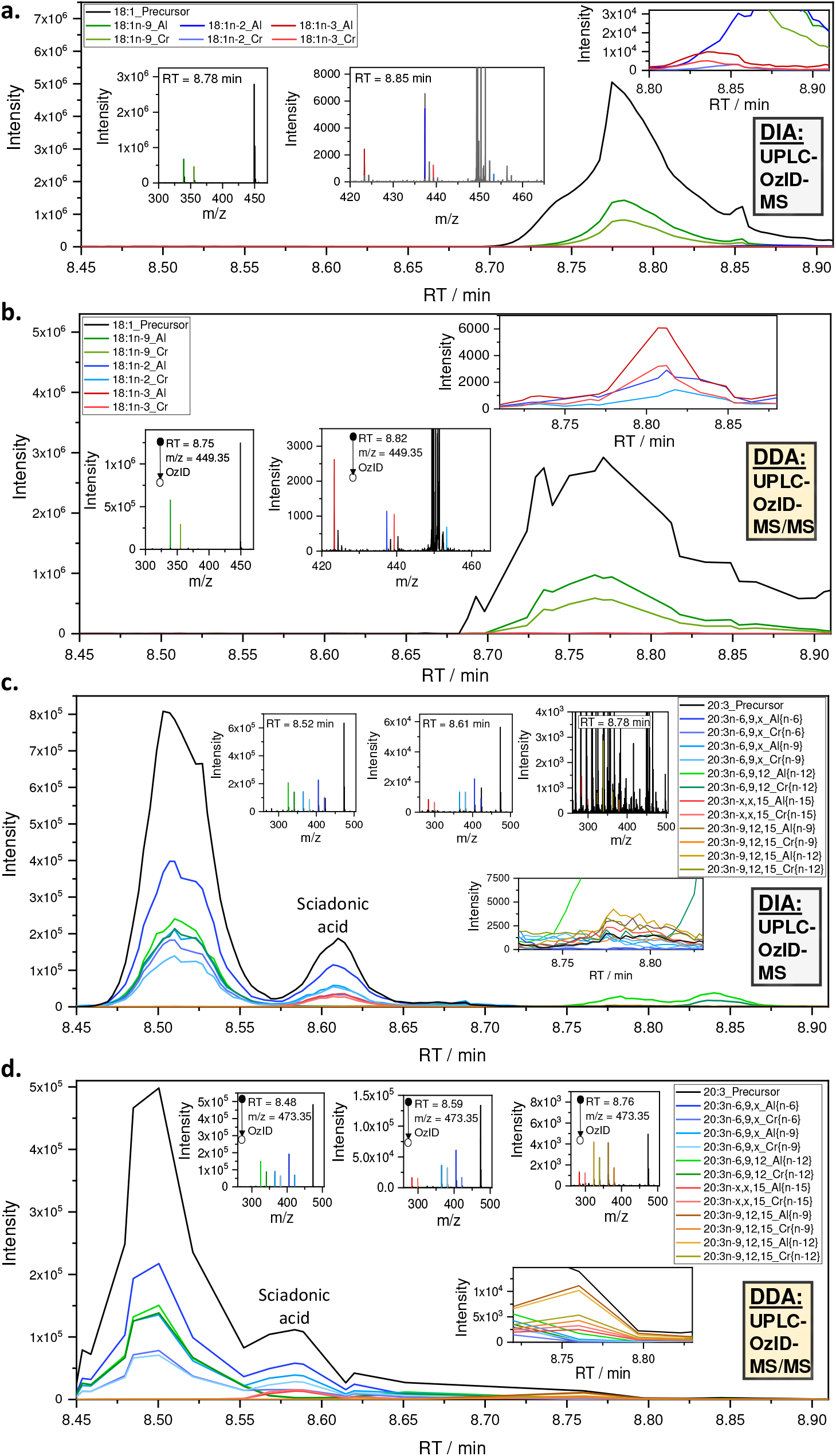
LC-MS data for selected fatty acids obtained by DIA UPLC-OzID-MS (**a**. and **c**.) and the associated DDA UPLC-OzID-MS/MS acquisition (**b**. and **d**.) from AMPP derivatized fatty acids from hydrolyzed human plasma (NIST 1950 SRM). Shown are extracted ion chromatograms and (tandem) mass spectra consistent with identification of FA 18:1*n*-9 (oleic acid), FA 18:1*n*-2, FA 18:1*n*-3, FA 20:3*n*-6,9,12 (dihomo-γ-linolenic acid), FA 20:3*n*-6,9,15 (sciadonic acid), and FA 20:3*n*-9,12,15 (mead acid). The extracted ion chromatograms of the data-dependent acquisition contain only a few points across each chromatographic peak, as multiple precursors are mass selected consecutively according to a highly segmented target list.

After acquisition, an exhaustive search for OzID product ions arising from all possible double bond positions can be conducted with subsequent automated filtering. Within this step, typically 100000 to 800000 transitions (instances of putative fatty acid assignments at associated retention time) are created for processing with Skyline Runner, which are automatically filtered to a drastically reduced number of transitions, typically totaling between 300 to 5000 transitions, depending on the parameters of the analysis. Alternatively, the search can be limited to a subset of defined fatty acid species for cases where replicate or related samples have been processed already. Multiple analysis steps within Skyline runner and custom-written python code are automated and controlled by a windows batch file. The target list generated in the previous step is used to guide the search for matching precursor and product signals at the respective retention times. To prevent loss of correctly identified species within the workflow, the output Skyline file retains duplicate transitions and potentially false positive assignments. Curation of the list of putative assignments at this stage enables removal of duplicates and false-positives based on the following criteria: i) visual inspection of extracted OzID product ion chromatograms allows deletion of duplicates (multiple transitions assigning the same chromatographic feature), ii) the signal-to-noise ratio of the OzID product ions in tandem OzID mass spectra are to be determined with a separate python program, allowing all species for which S/N<3 to be deleted from the Skyline file and those for which 3<S/N<10 to be tentatively identified (not subjected to quantification) and iii) the MS/MS view in Skyline can also be used to visually validate that all OzID product ions are derived from the respective precursor.

Relative quantification of fatty acids is performed using the DIA dataset, while the DDA dataset aids in validating this process and providing a means to correct the outcome, if required. While extracted ion chromatograms of OzID product ions are used to establish peak positions, a deconvolution of the precursor extracted ion chromatogram is used for relative isomer quantification as this minimizes biases arising from differences in ozonolysis efficiency. The deconvolution is facilitated by the automatic generation of an Excel file with prefilled parameters and graphs for fitting of Gaussians representing each isomer centered on the OzID transitions. For highly complex samples containing low-abundant coeluting isomers, several such species can only be quantified based on their OzID product ions. Thus, the relative quantity of these fatty acids represents a sophisticated best estimate with a mean accuracy of 4% of the reported value (based on the ratio of isomers present in a standard mixture of 37 fatty acids). Visualizations of relative abundances in bar charts and structural identifiers including systematic names and LipidMAPS IDs are generated by several python scripts and facilitate reporting of the findings as well as comparison with known fatty acid species (LipidMAPS Lipidomics Gateway online database).^36^

A best estimate of absolute quantities can finally be generated by Python scripts using direct infusion data (Fig. 1d). While absolute quantification is not a priority of this discovery workflow, it is advantageous to be able to access a measure of the contribution of each isomer to the total fatty acid pool. There are well recognized challenges in quantification in reversed-phase LC-MS of lipids due to changes in ionization efficiencies during gradient elution. To bypass this challenge, a parallel loop injection (direct infusion- or shotgun-ESI experiment without ozone) of 10 μL of the sample is undertaken. Isotope corrections are conducted on these data and the presence of the deuterated palmitic acid internal standard allows quantification of the fatty acid precursors at the sum composition level (i.e., the total number of carbons and degree of saturation, e.g., all FA 18:1 isomers) as illustrated in Fig. 3a. Parallel measurement of all fatty acid groups (without discrimination of isomers) by direct infusion ensures that ionization biases are reduced. These data enable relative quantitation of fatty acids at the sum composition level and reproduces the make-up of the reference mixture of fatty acids (based on reference FAs ranging from 12-24 carbons, Fig. S9). For absolute quantitation, a final python script concludes the workflow by drawing on the sum composition quantitation of replicate data against the internal standards, producing the final best estimate including standard deviation and coefficients of variation for each fatty acid.

**Figure 3:**
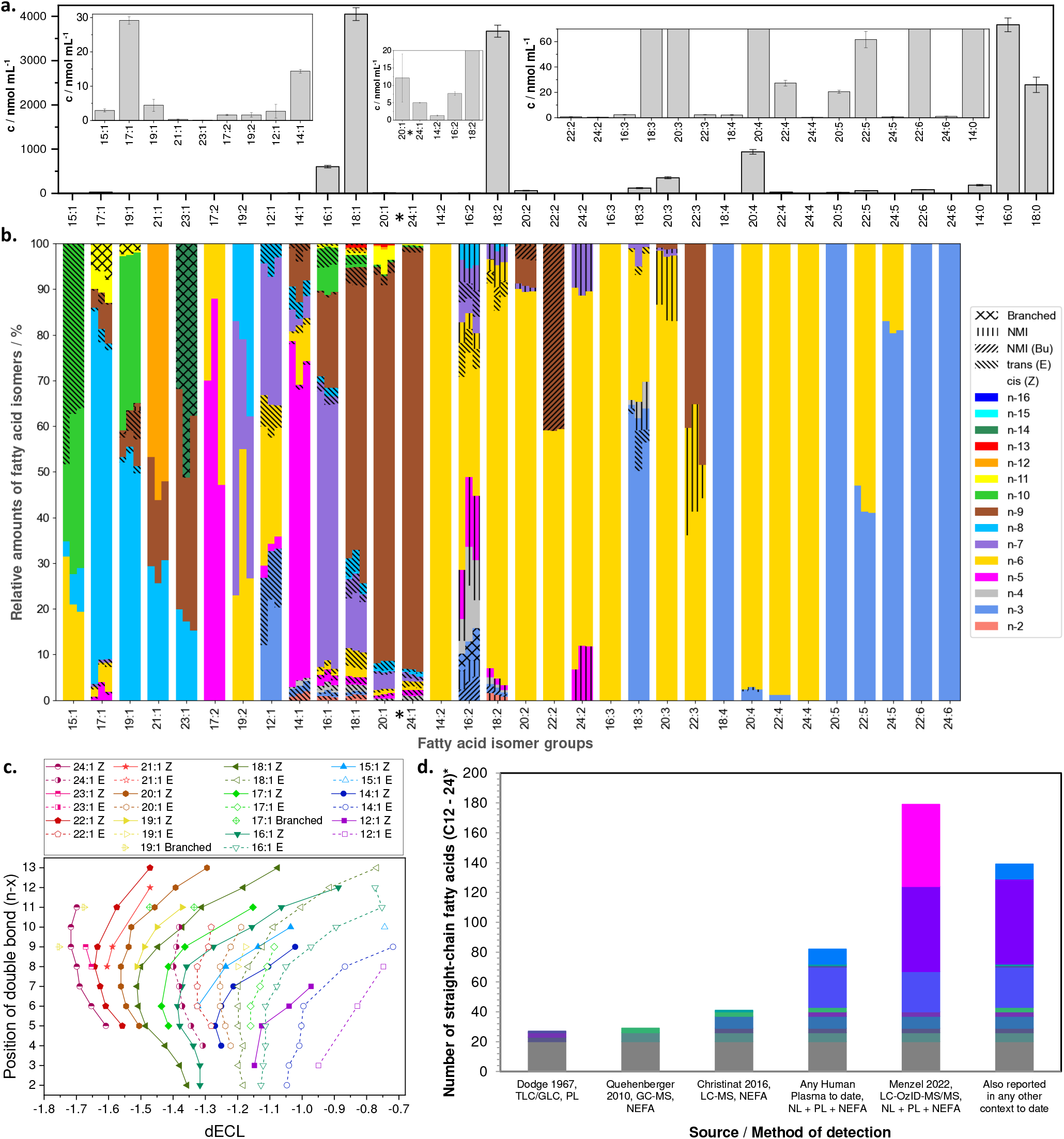
Analysis of fatty acids in human plasma (NIST 1950 standard reference material). **a**. Quantification of plasma fatty acids at the sum composition level based on direct infusion ESI-MS. Selected saturated fatty acid quantities are shown, for the full profile see the Supplementary Information, Section 3.3, Table S4. **b**. Color-coded bar chart showing the different fatty acid isomers and their relative abundance as determined by LC-OzID-MS. Each technical replicate is represented by a vertical segment of each bar. Non-methylene interrupted (NMI) fatty acids are highlighted by patterns distinguishing between butylene interrupted (Bu) fatty acids and others. **c**. Identified monounsaturated fatty acid species in the NIST 1950 human plasma and their relation between structure and retention time. Each data point represents a fatty acid isomer with double bond position as shown on the y-axis, plotted against dECL, the difference of the observed equivalent chain length (ECL) to the chain length of the associated saturated fatty acid. **d**. Comparison of the numbers of straight chain fatty acids (C12-24) that are reported in human plasma, detected by different analytical methods and summarized literature surveys. Colors highlight fatty acid species that are found in multiple contexts or only reported in one of the displayed contexts. *Docosenoic acids (FA 22:1) are excluded from the analysis due to unavoidable contamination from plastics additive erucamide (FA 22:1*n*-9).

### Benchmarking OzFAD using a pooled human plasma standard reference material

Human blood plasma is one of the most studied biological samples in biomedical research.^37^ To benchmark our discovery workflow, we used NIST 1950 human plasma as a well-studied, commercially available reference. This sample combines blood plasma of 100 American volunteers and is widely accepted as a reference material to standardize the field of lipidomics.^38,39^ After processing of three replicates with the OzFAD workflow, comparison of quantities of unsaturated fatty acids observed with certified values reported for this reference material^40^ reveals an accuracy of 63% (mean average deviation) for the OzFAD workflow (Supplementary Information, Fig. S23) versus accuracy (mean average deviation) of participating laboratories in the same interlaboratory study^41^ ranging from 11% to 62%. The range of reference values reflects the challenge of absolute quantification in lipidomics. Further enhancement of the quantification method described herein (e.g., through wider availability of fatty acid isomer standards) is beyond the scope of this study.

Fig. 2(a) and (c) show example extracted ion chromatograms from DIA LC-OzID-MS of lipids extracted, hydrolyzed and derivatized from the reference human plasma. As expected, the data readily identify abundant oleic acid (FA 18:1*n*-9) by retention time alignment (8.78 min) of the precursor ion at *m/z* 449.35 and the OzID product ions at *m/z* 339.21 and 355.20. Importantly, the selectivity and dynamic range of the OzFAD analysis reveals many more features corresponding to double bond regioisomers, for example FA 18:1*n*-2 and FA 18:1*n*-3 (where the *n*-x nomenclature indicates the position of the double bond with respect to the methyl terminus), despite being incompletely resolved by chromatography (Fig. 2a inset). These putative identifications are confirmed by the targeted DDA LC-OzID-MS/MS steps that uniquely assign the OzID product ions to the selected precursor ion for each double bond positional isomer. In addition, the greatly improved signal-to-noise ratio in the tandem mass spectrum enables reconstruction of the discrete retention time of each isomer (Fig. 2b). This is further highlighted by the identification of a fatty acid with composition FA 20:3, which coelutes with the significantly more abundant FA 18:1 isomer family. While the data-independent acquisition does not produce unambiguous chromatographic or mass spectral evidence of the double bond positions of FA 20:3 at 8.78 min (Fig. 2c, insets), the data-dependent tandem mass spectra locate multiple pairs of time-aligned OzID transitions that identify this fatty acid as FA 20:3*n*-9,12,15 (mead acid) (Fig. 2d, insets). In cases of fatty acids that are well-known, abundant and chromatographically resolved from other features, such as FA 20:3*n*-6,9,12 (dihomo-γ-linolenic acid, 8.50 min), the DIA LC-OzID-MS analysis (Fig. 2c and insets) may sufficiently support identification. In contrast, for unexpected features, such as FA 20:3*n*-6,9,15 (sciadonic acid, 8.60 min), confirmation by the characteristic OzID product ions in the DDA tandem MS acquisition is critical for confident identification.

Harnessing the full analytical power of the workflow, we identified a total of 186 fatty acids in the NIST 1950 SRM that meet the stringent criteria for identification, including the retention time range prediction and observation of the time-aligned precursor and OzID product ions in the tandem MS acquisition with a signal-to-noise ratio for the OzID product ions above 10 (Fig. 3). Combining the absolute quantification at the sum composition level (Fig. 3a) with the relative quantification of the isomers (Fig. 3b) it is possible to determine the concentration of all fatty acids identified by this workflow (Table S4). This analysis establishes the dynamic range of confidently assigned fatty acid species to five orders of magnitude, with oleic acid (FA 18:1*n*-9 *cis*) being detected at 2.5 ± 0.1 μmol mL^-1^, whereas FA 24:1n-6 *cis* is observed at 0.028 ± 0.006 nmol mL^-1^. Three replicates of the fatty acid analysis of NIST 1950 SRM pooled human plasma are displayed (Fig. 3a).

Each group of fatty acids, such as dodecenoic acids are represented as a segmented bar, showing the relative abundances of replicate 1 on the left and replicate 2 and 3 on the middle and right, respectively, of each bar. Both identification as well as relative quantification are reproducible, as any species found to be present above noise was detected in all three replicates. The visual representation highlights that variations between technical replicates are smaller in case of fatty acids with a high overall abundance, for example in the case of octadecenoic acids (FA 18:1) a mean coefficient of variation (COV) of the relative isomer abundance of 0.2 is determined, whereas variations are larger for fatty acids with low overall abundance, such as nonadecadienoic acids (FA 19:2) with a mean COV of 0.5.

Configuration of double bonds of monounsaturated fatty acids is assigned here by comparison of retention times with other fatty acids. The retention times of fixed-charge derivatized fatty acid isomers in reversed-phase chromatography are systematically affected by double bond position and configuration. The concept of equivalent chain length (ECL) or equivalent carbon number was introduced to rationalize trends in fatty acid methyl ester retention times in gas chromatography and to compare values between different chromatographic runs that may be affected by retention time shifts. We found that the difference between observed equivalent chain length and chain length of the respective saturated fatty acid (differential equivalent chain length, dECL) is a measure that allows comparison of fatty acid isomer retention behavior across varied chain lengths and double bond positions (Fig. 3c). The trends in dECL values of all monounsaturated fatty acids that we detected in human plasma enable assignment of fatty acid structures through extrapolation and interpolation of dECL values across fatty acids with varied chain lengths and double bond positions. In general, branched chain fatty acids elute earlier than straight chain fatty acids (comparing fatty acids with the same number of carbons overall), while fatty acids with *cis* double bonds (connected by solid lines in Fig. 3c) elute earlier than their *trans* counterparts (connected by dashed lines in Fig. 3c). For example, the configuration of FA 17:1*n*-11 *cis* (6Z-heptadecenoic acid) is confirmed by comparing its dECL value of -1.37 to the dECL values of other detected heptadecenoic acids as well as to hexadecenoic and octadecenoic acids considering their double bond position. Thereby, the other two instances of FA 17:1*n*-11 are identified as branched unsaturated fatty acids, although no assignment of the number and position of the branch points can be made by LC-OzID-MS. The equivalent carbon number concept was recently also employed to annotate lipids in reversed-phase liquid chromatography across multiple lipid classes.^42^ We found that *trans* fatty acids make up 6.5% of the total fatty acid content of the reference plasma.

We surveyed the literature to assess whether each of the fatty acids we identified in the standard reference material had been identified previously in human plasma, was reported in any other context, or represents a newly discovered fatty acid. The outcomes of this survey are shown in detail in Table S4 (Supplementary Information, section 3.3) and numbers of detected fatty acids are summarised in Fig. 3d. The bar chart in Fig. 3d displays the numbers of straight-chain fatty acids (C12-24) that were identified in three selected publications^37,43,44^ compared to the number identified in all publications that were part of our literature survey as well as the ones that were also identified in any biological context to the best of our knowledge. The color-coded areas represent fatty acids that were commonly found in the varied contexts. In addition, we identified five branched chain fatty acids with the limitation of not being able to determine the position of the branch point(s). Further, thirteen docosenoic acids were observed, which are excluded from the comparison due to the presence of erucamide and isomers of this erucic acid derivative in the associated Process Blank. Finally, an additional 33 bisunsaturated fatty acids are only tentatively identified due to incomplete chromatographic separation. Future work on longer chromatographic gradient methods for LC-OzID-MS is expected to increase the number of confidently assigned fatty acids. Given that some prior investigations of human plasma fatty acids had focused on non-esterified fatty acids (NEFA), we carried out a derivatization using a non-hydrolyzed lipid extract from the same pooled human plasma reference material (Fig. S24 and Table S5). This analysis led to a similar profile of fatty acid species, with only one additional isomer being detected in NEFA, that was not found in the total hydrolyzed lipid extract. However, several low-abundant species that were identified in the hydrolyzed lipid extract were not detected above the signal-to-noise threshold in the NEFA analysis. Thus, analysis of hydrolyzed lipid extracts with the OzFAD workflow provides an excellent foundation to help establish overall lipid diversity (Supplementary Information, section 3.3 and 3.4).

Considering the analysis of total fatty acids, our workflow led to the discovery of 55 novel fatty acid isomers (including FA 18:1*n*-2 *cis* and 31 novel *trans* fatty acids) that, to the best of our knowledge, have previously not been reported, while an additional 52 fatty acids have not been reported in human plasma, but have instead been identified in other biological sources (including FA 18:1*n*-3 *cis*). Among the identified fatty acids, twelve are non-methylene interrupted fatty acids including sciadonic acid (FA 20:3*n*-6,9,15 at 47 ± 8 nmol mL^-1^) and keteleeronic acid (FA 20:2*n*-9,15 at 2.4 ± 0.4 nmol mL^-1^). Polyunsaturated fatty acids that exhibit a pattern of unsaturation other than the canonical methylene-interrupted or conjugated sequences of double bonds have not previously been reported in human plasma but have been observed in other biological systems including pine nuts,^45^ sea urchins,^14^ mangos,^46^ marine food webs,^47^ dairy^48^ as well as human breast milk.^49^ Prior identification of non-methylene interrupted fatty acids in foodstuffs may point to these being present in reference plasma as a result of dietary intake. However, the co-detection of unsaturated fatty acids that would serve as potential intermediates in the biosynthesis of these species suggests the intriguing possibility that human lipid metabolism may also be capable of generating them. Therefore, to investigate the diversity of human lipid metabolism, we demonstrate the capability of the OzFAD workflow in lipids derived from human sebaceous secretions (vernix caseosa) and human-derived cell lines.

### OzFAD uncovers non-canonical fatty acids in vernix caseosa

Vernix caseosa, the white waxy layer that newborns are covered in at birth, is known for complexity and perversity of fatty acid structures with up to 167 chromatographic features previously attributed to distinct fatty acids but without complete structural assignment.^50,51^ The vernix caseosa fatty acid profile is characterized by very long chain fatty acids, extensive chain branching and unusual positions of unsaturation including high abundance of sapienic acid (FA 16:1*n*-10*cis*).^52,53^ Analysis of vernix caseosa was conducted according to the workflow described herein with the 4-iodo-AMPP derivatization agent utilized in place of AMPP (Fig. S27 and Table S6).^52^ *De novo* assignment of sites of unsaturation was conducted as previously described, with fatty acids assigned as branched or straight chains based on an analysis of retention time indices (Supplementary Information, Fig. S26). While assignment of the explicit site of methyl chain branching (e.g., *iso* or *anteiso*) is beyond the scope of this method, unsaturated fatty acids with straight chain and branched chain variants were well-resolved by chromatography allowing ready classification. For example, a branched fatty acid with 17 carbon atoms in total, a methyl branch in the penultimate position (*anteiso*) and a double bond in the Δ7 position (14Me-16:1*n*-9*cis*; (7Z)-14Me-hexadecenoic acid) would be classified by OzFAD as 17:1*n*-10 and due to the retention time alignment and differential equivalent chain length (dECL) analysis it would be assigned as branched. As both chain branching and double bond positions influence retention times, the unequivocal assignment of double bond positions will aid in the full structural identification of all fatty acids in complex samples such as vernix caseosa by future extensions of this workflow incorporating photodissociation of the photolabile 4-iodo-AMPP derivatization agent.^52^

Similar to the scale of discoveries in human plasma, our workflow revealed an unexpected diversity of double bond positions and patterns, including 18 straight chain saturated, 116 straight chain monounsaturated, 32 straight chain polyunsaturated, 30 branched saturated and 42 branched monounsaturated fatty acid species out of a total number of 238 fatty acids (Fig. 4 and Supplementary Information, section 3.5). Further, trends in relative abundances of monounsaturated fatty acids of increasing chain lengths (16-30 or 17-29 carbons, respectively) were observed, with *n*-5 to *n*-7 monounsaturates making up an increasing percentage of the respective isomer groups, whereas the proportion of *n*-9 to *n*-12 monounsaturates decreases accordingly. Such observations may be indicative of differential elongation rates depending on double bond positions (Supplementary Information, Fig. S27). In addition to monounsaturates, the sensitivity of the OzFAD approach enabled identification of several novel non-methylene interrupted polyunsaturated fatty acids, which could represent intermediates in a previously unknown metabolic pathway (Fig. 4 and Fig. S27). For example, Fig. 4 presents mass spectral evidence for the presence of FA 16:1*n*-3*cis* that could plausibly undergo Δ6 desaturation (e.g., *via* FADS2) to yield FA 16:2*n*-3,10. Subsequent elongation could yield FA 18:2*n*-3,10 followed by Δ5 desaturation (e.g., *via* FADS1) to give FA 18:3*n*-3,10,13. While mechanistic studies are required to demonstrate this pathway unequivocally, the unambiguous detection of all intermediates in this pathway provides a strong foundation for the hypothesis that human (or, alternatively, human microbiome^54^) lipid metabolism is capable of the biosynthesis of non-methylene interrupted fatty acids. Overall, the OzFAD workflow critically aids the discovery and investigation of unexpected double bond patterns that occur in low abundant fatty acids and coelute with known isomeric structures.

**Figure 4:**
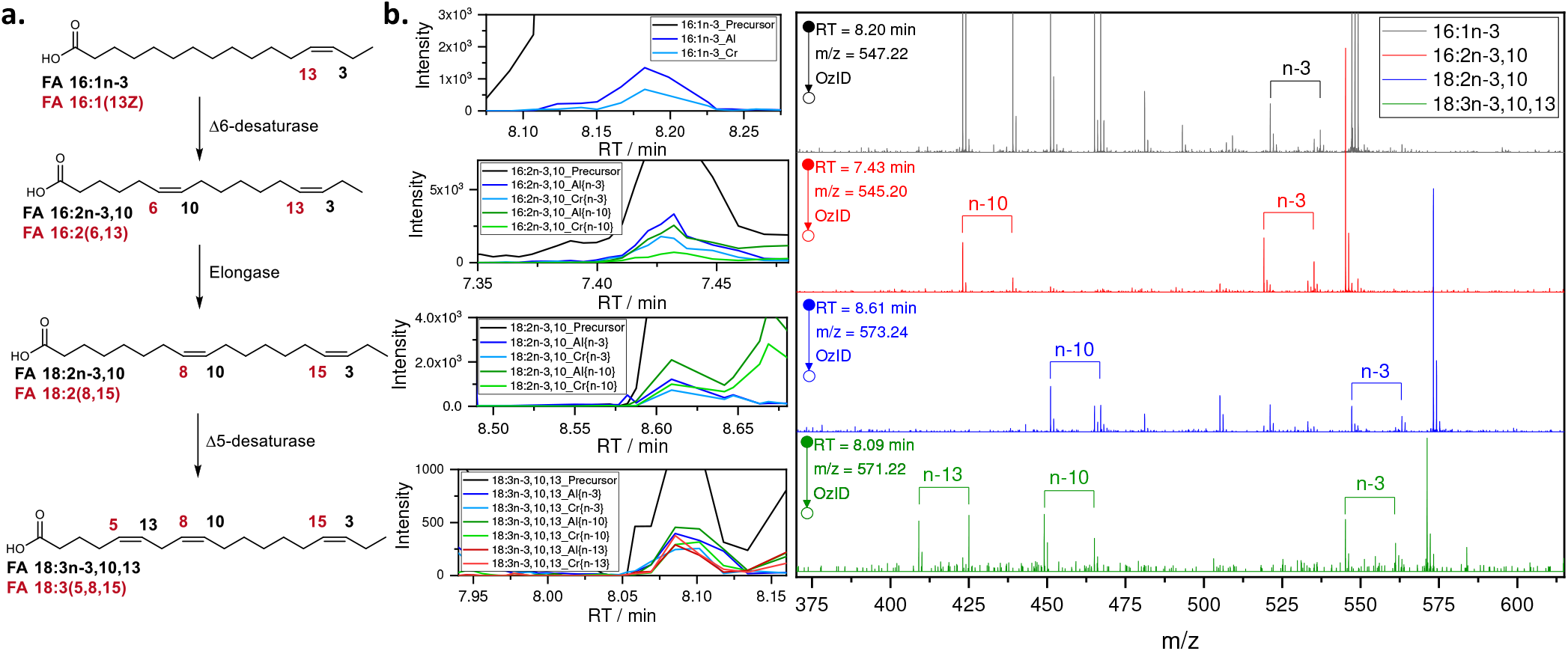
**a**. A proposed biosynthetic pathway rationalizing the discovery of non-methylene interrupted fatty acids in vernix caseosa, such as the never previously reported fatty acids FA 16:2*n*-3,10 (6,13-hexadecadienoic acid); FA 18:2*n*-3,10 (8,15-octadecadienoic acid) and FA 18:3*n*-3,10,13 (5,8,15-octadecatrienoic acid). The double bond configuration is tentatively assigned as *cis* in each case. **b**. Associated extracted ion chromatograms and OzID MS/MS spectra of the 4-I-AMPP derivatized fatty acids. For the complete fatty acid profile, refer to the Supplementary Information, section 3.5, Fig. S27 and Table S6.

### OzFAD for quantifying changes in lipid metabolism of cancer cell lines

The role of lipids and fatty acids in cancer metabolism is gaining increased interest.^55-58^ Here, we applied the OzFAD pipeline to characterize the total fatty acid profiles of three cancer cell lines (i.e., MCF7, LNCaP and LNCaP_SCD1*i*; SCD-1 inhibition: A939572) and investigate the effect of enzyme expression and inhibition of stearyl-CoA desaturase-1 (SCD-1 / Δ9-desaturase) (Supplementary Information, section 3.7). The MCF7 cell line originates from human breast cancer and has high expression of SCD-1, whereas LNCaP originates from a metastatic human prostate cancer that has previously been shown to exhibit non-canonical pathways of desaturation and elongation. Findings from the application of the OzFAD protocol to these cell lines are summarized in Fig. 5.

**Figure 5:**
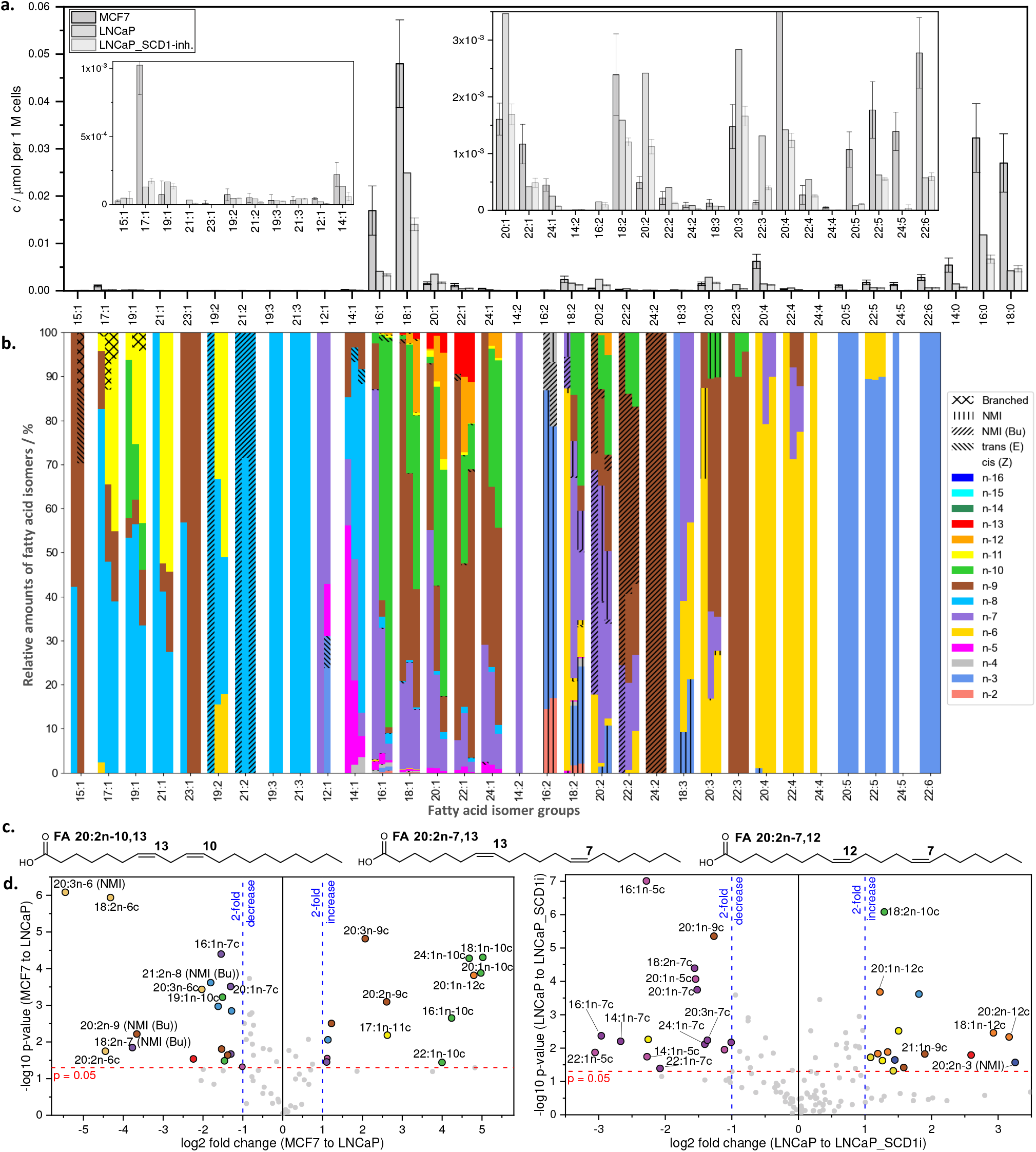
Fatty acid profiles of three cancer cell lines. **a**. Estimated fatty acid quantities by direct infusion ESI-MS. The error bars indicate the standard deviation of three biological replicates. Due to interference of ion suppressing contaminants in two replicates of LNCaP, only one is shown here, instead of the mean of three replicates. **b**. Relative quantification of fatty acid isomers in the three cell lines by LC-OzID-MS and LC-OzID-MS/MS. For each isomer group, a segmented bar is shown, where the segments on the left represent fatty acids in MCF7 cancer cell line extracts, the segments in the middle of each bar represent fatty acids in LNCaP cancer cell line extracts and segments on the right, respectively, represent LNCaP_SCD-1*i* cancer cell line extracts. Shown are mean values of relative abundances of three biological replicates of each cell line. For individual values and standard deviations, refer to the Supplementary Information. **c**. Example docosadienoic acids in either MCF7 or the LNCaP cell lines. **d**. Volcano plots visualizing the fold changes (and associated p-values) in relative fatty acid isomer abundances as compared between MCF7 and LNCaP (left volcano plot) and between LNCaP and LNCaP_SCD1*i* (volcano plot on the right). Selected isomers are labelled to show examples of large changes (exceeding two-fold changes, blue dashed lines) in relative isomer abundances within the respective fatty acid isomer groups with an associated p-value above 0.05 (red dashed line).

While the direct infusion data, shown in Fig. 5a, indicates differences in the abundance of fatty acid isomer groups, only the isomer-resolved LC-OzID-MS analysis (Fig. 5b) reveals significant underlying changes in the presence and abundance of distinct molecular species. Further, a comparison of the number of octadecenoic acids in LNCaP cancer cell lines identified in our previous work based on high-resolution direct-infusion OzID-MS to the LC-based OzFAD workflow indicates an increase from 7 species^23^ to 17 fatty acids (Table S9). The effect of lowered and inhibited Δ9-desaturation corresponds to the relative amounts of several species, as demonstrated by the volcano plots (Fig. 5c). Inhibition of SCD-1 activity shows the metabolic shift from *n*-5, *n*-7 and *n*-9 monounsaturates (e.g., FA 14:1*n*-5, FA 16:1*n*-7) to the *n*-10 and *n*-12 counterparts (FADS2 Δ6-desaturation products) and subsequent desaturation and elongation products (from sapienic acid FA 16:1*n*-10 to FA 22:3*n*-10,13,16), in line with previous findings.^23,59^ Interestingly, the pathway to FA 22:4*n*-7,10,13,16 appears significant in LNCaP cells, but not in MCF7 cells.

Some metabolic shifts may be masked by conventional analysis of fatty acids, whereas the wide dynamic range of the OzFAD workflow reveals even trace fatty acids in the media for cell culture. Here we analyzed fetal bovine serum (Fig. S28 and Table S7), to establish the fatty acid profile of the culture media. While bovine and human fatty acid profiles are known to be different,^60^ OzFAD analysis again uncovers an unprecedented diversity. Comparison of fatty acid profiles of the cells and the culture media (fetal bovine serum) indicates that, for example, the *n*-7,13 butylene interrupted pattern observed in FA 18:2, 20:2 and 22:2 is present in MCF7 cell extracts, but not in the media or in the LNCaP cells, revealing a distinct metabolic pathway. We make the hypothesis that a biosynthetic route *via* Δ5-desaturation (FADS1) of FA 18:1*n*-7 yields FA 18:2*n*-7,13 and likewise FA 20:1*n*-9 leads to FA 20:2*n*-9,15 (Supplementary Information, section 3.7, Fig. S29). Elongation of these leads to chain lengths up to 22 and 24 carbons, respectively.

The same mechanism for the formation of butylene-interrupted fatty acids could apply to odd chain fatty acids FA 19:1*n*-8,14 and FA 21:1*n*-8,14. All non-methylene interrupted fatty acids that are derived from Δ5-desaturation, including sciadonic acid (FA 20:3*n*-6,9,15), are present in significantly higher ratios in MCF7 cells compared to LNCaP cells. On the other hand, the non-methylene interrupted fatty acid FA 20:2*n*-7,12 is only found in LNCaP cells, but not in the culture media or in MCF7 cells, indicating the possibility of a novel Δ6 desaturation pathway from FA 20:1*n*-7. Detection of novel fatty acid isomers is critical to reveal substrate specificity of desaturases and elongases and to complete human lipid profiles including their biological drivers.^61-64^

## DISCUSSION

We have developed an end-to-end workflow for the *de novo* identification of fatty acids from complex biological samples. Application of this pipeline to human-derived lipids from blood plasma, cancer cells, and sebaceous waxes has significantly extended the number of fatty acid identifications in these well-studied systems. Compared to other strategies to elucidate double bonds, the OzFAD workflow features two distinct advantages. Firstly, the *de novo* data analysis relies on no assumptions of which fatty acids may be present in the sample, allowing the detection of unexpected sites of unsaturation, making the workflow a true discovery tool. The analysis proceeds *via* a stepwise process involving an automated, exhaustive isomer search and filtering, augmented by intermittent user input for quality control. Secondly, computationally-generated targets for targeted acquisition allows mass-selection of all precursors that are present without direct user intervention. Carrying out OzID on the mass-selected precursor ions yields confident assignments of all unsaturated fatty acids across five orders of magnitude with signal-to-noise values for the ozonolysis products above ten. This analysis, based upon the specific and predictable OzID product ion generation, enables tracking a multitude of coeluting fatty acid isomers even across vastly different concentrations, alleviating the need for complete chromatographic separation or for reference mass spectra. We aimed to combine the best of automation and human intelligence to streamline the discovery process, avoiding false positive or negative identification.

The application of retention time analysis on large pools of fatty acid isomers has enabled augmentation of the fatty acid structural assignment beyond double bond location to include double bond configuration and the identification of methyl branching. Currently, the method cannot explicitly assign sites of chain branching but this can be overcome in the future by the introduction of parallel analysis of samples using radical-directed dissociation^11^ or similar methods. Importantly, the effective application of the I-AMPP derivatization as well as the archetype AMPP derivative indicates that these methods can be run in parallel on the same derivatized extracts. While the workflow unmasks structural fatty acid diversity to a level not seen before, future work will focus on improving quantitation accuracy across the vast number of fatty acids that can now be identified. Central to these advances will be improved availability of reference standards, both standards for many of the novel fatty acids that have been presented here for the first time as well as isotopically labelled standards for each isomer group.

Application of the OzFAD tool has already underpinned the discovery of 107 fatty acids, not previously reported in human plasma. Of these, 55 fatty acids are, to the best of our knowledge, discovered here for the first time. All but one species that were identified in non-esterified fatty acids in human plasma were also found in the total fatty acid fraction (hydrolyzed lipids), indicating that the analysis of hydrolyzed lipids by this method can inform future in-depth studies of intact lipids by LC-OzID-MS/MS. Thereby, we anticipate an expansion of lipidome coverage on a similar scale compared to the discoveries shown herein. Many of the discoveries revealed here correspond to *trans* monounsaturates, whereas some non-methylene interrupted species were detected. Interestingly, sciadonic acid is found at significant concentrations in the human plasma standard reference material NIST 1950. The discovery of such non-canonical patterns of unsaturation in vernix caseosa and human cell lines indicates that hitherto undescribed lipid metabolism may be responsible for the synthesis or modification of lipids leading to production of such species. The depth of the analysis shown here sets new standards for the identification of fatty acid double bond isomers. Our findings and future work are expected to have far-reaching consequences for lipidomics and metabolomics studies involving cell culture, healthy and diseased human tissues, cosmetics, supplements, the human diet as well as plant, microorganism, and animal metabolism. The OzFAD workflow is a tool to tackle the challenge of lipidome coverage at a deeper level than previously achieved, adding to the toolkit of methods to study highly complex interactions, such as those between the human microbiome(s) and the human metabolism.

## METHODS

### Nomenclature

Fatty acids are described within this work using either systematic names (sciadonic acid is 5Z,11Z,14Z-eicosatrienoic acid; indicating the double bond positions from the carboxyl end of the fatty acid) or accepted shorthand notations using the *n*-nomenclature (sciadonic acid is either FA 20:3*n*-6,9,15 or briefly 20:3*n*-6,9,15; indicating double bond positions from the methyl terminus). Configuration of carbon-carbon double bonds is denoted using either E/Z, *trans/cis* or briefly t/c, such that oleic acid is either systematically 9Z-octadecenoic acid, or FA 18:1*n*-9*cis* (or briefly 18:1n-9c).

### Cell culture and lipid extraction

Human prostate cancer cell line LNCaP (clone FGC) was obtained from the American Type Culture Collection and authenticated by short-tandem repeat profiling at Cell Bank Australia (NSW, Australia) in July 2020. Cells were cultured in RPMI-1640 medium (Life Technologies) containing 10% (v/v) fetal bovine serum (FBS) and 2 mmol L^-1^ L-glutamine (Life Technologies) in 5% CO_2_ in a humidified atmosphere at 37 °C and subjected to regular mycoplasma testing. For SCD-1*i* treatment, cells were seeded in triplicate at 4.5 × 10^5^ cells/well in 6-well plates overnight, before being treated with 1 μM A939572 (Tocris) using dimethylsulfoxide as a vehicle and cultured for 72 h, then collected for lipid extraction. Human breast cancer MCF7 cells were grown in Dulbecco’s modified eagle medium (DMEM) medium complemented with 10% (v/v) fetal bovine serum (FBS; heat inactivated for 30 min at 56 °C) and 1% (v/v) penicillin/streptomycin in an incubator set to 37 °C with 5% CO2. The cell culture was profiled *via* short tandem repeat profiling and regularly tested for mycoplasma.

Lipids were extracted following a protocol by Matyash *et al*.^*35*^ and modified by Young, *et al*.^*23*^ Cells were trypsinized and counted with an automated TC-20 cell counter (BioRad) and 1 × 10^6^ (in the case of LNCaP and LNCaP_SCD1i: 5 × 10^6^ and subsequently adjusted solvent volumes) cells were pelleted in microfuge tubes (in triplicate). The cell pellets were washed twice with Dulbecco’s phosphate-buffered saline (DPBS) and stored at -80 °C. Lipids were extracted in 2 mL glass vials and a stock solvent mixture containing 370 μL methyl tert-butyl ether (MTBE) with 0.01% butylated hydroxytoluene (BHT). Cell pellets were dispersed in 110 μL methanol and 390 μL solvent mixture and vortex mixed for 20 sec with subsequent continuous agitation for 1 h. After addition of 100 μL ammonium acetate (150 mM) and vortex mixing for 20 sec, the phases were separated by centrifugation for 5 min at 2000 g. The organic phase was collected and stored at -80 °C.

### Lipid extraction from NIST1950 SRM (pooled human plasma)

After thawing and sonication for 5 mins, the NIST Human Plasma was found to be a mixture of solids dispersed in the liquid phase. To reliably assess the total fatty acid content of the human plasma sample, 800 μL of the total mixture were collected and divided equally into four vials and lipids were extracted as per the procedure based on Matyash et al.^35^ The lipid extract was then pooled and equally divided over four individual vials for further treatment, hydrolysis and subsequent derivatization. Three aliquots were taken from the vials for further analysis. Thus, the standard deviation of each of the three technical replicates of the NIST Human Plasma incorporates the statistical experimental error of hydrolysis, derivatization, instrumental analysis, and data analysis.

### Hydrolysis, fatty acid extraction and fixed-charge derivatization

The sensitivity of fatty acid detection by mass spectrometric methods increases significantly by derivatization with a fixed positive charge, such as 1-(4-(aminomethyl)phenyl)pyridinium (AMPP).^65,66^ Thus, either AMPP or the iodo-substituted derivatization agent 4-I-AMPP^52^ were used in this study to investigate the total fatty acid content of the samples.

To the vial containing the residue of the lipid extraction 0.2 mL methanol and 0.2 mL aqueous tetrabutylammonium hydroxide solution (40% w/w) were added, and the mixture was heated at 75 °C for two hours. Subsequently, 1.5 mL pentane, 1.5 mL water and 0.3 mL concentrated hydrochloric acid were added. The mixture was vortex stirred for 1 min. After allowing the phases to separate, the organic layer (top layer) was collected. 1.5 mL pentane were added to the residual aqueous phase and extraction was repeated. The organic phases were combined and 3 μL *N,N*-diisopropylethylamine, and 0.15 mL acetonitrile/dimethylformamide 4:1 (v/v) was added. The biphasic mixture was vortex stirred for 1 min, before the top layer was evaporated by a stream of nitrogen. To the vial containing the residual liquid from the hydrolysis procedure 10 μL of a freshly prepared 1 M aqueous solution of *N-*ethyl*-N’*-(dimethylaminopropyl)carbodiimide hydrochloride was added, before 40 μL of a 30 mM solution of 1-hydroxybenzotriazole nonahydrate in acetonitrile was added. The solution was vortex stirred for one min. 0.1 mL of a 20 mM solution of AMP^+^ (Cayman Chemical) in acetonitrile/dimethylformamide 4:1 (v/v) was added, and the mixture was vortex stirred for 1 min and heated to 65 °C for 30 mins. After allowing the solution to cool to ambient temperature, 1 mL methanol was added. The solution was diluted six-fold and all volatiles were removed by evaporation under a stream of nitrogen. The residue was dissolved in 1 mL methanol and the sample was queued for measurement.

### Instrumentation and details of acquisitions

Analysis was performed using a Waters Acquity (UPLC i-Class; CSH, C18 reversed phase column, 2.1 × 100 mm, Particle size 1.7 μm) liquid chromatography system coupled with a Waters SYNAPT G2S*i* (Z-Spray, T-Wave Ion Mobility; TOF) mass spectrometer, previously modified to allow for ozonolysis reactions to be undertaken in the traveling-wave ion-mobility cell of the instrument.^21^ Samples were prepared in 1.5 mL LC-MS vials with septum and kept in the autosampler of the LC system at 10 °C immediately prior to analysis. The column was kept at 60 °C during analysis. Liquid chromatography was performed with a linear gradient at a flow rate of 0.4 mL min^-1^. Mobile phase A is water with 0.1 % formic acid; mobile phase B is acetonitrile with 0.1% formic acid. The injection volume varies depending on the sample from 1 μL to 10 μL. Initially, the mobile phase was kept for 0.5 min at 20% B (80% A), then linearly increased to 90% B (16 min) and subsequently linearly increased to 100% B (17.35 min), held at 100% B until 20.35 min and finally reduced to 20% B until 21 min. The mass spectrometer was operated in positive mode, sensitivity mode, MS-mode and IMS-mode. The source temperature was set to 120 °C and the desolvation temperature to 550 °C. A capillary voltage of +2.5 kV was applied, with a sampling cone of 40 V and source offset set to 40 V. Cone gas was set to 100 L h^-1^, desolvation gas 900 L h^-1^ and the nebulizer was set to 6.5 bar. Gas controls were set to: trap 2.0 mL min^-1^, He-cell 180 mL min^-1^ and IMS 10 mL min^-1^. For ozonolysis, the IMS travelling wave velocity and height were 650 ms^-1^ and 28 V, while the transfer travelling wave velocity and height were 1000 ms^-1^ and 2 V, respectively. The ion mobility cell is filled with a mixture of ozone, oxygen and nitrogen, instead of just nitrogen. Oxygen (10 psi) is converted to ozone using a high-concentration ozone generator (Ozone solutions, TG-40), generating 200 - 250 gm^-3^ of ozone in oxygen at a flow of 400 mL min^-1^. A portion of the ozone in oxygen mixture is then introduced into the IMS gas flow using a needle valve, such that the pressure in the IMS region is ∼3 mbar. Excess ozone is catalytically destroyed using an unheated ozone destruct catalyst before being exhausted from the laboratory. Both ambient and in-situ produced ozone concentrations are monitored each by an ozone monitor (106-L and 106-H, respectively; 2B Technologies). A solution of 200 pg μL^-1^ leucine enkephalin in a mixture of 50% acetonitrile and 50% water with 0.1% formic acid is used as a Lockmass solution. Lockmass correction is performed upon loading of the raw LC-MS dataset into Skyline. Data is acquired at a scan time of 0.1 secs. For acquisition of direct infusion data, the column was removed from the chromatography system and each sample was injected by the autosampler into a 50:50 mixture of acetonitrile/water with a 0.1% formic acid content (loop injection) at a flow rate of 0.4 mL min^-1^. Data is acquired for 2 min. After each injection, a needle wash was performed.

### General data analysis

Data were processed and visualized with custom-written python source code, Skyline, MassLynx, Microsoft Excel and OriginPro 9.1G. For the automated data analysis using the workflow presented here, template files and settings files provided as Supplementary Materials are required. An installation of Skyline MS (Skyline or Skyline-Daily, 20.2; 64-bit), Skyline Runner, as well as a current version of python (e.g., python 3.9.2; 64-bit; IDE: e.g., Visual Studio Code), including pandas, openpyxl, scipy, numpy, matplotlib, brainpy, requests, bs4, subprocess, statistics, csv and datetime on a Windows PC were used herein. For both the analysis of direct infusion ESI-MS and LC-OzID-MS(/MS) data the same four-letter-codes are used to describe the derivatization agent (charged head group). An overview of available abbreviations and derivatization agents is shown in Fig. S1 (Supplementary Information, section 2). The workflow can be applied to other derivatization agents without modification. In this case, a new four-letter-code and the sum formula of the associated head group needs to be entered. An overview of the stages of the analysis and acquisition steps as well as a detailed step-by-step protocol is shown in Fig. S2 and Supplementary Information, section 2.

### Data analysis protocol for direct infusion by loop injection

The simultaneous detection of a mixture of positive charge-tagged fatty acids prevents ionization biases during electrospray ionization. The measurement, carried out without fragmentation or ozone-induced dissociation (OzID), reveals ratios of fatty acid isomer groups in a commercial standard mixture (Supplementary Information, section 3.1.1 and Fig. S9). We introduce a python script for the automated analysis of the direct infusion MS data, which includes isotope corrections to account for varied numbers of ^13^C and other isotopes, as well as isobaric overlap of unsaturated fatty acids that differ by 2 Da in m/z, e.g., DPA (22:5) and DHA (22:6). The script performs a baseline subtraction, corrects *m/z* values and allows for subtraction of a process blank. Prior to running the script to quantify fatty acids, the acquired data is averaged over the full width at half maximum of the eluted peak of the loop injection. The resulting mass spectrum is copied into the respective input excel file. This is carried out analogously for the respective Process Blank. Upon starting the python script, the four-letter-code of the derivatization agent is requested, before values within the defined *m/z* range of the mass spectrum is read from the file into lists. First, the baseline is numerically calculated and subtracted from the spectrum. Any value that is negative after baseline correction is set to zero. Second, an m/z correction is performed: The peaks associated to the derivatized palmitic acid (16:0) and stearic acid (18:0) are located and the average deviation of the *m/z* value of their peak maxima from the theoretical *m/z* is calculated. The *m/z* values of the entire spectrum are corrected by the resulting shift. After isotopic corrections, the output excel file is written containing both original and processed spectra and relative quantities, from which absolute quantities are calculated using the internal standard.

### Data analysis protocol for data independent acquisition; UPLC-OzID-MS

The analysis of each LC-OzID-MS and LC-OzID-MS/MS dataset is controlled by windows batch files, which invoke instances of Skyline runner and custom python source code. The automated and user interactive steps in this procedure are described in detail in the Supplementary Information, Fig. S2, section 2.2 and 2.4. Isomeric fixed-charge derivatized fatty acids elute at similar retention times in reversed-phase liquid chromatography, allowing predicted retention time ranges based on known fatty acid standards to be used as a criterion for identification. The first step in the automated data analysis workflow establishes such retention time ranges employing the observed retention time of palmitic and stearic acid. These saturated fatty acids are present in practically all biological samples, as they also constitute the main components of any process blank sample. The prediction is empirically based on observed retention times of analytical standards, but fatty acid structures are highly predictably correlated to their retention times, allowing extrapolation from observations using analytical standards to all related structures.

### Data analysis protocol for data dependent acquisition; UPLC-OzID-MS/MS

The analysis steps that follow the precursor analysis step (performed on the DIA dataset) are highly similar for both analysis of DIA and DDA datasets. The user is prompted to select one of three options. I) Library based search, which triggers a search for double bond positions that are specified in an excel file; II) d*e novo* exhaustive search, which triggers a search for any double bond position that is chemically feasible, including conjugated fatty acids, but excluding allylic double bonds (i.e., chemically unstable allenes) or III) streamlined analysis, which invokes a *de novo* exhaustive search for all fatty acids up to and including three double bonds, but carries out a library based search for all fatty acids with more than three double bonds. The library-based modes allow faster analysis either when the identity of fatty acid isomers in the sample is known, or when the dataset is to be probed for a defined set of fatty acid isomers. On the contrary, the analysis of highly complex fatty acid profiles with many unknown species is easier *via* the slower but more thorough *de novo* exhaustive search. Criteria for filtering the data are defined by the user through the Skyline template file (instrument mass resolving power) and through input requested by the workflow, such as intensity thresholds. Further criteria for retaining transitions are coelution of OzID product ions and precursor in case of DIA analysis. Only relying on data independent acquisition can lead to false positive and false negative identifications. Especially in the case of complex biological samples, fatty acids with different m/z values may coelute and produce OzID product ions that can theoretically arise from either precursor. Secondly, the extracted ion chromatogram of a potential OzID product ion in case of a low abundant species may only show an elevated baseline, instead of a clear chromatographic peak. These problems are overcome by the tandem MS acquisition, as mass selection of each precursor ensures that associated OzID product ions arise exclusively from the former and the signal-to-noise (S/N) ratio within the tandem mass spectrum is significantly improved over the data independent acquisition.

We further introduce a separate algorithm (python script) for the calculation of S/N ratios from mass spectral data extracted from either Skyline or other Software as tables (Fig. S3 and S4). This algorithm can combine multiple individual spectra or use one and determines the number and average of OzID product peak intensities as well as the number and average peak intensity of noise peaks. Every peak m/z that is not within close proximity (defined m/z tolerance) of the precursor isotopic pattern or any m/z value that could arise from a possible OzID product is considered an eligible noise peak. The algorithm reads an excel input file that can contain hundreds of MS spectral data exported from Skyline (DDA acquisition), detects, which spectra to combine and which to compute separately and writes the S/N values for each species in an output excel file. Each eligible noise peak and each OzID product peak is numerically integrated, and the S/N value is calculated from the integrals and numbers of peaks according to equation 1. Signal is calculated as the sum of the numerical integrals of all OzID product peaks of the fatty acid in question, divided by the number of the OzID product peaks (two in case of a monounsaturated fatty acid, twelve in case of a hexaunsaturated acid). Noise is calculated as the sum of the numerical integrals of all eligible noise peaks divided by the number of the eligible noise peaks. Note that the precursor peak integral is not contained in this calculation as its intensity could be derived from a different regioisomer and only the OzID product ions in the data dependent acquisition are directly diagnostic of the specific double bond isomer.

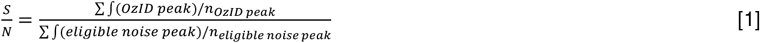

Relative quantification of isomeric fatty acids would be trivial if each isomer were completely separated from each other. Yet, reversed-phase UPLC does not fully resolve some isomeric species, which is often also true for gas-chromatographic methods. The conversion of double bonds to the characteristic product ions in *in-situ* OzID depends on the position of the double bond.^20^ Therefore, the OzFAD workflow automatically creates an excel file for the deconvolution of the precursor chromatogram of each fatty acid group (Fig. S6). The deconvolution parameters can easily be adjusted, and the saved document is the input for the following step of the automated analysis (Fig. S7). Finally, the workflow makes tentative assignments of double bond configuration based on the order of elution and generates systematic names, fatty acid descriptors and retrieves LIPID MAPS IDs and common names from the LIPID MAPS database (Table S1). The relative quantification is summarized both in a table and as a bar chart (Fig. S8). A segmented bar chart showing a comparison of three replicates, or three fatty acid profiles can be generated with a separate python script from the output of the main workflow.

Fatty acid isomers are excluded from the analysis, if the feature is found to arise from over-oxidation of the isomer with the respective double bond in a position of one carbon closer to the carboxyl end of the fatty acid (Supplementary Information, section 3.1.2). The 37 mix fatty acid standard analysis (Table S2) informs the degree of over-oxidation that leads to artefacts at the same retention time for the *n*-(*x*+1) position at max. 1.5% for *cis* double bonds and max. 3% for *trans* double bonds. Fatty acids are also excluded from the analysis, if an amount exceeding 75% of the fatty acid in the sample is found in the associated Process Blank (Supplementary Information, section 3.2). Using the internal standard per-deuterated palmitic acid, the relative quantities of isomers were corrected by the amounts found in the associated Process Blank, where appropriate (In cases of between 1 and 75% of the fatty acid isomer that is present in the sample is also found in the Process Blank).

## Data availability statement

The data that support the findings of this study are available from QUT research data finder at: _.

## Code availability statement

Python source code, windows batch files and associated files are publicly available at: github.com/jphmenzel/jpmlipidomics

## Acknowledgements

The authors acknowledge funding from the Australian Research Council (ARC, DP190101486) and Waters Corporation (through the ARC linkage program, LP180100238) and support from the Queensland University of Technology (QUT) including access to the Central Analytical Research Facility. The authors thank the research team of Josef Cvačka for kindly providing a 4-Iodo-AMPP derivatized fatty acid extract (total fatty acid content of a lipid extract) of vernix caseosa. We gratefully acknowledge discussions and laboratory assistance by Dr. David Marshall (QUT). We thank Dr. Pawel Sadowski (QUT) for assistance with the Skyline Mass Spectrometry environment. We gratefully acknowledge the provision of heat-inactivated fetal bovine serum from the Translational Research Institute, Woolloongabba. We acknowledge ongoing technical support for mass spectrometry by Dr. Tyren Dodgen, Waters Corporation. “Python” and the Python logos are trademarks or registered trademarks of the Python Software Foundation, used by the authors with permission from the foundation.

## Author contributions

J.P.M. carried out derivatization reactions with assistance of R.S.E.Y. B.L.J.P. and J.P.M. designed the method for instrumental acquisition of LC-OzID-MS data. J.P.M. performed direct infusion ESI-MS, UPLC-OzID-MS and UPLC-OzID-MS/MS instrumental analysis and wrote all python source code, windows batch files and performed the data analysis. MCF7 cancer cells were grown in cell culture by A.B. under the guidance of S.T.H. LNCaP and SCD1-inhibited LNCaP cancer cells were grown in cell culture by J.C. under the guidance of L.M.B. The manuscript was written by J.P.M. with contributions of all co-authors. The study was motivated and supervised by S.J.B. All authors have given approval to the final version of the manuscript.

## Ethics declarations

Vernix caseosa was collected by the group of Josef Cvačka, Institute of Organic Chemistry and Biochemistry, Academy of Sciences of the Czech Republic, Prague, Czech Republic, with written informed parental consent and the work was approved by the Ethics Committee of the General University Hospital, Prague (910/09 S-IV).

## Supplementary Information

This material is available free of charge online. (Materials, data analysis, supplementary results)

## 1. Materials

### 1.1 Materials

Acetonitrile (ACN, Fisher Chemical, Optima LC/MS suitable for UHPLC-UV), 0.1% Formic Acid in acetonitrile (Fisher Chemical, Optima LC/MS), *N,N*-Dimethylformamide (DMF, Sigma-Aldrich, for HPLC, >99.9%), *N,N*-Diisopropylethylamine (DIPEA, Sigma Aldrich, >99%), *N*-(3-Dimethylaminopropyl)-*N’*-ethylcarbodiimide hydrochloride (EDC*HCl; Sigma Aldrich, 98%), 1-Hydroxybenzotriazole hydrate (HOBt; Sigma Aldrich, >97.0%), n-Pentane (Sigma-Aldrich, for HPLC, >99.0%), Metabolites in Frozen Human Plasma (Standard Reference Material 1950, National Institutes of Standards and Technology, NIST), Methanol (MeOH, Fisher Chemical, Optima LC/MS), Methyl *t*-butyl ether (MTBE, RCI Labscan, HPLC), Tetrabutylammonium hydroxide solution (Sigma-Aldrich, 40 wt. % in H_2_O), Sodium Chloride (NaCl, Sigma-Aldrich, SigmaUltra minimum 99.5%), Supelco 37 Component FAME Mix (certified reference material, TraceCERT in dichloromethane, varied conc., Sigma-Aldrich), and 0.1% Formic Acid in Water (Fisher Chemical, Optima LC/MS) were used as received.

## 2. Data analysis with windows batch files, python scripts and Skyline MS

**Figure S1:**
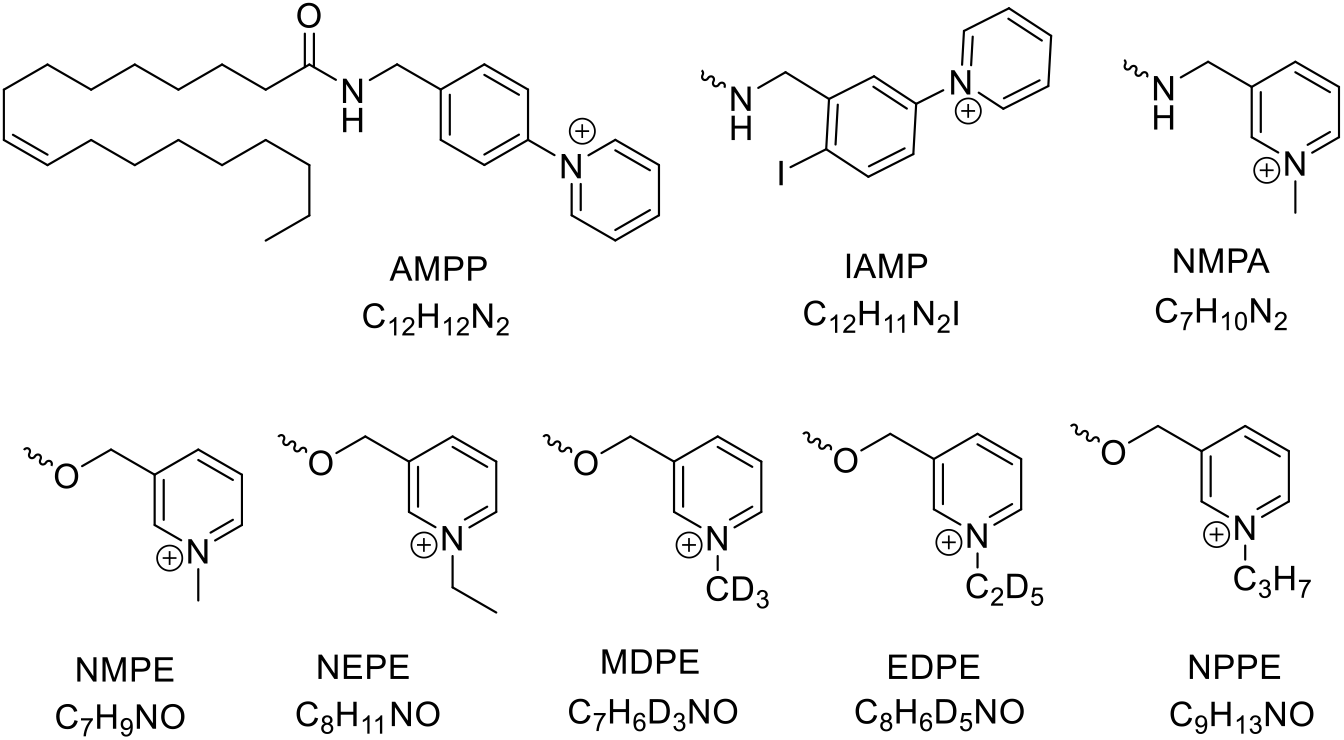
Four-letter-codes of derivatization agents and sum formula of the respective head groups (bound to the fatty acid either via an amide or ester bond) shown as derivatized oleic acid (C18:1n-9). When either of the derivatization agents is used in the workflow, only the four-letter-code shown is required for running the analysis, otherwise the sum formula of the head group is required.

### 2.1 Details on semi-automated data analysis with OzFAD algorithms

A general overview of the workflow is displayed in Figure S2.

**Figure S2:**
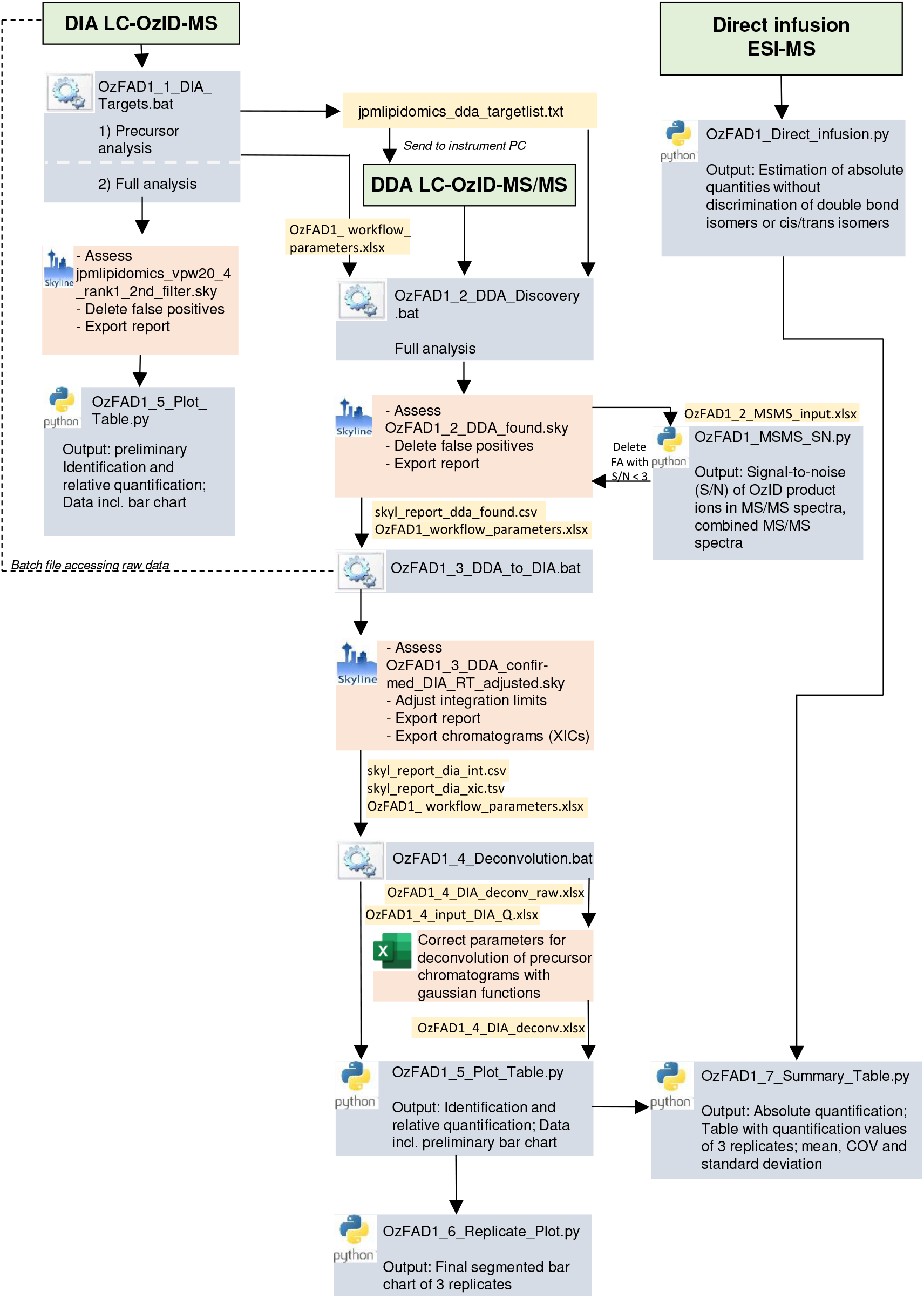
Overview of the analysis workflow including instrumental analysis and all steps that require user intervention. Instrumental analysis steps are shown in green boxes, scripts that perform automated analysis steps are shown in grey boxes. Symbols indicate whether all steps within the respective analysis step are automated by a batch file (which in turn may perform analysis steps using Skyline Runner and python scripts), whether the user is required to carry out an assessment step in a Skyline file that was generated by the previously executed script or whether the user is required to carry out manual review and adjustments in an excel file that was generated by the previously executed script.

### 2.2 Practical instructions on setting up the workflow OzFAD

The workflow consists of a collection of windows batch files and python files, which need to be in the correct directory to function together as intended. Further, some python packages are required. The workflow is written for use on a windows computer.

To get started, please make sure you have a python version and *Skyline* installed. Python code was edited with *Visual Studio Code* and windows batch files were edited with *Notepad++*. Several packages need to be installed for python programs to run without error messages (pandas, openpyxl, scipy, brainpy, numpy, matplotlib, bs4, requests, csv, statistics, subprocess). These can be installed in *Visual Studio code* using pip with commands such as: ‘pip install pandas’.

The folder structure required to use this workflow is shown in the following. Within the local user folder, create one folder called pythonprogramming and one called batchprogramming. In these, create folders according to the folder structure shown below and enter the files from the git-hub repository as shown. The Skyline runner executable is available from the Skyline webpages. There only needs to be one Import.log file in the folder workflow_log_files to get started. These log files show information on what Skyline (Runner) does during running of the workflow and can be used for troubleshooting in case a part of the workflow doesn’t finish correctly.

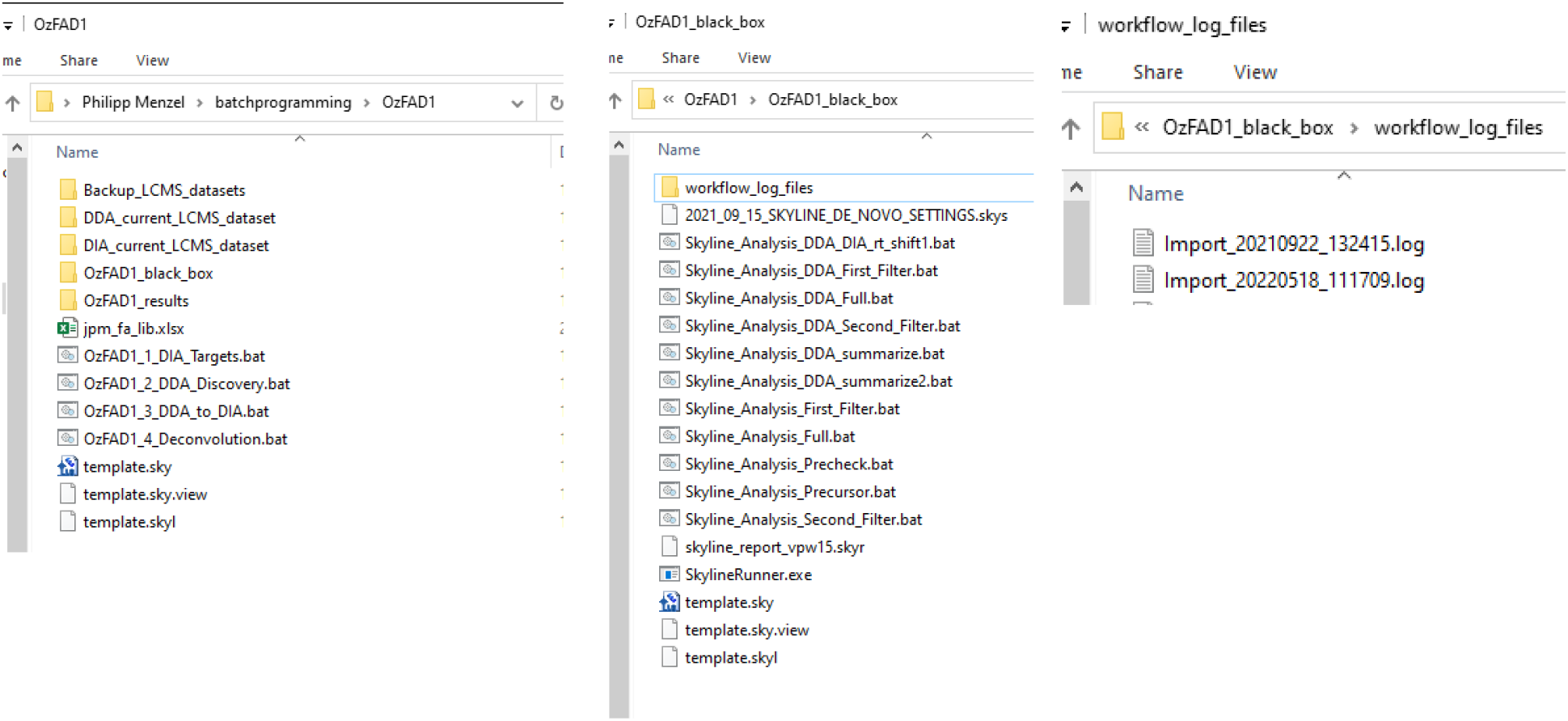
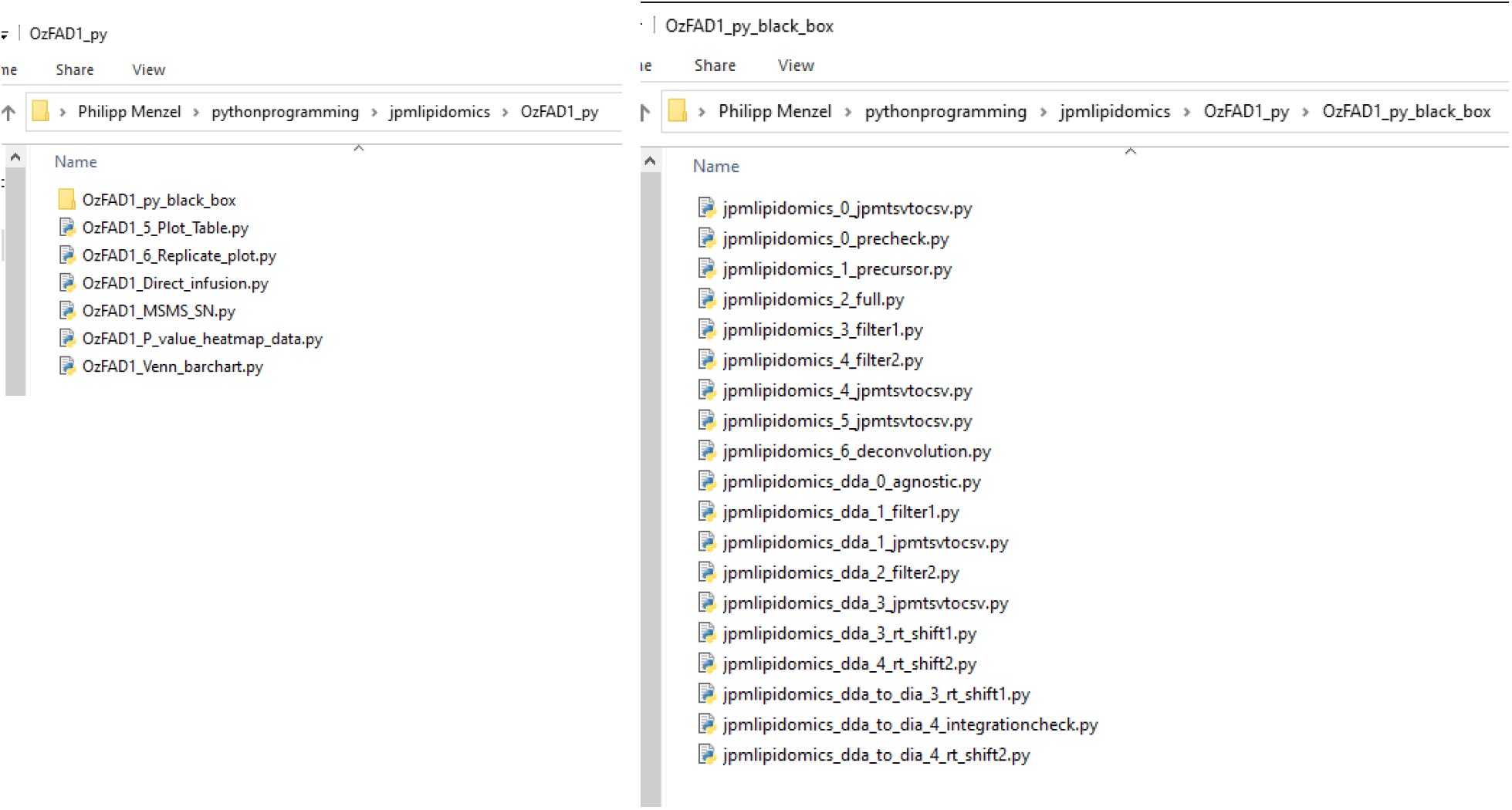

To successfully run the code, links within the batch files need to be updated to reflect the local folder structure. Open each batch file with Notepad++ and find and replace each instance of the local username to reflect your local folder structure as shown in the above screenshot. Perform this find and replace process also to update the files to your local Python version and path. Both needs to be updated in all batch files.

Finally, to run data through the workflow, always ensure that there is enough disk space available locally, that no instance of Skyline is opened as it may interfere with Skyline Runner and ensure that the raw files / folders are in the appropriate directories. Before running a batch file, make sure that the DIA LC-OzID-MS raw data file is in folder *‘DIA_current_LCMS_dataset’* and the DDA LC-OzID-MS/MS raw data file is in folder *‘DDA_current_LCMS_dataset’*. There should at any time only be one dataset each in these folders, respectively. The order in which the batch files are to be used is described in the Supplementary Information, section 2.1, Figure S2.

### 2.3 Algorithm for the determination of signal-to-noise values from OzID-MS/MS tandem mass spectra

The process to determine S/N values from OzID-MS/MS spectra is visualized below using the example of AMPP derivatized 9Z,15Z-tetracosadienoic acid (C24:2n-9,15) in MCF7 cells.

**Figure S3:**
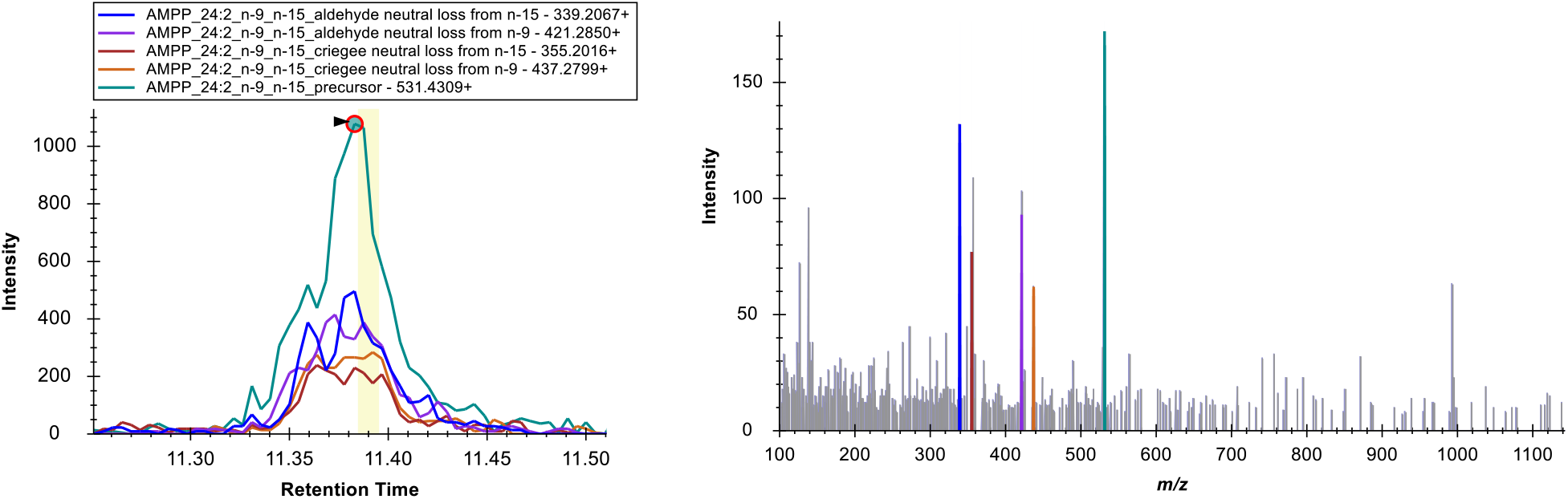
Left: Extracted ion chromatogram of C24:2n-9,15 from the data dependent acquisition (LC-OzID-MS/MS) of AMPP derivatized fatty acids in lipid extracts of MCF7 cells, replicate 1. Right: Associated OzID-MS/MS tandem mass spectrum at RT = 11.38 min. Displayed is each the visualization shown in Skyline, the legend and color coding from the chromatograms on the left applies to the mass spectrum on the right.

**Figure S4:**
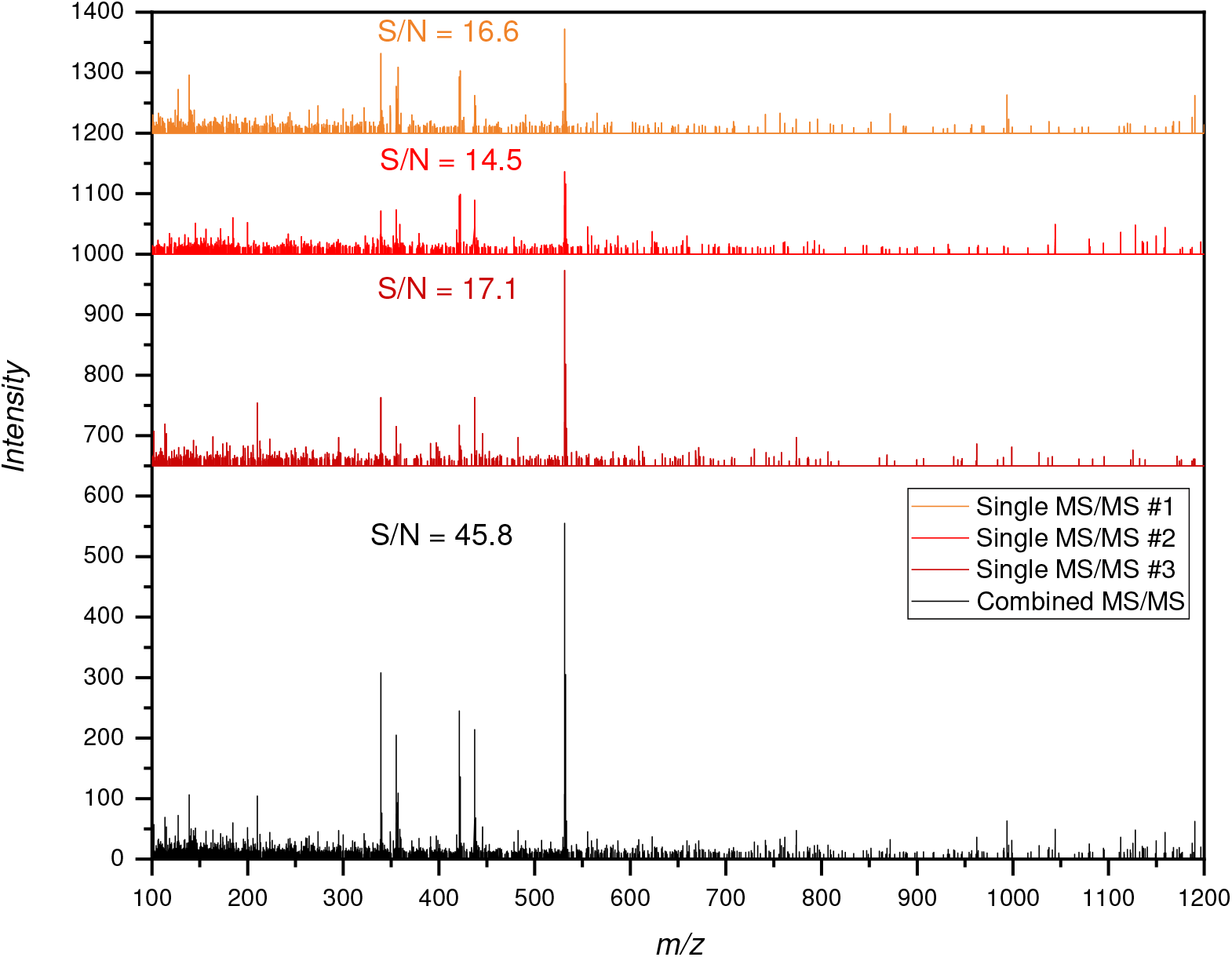
To demonstrate the procedure for the calculation of S/N values from OzID-MS/MS spectra, three MS/MS spectra from the chromatographic peak shown in Figure S4 were extracted and are displayed above the combined spectrum and S/N values for the fatty acid (C24:2n-9,15) are shown above each spectrum, respectively. The algorithm that calculates the S/N values reads an excel file containing the data of selected MS/MS spectra (data copied directly from Skyline to clipboard and into the excel input file). Multiple MS/MS spectra are combined by the algorithm and the combined spectra for each species are analyzed.

### 2.4 Detailed step-by-step guide to data analysis with the OzFAD (Ozone-enabled Fatty Acid Discovery) workflow

1. To ensure that fatty acid isomers are reliably identified and do not originate from background (glassware or pipette tips), always analyze a Process Blank associated with each set of samples in addition to the samples with the workflow. This step-by-step guide uses the datasets of the MCF7 cell line extract, replicate 1 to demonstrate the practical steps of the workflow.
2. Copy DIA LC-OzID-MS dataset into folder ‘DIA_current_LCMS_dataset’.
3. Start OzFAD_1_DIA_Targets.bat and enter input parameters as requested (example below).

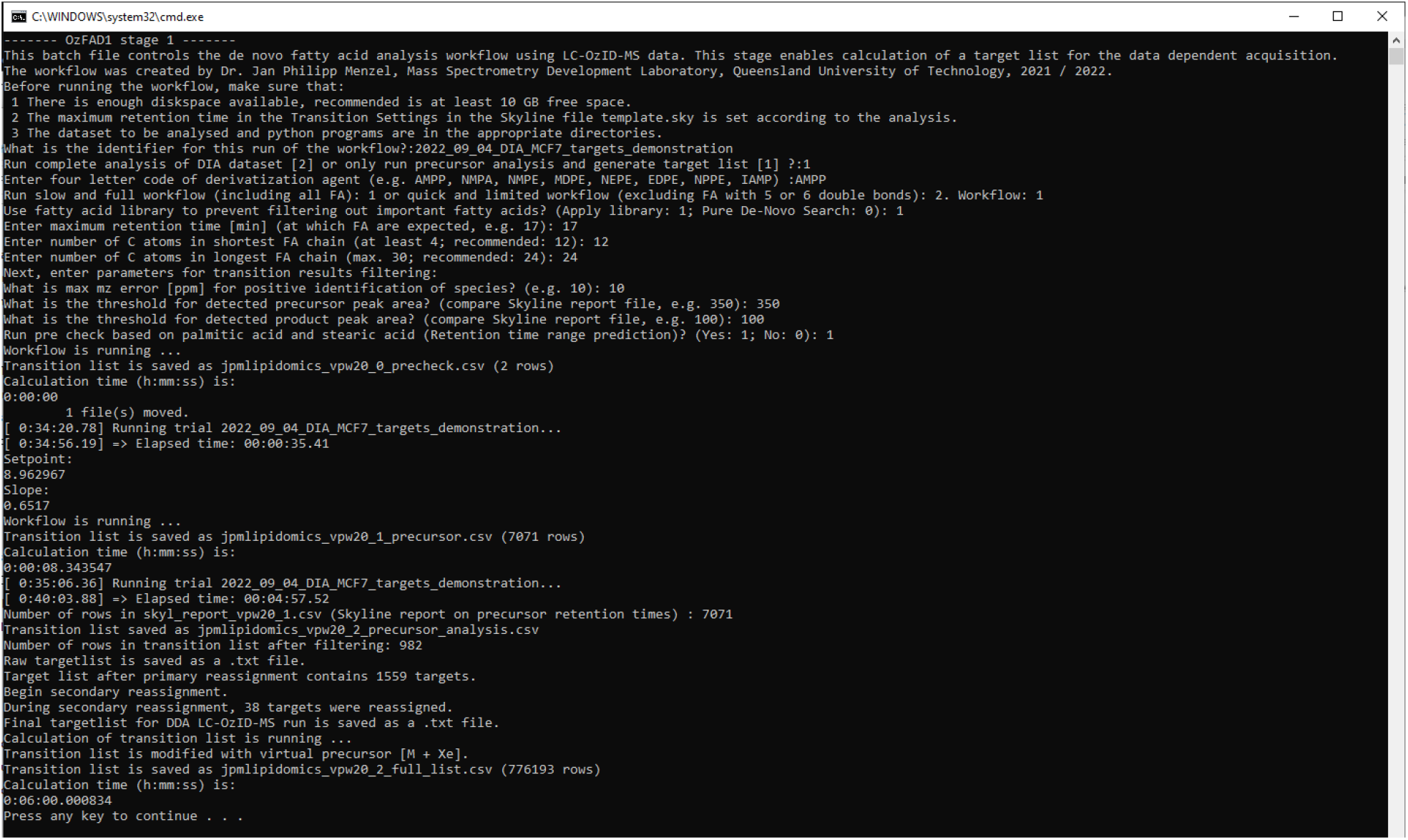
4. Rename target list and send to instrument PC. The separate raw target list is not to be used but serves as a reference to judge the target resampling algorithm if required.
5. Perform data-dependent acquisition (LC-OzID-MS/MS).
6. Copy DDA LC-OzID-MS/MS dataset into the respective folder.
7. Copy the target list generated in the previous step, named jpmlipidomics_dda_targetlist.txt into the folder OzFAD1. (For the purposes of demonstrating the steps in the workflow, a changed target list that is limited to fatty acids with chain length of 16 to 18 carbon atoms is used here.)
8. Copy the workflow parameters file generated in the previous step, named OzFAD1_workflow_parameters.xlsx into the folder OzFAD1. (An adapted file is used here with the same limitation (FA C16 - 18.)
9. Start batch file OzFAD1_2_DDA_Discovery.bat and choose workflow analysis mode. Three options are available: Full discovery, streamlined and library based. The library-based search can be used, when a selected number of defined isomers should be identified quickly (e.g., analysis of a replicate sample of one that has previously been characterized with the full or streamlined discovery workflow). The streamlined analysis is significantly faster than the full discovery mode, as it limits the search for polyunsaturated fatty acids with four or more double bonds to the ones that are listed in the fatty acid library. This reflects the diversity of natural fatty acid samples as there are more likely unexpected novel mono-, bis- and tris unsaturated fatty acids than tetra-, penta- or hexaunsaturated fatty acid isomers.

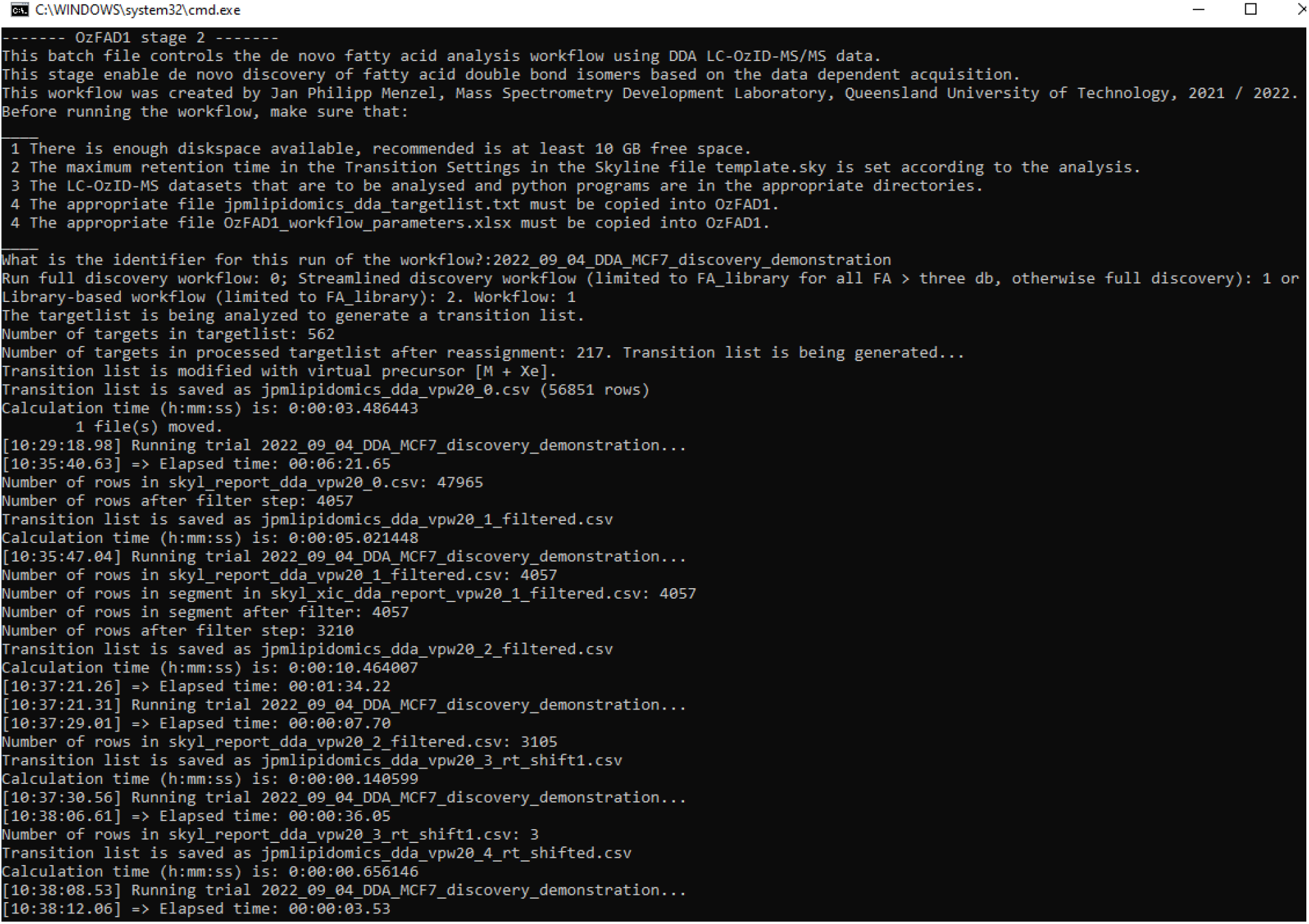
10. Open the Skyline file jpmlipidomics_dda_vpw20_2_filtered.sky in the folder that was created with the name of the identifier as entered above within the folder OzFAD1_results.
11. Manually filter the list of fatty acid species and delete all that do not clearly correspond to an isomer identifiable by the presence of the precursor fatty acid m/z and the respective product m/z values. The view of the extracted ion chromatograms and the MS/MS spectra at the relevant retention times in Skyline helps to quickly identify, which species to keep and which to delete, see below. Additionally, delete those features that arise from over-oxidation. These features are identified by coeluting OzID product ions of a double bond position one carbon closer to the methyl terminus that are not larger than 3% of the intensity of the latter.

**Figure S5:**
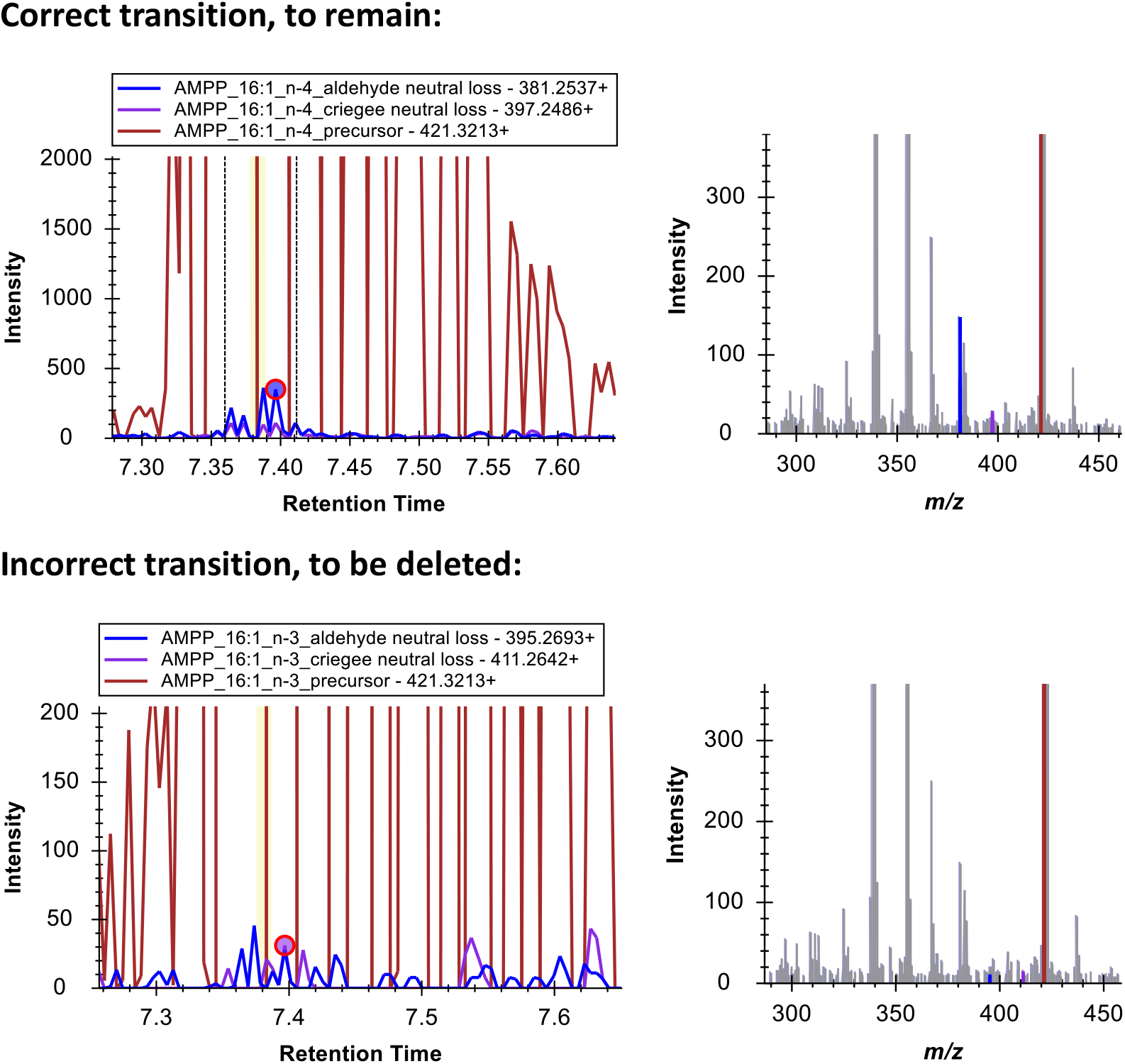
Chromatograms (data dependent acquisition) and tandem OzID mass spectra of a transition with OzID product ions above noise in comparison to a feature not significantly above noise.
12. For isomers that are low in abundance, export tandem mass spectra (OzID MS/MS; at least one spectrum, three recommended) from Skyline (from MS/MS view) as a Table and enter each into the input excel file for the S/N calculation script OzFAD1_MSMS_SN.py. Run script to determine the signal-to-noise ratio for each species and keep only those species in Skyline with S/N > 10 (S/N > 3 for those species that are to be tentatively identified, but not quantified). The screenshot of the script after being run in Visual Studio Code shows that FA 16:1n-3 is indeed not present above noise in MCF7 cell extracts as also seen in the Skyline view shown above. For this demonstration, all species other than hexadecenoic acids were deleted from the Skyline file. Export a Skyline report with the report template provided with this publication, the report csv file needs to be named skyl_report_dda_found.csv to enter the next stage of the workflow. The report can be changed into a transition list for reuse with Skyline on replicate datasets by changing the top row to the descriptions used in any other Skyline transition list generated by the workflow as well as deleting the last few columns while keeping explicit retention times and defining retention time windows.

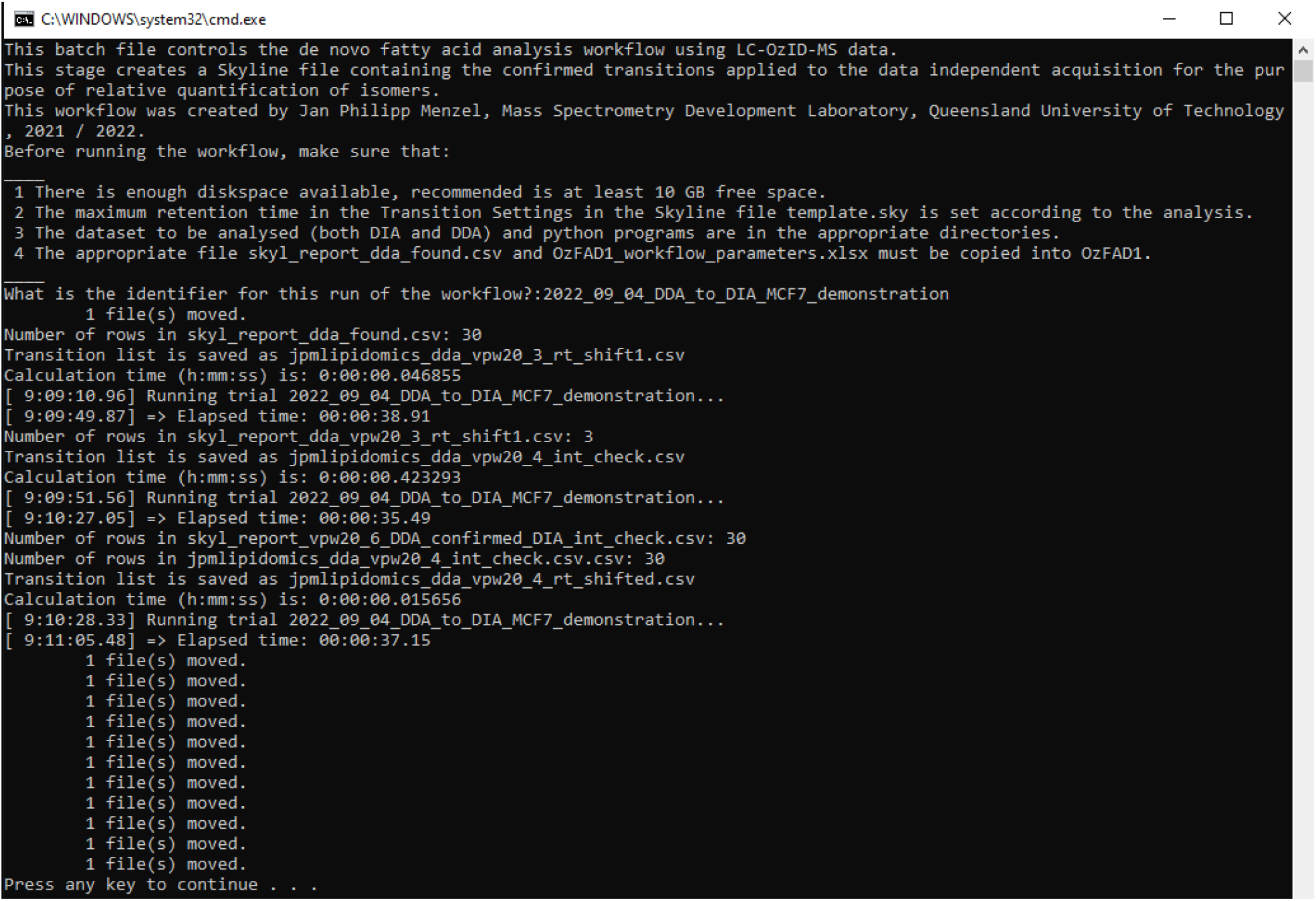
13. Copy the previous file OzFAD1_workflow parameter.xlsx and the file skyl_report_dda_found.csv into the folder OzFAD1 and run the script OzFAD1_3_DDA_to_DIA.bat as shown below.

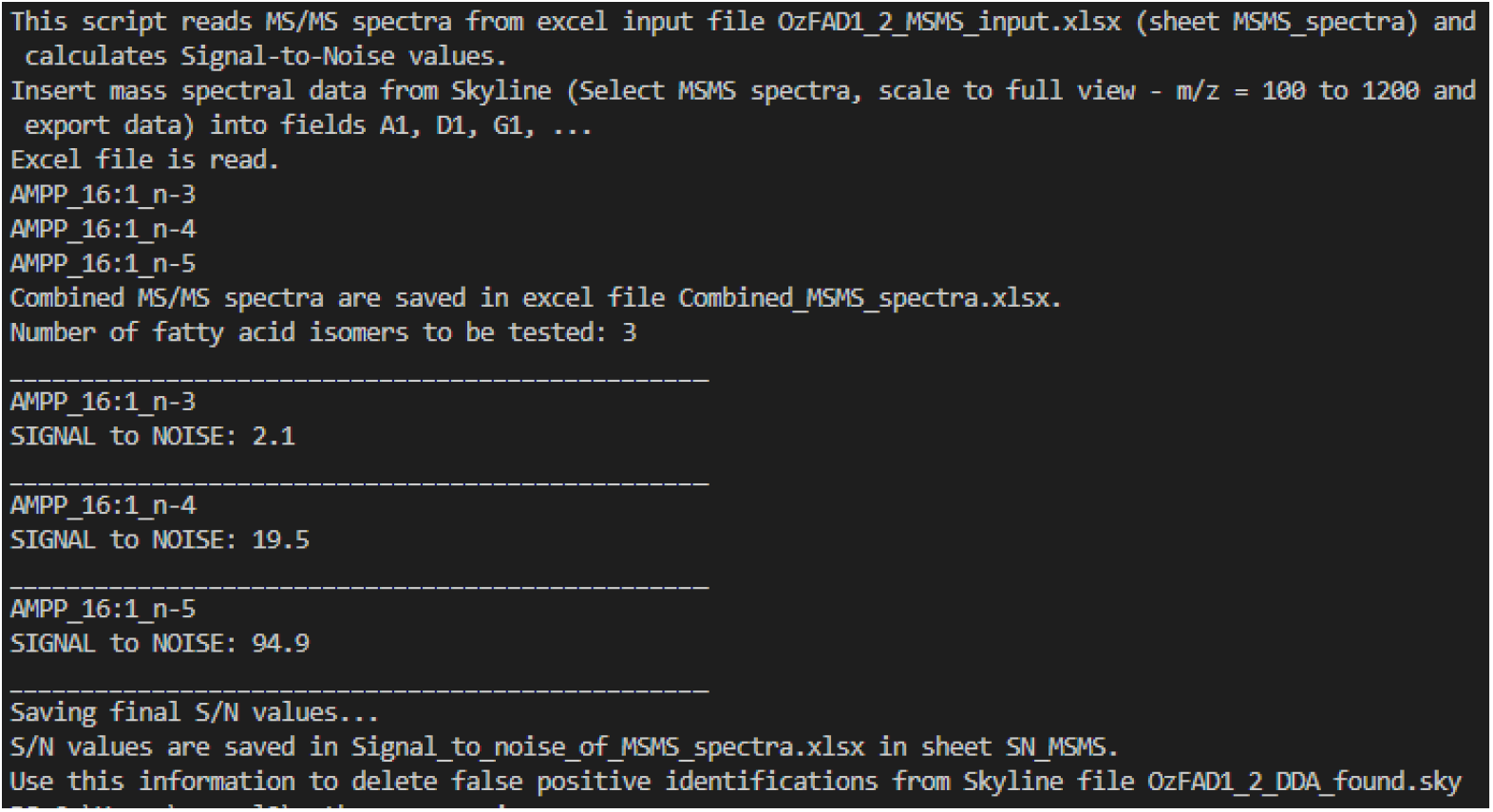
14. Open the Skyline file jpmlipidomics_vpw20_6_DDA_confirmed_DIA_rt_shifted.sky in the folder associated with this run of the workflow (in OzFAD1_results). Adjust the integration borders of each isomer. Ideally, the center of the OzID chromatograms would align with the center between the integration limits and the integration limits would capture the entire OzID chromatogram peak width. Export a report (same report template as used earlier) into the file skyl_report_dia_int.csv and export chromatograms into the file skyl_report_dia_xic.tsv. Further, carry out the analysis of the Process Blank equally to this step to identify, if some species need to be deleted from the sample due to their presence in both Sample and associated Process Blank in similar amounts.
15. Transfer the reports generated in step 14 as well as the file OzFAD1_workflow_parameters.xlsx into the folder OzFAD1 and run the batch file OzFAD1_4_Deconvolution.bat.

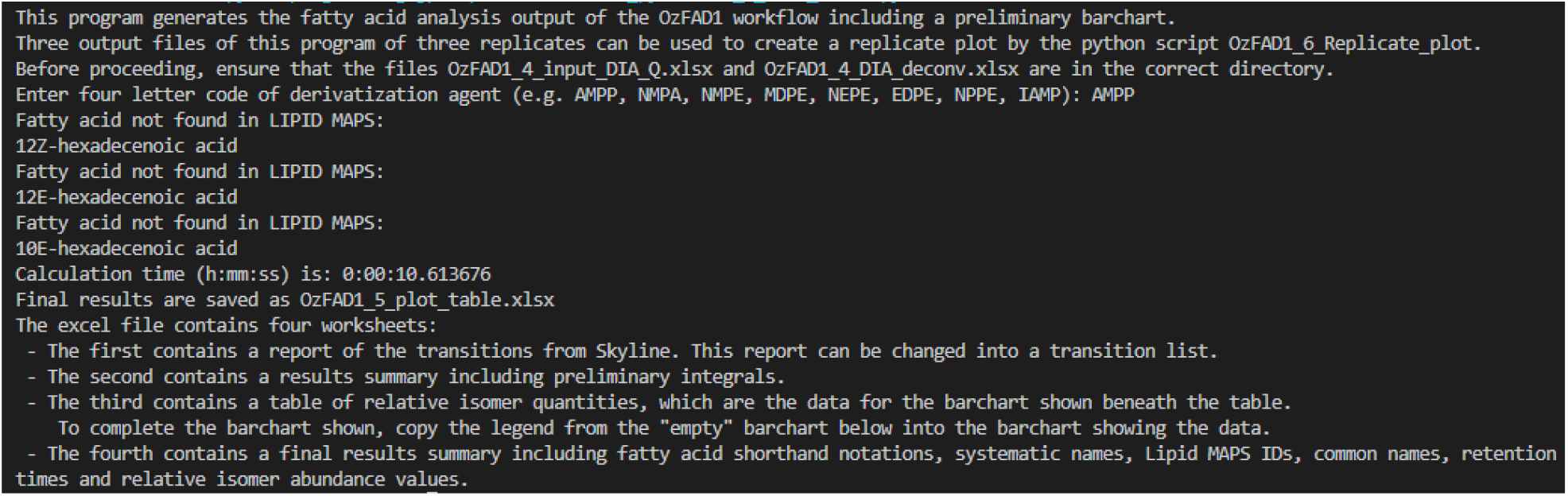
16. Open the file OzFAD1_4_DIA_deconv_raw.xlsx, go to sheet DC_16_1 (respectively all other sheets for each isomer group) and adjust the parameters for the fitting of gaussians to the precursor chromatogram (column B). The algorithm initially sets the ratios of the integrals of the gaussians according to the ratios of the OzID product ions (integrals shown in column E) as detected in the data independent acquisition. The width of each gaussian within one isomer group is set to be equal, field B4 can be used to adjust these. Column G indicates, which isomer feature corresponds to the one with maximum intensity. The retention time position of each gaussian is set according to the retention times of each feature in Skyline. To conclude this analysis step, some height and position parameters for some gaussians need to be adjusted to match the sum of the gaussians to the precursor chromatograms. Often there are many features that coelute too closely and are present at widely differing intensities. Consequently, deconvolution can then not be carried out and the ratio of the OzID Integrals needs to be used to provide the best estimate of the contribution of the respective isomer to the precursor chromatogram. The graphs below represent the initially automatically generated plot and the graph with manually refined values. In this example, the deconvolution can only be done based on the gaussian fit for the species FA 16:1*n*-7, FA 16:1*n*-9 and FA 16:1*n*-10.

**Figure S6:**
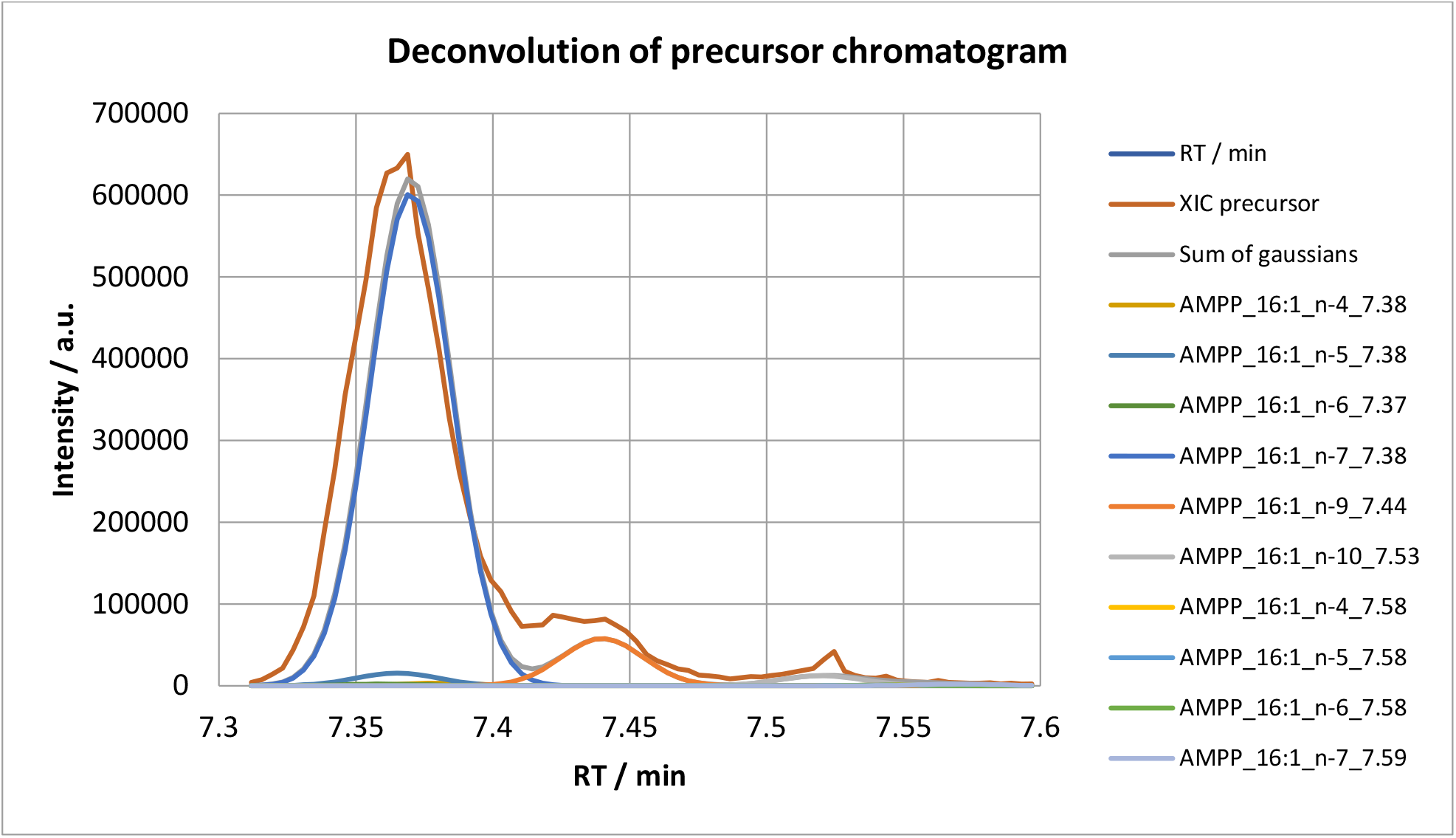
Computationally generated suggested start values for deconvolution of an example precursor chromatogram (FA 16:1 in MCF7, replicate 1) with gaussians representing the contribution of each fatty acid isomer.

**Figure S7:**
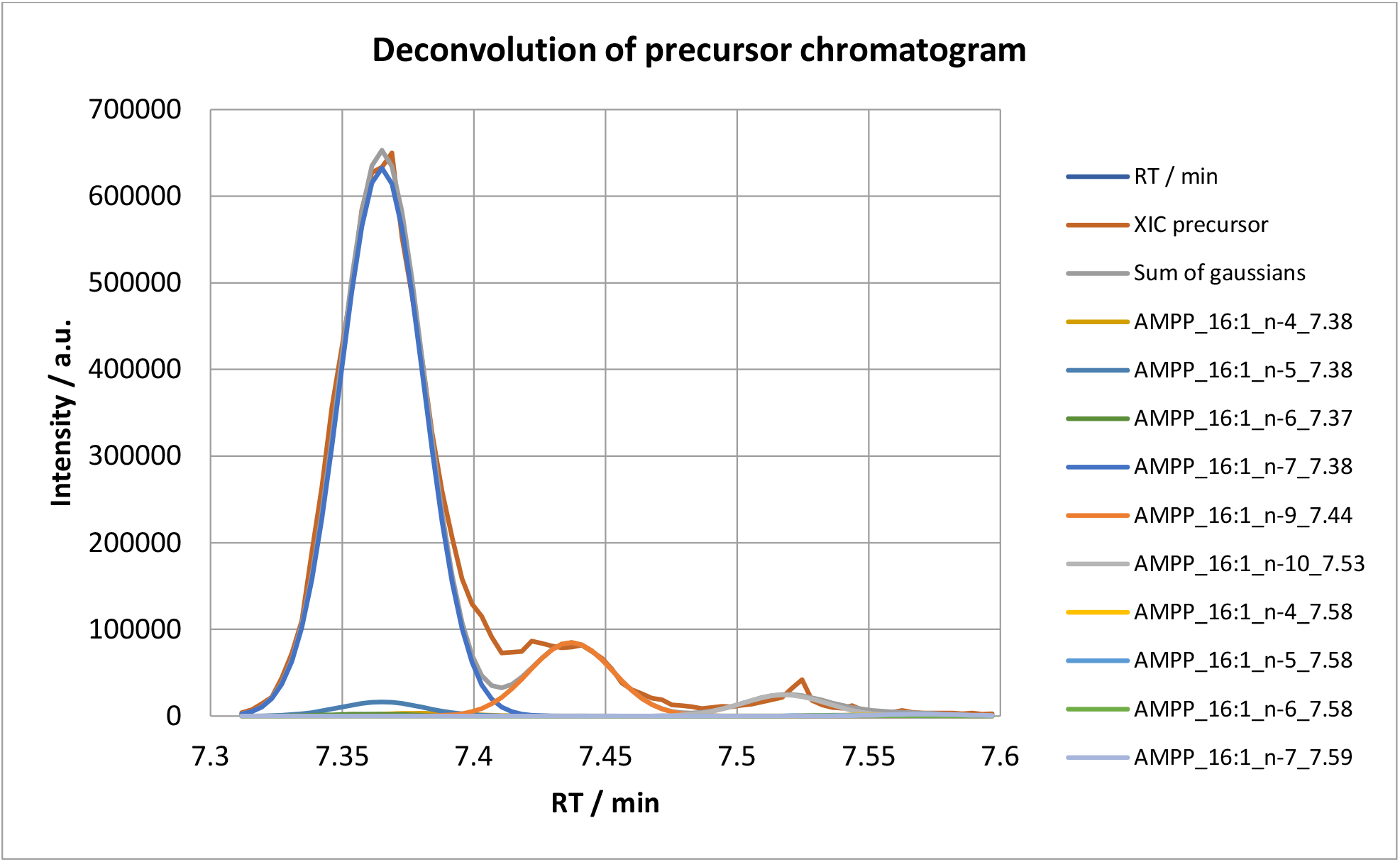
Deconvolution of an example precursor chromatogram (FA 16:1 in MCF7, replicate 1) after manual refining of parameters with gaussians representing the contribution of each fatty acid isomer.
17. Transfer the files OzFAD1_4_DIA_deconv_raw.xlsx (with adjusted parameters) and OzFAD1_4_input_DIA_Q.xlsx to the folder pythonprogramming and run the python script OzFAD1_5_Plot_Table.py, see below screenshot.

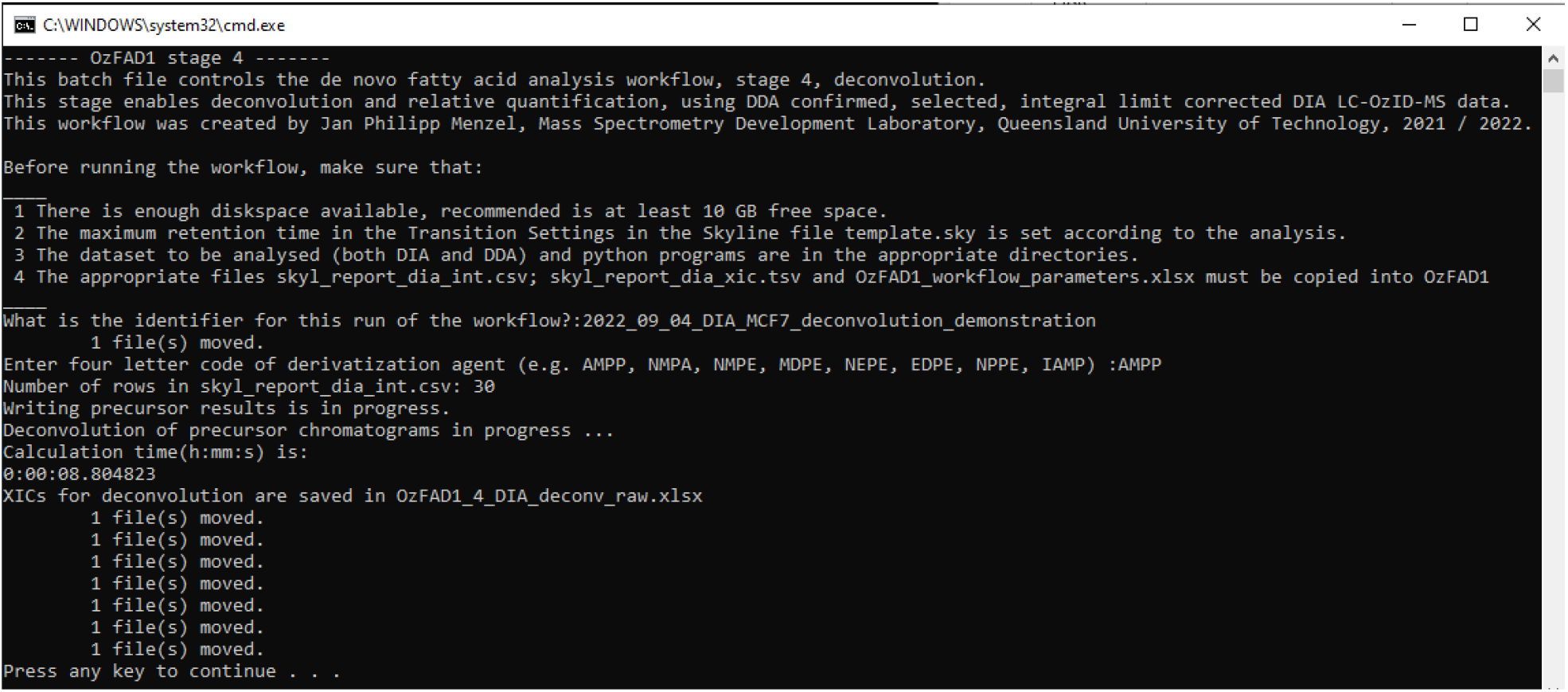

The excel file that is produced contains the following bar chart and Table based on the analysis of hexadecenoic acids in MCF7 cells, replicate 1. The values are not corrected for the Process Blank in case an isomer is present in the Process Blank in significant amounts. Sapienic acid is for example often detected in a Process Blank but may be present at much higher concentration in the sample under investigation.

**Table S1:** Results summary of hexadecenoic acids in MCF7 cells, replicate 1.

| Fatty acid (n-x) | FA shorthand | Systematic name | LipidMAPS ID | Common Name | RT / min | Relative isomer abundance / % |
| --- | --- | --- | --- | --- | --- | --- |
| C16:1n-4 | 16:1(12Z) | 12Z-hexadecenoic acid | Not found in LIPID MAPS. | — | 7.37 | 0.89 |
| C16:1n-5 | 16:1(11Z) | 11Z-hexadecenoic acid | LMFA01030262 | cis-Palmitvaccenic acid | 7.37 | 2.1 |
| C16:1n-6 | 16:1(10Z) | 10Z-hexadecenoic acid | LMFA01030058 | cis-10-palmitoleic acid | 7.37 | 0.3 |
| C16:1n-7 | 16:1(9Z) | 9Z-hexadecenoic acid | LMFA01030056 | cis-9-palmitoleic acid | 7.37 | 82 |
| C16:1n-9 | 16:1(7Z) | 7Z-hexadecenoic acid | LMFA01030055 | 7Z-palmitoleic acid | 7.39 | 11 |
| C16:1n-10 | 16:1(6Z) | 6Z-hexadecenoic acid | LMFA01030267 | Sapienic acid | 7.52 | 3 |
| C16:1n-4_(E) | 16:1(12E) | 12E-hexadecenoic acid | Not found in LIPID MAPS. | — | 7.52 | 0.17 |
| C16:1n-5_(E) | 16:1(11E) | 11E-hexadecenoic acid | LMFA01030261 | Lycopodic acid | 7.52 | 0.13 |
| C16:1n-6_(E) | 16:1(10E) | 10E-hexadecenoic acid | Not found in LIPID MAPS. | — | 7.52 | 0.1 |
| C16:1n-7_(E) | 16:1(9E) | 9E-hexadecenoic acid | LMFA01030057 | trans-9-palmitoleic acid | 7.54 | 0.33 |

**Figure S8:**
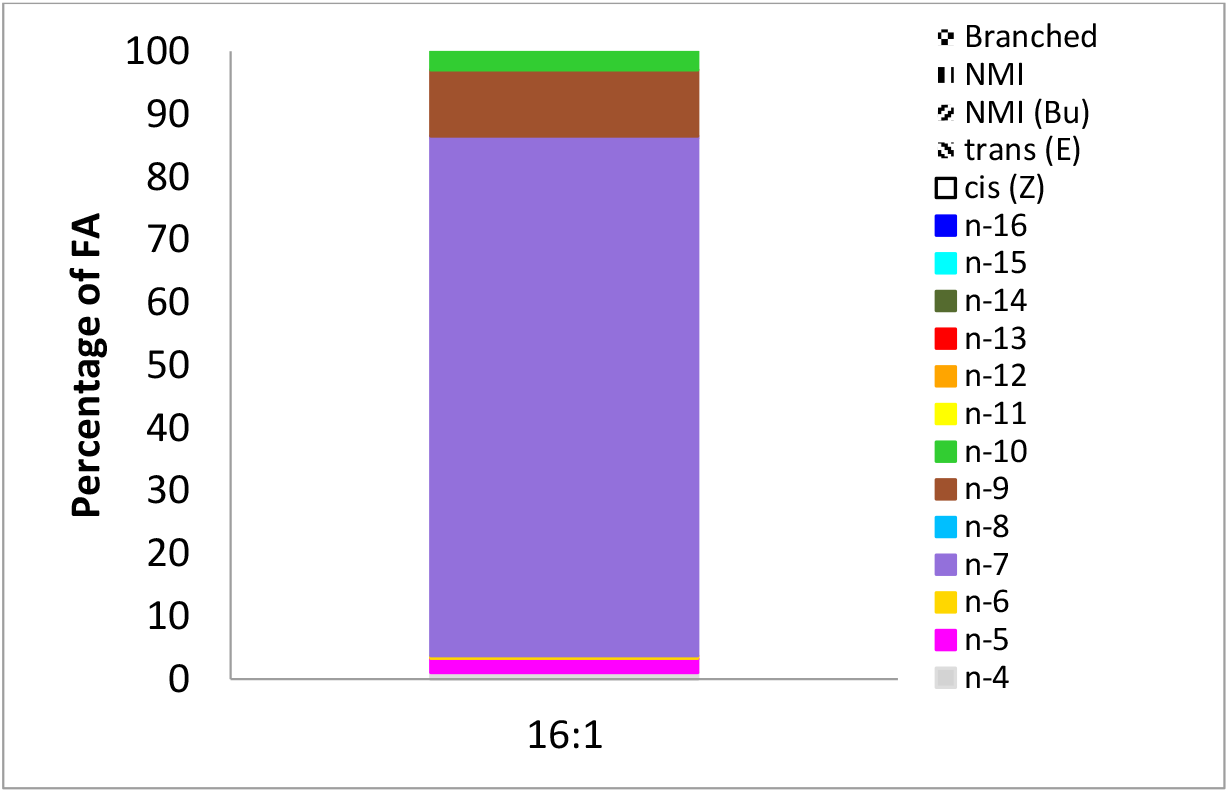
Bar chart for relative isomer abundance of hexadecenoic acids in MCF7 cells, replicate 1. The abundance of trans fatty acids is so low that the respective segments in the bar appear not clearly visible.

## 3. Results

### 3.1 Fatty acid standards – benchmarking quantification

The analytical standard SUPELCO FAME 37mix was used to benchmark the discovery workflow including the estimation of absolute quantities and relative quantities of fatty acids. 15 μL of the FAME 37mix solution dichloromethane were subjected to hydrolysis and AMPP derivatization equivalent to the other samples described in this work after evaporation of the dichloromethane under ambient conditions.

**Table S2:** Absolute and relative quantities of fatty acids derived from the respective fatty acid methyl esters (FAME) in the 15 μL of the analytical standard solution.

| Fatty acid species | Systematic Name | LIPID MAPS ID | n [nmol] |
| --- | --- | --- | --- |
| C14:1n-5 | 9Z-tetradecenoic acid | LMFA01030051 | 12.48 |
| C15:1n-5 | 10Z-pentadecenoic acid | LMFA01030259 | 11.79 |
| C16:1n-7 | 9Z-hexadecenoic acid | LMFA01030056 | 11.18 |
| C17:1n-7 | 10Z-heptadecenoic acid | LMFA01030283 | 10.62 |
| C18:1n-9 | 9Z-octadecenoic acid | LMFA01030002 | 20.24 |
| C18:1n-9t | 9E-octadecenoic acid | LMFA01030073 | 10.12 |
| C20:1n-9 | 11Z-eicosenoic acid | LMFA01030085 | 9.24 |
| C22:1n-9 | 13Z-docosenoic acid | LMFA01030089 | 8.51 |
| C24:1n-9 | 15Z-tetracosenoic acid | LMFA01030092 | 7.88 |
| C18:2n-6,9 | 9Z,12Z-octadecadienoic acid | LMFA01030120 | 10.19 |
| C18:2n-6t,9t | 9E,12E-octadecadienoic acid | LMFA01030123 | 10.19 |
| C20:2n-6,9 | 11Z,14Z-eicosadienoic acid | LMFA01031043 | 9.30 |
| C22:2n-6,9 | 13Z,16Z-docosadienoic acid | LMFA01030405 | 8.56 |
| C18:3n-3,6,9 | 9Z,12Z,15Z-octadecatrienoic acid | LMFA01030152 | 10.26 |
| C18:3n-6,9,12 | 6Z,9Z,12Z-octadecatrienoic acid | LMFA01030141 | 10.26 |
| C20:3n-3,6,9 | 11Z,14Z,17Z-eicosatrienoic acid | LMFA01030378 | 9.36 |
| C20:3n-6,9,12 | 8Z,11Z,14Z-eicosatrienoic acid | LMFA01030387 | 9.36 |
| C20:4n-6,9,12,15 | 5Z,8Z,11Z,14Z-eicosatetraenoic acid | LMFA01030001 | 9.42 |
| C20:5n-3,6,9,12,15 | 5Z,8Z,11Z,14Z,17Z-eicosapentaenoic acid | LMFA01030759 | 9.48 |
| C22:6n-3,6,9,12,15,18 | 4Z,7Z,10Z,13Z,16Z,19Z-docosahexaenoic acid | LMFA01030185 | 8.76 |
| C4:0 | butyric acid | LMFA01010004 | 58.75 |
| C6:0 | hexanoic acid | LMFA01010006 | 46.09 |
| C8:0 | octanoic acid | LMFA01010008 | 37.92 |
| C10:0 | decanoic acid | LMFA01010010 | 32.21 |
| C11:0 | undecanoic acid | LMFA01010011 | 14.98 |
| C12:0 | dodecanoic acid | LMFA01010012 | 27.99 |
| C13:0 | tridecanoic acid | LMFA01010013 | 13.14 |
| C14:0 | tetradecanoic acid | LMFA01010014 | 24.75 |
| C15:0 | pentadecanoic acid | LMFA01010015 | 11.70 |
| C16:0 | hexadecanoic acid | LMFA01010001 | 33.28 |
| C17:0 | heptadecanoic acid | LMFA01010017 | 10.55 |
| C18:0 | octadecanoic acid | LMFA01010041 | 20.10 |
| C20:0 | eicosanoic acid | LMFA01010020 | 18.37 |
| C21:0 | heneicosanoic acid | LMFA01010021 | 8.81 |
| C22:0 | docosanoic acid | LMFA01010022 | 16.92 |
| C23:0 | tricosanoic acid | LMFA01010023 | 8.14 |
| C24:0 | tetracosanoic acid | LMFA01010024 | 15.68 |

#### 3.1.1 Estimation of absolute quantities by direct infusion ESI-MS

**Figure S9:**
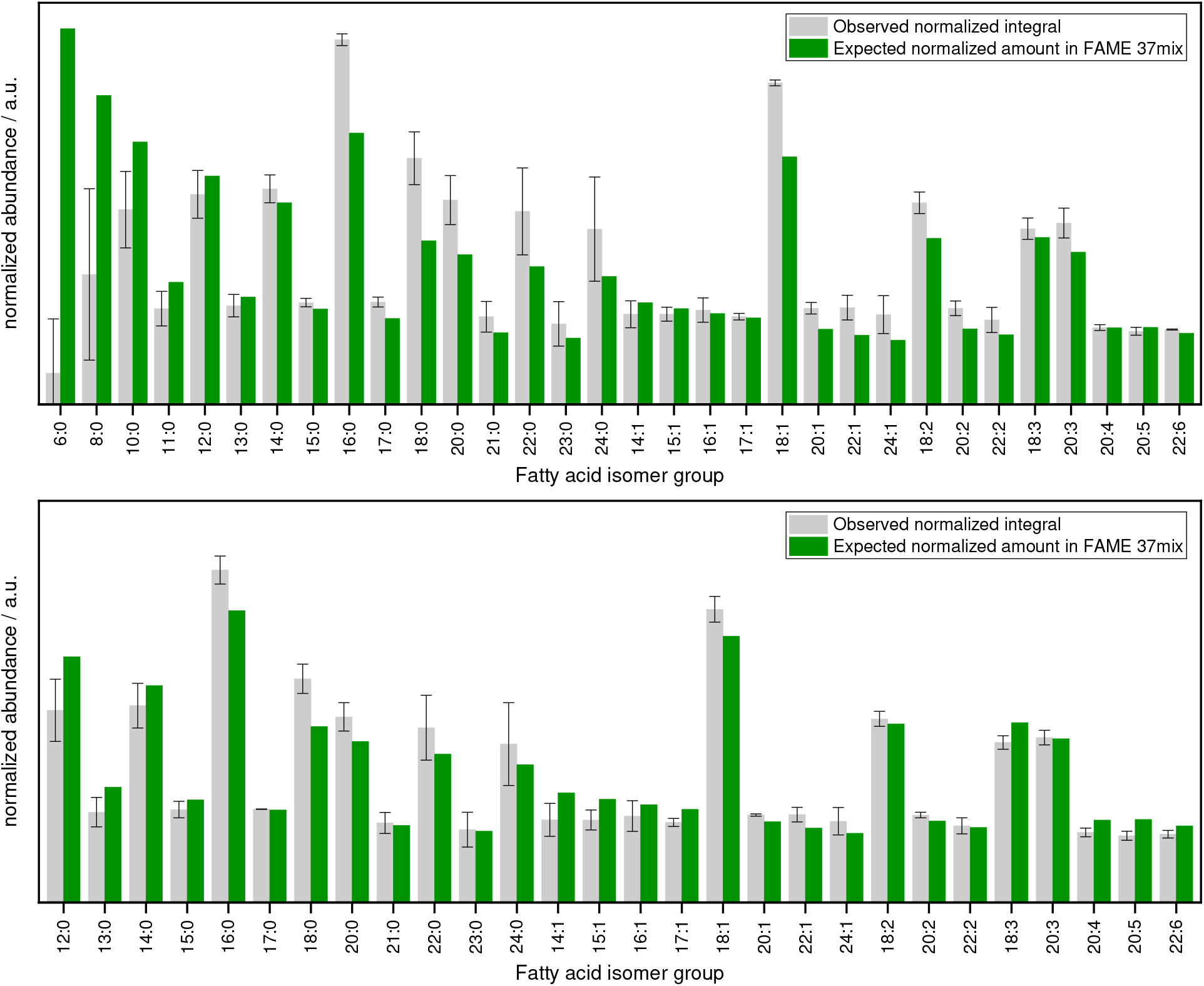
Comparison of observed relative quantities of non-isomeric fatty acids by direct infusion ESI-MS to the expected quantities of the FAME 37mix standard fatty acid solution. Each distribution was normalized to match the sum of the fatty acids of the distributions. Note that short chain fatty acids are underrepresented in the detected values likely due to partial evaporation during sample preparation and the hydrolysis and derivatization procedure. Above: all fatty acids included in the FAME 37mix standard sample are shown. Below: Fatty acids shorter than 12 carbon atoms are excluded from the normalization.

The mean of the coefficient of variation of the absolute quantities observed for fatty acids in the 37mix standard from C12 to C24 (Figure S9, lower bar chart) is 0.10 ± 0.07 (precision of quantitation – comparison of different fatty acid isomer groups – on average 10% of the determined value). The mean accuracy of quantification (mean absolute deviation between observed and expected normalized quantity divided by the expected normalized quantity) for the same fatty acid isomer groups is 15% ± 7%. Thus, quantities reported herein can serve as an approximate measure of the ratios and absolute values of fatty acid isomer groups present in the sample.

#### 3.1.2. LC-OzID-MS and LC-OzID-MS/MS analysis of the 37mix fatty acid standard

**Figure S10:**
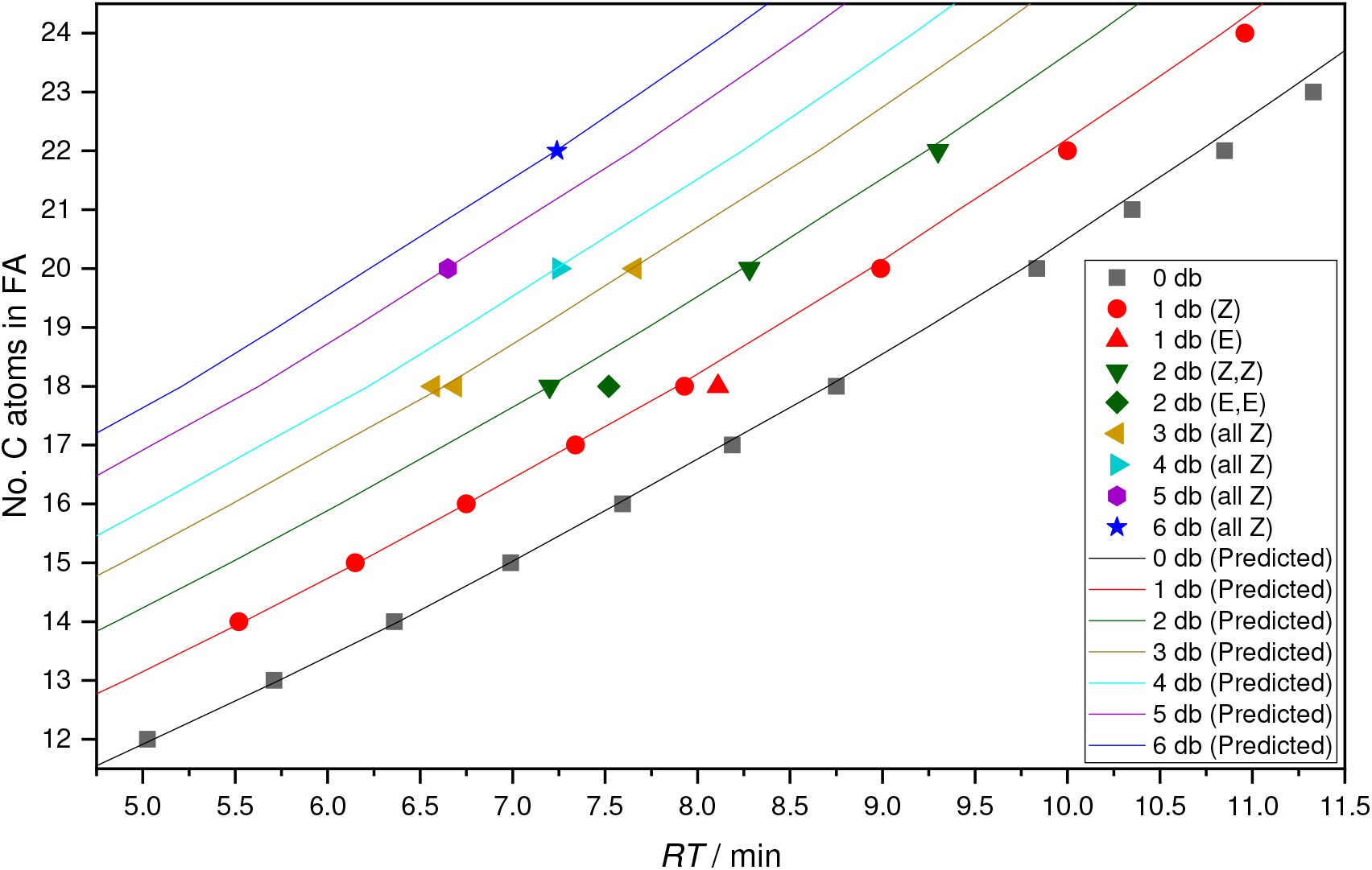
Retention times of AMPP derivatized fatty acids of the 37mix standard sample and predicted retention times of fatty acids with varied equivalent chain lengths and number of double bonds. The prediction method is empirically based on the elution of the fatty acids in this 37mix sample but applies as a general retention time range prediction with an estimated error <15% for all fatty acids. Generally, the retention time of palmitic and stearic acid are the input values for the retention time range prediction that is used within the workflow to limit the search for chromatographic features that represent fatty acid isomers. The tolerance of ± 15% of the RT value or ± 1 min, whichever is larger, is used to allow detection of branched and trans fatty acid isomers, which are chromatographically shifted.

**Figure S11:**
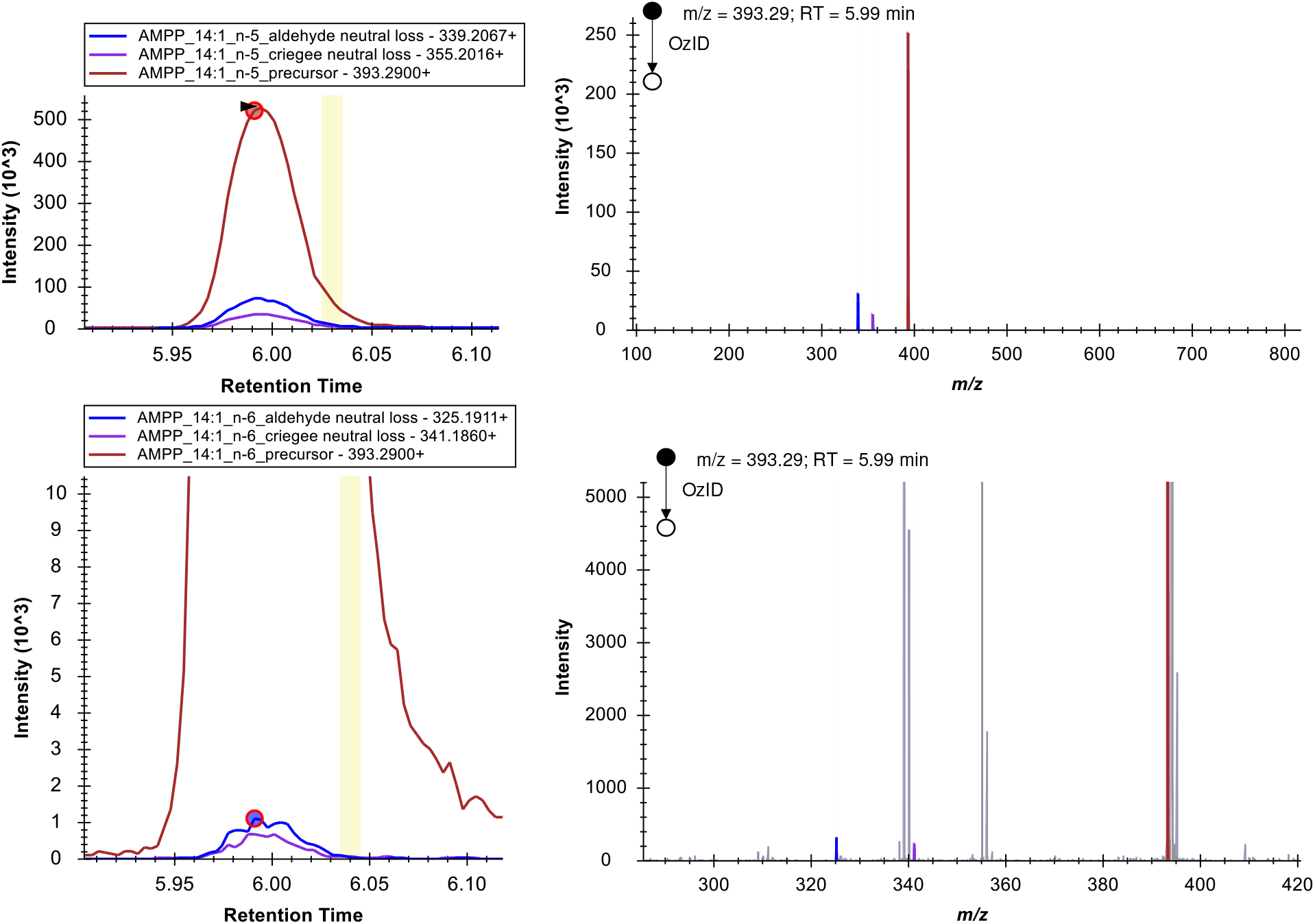
Chromatograms and mass spectra of the data dependent acquisition of the 37mix standard. Apart from the expected OzID product ions for FA 14:1*n*-5, a small degree of over-oxidation is observable as product ions with equal masses to the OzID product ions of the *n*-6 position.

**Figure S12:**
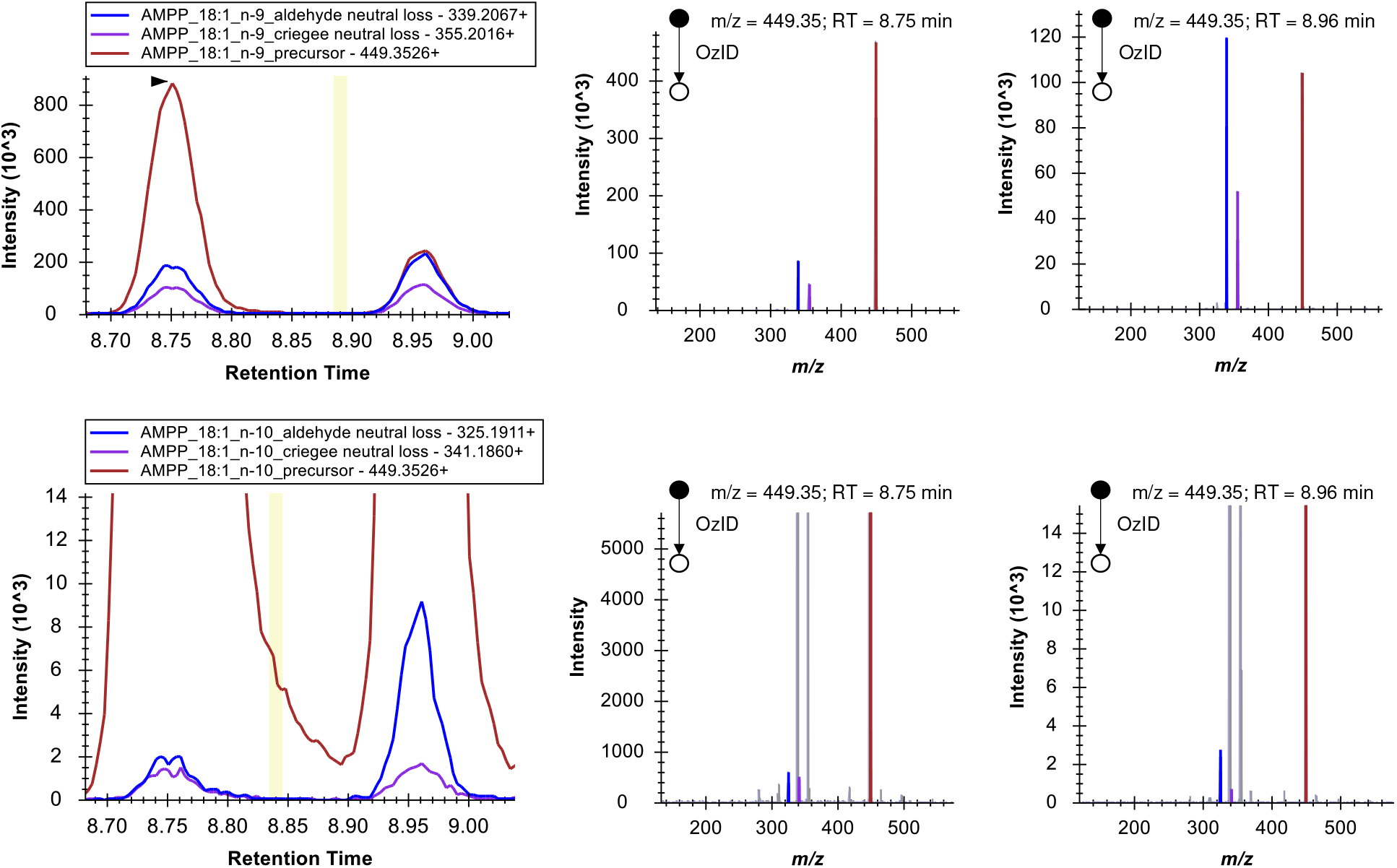
Chromatograms and mass spectra of the data dependent acquisition of the 37mix standard. Apart from the expected OzID product ions for FA 18:1*n*-9, a small degree of over-oxidation is observable as product ions with equal masses to the OzID product ions of the *n*-10 position.

**Figure S13:**
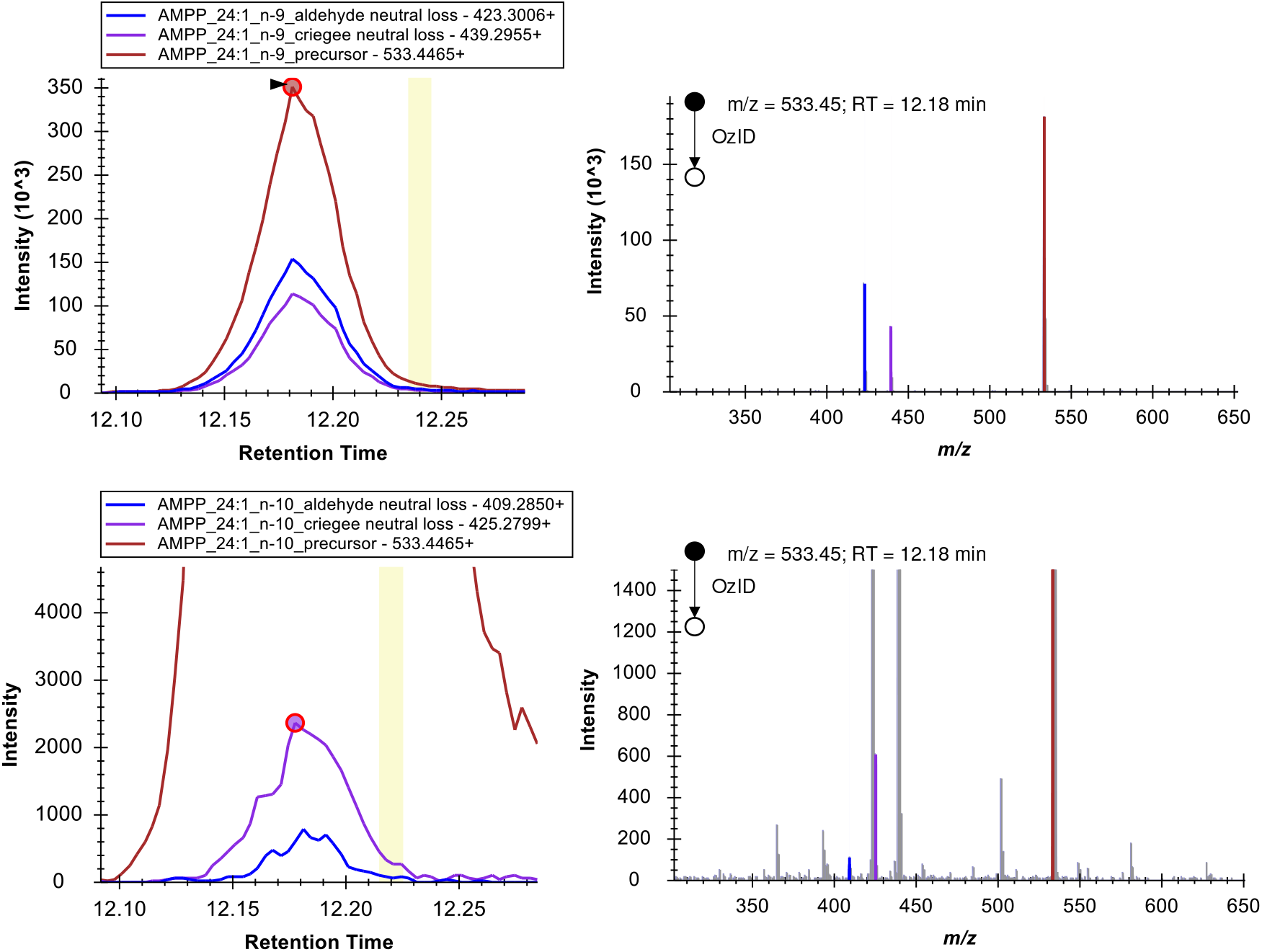
Chromatograms and mass spectra of the data dependent acquisition of the 37mix standard. Apart from the expected OzID product ions for FA 24:1*n*-9, a small degree of over-oxidation is observable as product ions with equal masses to the OzID product ions of the *n*-10 position. Note that due to the higher number of carbon atoms the +2 Isotope of the aldehyde product ion arising from ozonolysis on the n-9 position is isobaric with the criegee ion arising from over-oxidation.

**Figure S14:**
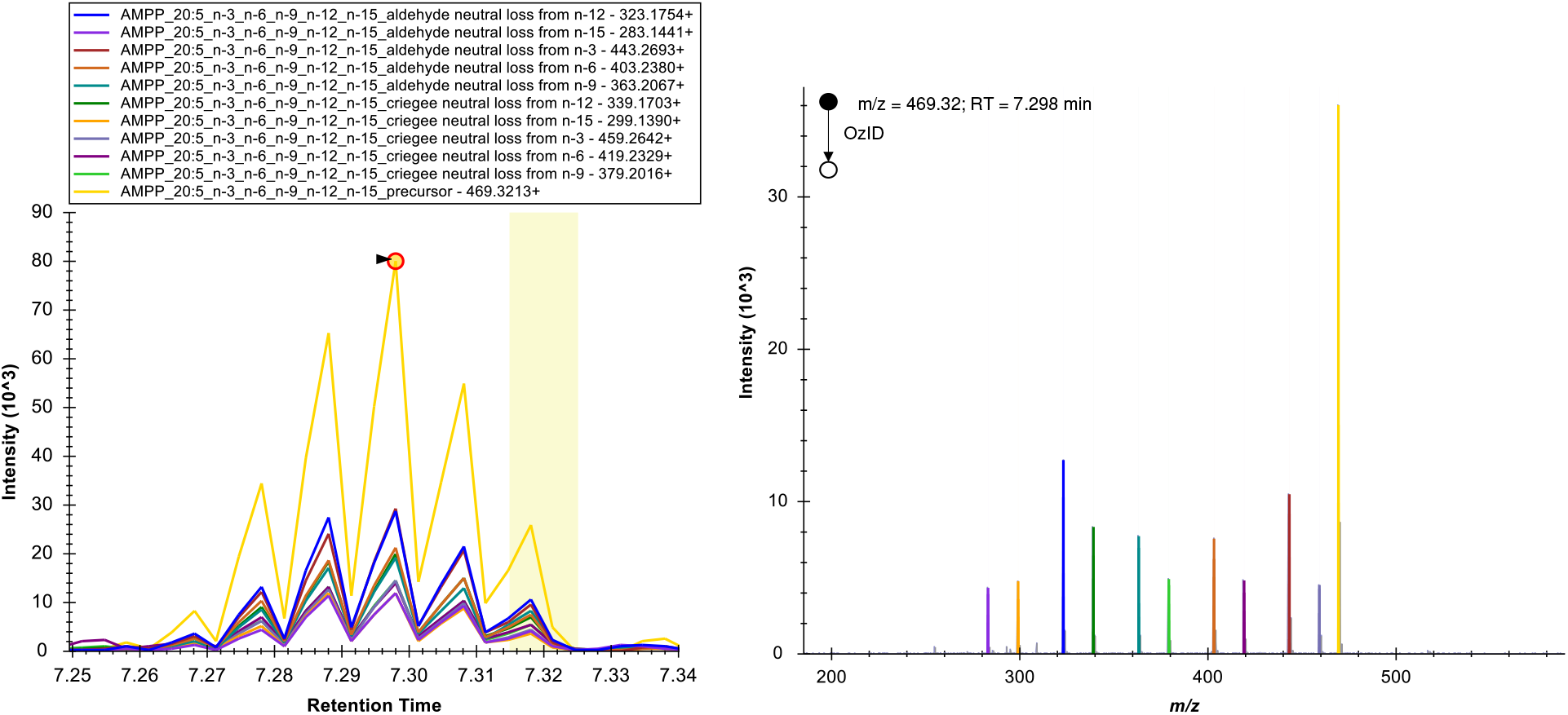
Chromatograms and mass spectra of the data dependent acquisition of the 37mix standard, showing FA 20:5*n*-3,6,9,12,15 (EPA).

**Figure S15:**
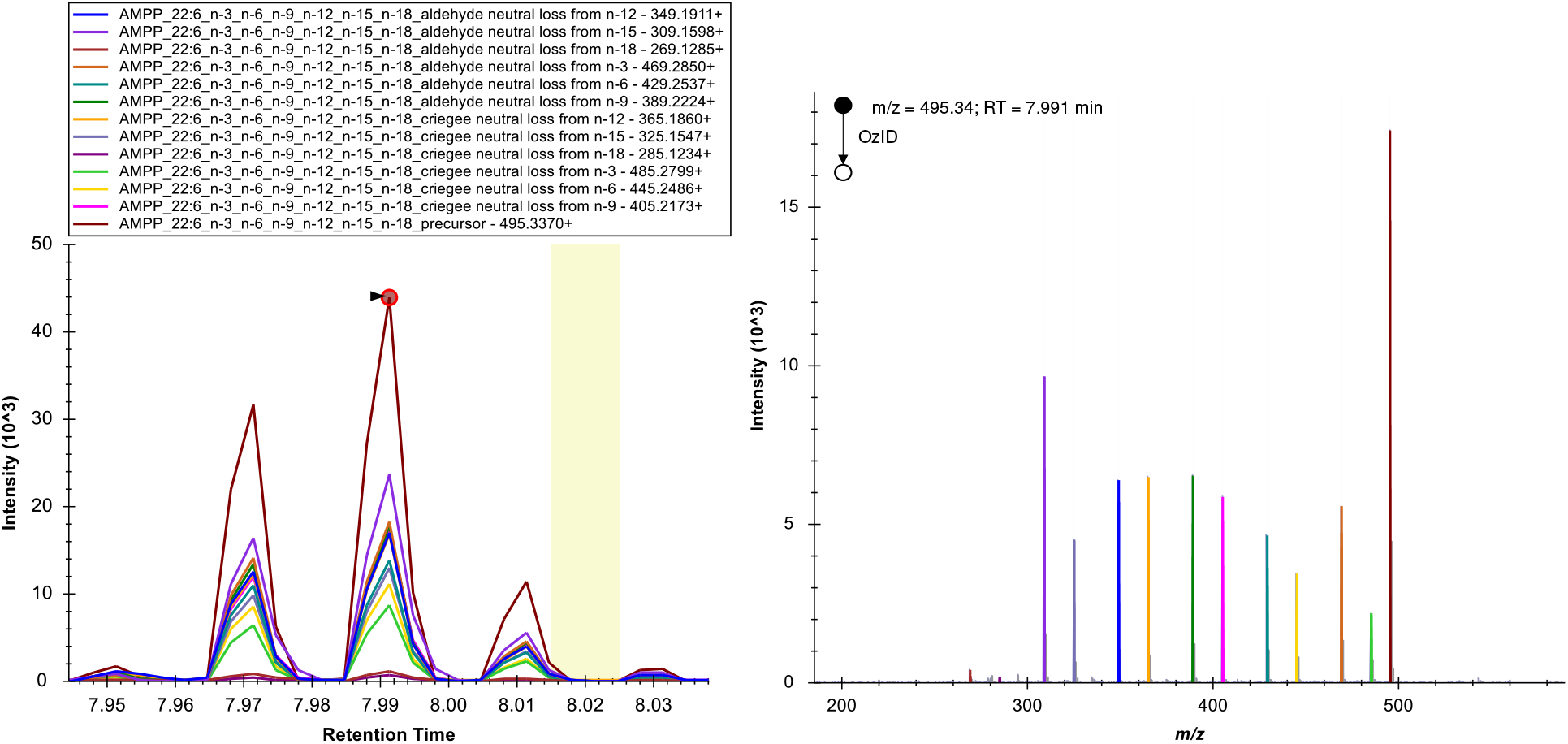
Chromatograms and mass spectra of the data dependent acquisition of the 37mix standard, showing FA 22:6*n*-3,6,9,12,15,18 (DHA). Note that ozone-induced dissociation from the Δ4 position (*n*-18) of the AMPP derivatized fatty acid appears to be drastically reduced in comparison to the other double bond positions.

**Table S3:** Degree of over-oxidation for monounsaturated fatty acids in the 37mix standard. The mean percentage of over-oxidation for monounsaturated cis fatty acids is 1.24, while the degree of over-oxidation is 2.84 for trans FA 18:1*n*-9. Values were calculated from the sums of OzID product ion peak integrals from the data dependent acquisition. Only fatty acid isomers were selected that featured an uninterrupted acquisition of the full peak.

| FA | Over-oxidation / % |
| --- | --- |
| FA 14:1n-5 | 1.51 |
| FA 15:1n-5 | 1.44 |
| FA 18:1n-9 | 1.22 |
| FA 18:1n-9t | 2.84 |
| FA 20:1n-9 | 1.16 |
| FA 22:1n-9 | 0.99 |
| FA 24:1n-9 | 1.13 |

All fatty acid isomer identifications in this work were validated to not be false positive discoveries based on the observed over-oxidation effect. For example, Fig. S13 shows the retention time shift between FA 19:1*n*-10 and FA 19:1*n*-11 in pooled human plasma, refer to Fig. 3 in the main paper, which evidences, apart from the observation of the ratio of the two fatty acids (I_C19:1n-1_)/(I_C19:1n-11_+I_C19:1n-10_) = 2%) that FA 19:1*n*-11 is a correctly identified isomer. Features that were excluded from the analysis as potential over-oxidation artefacts include FA 16:1*n*-8*cis*,^*1*^ FA 20:1*n*-10*cis*,^2^ and FA 24:1*n*-10*cis* in NIST 1950 human plasma.

**Figure S16:**
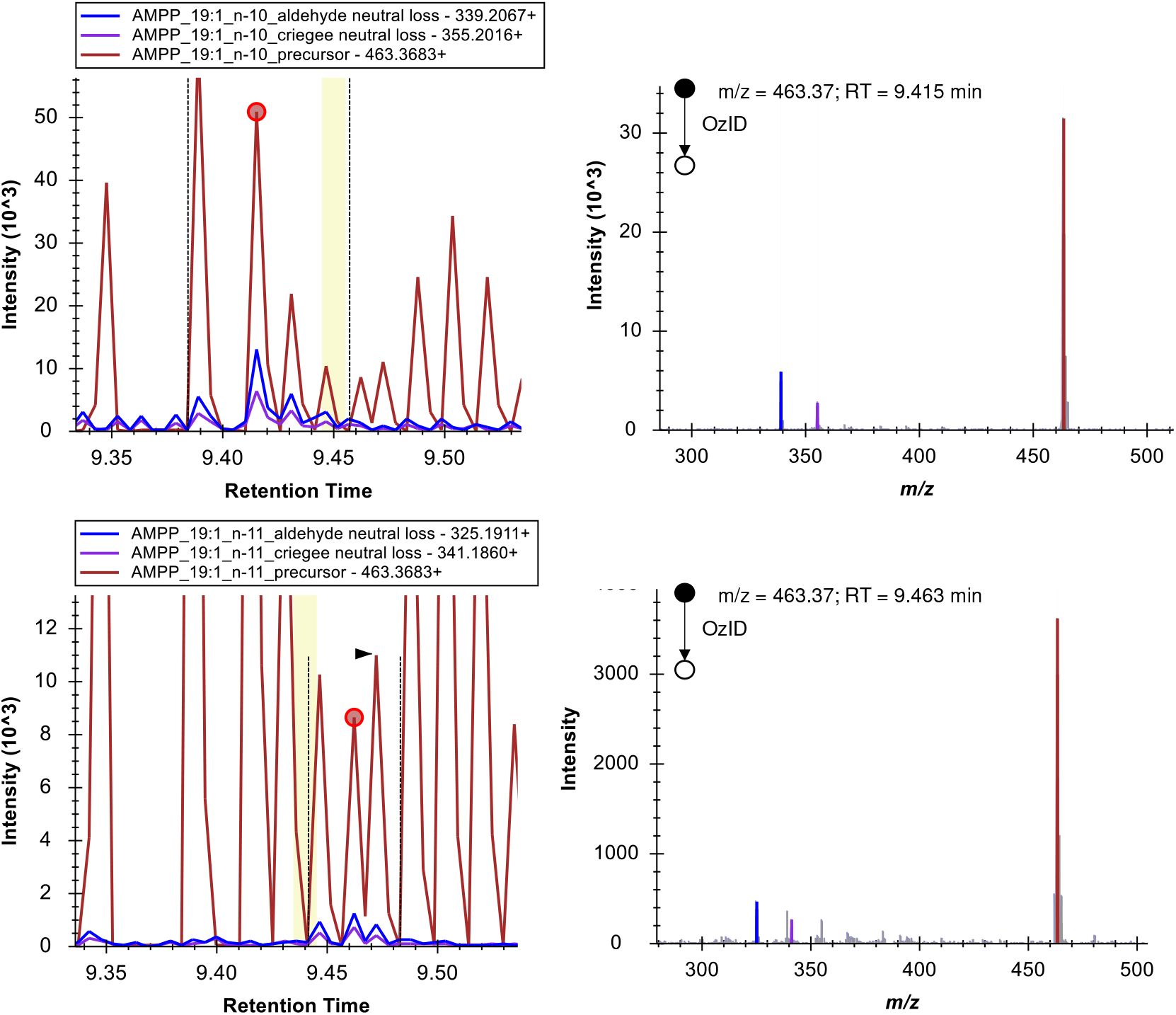
Extracted ion chromatograms (data dependent acquisition) and OzID-MS/MS spectra of FA 19:1*n*-10 (top chromatograms and associated MS/MS spectrum) and FA 19:1*n*-11 (bottom spectra) in NIST 1950 standard reference material (pooled human plasma).

**Figure S17:**
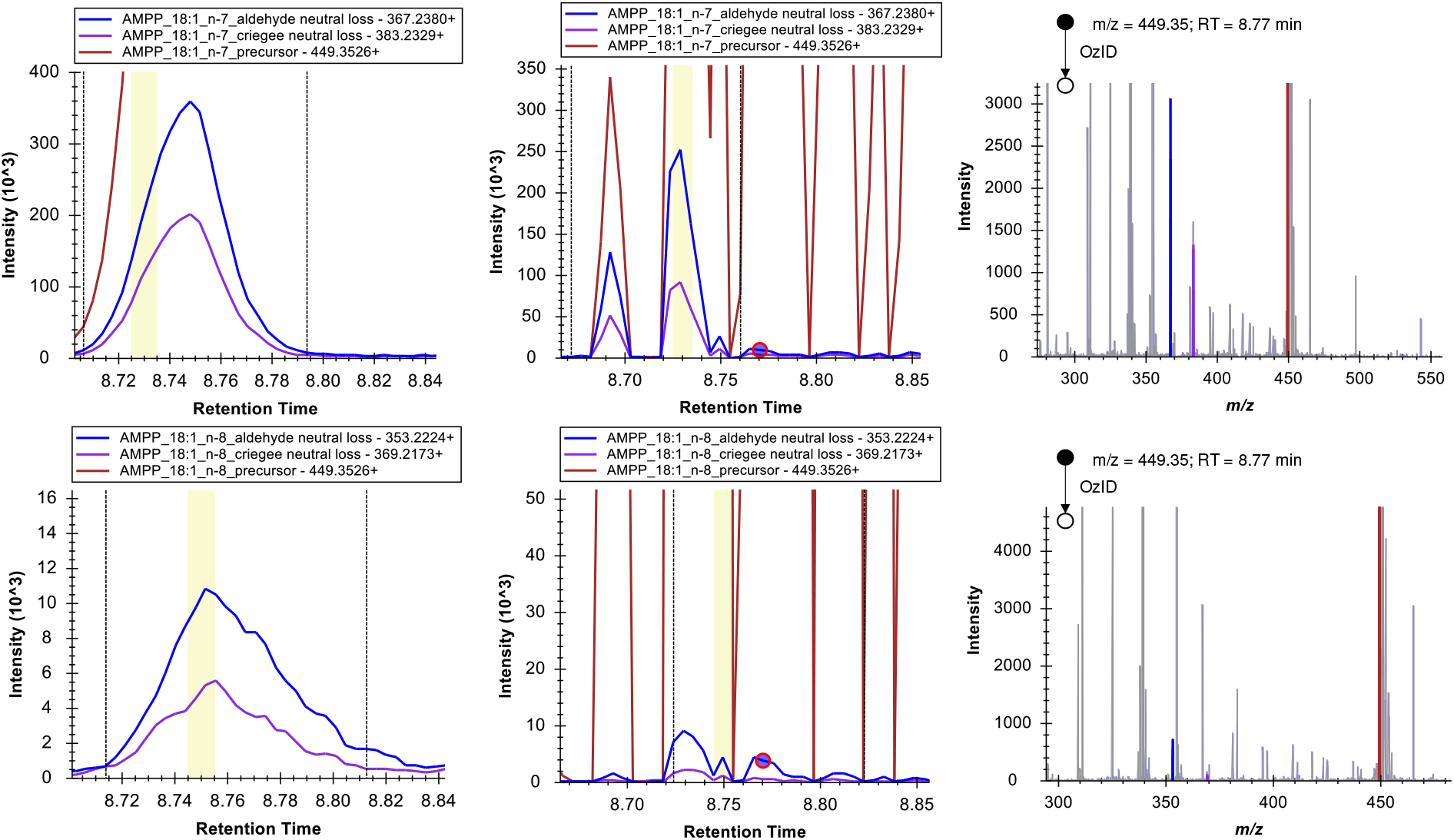
Extracted ion chromatograms (left: data independent acquisition; middle and right: data dependent acquisition) and OzID-MS/MS spectra of FA 18:1*n*-7*cis* (top chromatograms and associated MS/MS spectrum) and FA 18:1*n*-8*cis* (bottom spectra) in NIST 1950 standard reference material (pooled human plasma). The retention time shift and the ratio of OzID product ion abundance of FA 18:1*n*-8*cis* and FA 18:1*n*-7*cis* at 8.77 min show that the feature representing FA 18:1*n*-8*cis* is only partly caused by over-oxidation of FA 18:1*n*-7*cis*.

**Figure S18:**
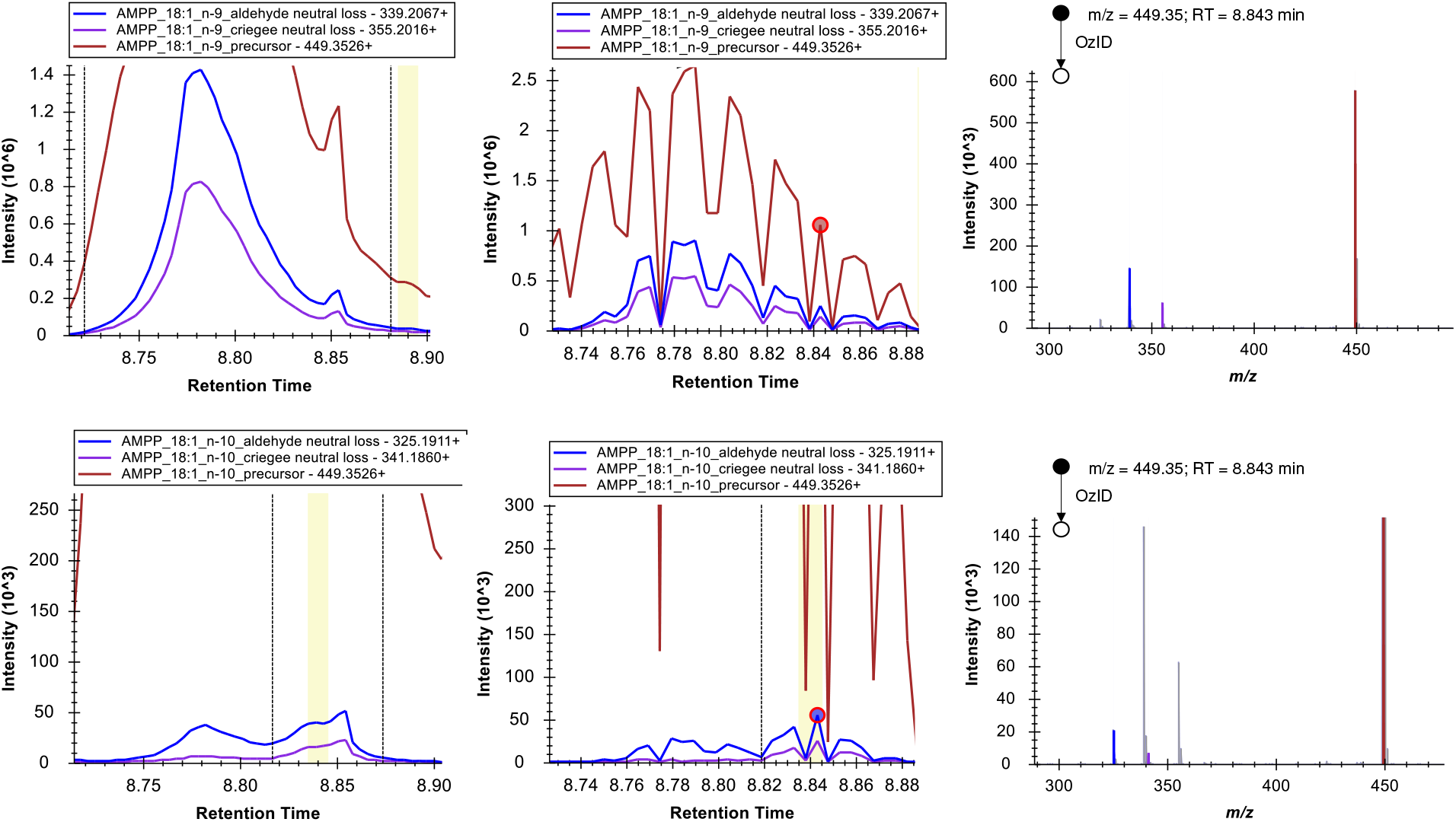
Extracted ion chromatograms (left: data independent acquisition; middle and right: data dependent acquisition) and OzID-MS/MS spectra of FA 18:1n-9*cis* (top chromatograms and associated MS/MS spectrum) and FA 18:1*n*-10*cis* (bottom spectra) in NIST 1950 standard reference material (pooled human plasma). The retention time shift and the ratio of OzID product ion abundance of FA 18:1*n*-9*cis* and FA 18:1*n*-10*cis* at 8.843 min show that the feature at RT = 8.85 min representing FA 18:1*n*-10*cis* is not caused by over-oxidation of FA 18:1*n*-9*cis*.

### 3.2 Process blank subtraction

Each lipid extraction and derivatization was accompanied by a process blank, a blank sample containing an equal amount of internal standard as the biological samples. The process blank is treated equally to the samples being taken through the workflow. In each process blank we observed a subset of small amounts of fatty acids, usually palmitic acid, stearic acid, erucic and oleic acid, among very minor amounts of other fatty acid species. We found that no method of cleaning glassware and tools could completely suppress the detection of remaining minor amounts of such fatty acids. Therefore, the respective Process blank is always acquired alongside the sample and a subtraction of the amounts of the species that are present at notable amounts relative to the biological sample is performed. One possible source of erucic acid appears to be the plastics additive erucamide,^3^ possibly introduced into the workflow through the use of plastic tips of Eppendorf pipettes. A characteristic peak at the m/z value of erucamide is observed in the Process Blank of for example human plasma, as well as a significant amount of the derivatized erucic acid. The latter is observed in the direct infusion, the LC-OzID-MS and LC-OzID-MS/MS acquisitions.

Regarding the analysis of human plasma, all docosenoic acids (22:1) were therefore excluded from the analysis. Further, the fatty acids FA 14:1*n*-8*cis*^4^ and FA 15:1*n*-9*cis*^4^ were excluded from the analysis, as similar amounts of these species were observed in the associated Process Blank. The relative amounts of several other species was adjusted by Process Blank subtraction. Example spectra for the Process Blank subtraction of Sapienic acid FA 16:1*n*-10*cis* are shown in Fig. S19.

**Figure S19:**
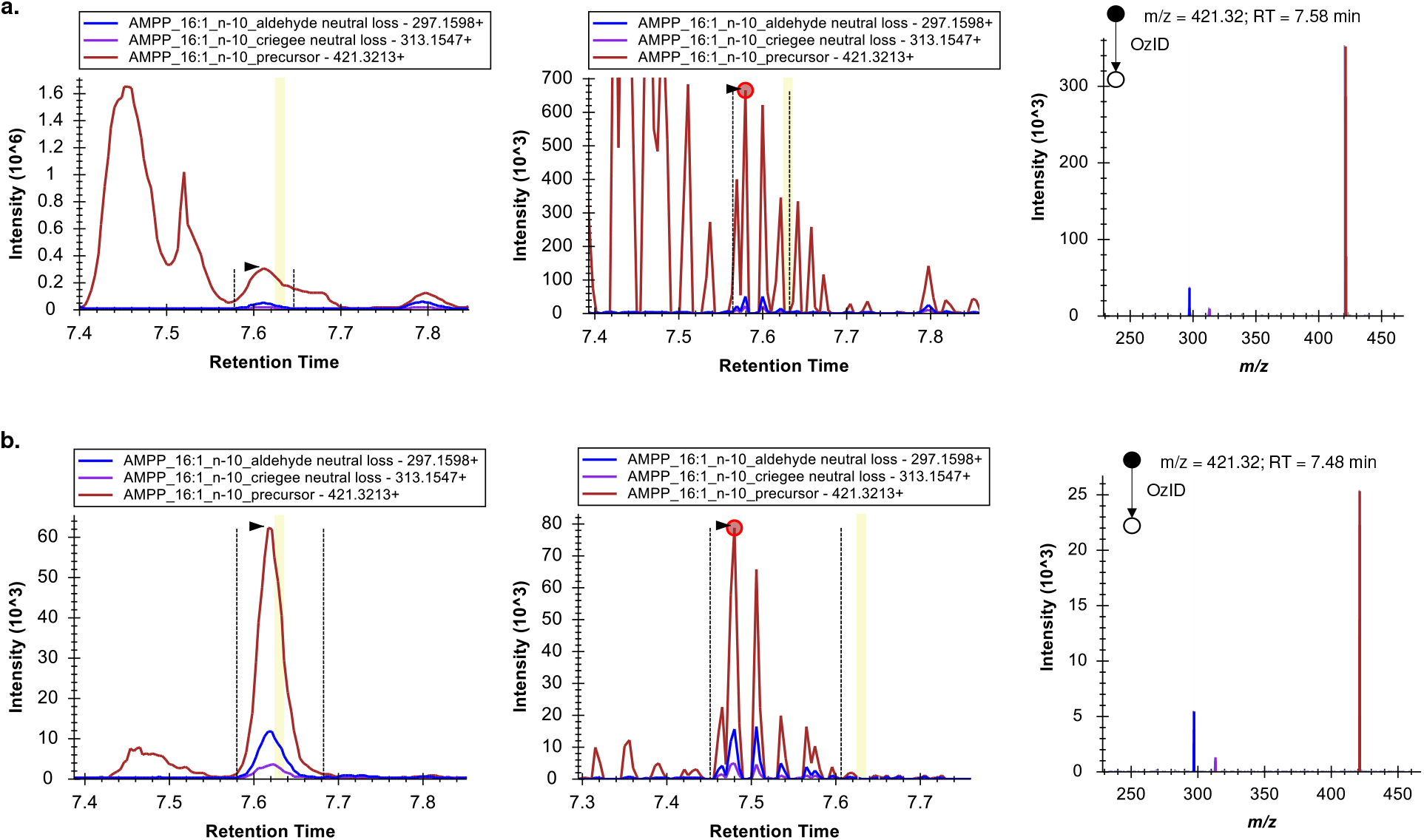
Comparison of Sapienic acid in (**a**.) pooled human plasma (NIST 1950 Standard Reference Material) and (**b**.) the associated Process Blank. For each sample, the chromatograms of the data independent acquisition are shown on the right, chromatograms of the data dependent acquisition in the middle and an associated MS/MS spectrum on the right (the red dot in the middle indicates the associated retention time). The intensity scale clearly indicates that the amount of hexadecenoic acids in the human plasma sample is far higher than in the Process Blank, whereas the intensity of the internal standard is nearly equal (omitted for clarity).

### 3.3 Fatty acids in hydrolyzed human plasma NIST1950 SRM

**Figure S20:**
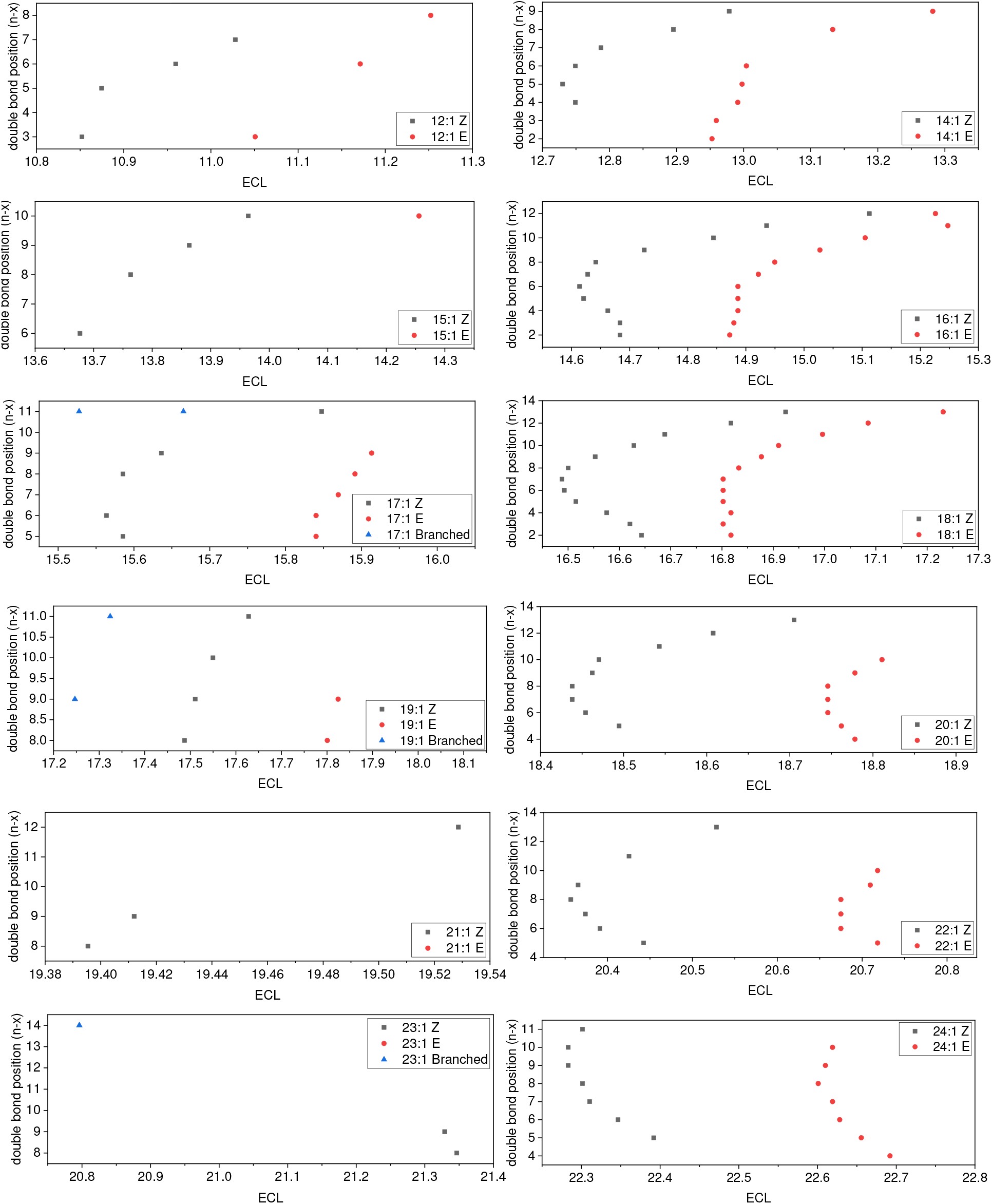
Equivalent chain lengths (ECL) of monounsaturated fatty acids detected by data dependent acquisition in NIST 1950 Standard Reference Material (Pooled Human Plasma). ECLs are calculated from observed retention times of the respective chromatographic peak of the OzID product chromatograms in the UPLC-OzID-MS/MS data. An expression was determined as a polynomial fit to the plot of the chain lengths of saturated fatty acids vs. their observed retention time in the same dataset as ECL = 0.0436 * (RT)^2 + 0.7393 * RT + 6.6932. For saturated fatty acids the equivalent chain length is defined as the number of carbon atoms in the fatty acyl chain, see also Figure S21. The equivalent chain lengths of monounsaturated fatty acids are used here to assign E/Z conformation and to assign branched vs. straight chain fatty acids.

**Figure S21:**
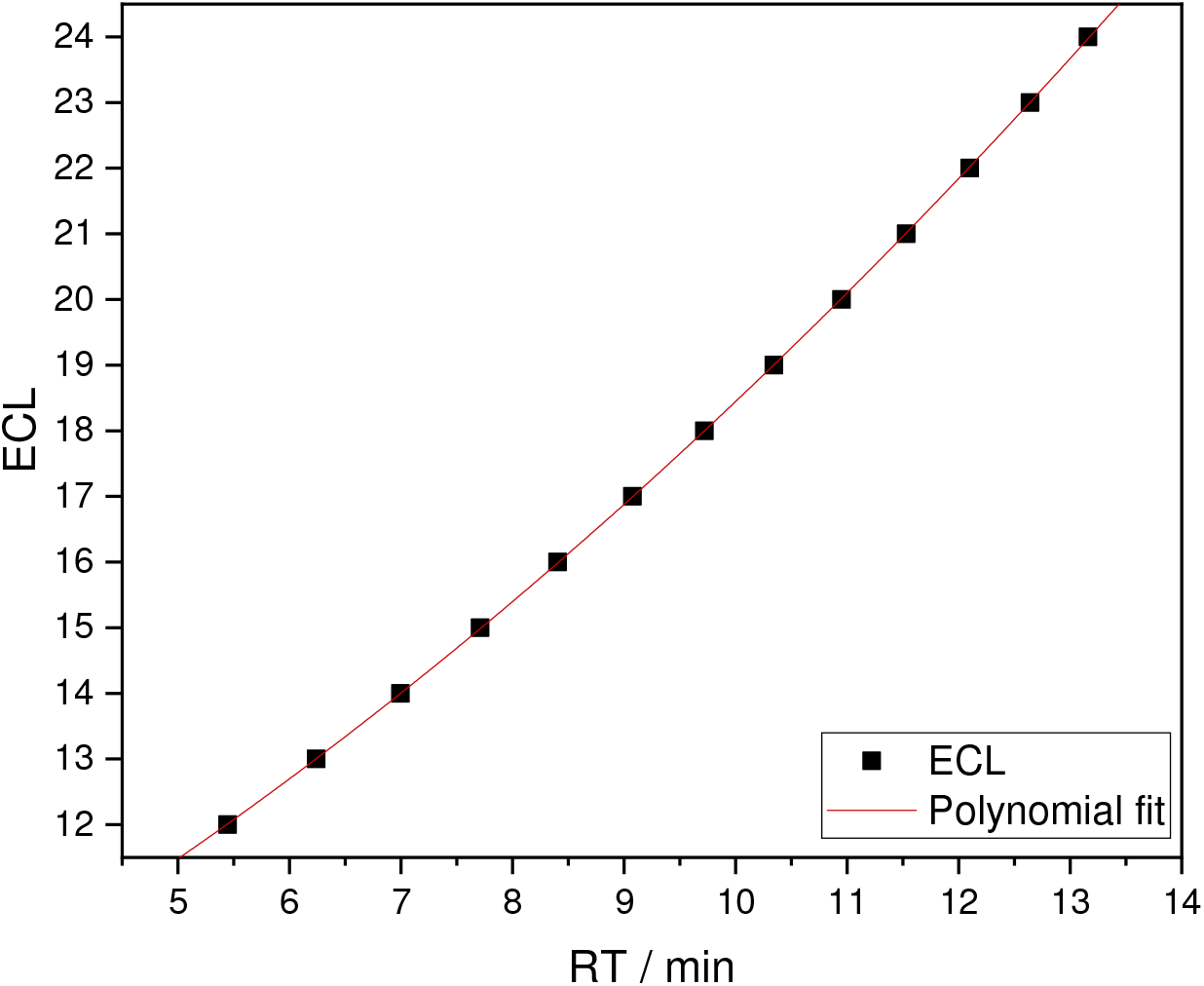
Polynomial fit of equivalent chain lengths vs. retention times of saturated fatty acids (straight chain) in NIST 1950 Standard Reference Material (Pooled Human Plasma), technical replicate 1 (ECL = 0.0436 * (RT)^2 + 0.7393 * RT + 6.6932).

**Figure S22:**
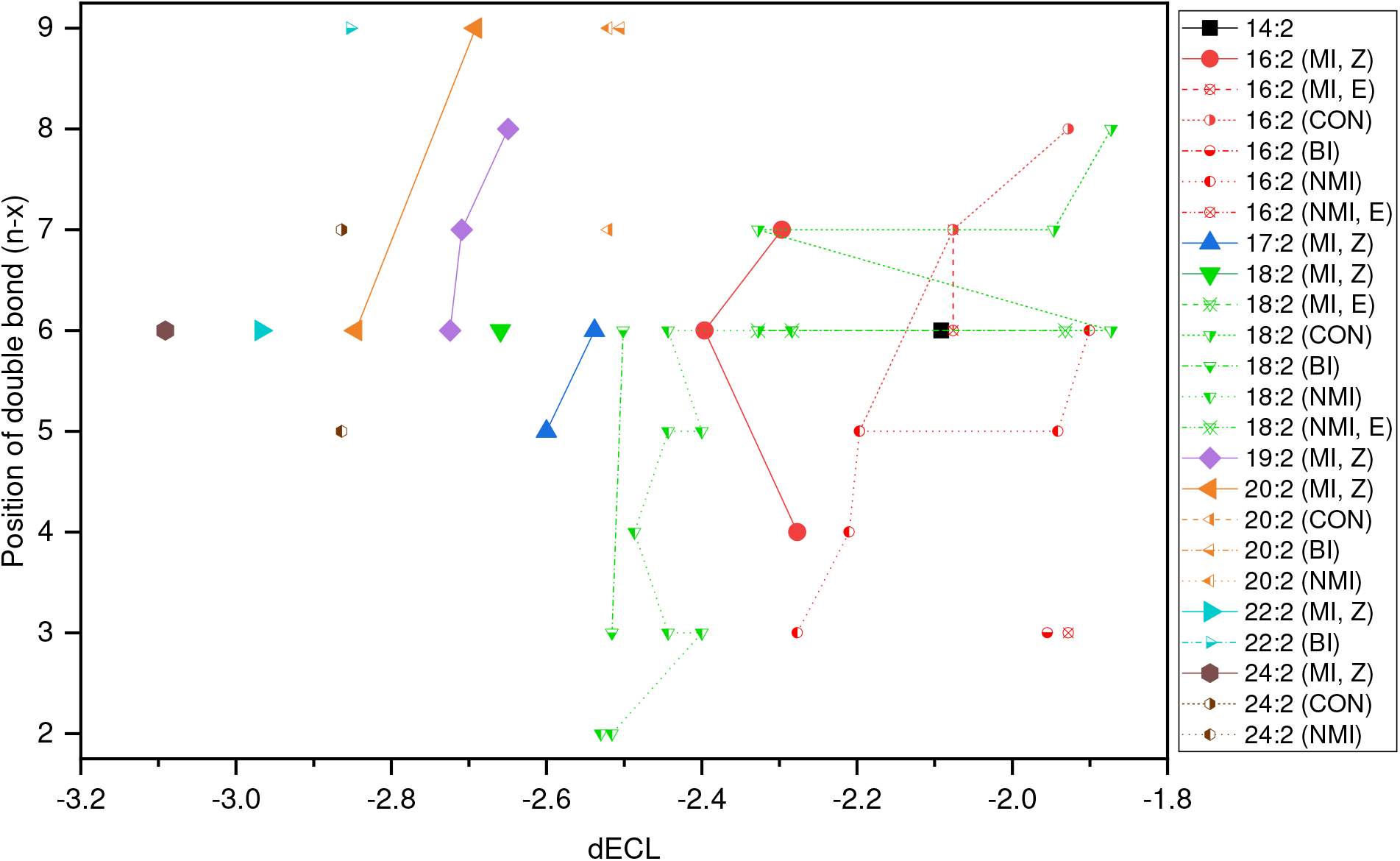
Analysis of differential equivalent chain lengths (dECL) for bisunsaturated fatty acids in NIST 1950 SRM. Canonical methylene-interrupted species (MI) are highlighted with larger symbols. Non-methylene interrupted fatty acids are assigned as conjugated (CON), butylene-interrupted (BI) or other non-methylene interrupted fatty acids (NMI). Double bond configurations cannot be established unequivocally for all fatty acid species. Note that some identifications of FA 18:2 and FA 16:2 isomers are made only tentatively. For determination of ECL and dECL values, the same equation that was used for monounsaturated fatty acids was employed here (ECL = 0.0436 * (RT)^2 + 0.7393 * RT + 6.6932).

**Figure S23:**
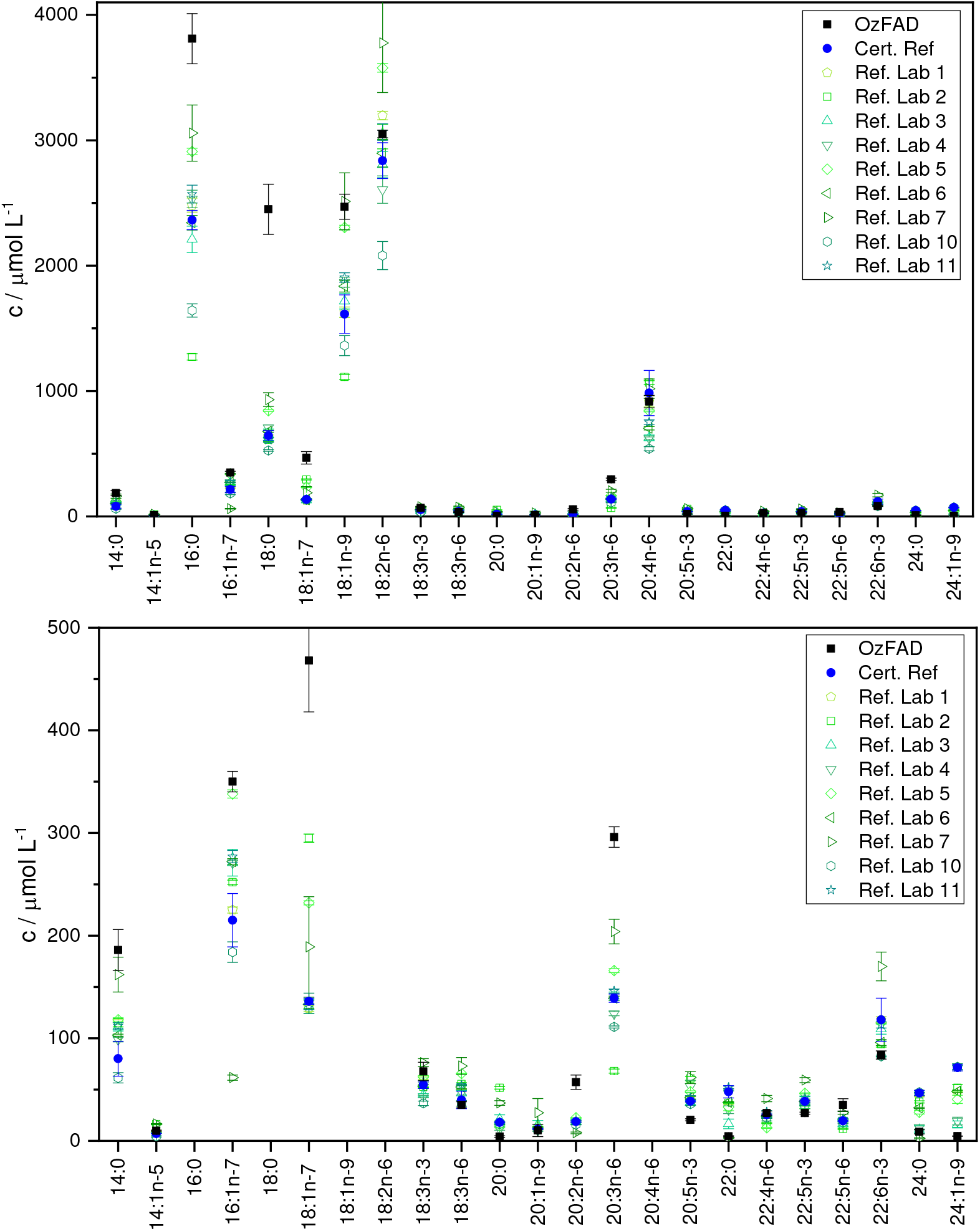
Comparison of quantities of fatty acids (total fatty acid content) in the NIST 1950 SRM quantified by the OzFAD workflow and values reported by the U.S. Centre for Disease Control and Prevention (CDC) based on an international interlaboratory analytical comparison study.^5^ The certified reference value is shown as blue filled circles and individual data from the participating laboratories in the interlaboratory study are shown as green open symbols. The two graphs show the same data scaled differently to allow visualization of fatty acids at high abundance (top) and low abundance (bottom).

The mean average accuracy of the fatty acids shown in Fig. S23 (mean of the percentage of the absolute deviation of the obtained quantity from the certified reference value) is 78% (63% for the unsaturated fatty acids shown). The same calculation for the reference laboratories within the study (mean average of the mean of the percentage of the absolute deviation of the reported quantity from the certified reference value) is 27% (25% for the unsaturated fatty acids shown). This shows that the data obtained through the OzFAD workflow can – without requiring libraries or reference standards – provide best estimates of fatty acid quantities, with the limitation of less accurate quantities as the ones that may be provided by laboratories specializing on the quantification of fatty acids (using fatty acid standards to perform quantification). Highly accurate quantification is outside the scope of the current work, which is focused on the discovery of novel isomers and completing fatty acid profiles in complex samples.

**Table S4:** Overview of fatty acids that are identified here in NIST1950 Standard Reference Material “Frozen Metabolites in Pooled Human Blood Plasma”. Shown are fatty acids detected in the total fatty acid fraction of the human plasma after lipid extraction, hydrolysis, fixed-charge derivatization and UPLC-OzID-MS (& MS/MS) analysis. Each fatty acid isomer is described according to the n nomenclature (column 1), where the double bond position is indicated as counted from the methyl end, according to the fatty acid shorthand and the systematic name, where the double bond position is indicated as counted from the carboxy end of the fatty acid. For the purpose of the automated database search, systematic names of polyunsaturated fatty acids are generated with cis double bonds only. If available at time of analysis (February 2022), the LipidMAPS ID with a link to the LipidMAPS webpage is included in column 4 as well as a common name (column 5). Retention times are rounded to three significant digits, and the relative isomer abundance is shown as a percentage of all fatty acid isomers with the same precursor mass. Signal-to-noise ratios are calculated from multiple selected and combined MS/MS spectra of each fatty acid isomer. The confidence of identification is high (H), when the double bond position can be unequivocally determined and low (L), when it can’t be determined, which exact isomer is present. For fatty acid isomers with low confidence of identification, only the presence of certain double bond positions can be ascertained, but not the identity (association of two double bond positions to one specific isomer) of the many coeluting bisunsaturated fatty acid isomers. The absolute quantification is shown in μmol L^-1^ of pooled human plasma. If the fatty acid isomer was identified in the literature (to the best of our knowledge), one reference is given in column 11. If the fatty acid was not identified in human plasma previously, but elsewhere, then a reference is given in column 12. The references are not exhaustive indications as to where the fatty acids have been reported, nor do the references necessarily show the first mention in the literature of the respective fatty acid isomer. If a fatty acid was not found in the literature at all, no reference is shown here, and the column is highlighted with a yellow background. For a few of these fatty acids, we propose common names, indicated in column 5 (italic font).

| Fatty acid (n-x) | FA shorthand ('FA' is omitted for clarity) | Systematic name | LipidMAPS ID | Common Name | RT / min | Relative isomer abundance / % | COV Rel. abundance | S/N OzID MS/MS | Confidence of identification (Hig h; Low: L) | Estim. abs. quantification / $\mu\text{mol L}^{-1}$ | Found in Human Plasma | Found elsewhere, if not in Human Plasma |
| --- | --- | --- | --- | --- | --- | --- | --- | --- | --- | --- | --- | --- |
| C12:0 | 12:0 | Dodecanoic acid | LMFA01010012 | Lauric acid | 5.45 | n.a. | - | - | H | 15 ± 2 | Y<sup>6</sup> | |
| C13:0 | 13:0 | Tridecanoic acid | LMFA01010013 | Tridecylic acid | 6.24 | n.a. | - | - | H | 3 ± 2 | Y<sup>7</sup> | |
| C14:0 | 14:0 | Tetradecanoic acid | LMFA01010014 | Myristic acid | 7.00 | n.a. | - | - | H | 190 ± 20 | Y<sup>6</sup> | |
| C15:0 | 15:0 | Pentadecanoic acid | LMFA01010015 | Pentadecylic acid | 7.71 | n.a. | - | - | H | 93 ± 8 | Y<sup>6</sup> | |
| C16:0 | 16:0 | Hexadecanoic acid | LMFA01010001 | Palmitic acid | 8.41 | n.a. | - | - | H | 3800 ± 200 | Y<sup>6</sup> | |
| C17:0 | 17:0 | Heptadecanoic acid | LMFA01010017 | Margaric acid | 9.08 | n.a. | - | - | H | 87 ± 5 | Y<sup>6</sup> | |
| C18:0 | 18:0 | Octadecanoic acid | LMFA01010041 | Stearic acid | 9.72 | n.a. | - | - | H | 2500 ± 200 | Y<sup>6</sup> | |
| C19:0 | 19:0 | Nonadecanoic acid | LMFA01010019 | Nonadecylic acid | 10.35 | n.a. | - | - | H | 2.0 ± 0.5 | | Y<sup>4</sup> |
| C20:0 | 20:0 | Eicosanoic acid | LMFA01010020 | Arachidic acid | 10.95 | n.a. | - | - | H | 4.1 ± 0.8 | Y<sup>6</sup> | |
| C21:0 | 21:0 | Heneicosanoic acid | LMFA01010021 | Heneicosylic acid | 11.53 | n.a. | - | - | H | 0.21 ± 0.05 | Y<sup>8</sup> | |
| C22:0 | 22:0 | Docosanoic acid | LMFA01010022 | Behenic acid | 12.10 | n.a. | - | - | H | 4.5 ± 0.5 | Y<sup>6</sup> | |
| C23:0 | 23:0 | Tricosanoic acid | LMFA01010023 | Tricosylic acid | 12.64 | n.a. | - | - | H | 1.9 ± 0.3 | Y<sup>6</sup> | |
| C24:0 | 24:0 | Tetracosanoic acid | LMFA01010024 | Lignoceric acid | 13.16 | n.a. | - | - | H | 9 ± 2 | Y<sup>6</sup> | |
| C12:1n-3 | 12:1(9Z) | 9Z-dodecenoic acid | LMFA01030229 | Lauroleic acid | 4.45 | 18 ± 5 | 0.3 | 115 | H | 0.4 ± 0.2 | | Y<sup>9</sup> |
| C12:1n-5 | 12:1(7Z) | 7Z-dodecenoic acid | LMFA01030228 | 7Z-dodecenoic acid | 4.46 | 2.3 ± 0.6 | 0.3 | 11 | H | 0.07 ± 0.06 | - | Y<sup>10</sup> |
| C12:1n-6 | 12:1(6Z) | 6Z-dodecenoic acid | LMFA01030227 | 6Z-dodecenoic acid | 4.54 | 26 ± 4 | 0.2 | 199 | H | 0.8 ± 0.7 | - | - |
| C12:1n-7 | 12:1(5Z) | 5Z-dodecenoic acid | LMFA01030226 | Lauroleic acid | 4.59 | 30 ± 2 | 0.07 | 254 | H | 0.8 ± 0.6 | Y<sup>11</sup> | |
| C12:1n-3 (E) | 12:1(9E) | 9E-dodecenoic acid | Not found in LIPID MAPS. | - | 4.62 | 13 ± 2 | 0.2 | 139 | H | 0.4 ± 0.3 | | Y<sup>12</sup> |
| C12:1n-6 (E) | 12:1(6E) | 6E-dodecenoic acid | Not found in LIPID MAPS. | - | 4.73 | 6 ± 1 | 0.2 | 44 | H | 0.2 ± 0.1 | - | - |
| C12:1n-8 (E) | 12:1(4E) | 4E-dodecenoic acid | Not found in LIPID MAPS. | - | 4.81 | 5 ± 2 | 0.4 | 29 | H | 0.13 ± 0.09 | - | Y<sup>13</sup> |
| C14:1n-5 | 14:1(9Z) | 9Z-tetradecenoic acid | LMFA01030051 | Myristoleic acid | 6.02 | 69 ± 6 | 0.09 | 2818 | H | 9.9 ± 0.5 | Y<sup>14</sup> | |
| C14:1n-6 | 14:1(8Z) | 8Z-tetradecenoic acid | LMFA01030050 | 8Z-tetradecenoic acid | 6.02 | 7 ± 5 | 0.7 | 59 | H | 1.0 ± 0.7 | - | - |
| C14:1n-4 | 14:1(10Z) | 10Z-tetradecenoic acid | Not found in LIPID MAPS. | - | 6.03 | 0.5 ± 0.5 | 0.9 | 30 | H | 0.08 ± 0.07 | - | - |
| C14:1n-7 | 14:1(7Z) | 7Z-tetradecenoic acid | LMFA01030249 | 7Z-tetradecenoic acid | 6.05 | 4 ± 2 | 0.5 | 296 | H | 0.5 ± 0.3 | Y<sup>2</sup> |  |
| C14:1n-3_(E) | 14:1(11E) | 11E-tetradecenoic acid | LMFA01030771 | Trans-tetradec-11-enoic acid | 6.21 | 1.8 ± 0.3 | 0.2 | 182 | H | 0.26 ± 0.05 |  | Y<sup>15</sup> |
| C14:1n-2_(E) | 14:1(12E) | 12E-tetradecenoic acid | Not found in LIPID MAPS. | - | 6.2 | 1.4 ± 0.6 | 0.4 | 36 | H | 0.2 ± 0.09 | - | - |
| C14:1n-9 | 14:1(5Z) | 5Z-tetradecenoic acid | LMFA01030248 | Physeteric acid | 6.21 | 8 ± 1 | 0.1 | 666 | H | 1.1 ± 0.2 | Y<sup>11</sup> | Y<sup>12</sup> |
| C14:1n-4_(E) | 14:1(10E) | 10E-tetradecenoic acid | Not found in LIPID MAPS. | - | 6.21 | 0.3 ± 0.1 | 0.3 | 28 | H | 0.05 ± 0.02 | - | - |
| C14:1n-5_(E) | 14:1(9E) | 9E-tetradecenoic acid | LMFA01030250 | Myristelaidic acid | 6.21 | 0.9 ± 0.2 | 0.2 | 31 | H | 0.12 ± 0.04 |  | Y<sup>12</sup> |
| C14:1n-6_(E) | 14:1(8E) | 8E-tetradecenoic acid | Not found in LIPID MAPS. | - | 6.23 | 1.2 ± 0.7 | 0.6 | 23 | H | 0.2 ± 0.1 | - | - |
| C14:1n-8_(E) | 14:1(6E) | 6E-tetradecenoic acid | Not found in LIPID MAPS. | - | 6.34 | 4.0 ± 0.9 | 0.2 | 690 | H | 0.6 ± 0.1 | - | - |
| C14:1n-9_(E) | 14:1(5E) | 5E-tetradecenoic acid | LMFA01030913 | - | 6.44 | 2 ± 2 | 0.9 | 73 | H | 0.3 ± 0.2 | - | - |
| C15:1n-6 | 15:1(9Z) | 9Z-pentadecenoic acid | LMFA01030866 | 15:1(9Z) | 6.75 | 24 ± 7 | 0.3 | 146 | H | 0.68 ± 0.09 |  | Y<sup>16</sup> |
| C15:1n-8 | 15:1(7Z) | 7Z-pentadecenoic acid | LMFA01030855 | 15:1(7Z) | 6.81 | 7 ± 3 | 0.5 | 67 | H | 0.2 ± 0.1 |  | Y<sup>16</sup> |
| C15:1n-10 | 15:1(5Z) | 5Z-pentadecenoic acid | LMFA01031197 | 5Z-pentadecenoic acid | 6.97 | 30 ± 10 | 0.4 | 72 | H | 0.9 ± 0.4 |  | Y<sup>12</sup> |
| C15:1n-10_(E) | 15:1(5E) | 5E-pentadecenoic acid | Not found in LIPID MAPS. | - | 7.19 | 41 ± 7 | 0.2 | 237 | H | 1.17 ± 0.07 | - | - |
| C16:1n-4 | 16:1(12Z) | 12Z-hexadecenoic acid | Not found in LIPID MAPS. | - | 7.45 | 0.5 ± 0.1 | 0.2 | 69 | H | 3.1 ± 0.7 | - | - |
| C16:1n-5 | 16:1(11Z) | 11Z-hexadecenoic acid | LMFA01030262 | cis-Palmitvaccenic acid | 7.45 | 1.3 ± 0.2 | 0.2 | 84 | H | 8 ± 1 | Y<sup>2</sup> |  |
| C16:1n-6 | 16:1(10Z) | 10Z-hexadecenoic acid | LMFA01030058 | cis-10-palmitoleic acid | 7.45 | 0.6 ± 0.2 | 0.3 | 37 | H | 4 ± 1 |  | Y<sup>1</sup> |
| C16:1n-7 | 16:1(9Z) | 9Z-hexadecenoic acid | LMFA01030056 | cis-9-palmitoleic acid | 7.45 | 58 ± 2 | 0.03 | 3212 | H | 350 ± 10 | Y<sup>6</sup> |  |
| C16:1n-9 | 16:1(7Z) | 7Z-hexadecenoic acid | LMFA01030766 |  | 7.52 | 20 ± 1 | 0.05 | 4275 | H | 120 ± 10 | Y<sup>2</sup> |  |
| C16:1n-2 | 16:1(14Z) | 14Z-hexadecenoic acid | Not found in LIPID MAPS. | - | 7.49 | 0.09 ± 0.03 | 0.3 | 25 | H | 0.6 ± 0.2 | - | - |
| C16:1n-3 | 16:1(13Z) | 13Z-hexadecenoic acid | LMFA01030264 | 13Z-hexadecenoic acid | 7.46 | 0.36 ± 0.09 | 0.3 | 147 | H | 2.2 ± 0.6 |  | Y<sup>1</sup> |
| C16:1n-2_(E) | 16:1(14E) | 14E-hexadecenoic acid | Not found in LIPID MAPS. | - | 7.61 | 0.7 ± 0.1 | 0.1 | 398 | H | 4.3 ± 0.6 | - | - |
| C16:1n-3_(E) | 16:1(13E) | 13E-hexadecenoic acid | Not found in LIPID MAPS. | - | 7.61 | 0.7 ± 0.2 | 0.3 | 418 | H | 4 ± 1 | - | - |
| C16:1n-4_(E) | 16:1(12E) | 12E-hexadecenoic acid | Not found in LIPID MAPS. | - | 7.61 | 1.1 ± 0.1 | 0.09 | 763 | H | 7 ± 1 | - | - |
| C16:1n-5_(E) | 16:1(11E) | 11E-hexadecenoic acid | LMFA01030261 | Lycopodic acid | 7.61 | 1.1 ± 0.2 | 0.2 | 707 | H | 6 ± 1 |  | Y<sup>12</sup> |
| C16:1n-6_(E) | 16:1(10E) | 10E-hexadecenoic acid | Not found in LIPID MAPS. | - | 7.62 | 1.1 ± 0.1 | 0.09 | 833 | H | 7 ± 1 |  | Y<sup>12</sup> |
| C16:1n-10 | 16:1(6Z) | 6Z-hexadecenoic acid | LMFA01030267 | Sapienic acid | 7.61 | 5.7 ± 0.6 | 0.1 | 2443 | H | 35 ± 5 | Y<sup>17</sup> |  |
| C16:1n-11 | 16:1(5Z) | 5Z-hexadecenoic acid | LMFA01030854 | 16:1(5Z) | 7.65 | 0.3 ± 0.1 | 0.4 | 85 | H | 1.7 ± 0.6 |  | Y<sup>18</sup> |
| C16:1n-7_(E) | 16:1(9E) | 9E-hexadecenoic acid | LMFA01030057 | trans-9-palmitoleic acid | 7.63 | 1.9 ± 0.2 | 0.1 | 1709 | H | 11.3 ± 0.6 | Y<sup>14</sup> |  |
| C16:1n-8_(E) | 16:1(8E) | 8E-hexadecenoic acid | LMFA01031190 | 8E-hexadecenoic acid | 7.65 | 1.7 ± 0.3 | 0.2 | 1992 | H | 10 ± 2 |  | Y<sup>12</sup> |
| C16:1n-9_(E) | 16:1(7E) | 7E-hexadecenoic acid | LMFA01030808 |  | 7.77 | 0.49 ± 0.05 | 0.1 | 886 | H | 3.0 ± 0.4 |  | Y<sup>19</sup> |
| C16:1n-10_(E) | 16:1(6E) | 6E-hexadecenoic acid | LMFA01031082 | 6E-Hexadecenoic acid | 7.79 | 4.1 ± 0.4 | 0.1 | 4257 | H | 25 ± 2 | Y<sup>17</sup> |  |
| C16:1n-12 | 16:1(4Z) | 4Z-hexadecenoic acid | Not found in LIPID MAPS. | - | 7.79 | 0.04 ± 0.01 | 0.2 | 25 | H | 0.26 ± 0.08 |  | Y<sup>20</sup> |
| C16:1n-11_(E) | 16:1(5E) | 5E-hexadecenoic acid | Not found in LIPID MAPS. | - | 7.86 | 0.58 ± 0.09 | 0.2 | 850 | H | 3.5 ± 0.4 | - | - |
| C16:1n-12_(E) | 16:1(4E) | 4E-hexadecenoic acid | Not found in LIPID MAPS. | - | 7.86 | 0.017 ± 0.006 | 0.4 | 41 | H | 0.11 ± 0.04 | - | - |
| C17:1n-6 | 17:1(11Z) | 11Z-heptadecenoic acid | LMFA01030856 | 17:1(11Z) | 8.12 | 4 ± 2 | 0.5 | 96 | H | 1.1 ± 0.6 | Y<sup>2</sup> |  |
| C17:1n-11_br | 17:1(6)_br |  |  |  | 8.1 | 7 ± 1 | 0.2 | 57 | H | 1.9 ± 0.4 |  |  |
| C17:1n-5 | 17:1(12Z) | 12Z-heptadecenoic acid | LMFA01030907 | 17:1(12Z) | 8.13 | 1 ± 1 | 0.7 | 15 | H | 0.4 ± 0.4 |  | Y<sup>21</sup> |
| C17:1n-8 | 17:1(9Z) | 9Z-heptadecenoic acid | LMFA01030060 | Margaroleic acid | 8.13 | 73 ± 8 | 0.1 | 3787 | H | 21 ± 1 | Y<sup>2</sup> |  |
| C17:1n-9 | 17:1(8Z) | 8Z-heptadecenoic acid | LMFA01030288 | Civetac acid | 8.15 | 6 ± 2 | 0.3 | 409 | H | 1.7 ± 0.7 | Y<sup>2</sup> |  |
| C17:1n-11_br | 17:1(6)_br |  |  |  | 8.17 | 7 ± 1 | 0.2 | 70 | L | 1.9 ± 0.4 |  |  |
| C17:1n-5_(E) | 17:1(12E) | 12E-heptadecenoic acid | Not found in LIPID MAPS. | - | 8.3 | 0.6 ± 0.2 | 0.3 | 17 | H | 0.18 ± 0.07 | - | - |
| C17:1n-6_(E) | 17:1(11E) | 11E-heptadecenoic acid | Not found in LIPID MAPS. | — | 8.3 | 0.6 ± 0.3 | 0.5 | 21 | H | 0.16 ± 0.09 | - | - |
| C17:1n-7_(E) | 17:1(10E) | 10E-heptadecenoic acid | LMFA01030282 | 10E-heptadecenoic acid | 8.31 | 0.7 ± 0.07 | 0.1 | 48 | H | 0.21 ± 0.03 | - | - |
| C17:1n-8_(E) | 17:1(9E) | 9E-heptadecenoic acid | LMFA01030289 | 9E-heptadecenoic acid | 8.32 | 2 ± 1 | 0.6 | 53 | H | 0.5 ± 0.3 | - | - |
| C17:1n-9_(E) | 17:1(8E) | 8E-heptadecenoic acid | LMFA01030287 | 8E-heptadecenoic acid | 8.34 | 1 ± 1 | 0.9 | 13 | H | 0.3 ± 0.4 | - | - |
| C17:1n-11 | 17:1(6Z) | 6Z-heptadecenoic acid | LMFA01031186 | — | 8.3 | 4.6 ± 0.5 | 0.1 | 89 | L | 1.3 ± 0.2 |  | Y<sup>4</sup> |
| C18:1n-6 | 18:1(12Z) | 12Z-octadecenoic acid | LMFA01030078 | cis-12-oleic acid | 8.77 | 2.0 ± 0.3 | 0.2 | 878 | H | 80 ± 10 |  | Y<sup>12</sup> |
| C18:1n-7 | 18:1(11Z) | 11Z-octadecenoic acid | LMFA01030076 | cis-vaccenic acid | 8.77 | 11.5 ± 0.9 | 0.08 | 4440 | H | 470 ± 50 | Y<sup>14</sup> |  |
| C18:1n-8 | 18:1(10Z) | 10Z-octadecenoic acid | LMFA01030074 | cis-10-oleic acid | 8.79 | 0.3 ± 0.1 | 0.4 | 59 | H | 11 ± 5 |  | Y<sup>1</sup> |
| C18:1n-3 | 18:1(15Z) | 15Z-octadecenoic acid | LMFA01030291 | 15Z-octadecenoic acid | 8.8 | 0.21 ± 0.07 | 0.3 | 93 | H | 9 ± 3 |  | Y<sup>12</sup> |
| C18:1n-4 | 18:1(14Z) | 14Z-octadecenoic acid | LMFA01030882 | 18:1(14Z) | 8.79 | 0.07 ± 0.01 | 0.1 | 6 | H | 2.9 ± 0.5 |  | Y<sup>12</sup> |
| C18:1n-5 | 18:1(13Z) | 13Z-octadecenoic acid | LMFA01030290 | 13Z-octadecenoic acid | 8.79 | 0.24 ± 0.04 | 0.2 | 32 | H | 10 ± 2 |  | Y<sup>12</sup> |
| C18:1n-10 | 18:1(8Z) | 8Z-octadecenoic acid | LMFA01030070 | cis-8-oleic acid | 8.81 | 0.69 ± 0.05 | 0.07 | 675 | H | 28 ± 2 | Y<sup>2</sup> |  |
| C18:1n-2 | 18:1(16Z) | 16Z-octadecenoic acid | LMFA01030292 | 16Z-octadecenoic acid | 8.83 | 0.2 ± 0.2 | 1.0 | 65 | H | 9 ± 7 | - | - |
| C18:1n-5_(E) | 18:1(13E) | 13E-octadecenoic acid | LMFA01030880 | 18:1(13E) | 8.95 | 1.5 ± 0.1 | 0.07 | 1149 | H | 61 ± 4 |  | Y<sup>12</sup> |
| C18:1n-6_(E) | 18:1(12E) | 12E-octadecenoic acid | LMFA01030079 | trans-12-elaidic acid | 8.95 | 3.77 ± 0.06 | 0.02 | 4093 | H | 153 ± 6 | Y<sup>22</sup> |  |
| C18:1n-12 | 18:1(6Z) | 6Z-octadecenoic acid | LMFA01030066 | Petroselinic acid | 8.95 | 0.25 ± 0.04 | 0.2 | 124 | H | 10 ± 2 | Y<sup>7</sup> |  |
| C18:1n-9 | 18:1(9Z) | 9Z-octadecenoic acid | LMFA01030002 | Oleic acid | 8.79 | 61 ± 4 | 0.07 | 3575 | H | 2500 ± 100 | Y<sup>6</sup> |  |
| C18:1n-11 | 18:1(7Z) | 7Z-octadecenoic acid | LMFA01030068 | 7Z-octadecenoic acid | 8.85 | 0.32 ± 0.02 | 0.06 | 521 | H | 13 ± 1 |  | Y<sup>1</sup> |
| C18:1n-7_(E) | 18:1(11E) | 11E-octadecenoic acid | LMFA01030077 | Trans-vaccenic acid | 8.96 | 3 ± 1 | 0.3 | 4004 | H | 140 ± 60 | Y<sup>23</sup> |  |
| C18:1n-2_(E) | 18:1(16E) | 16E-octadecenoic acid | LMFA01030081 | 16E-octadecenoic acid | 8.95 | 0.91 ± 0.05 | 0.06 | 381 | H | 37 ± 3 |  | Y<sup>12</sup> |
| C18:1n-3_(E) | 18:1(15E) | 15E-octadecenoic acid | LMFA01030080 | 15E-octadecenoic acid | 8.95 | 0.51 ± 0.02 | 0.04 | 354 | H | 21 ± 1 |  | Y<sup>12</sup> |
| C18:1n-9_(E) | 18:1(9E) | 9E-octadecenoic acid | LMFA01030073 | 9-elaidic acid | 8.95 | 3.9 ± 0.4 | 0.1 | 5495 | H | 160 ± 20 | Y<sup>23</sup> |  |
| C18:1n-8_(E) | 18:1(10E) | 10E-octadecenoic acid | LMFA01030075 | 10E-octadecenoic acid | 8.95 | 4 ± 1 | 0.3 | 4004 | H | 150 ± 60 | Y<sup>22</sup> |  |
| C18:1n-10_(E) | 18:1(8E) | 8E-octadecenoic acid | LMFA01030071 | trans-8-elaidic acid | 8.98 | 2.06 ± 0.07 | 0.03 | 2988 | H | 84 ± 3 |  | Y<sup>12</sup> |
| C18:1n-4_(E) | 18:1(14E) | 14E-octadecenoic acid | LMFA01030881 | 18:1(14E) | 8.95 | 1.5 ± 0.2 | 0.1 | 860 | H | 59 ± 7 |  | Y<sup>12</sup> |
| C18:1n-13 | 18:1(5Z) | 5Z-octadecenoic acid | LMFA01030296 | 5Z-octadecenoic acid | 8.99 | 0.54 ± 0.04 | 0.07 | 753 | H | 21.8 ± 0.7 | Y<sup>22</sup> |  |
| C18:1n-11_(E) | 18:1(7E) | 7E-octadecenoic acid | LMFA01030069 | 7E-octadecenoic acid | 9.06 | 0.5 ± 0.2 | 0.4 | 1298 | H | 20 ± 9 | - | - |
| C18:1n-12_(E) | 18:1(6E) | 6E-octadecenoic acid | LMFA01030067 | Petroselaidic acid | 9.14 | 0.5 ± 0.2 | 0.4 | 2193 | H | 20 ± 10 |  | Y<sup>24</sup> |
| C18:1n-13_(E) | 18:1(5E) | 5E-octadecenoic acid | LMFA01030065 | Thalictic acid | 9.22 | 0.34 ± 0.04 | 0.1 | 831 | H | 14 ± 2 | Y<sup>22</sup> |  |
| C19:1n-11_br | 19:1(8)_br |  | Not found in LIPID MAPS. | — | 9.28 | 2.0 ± 0.4 | 0.2 | 42 | H | 0.1 ± 0.05 |  |  |
| C19:1n-9_br | 19:1(10)_br |  |  |  | 9.28 | 0.79 ± 0.07 | 0.09 | 53 | H | 0.04 ± 0.01 | Y<sup>14</sup> |  |
| C19:1n-8 | 19:1(11Z) | 11Z-nonadecenoic acid | LMFA01030887 | 19:1(11Z) | 9.40 | 52 ± 2 | 0.04 | 1463 | H | 2 ± 1 | Y<sup>2</sup> |  |
| C19:1n-9 | 19:1(10Z) | 10Z-nonadecenoic acid | LMFA01030362 | 10Z-nonadecenoic acid | 9.41 | 4 ± 1 | 0.2 | 76 | H | 0.19 ± 0.04 | Y<sup>14</sup> |  |
| C19:1n-10 | 19:1(9Z) | 9Z-nonadecenoic acid | LMFA01030886 | 19:1(9Z) | 9.44 | 35 ± 3 | 0.09 | 503 | H | 1.6 ± 0.7 | Y<sup>2</sup> |  |
| C19:1n-11 | 19:1(8Z) | 8Z-nonadecenoic acid | Not found in LIPID MAPS. | — | 9.49 | 0.4 ± 0.05 | 0.1 | 45 | H | 0.017 ± 0.007 | - | - |
| C19:1n-8_(E) | 19:1(11E) | 11E-nonadecenoic acid | Not found in LIPID MAPS. | — | 9.6 | 1.4 ± 0.2 | 0.1 | 52 | H | 0.06 ± 0.02 | - | - |
| C19:1n-9_(E) | 19:1(10E) | 10E-nonadecenoic acid | LMFA01030361 | 10E-nonadecenoic acid | 9.61 | 4 ± 3 | 0.8 | 32 | H | 0.14 ± 0.09 | - | - |
| C20:1n-7 | 20:1(13Z) | 13Z-eicosenoic acid | LMFA01030367 | Paullinic acid | 10 | 3.3 ± 0.3 | 0.09 | 420 | H | 0.4 ± 0.2 | Y<sup>2</sup> |  |
| C20:1n-8 | 20:1(12Z) | 12Z-eicosenoic acid | Not found in LIPID MAPS. | — | 10 | 0.43 ± 0.09 | 0.2 | 23 | H | 0.06 ± 0.04 | - | - |
| C20:1n-9 | 20:1(11Z) | 11Z-eicosenoic acid | LMFA01030085 | cis-gondoic acid | 10.01 | 84 ± 2 | 0.02 | 7721 | H | 10 ± 6 | Y<sup>8</sup> |  |
| C20:1n-5 | 20:1(15Z) | 15Z-eicosenoic acid | LMFA01030369 | 15Z-eicosenoic acid | 10.03 | 0.5 ± 0.2 | 0.4 | 27 | H | 0.05 ± 0.02 |  | Y<sup>1</sup> |
| C20:1n-6 | 20:1(14Z) | 14Z-eicosenoic acid | LMFA01030087 | 14Z-eicosenoic acid | 10 | 0.7 ± 0.1 | 0.2 | 66 | H | 0.08 ± 0.05 |  | Y<sup>25</sup> |
| C20:1n-11 | 20:1(9Z) | 9Z-eicosenoic acid | LMFA01030084 | cis-Gadoleic acid | 10.06 | 4 ± 1 | 0.2 | 792 | H | 0.6 ± 0.4 |  | Y<sup>12</sup> |
| C20:1n-12 | 20:1(8Z) | 8Z-eicosenoic acid | LMFA01030372 | 8Z-eicosenoic acid | 10.1 | 0.4 ± 0.2 | 0.5 | 72 | H | 0.05 ± 0.04 | Y<sup>22</sup> |  |
| C20:1n-6 (E) | 20:1(14E) | 14E-eicosenoic acid | Not found in LIPID MAPS. | — | 10.19 | 0.48 ± 0.09 | 0.2 | 113 | H | 0.06 ± 0.04 | - | - |
| C20:1n-13 | 20:1(7Z) | 7Z-eicosenoic acid | LMFA01030858 | 20:1(7Z) | 10.16 | 0.5 ± 0.2 | 0.4 | 91 | H | 0.06 ± 0.05 |  | Y<sup>16</sup> |
| C20:1n-8 (E) | 20:1(12E) | 12E-eicosenoic acid | Not found in LIPID MAPS. | — | 10.19 | 1.7 ± 0.5 | 0.3 | 568 | H | 0.2 ± 0.1 | - | - |
| C20:1n-9 (E) | 20:1(11E) | 11E-eicosenoic acid | LMFA01030086 | trans-gondoic acid | 10.21 | 2.0 ± 0.3 | 0.2 | 692 | H | 0.2 ± 0.1 |  | Y<sup>26</sup> |
| C20:1n-10 (E) | 20:1(10E) | 10E-eicosenoic acid | Not found in LIPID MAPS. | — | 10.23 | 0.52 ± 0.06 | 0.1 | 156 | H | 0.06 ± 0.04 | - | - |
| C20:1n-4 (E) | 20:1(16E) | 16E-eicosenoic acid | Not found in LIPID MAPS. | — | 10.21 | 0.36 ± 0.02 | 0.06 | 74 | H | 0.04 ± 0.03 | - | - |
| C20:1n-5 (E) | 20:1(15E) | 15E-eicosenoic acid | Not found in LIPID MAPS. | — | 10.20 | 0.42 ± 0.06 | 0.1 | 50 | H | 0.05 ± 0.03 | - | - |
| C20:1n-7 (E) | 20:1(13E) | 13E-eicosenoic acid | Not found in LIPID MAPS. | — | 10.19 | 0.57 ± 0.08 | 0.1 | 151 | H | 0.07 ± 0.05 | - | - |
| C21:1n-8 | 21:1(13Z) | 13Z-heneicosenoic acid | Not found in LIPID MAPS. | — | 10.58 | 29 ± 3 | 0.1 | 64 | H | 0.1 ± 0.03 | - | - |
| C21:1n-9 | 21:1(12Z) | 12Z-heneicosenoic acid | LMFA01030398 | 12Z-heneicosenoic acid | 10.59 | 20 ± 4 | 0.2 | 68 | H | 0.07 ± 0.03 | Y<sup>27</sup> |  |
| C21:1n-12 | 21:1(9Z) | 9Z-heneicosenoic acid | Not found in LIPID MAPS. | — | 10.66 | 52 ± 5 | 0.1 | 67 | H | 0.17 ± 0.07 |  | Y<sup>12</sup> |
| C22:1n-9 | 22:1(13Z) | 13Z-docosenoic acid | LMFA01030089 | cis-erucic acid | 11.16 | n.a. | 0.1 | 3307 | H | - | Y<sup>8</sup> |  |
| C22:1n-6 | 22:1(16Z) | 16Z-docosenoic acid | Not found in LIPID MAPS. | — | 11.17 | n.a. | 0.1 | 21 | H | - | - | - |
| C22:1n-7 | 22:1(15Z) | 15Z-docosenoic acid | LMFA01030402 | 15Z-docosenoic acid | 11.16 | n.a. | 0.1 | 125 | H | - | Y<sup>2</sup> |  |
| C22:1n-8 | 22:1(14Z) | 14Z-docosenoic acid | Not found in LIPID MAPS. | — | 11.15 | n.a. | 0.1 | 9 | H | - | - | - |
| C22:1n-11 | 22:1(11Z) | 11Z-docosenoic acid | LMFA01030088 | cis-cetoleic acid | 11.19 | n.a. | 0.1 | 109 | H | - |  | Y<sup>28</sup> |
| C22:1n-5 | 22:1(17Z) | 17Z-docosenoic acid | LMFA04000085 | 17Z-Docosenoic acid | 11.2 | n.a. | 0.1 | 9 | H | - |  | Y<sup>1</sup> |
| C22:1n-13 | 22:1(9Z) | 9Z-docosenoic acid | LMFA01030911 | 22:1(9Z) | 11.25 | n.a. | 0.1 | 103 | H | - |  | Y<sup>12</sup> |
| C22:1n-7 (E) | 22:1(15E) | 15E-docosenoic acid | Not found in LIPID MAPS. | — | 11.34 | n.a. | 0.1 | 27 | H | - | - | - |
| C22:1n-8 (E) | 22:1(14E) | 14E-docosenoic acid | Not found in LIPID MAPS. | — | 11.34 | n.a. | 0.1 | 108 | H | - | - | - |
| C22:1n-6 (E) | 22:1(16E) | 16E-docosenoic acid | Not found in LIPID MAPS. | — | 11.34 | n.a. | 0.1 | 30 | H | - | - | - |
| C22:1n-9 (E) | 22:1(13E) | 13E-docosenoic acid | LMFA01030090 | trans-brassicidic acid | 11.34 | n.a. | 0.1 | 113 | H | - |  | Y<sup>29</sup> |
| C22:1n-10 (E) | 22:1(12E) | 12E-docosenoic acid | Not found in LIPID MAPS. | — | 11.36 | n.a. | 0.1 | 74 | H | - | - | - |
| C22:1n-5 (E) | 22:1(17E) | 17E-docosenoic acid | Not found in LIPID MAPS. | — | 11.36 | n.a. | 0.1 | 62 | H | - | - | - |
| C23:1n-14 br | 23:1(9 br) |  |  |  | 11.41 | 40 ± 10 | 0.3 | 36 | H | 0.05 ± 0.02 |  |  |
| C23:1n-9 | 23:1(14Z) | 14Z-tricosenoic acid | LMFA01030413 | 14Z-tricosenoic acid | 11.71 | 42 ± 9 | 0.2 | 106 | H | 0.05 ± 0.01 |  | Y<sup>12</sup> |
| C23:1n-8 | 23:1(15Z) | 15Z-tricosenoic acid | Not found in LIPID MAPS. | — | 11.72 | 17 ± 2 | 0.1 | 34 | H | 0.022 ± 0.007 | - | - |
| C24:1n-9 | 24:1(15Z) | 15Z-tetracosenoic acid | LMFA01030092 | Nervonic acid | 12.25 | 91.5 ± 0.4 | 0.004 | 4095 | H | 4.6 ± 0.1 | Y<sup>8</sup> |  |
| C24:1n-7 | 24:1(17Z) | 17Z-tetracosenoic acid | LMFA01031084 | 17Z-Tetracosenoic acid | 12.26 | 0.6 ± 0.2 | 0.4 | 64 | H | 0.027 ± 0.008 |  | Y<sup>30</sup> |
| C24:1n-8 | 24:1(16Z) | 16Z-tetracosenoic acid | Not found in LIPID MAPS. | — | 12.26 | 0.5 ± 0.2 | 0.4 | 19 | H | 0.023 ± 0.008 | - | - |
| C24:1n-11 | 24:1(13Z) | 13Z-tetracosenoic acid | Not found in LIPID MAPS. | — | 12.26 | 0.41 ± 0.07 | 0.2 | 21 | H | 0.021 ± 0.004 | - | - |
| C24:1n-5 | 24:1(19Z) | 19Z-tetracosenoic acid | LMFA01031022 | 19Z-Tetracosenoic acid | 12.31 | 0.2 ± 0.1 | 0.7 | 9 | H | 0.008 ± 0.005 |  | Y<sup>1</sup> |
| C24:1n-6 | 24:1(18Z) | 18Z-tetracosenoic acid | Not found in LIPID MAPS. | — | 12.28 | 0.6 ± 0.1 | 0.2 | 51 | H | 0.028 ± 0.006 | - | - |
| C24:1n-5 (E) | 24:1(19E) | 19E-tetracosenoic acid | Not found in LIPID MAPS. | — | 12.45 | 1.11 ± 0.04 | 0.04 | 173 | H | 0.056 ± 0.001 |  | Y<sup>12</sup> |
| C24:1n-6 (E) | 24:1(18E) | 18E-tetracosenoic acid | Not found in LIPID MAPS. | — | 12.44 | 1.3 ± 0.2 | 0.2 | 189 | H | 0.067 ± 0.009 |  | Y<sup>12</sup> |
| C24:1n-7 (E) | 24:1(17E) | 17E-tetracosenoic acid | Not found in LIPID MAPS. | — | 12.43 | 1.1 ± 0.3 | 0.3 | 176 | H | 0.05 ± 0.02 |  | Y<sup>12</sup> |
| C24:1n-10 (E) | 24:1(14E) | 14E-tetracosenoic acid | Not found in LIPID MAPS. | — | 12.43 | 0.41 ± 0.02 | 0.05 | 90 | H | 0.02 ± 0.001 | - | - |
| C24:1n-4 (E) | 24:1(20E) | 20E-tetracosenoic acid | Not found in LIPID MAPS. | — | 12.47 | 0.96 ± 0.06 | 0.06 | 184 | H | 0.048 ± 0.003 | - | - |
| C24:1n-8 (E) | 24:1(16E) | 16E-tetracosenoic acid | Not found in LIPID MAPS. | — | 12.42 | 0.4 ± 0.04 | 0.1 | 100 | H | 0.02 ± 0.002 | - | - |
| C24:1n-9 (E) | 24:1(15E) | 15E-tetracosenoic acid | LMFA01030093 | trans-selacholeic acid | 12.43 | 1.1 ± 0.07 | 0.06 | 262 | H | 0.055 ± 0.002 |  | Y<sup>30</sup> |
| C14:2n-6,9 | 14:2(5,8) | 5,8-tetradecadienoic acid | LMFA01030257 | Goshuyic acid | 5.36 | = | = | 133 | H | 1.3 ± 0.2 | γ<sup>11</sup> |  |
| C16:2n-6,9 | 16:2(7,10) | 7Z,10Z-hexadecadienoic acid | LMFA01030807 | 7Z,10Z-hexadecadienoic acid | 6.7 | 32 ± 9 | 0.3 | 553 | H | 2.5 ± 0.5 |  | γ<sup>31</sup> |
| C16:2n-7,10 | 16:2(6,9) | 6Z,9Z-hexadecadienoic acid | LMFA01030273 | 6Z,9Z-hexadecadienoic acid | 6.79 |  |  | 81 | L |  |  |  |
| C16:2n-3,10 | 16:2(6,13) | 6,13-hexadecadienoic acid | Not found in LIPID MAPS. | — | 6.79 |  |  | 104 | L |  |  |  |
| C16:2n-4,7 | 16:2(7,12) | 9,12-hexadecadienoic acid | LMFA01030275 | Palmitolinoleic acid | 6.81 |  |  | 103 | L |  |  | γ<sup>31</sup> |
| C16:2n-4,9 | 16:2(7,12) | 7,12-hexadecadienoic acid | Not found in LIPID MAPS. | — | 6.84 |  |  | 95 | L |  |  |  |
| C16:2n-5,7 | 16:2(9,11) | 9,11-hexadecadienoic acid | Not found in LIPID MAPS. | — | 6.85 |  |  | 207 | L |  |  |  |
| C16:2n-5,9 | 16:2(7,11) | 7,11-hexadecadienoic acid | Not found in LIPID MAPS. | — | 6.85 |  |  | 195 | L |  |  |  |
| C16:2n-6,9_(E) | 16:2(7,10) | 7,10-hexadecadienoic acid | Not found in LIPID MAPS. | — | 6.94 |  |  | 124 | L |  |  |  |
| C16:2n-7,9 | 16:2(7,9) | 7,9-hexadecadienoic acid | Not found in LIPID MAPS. | — | 6.94 |  |  | 167 | L |  |  |  |
| C16:2n-7,10_(E) | 16:2(6,9) | 6,9-hexadecadienoic acid | Not found in LIPID MAPS. | — | 6.94 |  |  | 71 | L |  |  |  |
| C16:2n-3,9 | 16:2(7,13) | 7,13-hexadecadienoic acid | Not found in LIPID MAPS. | — | 7.03 |  |  | 16 | L |  |  |  |
| C16:2n-3,10_(E) | 16:2(6,13) | 6,13-hexadecadienoic acid | Not found in LIPID MAPS. | — | 7.05 |  |  | 47 | L |  |  |  |
| C16:2n-5,10 | 16:2(6,11) | 6,11-hexadecadienoic acid | Not found in LIPID MAPS. | — | 7.04 |  |  | 110 | L |  |  |  |
| C16:2n-8,10 | 16:2(6,8) | 6,8-hexadecadienoic acid | Not found in LIPID MAPS. | — | 7.05 |  |  | 54 | L |  |  |  |
| C16:2n-6,10 | 16:2(6,10) | 6,10-hexadecadienoic acid | Not found in LIPID MAPS. | — | 7.07 | 3.9 ± 0.7 | 0.2 | 69 | L | 0.29 ± 0.03 |  |  |
| 17:2n5,8 | 17:2(9,12) | 9Z,12Z-heptadecadienoic acid | Not found in LIPID MAPS. | — | 7.28 | 70 ± 20 | 0.3 | 75 | H | 1.1 ± 0.2 |  | γ<sup>32</sup> |
| C17:2n-6,9 | 17:2(8,11) | 8Z,11Z-heptadecadienoic acid | LMFA01031037 | Norlinoleic acid | 7.34 | 30 ± 20 | 0.6 | 33 | H | 0.5 ± 0.4 |  | γ<sup>33</sup> |
| C18:2n-6,9 | 18:2(9,12) | 9Z,12Z-octadecadienoic acid | LMFA01030120 | Linoleic acid | 7.96 | 83 ± 2 | 0.02 | 3292 | H | 3050 ± 30 | γ<sup>6</sup> |  |
| C18:2n-2,9 | 18:2(9,16) | 9,16-octadecadienoic acid | Not found in LIPID MAPS. | — | 8.05 |  |  | 1368 | L | - |  |  |
| C18:2n-2,10 | 18:2(8,16) | 8,16-octadecadienoic acid | Not found in LIPID MAPS. | — | 8.06 |  |  | 20 | L | - |  |  |
| C18:2n-3,9 | 18:2(9,15) | 9,15-octadecadienoic acid | Not found in LIPID MAPS. | Mangiferic acid | 8.06 |  |  | 1916 | L | - | γ<sup>22</sup> |  |
| C18:2n-4,9 | 18:2(9,14) | 9,14-octadecadienoic acid | Not found in LIPID MAPS. | — | 8.08 |  |  | 2432 | L | - | γ<sup>22</sup> |  |
| C18:2n-5,9 | 18:2(9,13) | 9,13-octadecadienoic acid | Not found in LIPID MAPS. | — | 8.11 |  |  | 2344 | L | - | γ<sup>22</sup> |  |
| C18:2n-6,12 | 18:2(6,12) | 6,12-octadecadienoic acid | Not found in LIPID MAPS. | — | 8.07 |  |  | 857 | L | - |  |  |
| C18:2n-3,10 | 18:2(8,15) | 8,15-octadecadienoic acid | Not found in LIPID MAPS. | — | 8.14 |  |  | 307 | L | - |  |  |
| C18:2n-3,13 | 18:2(5,15) | 5,15-octadecadienoic acid | Not found in LIPID MAPS. | — | 8.11 |  |  | 105 | L | - |  |  |
| C18:2n-5,10 | 18:2(8,13) | 8,13-octadecadienoic acid | Not found in LIPID MAPS. | — | 8.14 |  |  | 843 | L | - |  |  |
| C18:2n-6,13 | 18:2(5,12) | 5,12-octadecadienoic acid | LMFA01030110 | 5Z,12Z-octadecadienoic acid | 8.11 |  |  | 401 | L | - |  |  |
| C18:2n-6,8 | 18:2(10,12) | 10,12-octadecadienoic acid | LMFA01030124 |  | 8.19 |  |  | 1451 | L | - | γ<sup>23</sup> |  |
| C18:2n-6,9_(E) | 18:2(9,12) | 9,12-octadecadienoic acid | LMFA02000352 |  | 8.19 |  |  | 2309 | L | - | γ<sup>22</sup> |  |
| C18:2n-7,9 | 18:2(9,1<br>1) | 9,11-octadecadienoic<br>acid | LMFA01030117 | CLA | 8.19 |  |  | 3545 | L | - | γ<sup>22</sup> |  |
| C18:2n-6,10 | 18:2(8,1<br>2) | 8,12-octadecadienoic<br>acid | Not found in LIPID<br>MAPS. | — | 8.22 |  |  | 1247 | L | - |  |  |
| C18:2n-6,12_(E) | 18:2(6,1<br>2) | 6,12-octadecadienoic<br>acid | Not found in LIPID<br>MAPS. | — | 8.22 |  |  | 654 | L | - |  |  |
| C18:2n-7,9_(E) | 18:2(9,1<br>1) | 9,11-octadecadienoic<br>acid | LMFA01030118 | Rumenic acid | 8.45 |  |  | 257 | L | - |  |  |
| C18:2n-8,10 | 18:2(8,1<br>0) | 8,10-octadecadienoic<br>acid | Not found in LIPID<br>MAPS. | — | 8.5 |  |  | 43 | L | - |  |  |
| C18:2n-6,8_(E) | 18:2(10,<br>12) | 10,12-octadecadienoic<br>acid | LMFA01031059 | 10Z,12E-Octadecadienoic<br>acid | 8.5 |  |  | 9 | L | - |  |  |
| C18:2n-6,9_(E) | 18:2(9,1<br>2) | 9E,12E-octadecadienoic<br>acid | LMFA02000352 | Linoelaidic acid | 8.46 |  |  | 155 | L | - | γ<sup>22</sup> |  |
| C19:2n-6,9 | 19:2(10,<br>13) | 10Z,13Z-nonadecadienoic<br>acid | LMFA01030129 | 10Z,13Z-nonadecadienoic<br>acid | 8.6 | 30 ±<br>20 | 0.6 | 54 | H | 0.6 ±<br>0.5 | γ<sup>27</sup> |  |
| C19:2n-7,10 | 19:2(9,1<br>2) | 9Z,12Z-nonadecadienoic<br>acid | Not found in LIPID<br>MAPS. | — | 8.61 | 40 ±<br>20 | 0.5 | 69 | H | 0.7 ±<br>0.4 | - | γ<sup>34</sup> |
| C19:2n-8,11 | 19:2(8,1<br>1) | 8Z,11Z-nonadecadienoic<br>acid | Not found in LIPID<br>MAPS. | — | 8.65 | 30 ±<br>10 | 0.4 | 52 | H | 0.37 ±<br>0.08 | - | - |
| C20:2n-6,9 | 20:2(11,<br>14) | 11Z,14Z-eicosadienoic acid | LMFA01031043 | Dihomolinoleic<br>acid | 9.18 | 89.7 ±<br>0.4 | 0.005 | 260 | H | 57 ± 7 | γ<sup>6</sup> |  |
| C20:2n-9,12 | 20:2(8,1<br>1) | 8Z,11Z-eicosadienoic acid | LMFA01030377 | 8Z,11Z-eicosadienoic acid | 9.28 | 5.7 ±<br>0.2 | 0.04 | 1318 | H | 3.7 ±<br>0.6 | γ<sup>8</sup> |  |
| C20:2n-7,9 | 20:2(11,<br>13) | 11,13-eicosadienoic acid | Not found in LIPID<br>MAPS. | — | 9.39 | 1.1 ±<br>0.3 | 0.3 | 298 | H | 0.7 ±<br>0.2 | - | - |
| C20:2n-9,13 | 20:2(7,1<br>1) | 7,11-eicosadienoic<br>acid | LMFA01031035 | Dihomotaxoleic<br>acid | 9.39 | 0.3 ±<br>0.1 | 0.3 | 100 | H | 0.2 ±<br>0.07 |  | γ<sup>35</sup> |
| C20:2n-9,15 | 20:2(5,1<br>1) | 5,11-eicosadienoic<br>acid | LMFA01030865 | Keteleeronic acid | 9.40 | 3.2 ±<br>0.2 | 0.06 | 597 | H | 2.0 ±<br>0.4 |  | γ<sup>28</sup> |
| C22:2n-6,9 | 22:2(13,<br>16) | 13Z,16Z-docosadienoic<br>acid | LMFA01030405 | 13Z,16Z-docosadienoic<br>acid | 10.36 | 59.1 ±<br>0.2 | 0.003 | 324 | H | 0.44 ±<br>0.04 | γ<sup>6</sup> |  |
| C22:2n-9,15 | 22:2(7,1<br>3) | 7,13-docosadienoic<br>acid | LMFA04000061 | 22:2(7Z,13Z) | 10.43 | 40.9 ±<br>0.2 | 0.005 | 288 | H | 0.3 ±<br>0.03 |  | γ<sup>16</sup> |
| C24:2n-6,9 | 24:2(15,<br>18) | 15Z,18Z-tetracosadienoic<br>acid | LMFA01031071 | 15Z,18Z-Tetracosadienoic<br>acid | 11.47 |  |  | 415 | H | 0.3 ±<br>0.07 | γ<sup>8</sup> |  |
| C24:2n-7,9 | 24:2(15,<br>17) | 15,17-tetracosadienoic<br>acid | Not found in LIPID<br>MAPS. | — | 11.6 | 10.5 ±<br>0.9 | 0.09 | 148 | H | 0.04 ±<br>0.01 | - | - |
| C24:2n-5,9 | 24:2(15,<br>19) | 15,19-tetracosadienoic<br>acid | Not found in LIPID<br>MAPS. | — | 11.6 | 10 ± 3 | 0.3 | 165 | H | 0.04 ±<br>0.01 | - | - |
| C16:3n-6,9,12 | 16:3(4,<br>7,10) | 4,7,10-hexadecatrienoic<br>acid | LMFA01030134 | 4,7,10-hexadecatrienoic<br>acid | 6.25 |  |  | 145 | H | 2.4 ±<br>0.2 |  | γ<sup>31</sup> |
| C18:3n-3,6,9 | 18:3(9,<br>12,15) | 9Z,12Z,15Z-octadecatrienoic<br>acid | LMFA01030152 | alpha-Linolenic<br>acid | 7.24 | 56 ± 6 | 0.1 | 2118 | H | 68 ± 9 | γ<sup>6</sup> |  |
| C18:3n-3,7,9 | 18:3(9,<br>11,15) | 9,11,15-octadecatrienoic<br>acid | Not found in LIPID<br>MAPS. | — | 7.41 | 3 ± 2 | 0.7 | 141 | H | 3 ± 3 | - | - |
| C18:3n-4,7,9 | 18:3(9,<br>11,14) | 9,11,14-octadecatrienoic<br>acid | Not found in LIPID<br>MAPS. | — | 7.31 | 4 ± 2 | 0.6 | 103 | H | 4 ± 3 | - | - |
| C18:3n-6,9,12 | 18:3(6,<br>9,12) | 6Z,9Z,12Z-octadecatrienoic<br>acid | LMFA01030141 | gamma-Linolenic<br>acid | 7.38 | 29 ± 3 | 0.1 | 1843 | H | 35 ± 3 | γ<sup>6</sup> |  |
| C18:3n-3,6,9_(E) | 18:3(9,<br>12,15) | 9,12,15-octadecatrienoic<br>acid |  |  | 7.45 | 4 ± 4 | 0.9 | 145 | H | 5 ± 5 | - | - |
| C18:3n-6,9,12_(E) | 18:3(6,<br>9,12) | 6,9,12-octadecatrienoic<br>acid | Not found in LIPID<br>MAPS. | — | 7.58 | 1.4 ±<br>0.5 | 0.4 | 343 | H | 1.7 ±<br>0.7 | - | - |
| C18:3n-7,10,13 | 18:3(5,<br>8,11) | 5,8,11-octadecatrienoic<br>acid | Not found in LIPID<br>MAPS. | — | 7.58 | 2 ± 2 | 0.9 | 98 | H | 3 ± 3 | - | γ<sup>36</sup> |
| C18:3n-3,6,9_(E) | 18:3(9,<br>12,15) | 9,12,15-octadecatrienoic<br>acid |  |  | 7.58 | 4 ± 4 | 0.9 | 51 | H | 5 ± 5 | - | - |
| C20:3n-6,9,12 | 20:3(8,<br>11,14) | 8Z,11Z,14Z-eicosatrienoic acid | LMFA01030387 |  | 8.51 | 84 ± 2 | 0.02 | 797 | H | 300 ±<br>10 | γ<sup>6</sup> |  |
| C20:3n-6,9,15 | 20:3(5,<br>11,14) | 5Z,11Z,14Z-eicosatrienoic acid | LMFA01030380 | Sciadonic acid | 8.60 | 13 ± 1 | 0.08 | 1085 | H | 47 ± 8 |  | γ<sup>28</sup> |
| C20:3n-7,10,12 | 20:3(8,<br>10,13) | 8,10,13-eicosatrienoic acid | Not found in LIPID<br>MAPS. | — | 8.68 | 0.7 ±<br>0.4 | 0.6 | 275 | H | 3 ± 1 | - | - |
| C20:3n-7,10,13 | 20:3(7,<br>10,13) | 7,10,13-eicosatrienoic acid | LMFA01030384 | 7Z,10Z,13Z-eicosatrienoic acid | 8.66 | 0.5 ±<br>0.2 | 0.4 | 149 | H | 1.7 ±<br>0.6 | - | γ<sup>37</sup> |
| C20:3n-9,12,15 | 20:3(5,<br>8,11) | 5Z,8Z,11Z-eicosatrienoic acid | LMFA01030381 | Mead acid | 8.79 | 1.2 ±<br>0.2 | 0.2 | 373 | H | 4.1 ±<br>0.8 | γ<sup>8</sup> |  |
| C22:3n-6,9,15 | 22:3(7,13,16) | 7Z,13Z,16Z-docosatrenoic acid | LMFA04000062 | 22:3(7Z,13Z,16Z) | 9.65 | 18 ± 9 | 0.5 | 111 | H | 0.4 ± 0.2 |  | Y<sup>16</sup> |
| C22:3n-6,9,12 | 22:3(10,13,16) | 10Z,13Z,16Z-docosatrenoic acid | Not found in LIPID MAPS. | — | 9.64 | 41 ± 4 | 0.1 | 139 | H | 1.0 ± 0.2 | Y<sup>2</sup> |  |
| C22:3n-9,12,15 | 22:3(7,10,13) | 7Z,10Z,13Z-docosatrenoic acid | LMFA01030411 | Dihomo Mead's acid | 9.79 | 41 ± 7 | 0.2 | 115 | H | 1.0 ± 0.2 |  | Y<sup>38</sup> |
| C18:4n-3,6,9,12 | 18:4(6,9,12,15) | 6Z,9Z,12Z,15Z-octadecatetraenoic acid | LMFA01030357 | Stearidonic acid | 6.66 | — | — | 263 | H | 2.1 ± 0.4 | Y<sup>6</sup> |  |
| C20:4n-6,9,12,15 | 20:4(5,8,11,14) | 5Z,8Z,11Z,14Z-eicosatetraenoic acid | LMFA01030001 | Arachidonic acid | 8.01 | 97.5 ± 0.3 | 0.003 | 266 | H | 920 ± 50 | Y<sup>6</sup> |  |
| C20:4n-3,6,9,12 | 20:4(8,11,14,17) | 8Z,11Z,14Z,17Z-eicosatetraenoic acid | LMFA01030818 | omega-3-Arachidonic acid | 7.82 | 2.0 ± 0.3 | 0.2 | 557 | H | 19 ± 3 | Y<sup>14</sup> |  |
| C20:4n-3,6,9,15 | 20:4(5,11,14,17) | 5Z,11Z,14Z,17Z-eicosatetraenoic acid | LMFA01030394 | Juniperonic acid | 7.9 | 0.55 ± 0.08 | 0.2 | 396 | H | 5 ± 1 |  | Y<sup>39</sup> |
| C22:4n-6,9,12,15 | 22:4(7,10,13,16) | 7Z,10Z,13Z,16Z-docosatetraenoic acid | LMFA01030178 | Adrenic Acid | 9.03 | 98.9 ± 0.03 | 0.0003 | 494 | H | 27 ± 2 | Y<sup>6</sup> |  |
| C22:4n-3,6,9,12 | 22:4(10,13,16,19) | 10Z,13Z,16Z,19Z-docosatetraenoic acid | Not found in LIPID MAPS. | — | 8.97 | 1.13 ± 0.03 | 0.03 | 33 | H | 0.31 ± 0.03 | - | - |
| C24:4n-6,9,12,15 | 24:4(9,12,15,18) | 9Z,12Z,15Z,18Z-tetracosatetraenoic acid | LMFA01030819 | 9Z,12Z,15Z,18Z-Tetracosatetraenoic acid | 10.07 | — | — | 16 | H | 0.29 ± 0.01 | Y<sup>8</sup> |  |
| C20:5n-3,6,9,12,15 | 20:5(5,8,11,14,17) | 5Z,8Z,11Z,14Z,17Z-eicosapentaenoic acid | LMFA01030759 | EPA | 7.32 | — | — | 157 | H | 21 ± 1 | Y<sup>6</sup> |  |
| C22:5n-3,6,9,12,15 | 22:5(7,10,13,16,19) | 7Z,10Z,13Z,16Z,19Z-docosapentaenoic acid | LMFA04000044 | DPA | 8.35 | 43 ± 3 | 0.07 | 331 | H | 27 ± 1 | Y<sup>8</sup> |  |
| C22:5n-6,9,12,15,18 | 22:5(4,7,10,13,16) | 4Z,7Z,10Z,13Z,16Z-docosapentaenoic acid | LMFA04000064 | 22:5(4Z,7Z,10Z,13Z,16Z) | 8.70 | 57 ± 3 | 0.05 | 483 | H | 35 ± 6 | Y<sup>8</sup> |  |
| C24:5n-3,6,9,12,15 | 24:5(9,12,15,18,21) | 9Z,12Z,15Z,18Z,21Z-tetracosapentaenoic acid | LMFA01030821 | 9Z,12Z,15Z,18Z,21Z-tetracosapentaenoic acid | 9.43 | 81 ± 1 | 0.01 | 40 | H | 0.6 ± 0.2 | Y<sup>7</sup> |  |
| C24:5n-6,9,12,15,18 | 24:5(6,9,12,15,18) | 6Z,9Z,12Z,15Z,18Z-tetracosapentaenoic acid | LMFA01030820 | 6Z,9Z,12Z,15Z,18Z-tetracosapentaenoic acid | 9.58 | 19 ± 1 | 0.05 | 13 | H | 0.13 ± 0.04 | Y<sup>7</sup> |  |
| C22:6n-3,6,9,12,15,18 | 22:6(4,7,10,13,16,19) | 4Z,7Z,10Z,13Z,16Z,19Z-docosahexaenoic acid | LMFA01030185 | DHA | 8.02 | — | — | 187 | H | 84 ± 4 | Y<sup>6</sup> |  |
| C24:6n-3,6,9,12,15,18 | 24:6(6,9,12,15,18,21) | 6Z,9Z,12Z,15Z,18Z,21Z-tetracosahexaenoic acid | LMFA01030822 | THA | 8.96 | — | — | 111 | H | 1.1 ± 0.3 | Y<sup>8</sup> |  |

### 3.4 Non-esterified fatty acids in human plasma NIST 1950 SRM

**Figure S24:**
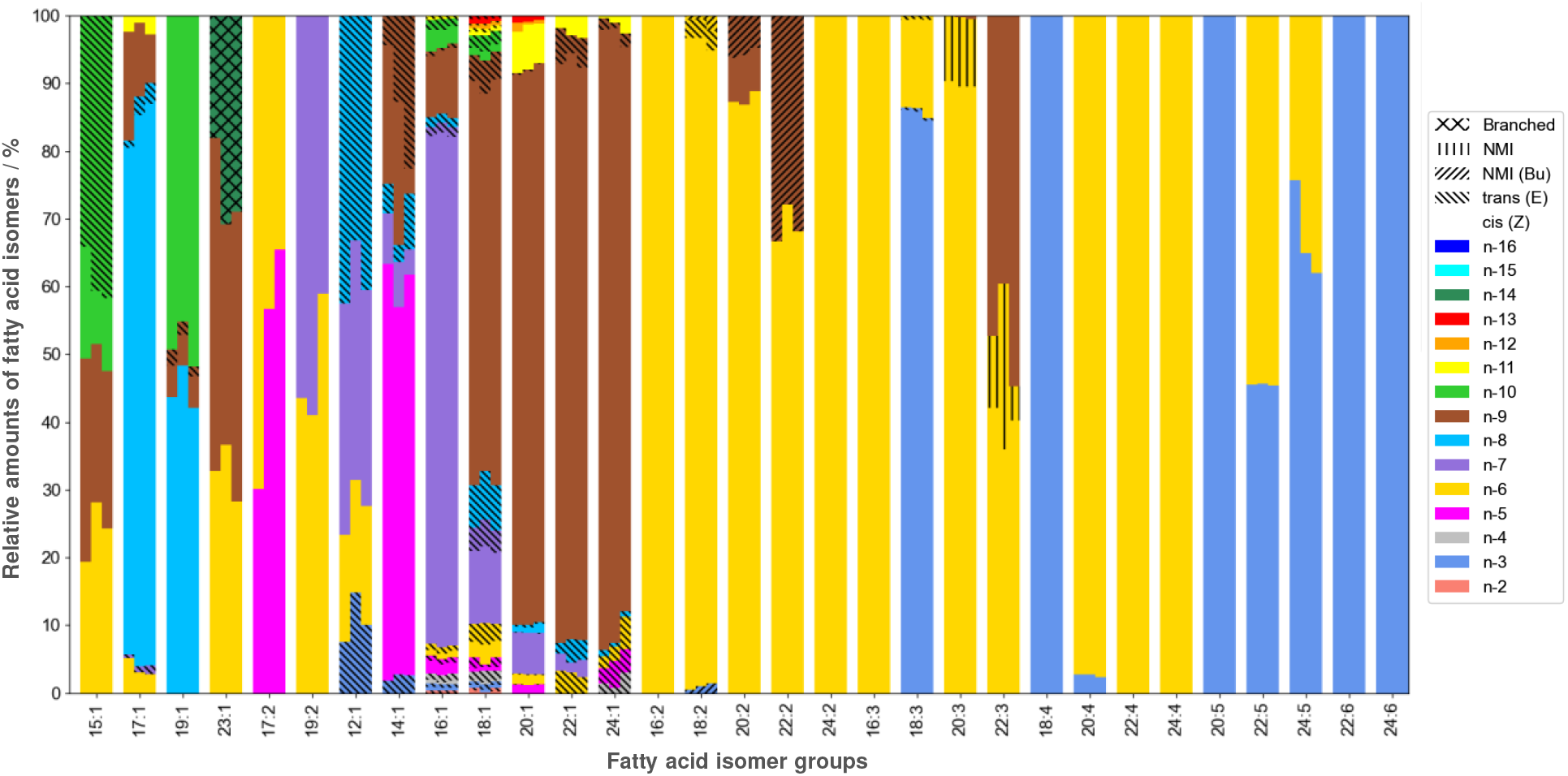
Relative abundance of fatty acid isomers by isomer group in non-esterified fatty acids (NEFA) in NIST 1950 SRM.

**Table S5:** Identification and relative quantification of non-esterified fatty acids (NEFA) in NIST 1950 human plasma standard reference material. The same lipid extract as for the analysis of the total fatty acid content was used here but hydrolysis was omitted for this analysis. Found species and their relative abundance are summarized in Figure S24.

| Fatty acid (n-x) | FA shorthand | Systematic name | LipidMAPS ID | Common Name | RT / min | Relative isomer abundance / % | COV (Rel. abundance) |
| --- | --- | --- | --- | --- | --- | --- | --- |
| C12:0 | 12:0 | dodecanoic acid | LMFA01010012 | Lauric acid | 5.44 |  |  |
| C13:0 | 13:0 | tridecanoic acid | LMFA01010013 | Tridecylic acid | 8.93 |  |  |
| C14:0 | 14:0 | tetradecanoic acid | LMFA01010014 | Myristic acid | 6.99 |  |  |
| C15:0 | 15:0 | pentadecanoic acid | LMFA01010015 | Pentadecylic acid | 7.7 |  |  |
| C16:0 | 16:0 | hexadecanoic acid | LMFA01010001 | Palmitic acid | 8.39 |  |  |
| C17:0 | 17:0 | heptadecanoic acid | LMFA01010017 | Margaric acid | 9.07 |  |  |
| C18:0 | 18:0 | octadecanoic acid | LMFA01010041 |  | 9.71 |  |  |
| C20:0 | 20:0 | eicosanoic acid | LMFA01010020 | Arachidic acid | 10.95 |  |  |
| C21:0 | 21:0 | heneicosanoic acid | LMFA01010021 | Heneicosylic acid | 11.54 |  |  |
| C22:0 | 22:0 | docosanoic acid | LMFA01010022 | Behenic acid | 12.09 |  |  |
| C23:0 | 23:0 | tricosanoic acid | LMFA01010023 | Tricosylic acid | 12.19 |  |  |
| C24:0 | 24:0 | tetracosanoic acid | LMFA01010024 | Lignoceric acid | 13.16 |  |  |
| C12:1n-3 (E) | 12:1(9E) | 9E-dodecenoic acid |  |  | 4.63 | 11 ± 4 | 0.4 |
| C12:1n-6 | 12:1(6Z) | 6Z-dodecenoic acid | LMFA01030227 | 6Z-dodecenoic acid | 4.55 | 16.6 ± 0.8 | 0.05 |
| C12:1n-7 | 12:1(5Z) | 5Z-dodecenoic acid | LMFA01030226 | Lauroleinic acid | 4.61 | 34 ± 2 | 0.06 |
| C12:1n-8 (E) | 12:1(4E) | 4E-dodecenoic acid | Not found in LIPID MAPS. |  | 4.8 | 39 ± 5 | 0.1 |
| C14:1n-5 | 14:1(9Z) | 9Z-tetradecenoic acid | LMFA01030051 | Myristoleic acid | 6.02 | 58 ± 4 | 0.07 |
| C14:1n-7 | 14:1(7Z) | 7Z-tetradecenoic acid | LMFA01030249 | 7Z-tetradecenoic acid | 6.03 | 6 ± 2 | 0.3 |
| C14:1n-3 (E) | 14:1(11E) | 11E-tetradecenoic acid |  |  | 6.19 | 2.4 ± 0.6 | 0.3 |
| C14:1n-9 | 14:1(5Z) | 5Z-tetradecenoic acid | LMFA01030248 | Physeteric acid | 6.21 | 20 ± 10 | 0.7 |
| C14:1n-8 (E) | 14:1(6E) | 6E-tetradecenoic acid | Not found in LIPID MAPS. |  | 6.31 | 5 ± 3 | 0.6 |
| C14:1n-9 (E) | 14:1(5E) | 5E-tetradecenoic acid | LMFA01060188 |  | 6.45 | 13 ± 9 | 0.7 |
| C15:1n-6 | 15:1(9Z) | 9Z-pentadecenoic acid | LMFA01030866 | 15:1(9Z) | 6.73 | 24 ± 4 | 0.2 |
| C15:1n-9 | 15:1(6Z) | 6Z-pentadecenoic acid | LMFA01031184 | 6Z-pentadecenoic acid | 6.88 | 26 ± 4 | 0.2 |
| C15:1n-10 | 15:1(5Z) | 5Z-pentadecenoic acid | LMFA01031197 | 5Z-pentadecenoic acid | 6.94 | 12 ± 4 | 0.3 |
| C15:1n-10 (E) | 15:1(5E) | 5E-pentadecenoic acid | Not found in<br>LIPID MAPS. | — | 7.17 | 39 ± 4 | 0.1 |
| C16:1n-4 | 16:1(12Z) | 12Z-hexadecenoic acid | Not found in<br>LIPID MAPS. | — | 7.44 | 0.48 ± 0.08 | 0.2 |
| C16:1n-5 | 16:1(11Z) | 11Z-hexadecenoic acid | LMFA01030262 | cis-Palmitavaccenic acid | 7.44 | 1.6 ± 0.2 | 0.1 |
| C16:1n-6 | 16:1(10Z) | 10Z-hexadecenoic acid | LMFA01030058 | cis-10-palmitoleic acid | 7.44 | 0.84 ± 0.02 | 0.02 |
| C16:1n-7 | 16:1(9Z) | 9Z-hexadecenoic acid | LMFA01030056 | cis-9-palmitoleic acid | 7.44 | 75.4 ± 0.6 | 0.008 |
| C16:1n-9 | 16:1(7Z) | 7Z-hexadecenoic acid | LMFA01030766 | — | 7.46 | 9.5 ± 0.7 | 0.07 |
| C16:1n-3 | 16:1(13Z) | 13Z-hexadecenoic acid | LMFA01030264 | 13Z-hexadecenoic acid | 7.44 | 0.4 ± 0.1 | 0.2 |
| C16:1n-2 (E) | 16:1(14E) | 14E-hexadecenoic acid | Not found in<br>LIPID MAPS. | — | 7.62 | 0.35 ± 0.06 | 0.2 |
| C16:1n-3 (E) | 16:1(13E) | 13E-hexadecenoic acid | Not found in<br>LIPID MAPS. | — | 7.59 | 0.53 ± 0.09 | 0.2 |
| C16:1n-4 (E) | 16:1(12E) | 12E-hexadecenoic acid | Not found in<br>LIPID MAPS. | — | 7.6 | 1 ± 0.1 | 0.1 |
| C16:1n-5 (E) | 16:1(11E) | 11E-hexadecenoic acid | LMFA01030261 | Lycopodic acid | 7.59 | 0.84 ± 0.02 | 0.02 |
| C16:1n-6 (E) | 16:1(10E) | 10E-hexadecenoic acid | Not found in<br>LIPID MAPS. | — | 7.59 | 0.87 ± 0.02 | 0.02 |
| C16:1n-10 | 16:1(6Z) | 6Z-hexadecenoic acid | LMFA01030267 | Sapienic acid | 7.53 | 2.8 ± 0.3 | 0.1 |
| C16:1n-11 | 16:1(5Z) | 5Z-hexadecenoic acid | LMFA01030854 | 16:1(5Z) | 7.62 | 0.05 ± 0.02 | 0.4 |
| C16:1n-7 (E) | 16:1(9E) | 9E-hexadecenoic acid | LMFA01030057 | trans-9-palmitoleic acid | 7.62 | 1.43 ± 0.06 | 0.04 |
| C16:1n-8 (E) | 16:1(8E) | 8E-hexadecenoic acid | — | — | 7.62 | 1.38 ± 0.05 | 0.04 |
| C16:1n-9 (E) | 16:1(7E) | 7E-hexadecenoic acid | LMFA01030808 | Hypogeic acid | 7.65 | 0.6 ± 0.1 | 0.2 |
| C16:1n-10 (E) | 16:1(6E) | 6E-hexadecenoic acid | LMFA01031082 | 6E-Hexadecenoic acid | 7.78 | 1.4 ± 0.2 | 0.1 |
| C16:1n-11 (E) | 16:1(5E) | 5E-hexadecenoic acid | Not found in<br>LIPID MAPS. | — | 7.88 | 0.45 ± 0.08 | 0.2 |
| C17:1n-6 | 17:1(11Z) | 11Z-heptadecenoic acid | LMFA01030856 | 17:1(11Z) | 8.12 | 4 ± 1 | 0.3 |
| C17:1n-8 | 17:1(9Z) | 9Z-heptadecenoic acid | LMFA01030060 | Margaroleic acid | 8.12 | 80 ± 4 | 0.05 |
| C17:1n-9 | 17:1(8Z) | 8Z-heptadecenoic acid | LMFA01030288 | Civetic acid | 8.12 | 11 ± 4 | 0.4 |
| C17:1n-7 (E) | 17:1(10E) | 10E-heptadecenoic acid | — | — | 8.29 | 1.0 ± 0.4 | 0.4 |
| C17:1n-8 (E) | 17:1(9E) | 9E-heptadecenoic acid | LMFA01030289 | 9E-heptadecenoic acid | 8.31 | 2 ± 1 | 0.4 |
| C17:1n-11 | 17:1(6Z) | 6Z-heptadecenoic acid | LMFA01031186 | 6Z-heptadecenoic acid | 8.29 | 2.0 ± 0.9 | 0.5 |
| C18:1n-6 | 18:1(12Z) | 12Z-octadecenoic acid | LMFA01030078 | cis-12-oleic acid | 8.77 | 2.5 ± 0.3 | 0.1 |
| C18:1n-7 | 18:1(11Z) | 11Z-octadecenoic acid | LMFA01030076 | cis-vaccenic acid | 8.77 | 10.9 ± 0.4 | 0.04 |
| C18:1n-7 (E) | 18:1(11E) | 11E-octadecenoic acid | LMFA01030077 | trans-vaccenic acid | 8.96 | 3.6 ± 0.4 | 0.1 |
| C18:1n-3 | 18:1(15Z) | 15Z-octadecenoic acid | LMFA01030291 | 15Z-octadecenoic acid | 8.79 | 0.35 ± 0.02 | 0.06 |
| C18:1n-4 | 18:1(14Z) | 14Z-octadecenoic acid | LMFA01030882 | 18:1(14Z) | 8.78 | 0.18 ± 0.04 | 0.2 |
| C18:1n-5 | 18:1(13Z) | 13Z-octadecenoic acid | LMFA01030290 | 13Z-octadecenoic acid | 8.78 | 0.54 ± 0.06 | 0.1 |
| C18:1n-10 | 18:1(8Z) | 8Z-octadecenoic acid | LMFA01030070 | cis-8-oleic acid | 8.81 | 1.2 ± 0.2 | 0.2 |
| C18:1n-2 | 18:1(16Z) | 16Z-octadecenoic acid | LMFA01030292 | 16Z-octadecenoic acid | 8.82 | 0.2 ± 0.2 | 1.0 |
| C18:1n-5 (E) | 18:1(13E) | 13E-octadecenoic acid | LMFA01030880 | 18:1(13E) | 8.96 | 1.1 ± 0.6 | 0.5 |
| C18:1n-6 (E) | 18:1(12E) | 12E-octadecenoic acid | LMFA01030079 | trans-12-elaidic acid | 8.96 | 2.8 ± 0.3 | 0.1 |
| C18:1n-12 | 18:1(6Z) | 6Z-octadecenoic acid | LMFA01030066 | Petroselinic acid | 8.96 | 0.4 ± 0.08 | 0.2 |
| C18:1n-9 | 18:1(9Z) | 9Z-octadecenoic acid | LMFA01030002 | Oleic acid | 8.78 | 58 ± 2 | 0.03 |
| C18:1n-11 | 18:1(7Z) | 7Z-octadecenoic acid | LMFA01030068 | 7Z-octadecenoic acid | 8.83 | 0.48 ±<br>0.001 | 0.002 |
| C18:1n-2 (E) | 18:1(16E) | 16E-octadecenoic acid | LMFA01030081 | 16E-octadecenoic acid | 8.95 | 0.4 ± 0.2 | 0.6 |
| C18:1n-3 (E) | 18:1(15E) | 15E-octadecenoic acid | LMFA01030080 | 15E-octadecenoic acid | 8.96 | 0.7 ± 0.1 | 0.2 |
| C18:1n-9 (E) | 18:1(9E) | 9E-octadecenoic acid | LMFA01030073 | 9-elaidic acid | 8.96 | 4.3 ± 0.6 | 0.1 |
| C18:1n-8 (E) | 18:1(10E) | 10E-octadecenoic acid | LMFA01030075 | 10E-octadecenoic acid | 8.96 | 6.7 ± 0.5 | 0.08 |
| C18:1n-10 (E) | 18:1(8E) | 8E-octadecenoic acid | LMFA01030071 | trans-8-elaidic acid | 8.97 | 2.0 ± 0.2 | 0.1 |
| C18:1n-4 (E) | 18:1(14E) | 14E-octadecenoic acid | LMFA01030881 | 18:1(14E) | 8.96 | 1.5 ± 0.2 | 0.1 |
| C18:1n-13 | 18:1(5Z) | 5Z-octadecenoic acid | LMFA01030296 | 5Z-octadecenoic acid | 8.99 | 0.6 ± 0.2 | 0.3 |
| C18:1n-11 (E) | 18:1(7E) | 7E-octadecenoic acid | LMFA01030069 | 7E-octadecenoic acid | 9.04 | 0.5 ± 0.1 | 0.2 |
| C18:1n-12 (E) | 18:1(6E) | 6E-octadecenoic acid | LMFA01030067 | Petroselaidic acid | 9.14 | 0.27 ± 0.04 | 0.2 |
| C18:1n-13 (E) | 18:1(5E) | 5E-octadecenoic acid | LMFA01030065 | Thalictric acid | 9.21 | 0.46 ± 0.04 | 0.09 |
| C19:1n-8 | 19:1(11Z) | 11Z-nonadecenoic acid | LMFA01030887 | 19:1(11Z) | 9.42 | 45 ± 3 | 0.07 |
| C19:1n-9 | 19:1(10Z) | 10Z-nonadecenoic acid | LMFA01030362 | 10Z-nonadecenoic acid | 9.42 | 4.62 ± 0.09 | 0.02 |
| C19:1n-10 | 19:1(9Z) | 9Z-nonadecenoic acid | LMFA01030886 | 19:1(9Z) | 9.42 | 49 ± 3 | 0.06 |
| C19:1n-9 (E) | 19:1(10E) | 10E-nonadecenoic acid | LMFA01030361 | 10E-nonadecenoic acid | 9.61 | 1.9 ± 0.5 | 0.3 |
| C20:1n-7 | 20:1(13Z) | 13Z-eicosenoic acid | LMFA01030367 | Paullinic acid | 9.99 | 6.04 ± 0.01 | 0.002 |
| C20:1n-8 | 20:1(12Z) | 12Z-eicosenoic acid | Not found in<br>LIPID MAPS. | — | 9.99 | 0.9 ± 0.4 | 0.4 |
| C20:1n-9 | 20:1(11Z) | 11Z-eicosenoic acid | LMFA01030085 | cis-gondoic acid | 9.99 | 81.8 ± 0.6 | 0.007 |
| C20:1n-5 | 20:1(15Z) | 15Z-eicosenoic acid | LMFA01030369 | 15Z-eicosenoic acid | 9.99 | 1.17 ± 0.03 | 0.03 |
| C20:1n-6 | 20:1(14Z) | 14Z-eicosenoic acid | LMFA01030087 | 14Z-eicosenoic acid | 9.99 | 1.41 ± 0.03 | 0.02 |
| C20:1n-11 | 20:1(9Z) | 9Z-eicosenoic acid | LMFA01030084 | cis-Gadoleic acid | 10.02 | 6.2 ± 0.4 | 0.07 |
| C20:1n-12 | 20:1(8Z) | 8Z-eicosenoic acid | LMFA01030372 | 8Z-eicosenoic acid | 10.07 | 0.8 ± 0.5 | 0.6 |
| C20:1n-6_(E) | 20:1(14E) | 14E-eicosenoic acid | Not found in LIPID MAPS. | — | 10.15 | 0.14 ± 0.06 | 0.4 |
| C20:1n-13 | 20:1(7Z) | 7Z-eicosenoic acid | LMFA01030858 | 20:1(7Z) | 10.15 | 0.8 ± 0.2 | 0.3 |
| C20:1n-8_(E) | 20:1(12E) | 12E-eicosenoic acid | Not found in LIPID MAPS. | — | 10.15 | 0.3 ± 0.1 | 0.3 |
| C20:1n-9_(E) | 20:1(11E) | 11E-eicosenoic acid | LMFA01030086 | trans-gondoic acid | 10.15 | 0.3 ± 0.1 | 0.4 |
| C20:1n-5_(E) | 20:1(15E) | 15E-eicosenoic acid | Not found in LIPID MAPS. | — | 10.18 | 0.05 ± 0.04 | 0.8 |
| C20:1n-7_(E) | 20:1(13E) | 13E-eicosenoic acid | Not found in LIPID MAPS. | — | 10.16 | 0.1 ± 0.04 | 0.4 |
| C22:1n-9 | 22:1(13Z) | 13Z-docosenoic acid | LMFA01030089 | cis-erucic acid | 11.15 | 86 ± 1 | 0.01 |
| C22:1n-7 | 22:1(15Z) | 15Z-docosenoic acid | LMFA01030402 | 15Z-docosenoic acid | 11.15 | 2.1 ± 0.6 | 0.3 |
| C22:1n-11 | 22:1(11Z) | 11Z-docosenoic acid | LMFA01030088 | cis-cetoleic acid | 11.17 | 2.6 ± 0.8 | 0.3 |
| C22:1n-8_(E) | 22:1(14E) | 14E-docosenoic acid | Not found in LIPID MAPS. | — | 11.32 | 3 ± 1 | 0.4 |
| C22:1n-6_(E) | 22:1(16E) | 16E-docosenoic acid | Not found in LIPID MAPS. | — | 11.32 | 2.9 ± 0.5 | 0.2 |
| C22:1n-9_(E) | 22:1(13E) | 13E-docosenoic acid | LMFA01030090 | trans-brassicic acid | 11.32 | 4 ± 1 | 0.2 |
| C23:1n-14_Br | 23:1(9Z)_Br |  |  |  | 11.41 | 26 ± 7 | 0.3 |
| C23:1n-9 | 23:1(14Z) | 14Z-tricosenoic acid | LMFA01030413 | 14Z-tricosenoic acid | 11.7 | 42 ± 8 | 0.2 |
| C24:1n-9 | 24:1(15Z) | 15Z-tetracosenoic acid | LMFA01030092 | Nervonic acid | 12.24 | 89 ± 5 | 0.06 |
| C24:1n-11 | 24:1(13Z) | 13Z-tetracosenoic acid | Not found in LIPID MAPS. | — | 12.24 | 1 ± 1 | 0.8 |
| C24:1n-5_(E) | 24:1(19E) | 19E-tetracosenoic acid |  |  | 12.42 | 3.2 ± 0.9 | 0.3 |
| C24:1n-6_(E) | 24:1(18E) | 18E-tetracosenoic acid | Not found in LIPID MAPS. | — | 12.42 | 3 ± 2 | 0.7 |
| C24:1n-4_(E) | 24:1(20E) | 20E-tetracosenoic acid | Not found in LIPID MAPS. | — | 12.49 | 2 ± 1 | 0.6 |
| C24:1n-8_(E) | 24:1(16E) | 16E-tetracosenoic acid | Not found in LIPID MAPS. | — | 12.41 | 0.7 ± 0.2 | 0.3 |
| C24:1n-9_(E) | 24:1(15E) | 15E-tetracosenoic acid | LMFA01030093 | trans-selacholeic acid | 12.42 | 1.5 ± 0.7 | 0.5 |
| C16:2n-6,9 | 16:2(7,10) | 7Z,10Z-hexadecadienoic acid | LMFA01030807 | 7Z,10Z-hexadecadienoic acid | 6.69 | — | — |
| C17:2n-5,8 | 17:2(9,12) | 9Z,12Z-heptadecadienoic acid | Not found in LIPID MAPS. | — | 7.3 | 50 ± 20 | 0.4 |
| C17:2n-6,9 | 17:2(8,11) | 8Z,11Z-heptadecadienoic acid | LMFA01031037 | Norlinoleic acid | 7.31 | 50 ± 20 | 0.4 |
| C18:2n-6,9 | 18:2(9,12) | 9Z,12Z-octadecadienoic acid | LMFA01030120 | Linoleic acid | 7.96 | 95 ± 2 | 0.02 |
| C18:2n-2,9 | 18:2(9,16) | 9Z,16Z-octadecadienoic acid | Not found in LIPID MAPS. | — | 8.06 | — | — |
| C18:2n-3,9 | 18:2(9,15) | 9Z,15Z-octadecadienoic acid | Not found in LIPID MAPS. | — | 8.06 | — | — |
| C18:2n-4,9 | 18:2(9,14) | 9Z,14Z-octadecadienoic acid | Not found in LIPID MAPS. | — | 8.06 | — | — |
| C18:2n-5,9 | 18:2(9,13) | 9Z,13Z-octadecadienoic acid | Not found in LIPID MAPS. | — | 8.07 | — | — |
| C18:2n-6,12 | 18:2(6,12) | 6Z,12Z-octadecadienoic acid | Not found in LIPID MAPS. | — | 8.22 | — | — |
| C18:2n-3,10 | 18:2(8,15) | 8Z,15Z-octadecadienoic acid | Not found in LIPID MAPS. | — | 8.1 | — | — |
| C18:2n-6,13 | 18:2(5,12) | 5Z,12Z-octadecadienoic acid | LMFA01030110 | 5Z,12Z-octadecadienoic acid | 8.32 | — | — |
| C18:2n-6,8 | 18:2(10,12) | 10Z,12Z-octadecadienoic acid | LMFA01030124 | 10Z,12Z-octadecadienoic acid | 8.32 | — | — |
| C18:2n-6,9_(E) | 18:2(9,12) | 9Z,12E-octadecadienoic acid | LMFA02000352 | — | 8.17 | — | — |
| C18:2n-7,9 | 18:2(9,11) | 9Z,11Z-octadecadienoic acid | LMFA01030117 | Ricinenic acid | 8.17 | — | — |
| C18:2n-6,10 | 18:2(8,12) | 8Z,12Z-octadecadienoic acid | Not found in LIPID MAPS. | — | 8.17 | — | — |
| C18:2n-6,12_(E) | 18:2(6,12) | 6Z,12E-octadecadienoic acid | Not found in LIPID MAPS. | — | 8.31 | — | — |
| C19:2n-6,9 | 19:2(10,13) | 10Z,13Z-nonadecadienoic acid | LMFA01030129 | 10Z,13Z-nonadecadienoic acid | 8.58 | 50 ± 10 | 0.2 |
| C19:2n-7,10 | 19:2(9,12) | 9Z,12Z-nonadecadienoic acid | Not found in LIPID MAPS. | — | 8.58 | 50 ± 10 | 0.2 |
| C20:2n-6,9 | 20:2(11,14) | 11Z,14Z-eicosadienoic acid | LMFA01031043 | Dihomolinoleic acid | 9.19 | 88 ± 1 | 0.01 |
| C20:2n-9,12 | 20:2(8,11) | 8Z,11Z-eicosadienoic acid | LMFA01030377 | 8Z,11Z-eicosadienoic acid | 9.27 | 6.7 ± 0.5 | 0.07 |
| C20:2n-9,15 | 20:2(5,11) | 5Z,11Z-eicosadienoic acid | LMFA01030865 | Keteleeronic acid | 9.4 | 5.6 ± 0.8 | 0.1 |
| C22:2n-6,9 | 22:2(13,16) | 13Z,16Z-docosadienoic acid | LMFA01030405 | 13Z,16Z-docosadienoic acid | 10.35 | 69 ± 3 | 0.04 |
| C22:2n-9,15 | 22:2(7,13) | 7Z,13Z-docosadienoic acid | LMFA04000061 | 22:2(7Z,13Z) | 10.43 | 31 ± 3 | 0.1 |
| C24:2n-6,9 | 24:2(15,18) | 15Z,18Z-tetracosadienoic acid | LMFA01031071 | 15Z,18Z-Tetracosadienoic acid | 11.47 | — | — |
| C24:2n-7,9 | 24:2(15,17) | 15Z,17Z-tetracosadienoic acid | Not found in LIPID MAPS. | — | 11.61 | — | — |
| C24:2n-5,9 | 24:2(15,19) | 15Z,19Z-tetracosadienoic acid | Not found in LIPID MAPS. | — | 11.62 | — | — |
| C16:3n-6,9,12 | 16:3(4,7,10) | 4Z,7Z,10Z-hexadecatrienoic acid | Not found in LIPID MAPS. | — | 6.25 | — | — |
| C18:3n-3,6,9 | 18:3(9,12,15) | 9Z,12Z,15Z-octadecatrienoic acid | LMFA01030152 | alpha-Linolenic acid | 7.24 | 85.4 ± 0.9 | 0.01 |
| C18:3n-6,9,12 | 18:3(6,9,12) | 6Z,9Z,12Z-octadecatrienoic acid | LMFA01030141 | gamma-Linolenic acid | 7.37 | 13.5 ± 0.8 | 0.06 |
| C18:3n-3,6,9_E | 18:3(9,12,15) | 9Z,12Z,15E-octadecatrienoic acid | LMFA01030353 | 9Z,12Z,15E-octadecatrienoic acid | 7.45 | 0.51 ± 0.05 | 0.1 |
| C18:3n-6,9,12_E | 18:3(6,9,12) | 6Z,9Z,12E-octadecatrienoic acid | Not found in LIPID MAPS. | — | 7.54 | 0.59 ± 0.06 | 0.1 |
| C20:3n-6,9,12 | 20:3(8,11,14) | 8Z,11Z,14Z-eicosatrienoic acid | LMFA01030387 | — | 8.5 | 89.8 ± 0.5 | 0.006 |
| C20:3n-6,9,15 | 20:3(5,11,14) | 5Z,11Z,14Z-eicosatrienoic acid | LMFA01030380 | Sciadonic acid | 8.59 | 9.9 ± 0.3 | 0.03 |
| C20:3n-9,12,15 | 20:3(5,8,11) | 5Z,8Z,11Z-eicosatrienoic acid | LMFA01030381 | Mead acid | 8.77 | 0.3 ± 0.2 | 0.7 |
| C22:3n-6,9,15 | 22:3(7,13,16) | 7Z,13Z,16Z-docosatrienoic acid | LMFA04000062 | 22:3(7Z,13Z,16Z) | 9.63 | 10 ± 10 | 0.8 |
| C22:3n-6,9,12 | 22:3(10,13,16) | 10Z,13Z,16Z-docosatrienoic acid | Not found in LIPID MAPS. | — | 9.63 | 39 ± 3 | 0.08 |
| C22:3n-9,12,15 | 22:3(7,10,13) | 7Z,10Z,13Z-docosatrienoic acid | LMFA01030411 | Dihomo Mead's acid | 9.77 | 47 ± 8 | 0.2 |
| C18:4n-3,6,9,12 | 18:4(6,9,12,15) | 6Z,9Z,12Z,15Z-octadecatetraenoic acid | LMFA01030357 | Stearidonic acid | 6.65 | — | — |
| C20:4n-6,9,12,15 | 20:4(5,8,11,14) | 5Z,8Z,11Z,14Z-eicosatetraenoic acid | LMFA01030001 | Arachidonic acid | 8.02 | 97.4 ± 0.2 | 0.002 |
| C20:4n-3,6,9,12 | 20:4(8,11,14,17) | 8Z,11Z,14Z,17Z-eicosatetraenoic acid | LMFA01030818 | omega-3-Arachidonic acid | 7.81 | 2.6 ± 0.2 | 0.08 |
| C20:4n-3,6,9,15 | 20:4(5,11,14,17) | 5Z,11Z,14Z,17Z-eicosatetraenoic acid | LMFA01030394 | Juniperonic acid | 7.88 | < 1 | — |
| C22:4n-6,9,12,15 | 22:4(7,10,13,16) | 7Z,10Z,13Z,16Z-docosatetraenoic acid | LMFA01030178 | Adrenic Acid | 9.03 | — | — |
| C24:4n-6,9,12,15 | 24:4(9,12,15,18) | 9Z,12Z,15Z,18Z-tetracosatetraenoic acid | LMFA01030819 | 9Z,12Z,15Z,18Z-Tetracosatetraenoic acid | 10.06 | — | — |
| C20:5n-3,6,9,12,15 | 20:5(5,8,11,14,17) | 5Z,8Z,11Z,14Z,17Z-eicosapentaenoic acid | LMFA01030759 | EPA | 7.32 | — | — |
| C22:5n-3,6,9,12,15 | 22:5(7,10,13,16,19) | 7Z,10Z,13Z,16Z,19Z-docosapentaenoic acid | LMFA04000044 | DPA | 8.34 | 45.5 ± 0.1 | 0.002 |
| C22:5n-6,9,12,15,18 | 22:5(4,7,10,13,16) | 4Z,7Z,10Z,13Z,16Z-docosapentaenoic acid | LMFA04000064 | 22:5(4Z,7Z,10Z,13Z,16Z) | 8.69 | 54.5 ± 0.1 | 0.002 |
| C24:5n-3,6,9,12,15 | 24:5(9,12,15,18,21) | 9Z,12Z,15Z,18Z,21Z-tetracosapentaenoic acid | LMFA01030821 | 9Z,12Z,15Z,18Z,21Z-tetracosapentaenoic acid | 9.42 | 68 ± 7 | 0.1 |
| C24:5n-6,9,12,15,18 | 24:5(6,9,12,15,18) | 6Z,9Z,12Z,15Z,18Z-tetracosapentaenoic acid | LMFA01030820 | 6Z,9Z,12Z,15Z,18Z-tetracosapentaenoic acid | 9.57 | 33 ± 7 | 0.2 |
| C22:6n-3,6,9,12,15,18 | 22:6(4,7,10,13,16,19) | 4Z,7Z,10Z,13Z,16Z,19Z-docosahexaenoic acid | LMFA01030185 | DHA | 8.01 | — | — |
| C24:6n-3,6,9,12,15,18 | 24:6(6,9,12,15,18,21) | 6Z,9Z,12Z,15Z,18Z,21Z-tetracosahexaenoic acid | LMFA01030822 | THA | 8.88 | — | — |

### 3.5 Fatty acids in vernix caseosa

Sample collection, lipid extraction, hydrolysis, and fixed charge derivatization (4-I-AMPP) were described previously. The derivatized fatty acids in Methanol were kept at -18°C prior to analysis with the methods described in this study. For each direct infusion ESI-MS measurement and LC-OzID-MS analyses (both untargeted and targeted analysis) 10 μL were injected.

**Figure S25:**
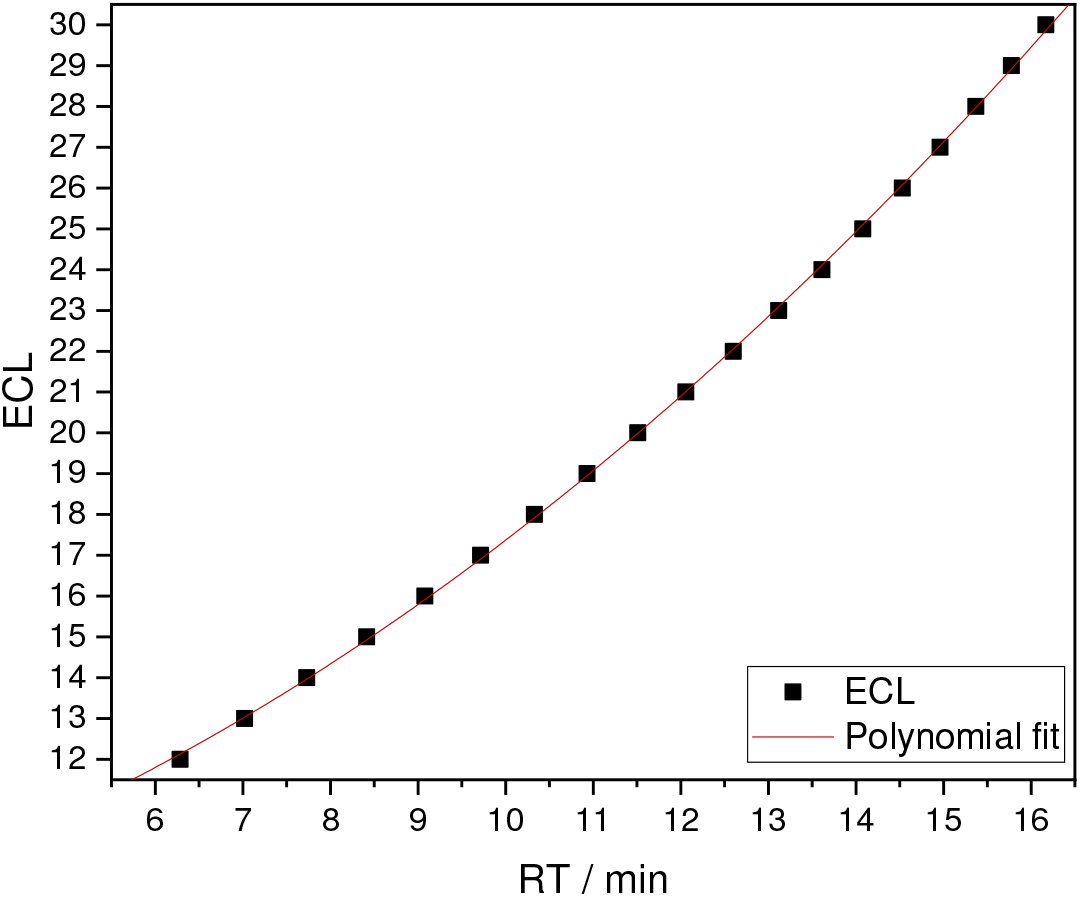
Polynomial fit of equivalent chain lengths vs. retention times of saturated fatty acids (straight chain) in Vernix caseosa (ECL = 0.0621 * (RT)^2 + 0.39907 * RT + 7.1711).

**Figure S26:**
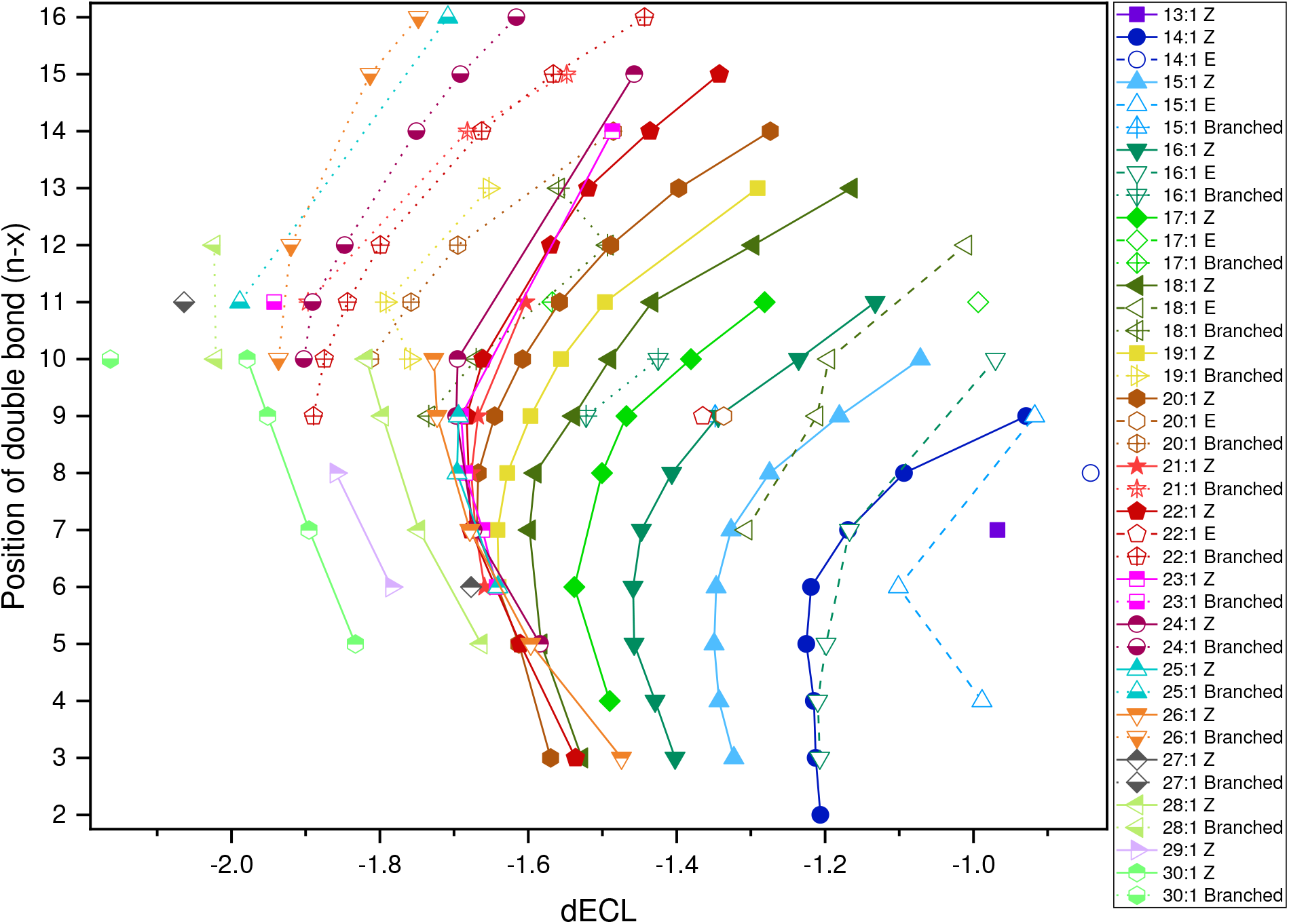
Double bond positions of 4-I-AMPP derivatized fatty acid isomers found in vernix caseosa plotted against their dECL values as determined with the polynomial fit shown in Figure S25.

**Figure S27:**
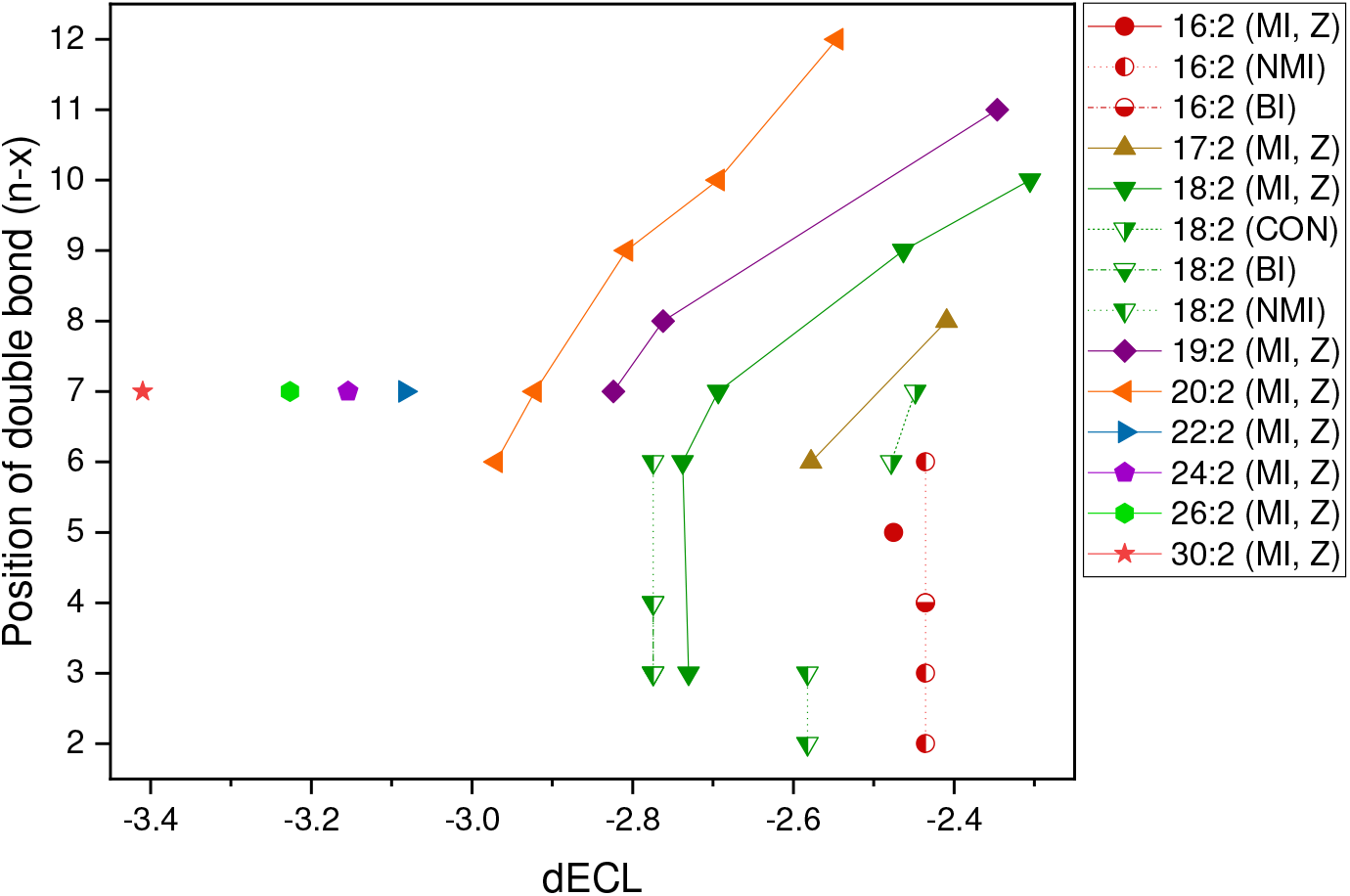
Analysis of differential equivalent chain lengths (dECL) for bisunsaturated fatty acids in vernix caseosa extracts. Non-methylene interrupted fatty acids are assigned as conjugated (CON), butylene-interrupted (BU) or other non-methylene interrupted fatty acids (NMI). Double bond configuration cannot be established unequivocally for all fatty acid species, but the analysis allows a tentative assignment as cis for the majority of fatty acids that are observed. For determination of ECL and dECL values the equation (ECL = 0.0621 * (RT)^2 + 0.39907 * RT + 7.1711) was determined from retention times of saturated fatty acids in the same sample.

**Figure S28:**
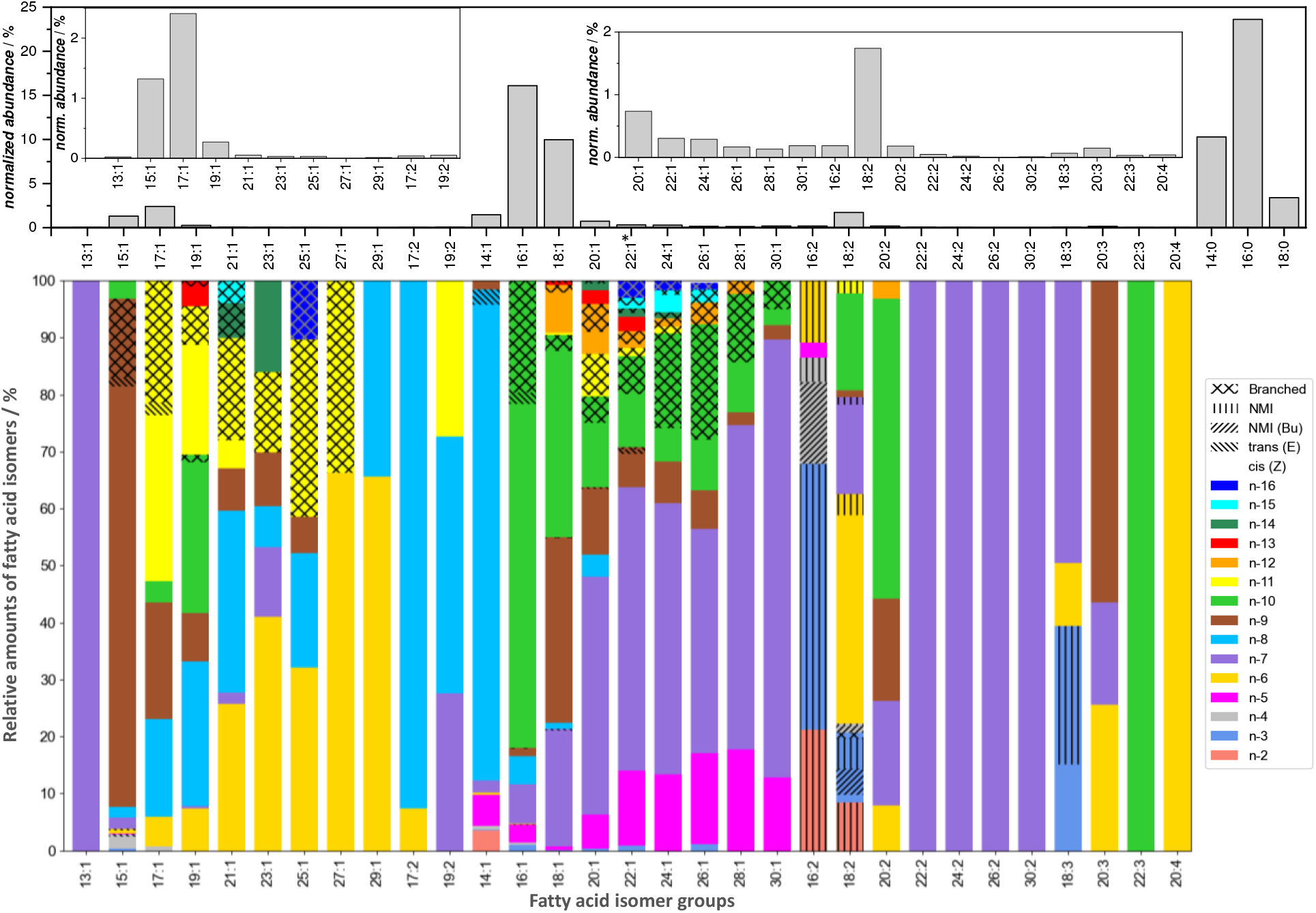
Unsaturated fatty acids and their relative abundance in vernix caseosa. A lipid extract of vernix caseosa was hydrolyzed and derivatized with 4-I-AMPP prior to UPLC-OzID-MS analysis. Positions of branch points cannot be determined by Ozone-induced dissociation and are not reported here. Above: Relative quantification of fatty acid isomer groups without discrimination of isomers. Below: Relative quantification of fatty acids for each isomer group based on LC-OzID-MS and LC-OzID-MS/MS. *The plastics additive erucamide and erucic acid derived from it are detected both in the vernix caseosa extract as well as the associated Process Blank. The data shown here is subtracted with the amounts detected in the Process Blank.

**Table S6:** Identified fatty acid species in a vernix caseosa lipid extract (after hydrolysis and fixed-charge derivatization with 4-I-AMPP). Fatty isomers that, to the best of our knowledge, have not been reported before, are highlighted with a yellow background, while those that we also discovered in pooled human plasma are highlighted with a grey background.

| Fatty acid (n-x) | FA shorthand | Systematic name | LipidMAPS ID | Common Name | RT / min | Confidence of identification (High: H; Low: L) | Relative isomer abundance / % of each isomer group | Estimated relative fatty acid abundance / % of the total fatty acid content |
| --- | --- | --- | --- | --- | --- | --- | --- | --- |
| C12:0_br | 12:0_br | = | = | = | 5.84 | H | 4.7 | 0.035 |
| C12:0_br | 12:0_br | = | = | = | 6.05 | H | 10.7 | 0.079 |
| C12:0_br | 12:0_br | = | = | = | 6.15 | H | 33.8 | 0.25 |
| C12:0 | 12:0 | Dodecanoic acid | LMFA01010012 | Lauric acid | 6.29 | H | 50.9 | 0.38 |
| C13:0_br | 13:0_br | = | = | = | 6.57 | H | 2.1 | 0.037 |
| C13:0_br | 13:0_br | = | = | = | 6.66 | H | 21.9 | 0.38 |
| C13:0_br | 13:0_br | = | = | = | 6.80 | H | 53.6 | 0.94 |
| C13:0_br | 13:0_br | = | = | = | 6.89 | H | 2.3 | 0.040 |
| C13:0 | 13:0 | Tridecanoic acid | LMFA01010013 | Tridecylic acid | 7.02 | H | 20.2 | 0.35 |
| C14:0_br | 14:0_br | = | = | = | 7.60 | H | 41.4 | 4.3 |
| C14:0 | 14:0 | Tetradecanoic acid | LMFA01010014 | Myristic acid | 7.73 | H | 58.6 | 6.1 |
| C15:0_br | 15:0_br | = | = | = | 8.05 | H | 3.4 | 0.36 |
| C15:0_br | 15:0_br | = | = | = | 8.21 | H | 38.9 | 4.1 |
| C15:0_br | 15:0_br | = | = | = | 8.28 | H | 5.4 | 0.57 |
| C15:0 | 15:0 | Pentadecanoic acid | LMFA01010015 | Pentadecylic acid | 8.42 | H | 52.3 | 5.5 |
| C16:0_br | 16:0_br | = | = | = | 8.86 | H | 4.6 | 1.1 |
| C16:0_br | 16:0_br | = | = | = | 8.95 | H | 16.8 | 4.0 |
| C16:0 | 16:0 | Hexadecanoic acid | LMFA01010001 | Palmitic acid | 9.08 | H | 78.7 | 18.6 |
| C17:0_br | 17:0_br | = | = | = | 9.38 | H | 12.1 | 0.53 |
| C17:0_br | 17:0_br | = | = | = | 9.52 | H | 54.1 | 2.4 |
| C17:0_br | 17:0_br | = | = | = | 9.59 | H | 2.2 | 0.097 |
| C17:0 | 17:0 | Heptadecanoic acid | LMFA01010017 | Margaric acid | 9.72 | H | 31.6 | 1.4 |
| C18:0_br | 18:0_br | = | = | = | 10.20 | H | 14.4 | 0.49 |
| C18:0 | 18:0 | Octadecanoic acid | LMFA01010041 | Stearic acid | 10.33 | H | 85.6 | 2.9 |
| C19:0_br | 19:0_br | = | = | = | 10.75 | H | n.a. |  |
| C19:0_br | 19:0_br | = | = | = | 10.80 | H | n.a. |  |
| C19:0 | 19:0 | Nonadecanoic acid | LMFA01010019 | Nonadecylic acid | 10.93 | H | n.a. (IS) |  |
| C20:0_br | 20:0_br | = | = | = | 11.39 | H | 73.1 | 1.1 |
| C20:0 | 20:0 | Eicosanoic acid | LMFA01010020 | Arachidic acid | 11.51 | H | 26.9 | 0.39 |
| C21:0_br | 21:0_br | = | = | = | 11.90 | H | 89.4 | 0.42 |
| C21:0 | 21:0 | Heneicosanoic acid | LMFA01010021 | Heneicosylic acid | 12.06 | H | 10.6 | 0.05 |
| C22:0_br | 22:0_br | = | = | = | 12.49 | H | 77.7 | 1.1 |
| C22:0 | 22:0 | Docosanoic acid | LMFA01010022 | Behenic acid | 12.60 | H | 22.3 | 0.31 |
| C23:0_br | 23:0_br | = | = | = | 12.98 | H | 75.2 | 0.49 |
| C23:0 | 23:0 | Tricosanoic acid | LMFA01010023 | Tricosylic acid | 13.12 | H | 24.8 | 0.16 |
| C24:0_br | 24:0_br | = | = | = | 13.50 | H | 67.5 | 1.8 |
| C24:0 | 24:0 | Tetracosanoic acid | LMFA01010024 | Lignoceric acid | 13.61 | H | 32.5 | 0.85 |
| C25:0_br | 25:0_br | = | = | = | 13.95 | H | 78.9 | 0.76 |
| C25:0 | 25:0 | Pentacosanoic acid | LMFA01010025 | Hyenic acid | 14.08 | H | 21.1 | 0.20 |
| C26:0_br | 26:0_br | = | = | = | 14.43 | H | 63.3 | 0.71 |
| C26:0 | 26:0 | Hexacosanoic acid | LMFA01010026 | Cerotic acid | 14.53 | H | 36.7 | 0.41 |
| C27:0_br | 27:0_br | = | = | = | 14.83 | H | 76.9 | 0.14 |
| C27:0 | 27:0 | Heptacosanoic acid | LMFA01010027 | Carboceric acid | 14.96 | H | 23.1 | 0.042 |
| C28:0_br | 28:0_br | = | = | = | 15.28 | H | 45.8 | 0.087 |
| C28:0 | 28:0 | Octacosanoic acid | LMFA01010028 | Montanic acid | 15.37 | H | 54.2 | 0.10 |
| C29:0_br | 29:0_br | = | = | = | 15.66 | H | 71.2 | 0.028 |
| C29:0 | 29:0 | Nonacosanoic acid | LMFA01010029 | Nonacosanoic acid | 15.78 | H | 28.8 | 0.011 |
| C30:0_br | 30:0_br | = | = | = | 16.08 | H | 48.0 | 0.024 |
| C30:0 | 30:0 | Triacontanoic acid | LMFA01010030 | Melissic acid | 16.17 | H | 52.0 | 0.027 |
| C13:1n-7 | 13:1(6Z) | 6Z-tridecenoic acid | Not found in LIPID MAPS. | = | 6.19 | H | 100 | 0.024 |
| C14:1n-4 | 14:1(10Z) | 10Z-tetradecenoic acid | Not found in LIPID MAPS. | = | 6.82 | H | 0.63 | 0.0092 |
| C14:1n-5 | 14:1(9Z) | 9Z-tetradecenoic acid | LMFA01030051 | Myristoleic acid | 6.82 | H | 5.5 | 0.081 |
| C14:1n-6 | 14:1(8Z) | 8Z-tetradecenoic acid | LMFA01030050 | 8Z-tetradecenoic acid | 6.82 | H | 0.34 | 0.0050 |
| C14:1n-2 | 14:1(12Z) | 12Z-tetradecenoic acid | Not found in LIPID MAPS. | = | 6.82 | H | 3.5 | 0.051 |
| C14:1n-3 | 14:1(11Z) | 11Z-tetradecenoic acid | LMFA01030770 | cis-tetradec-11-enoic acid | 6.84 | H | 0.21 | 0.0031 |
| C14:1n-7 | 14:1(7Z) | 7Z-tetradecenoic acid | LMFA01030249 | 7Z-tetradecenoic acid | 6.84 | H | 2.2 | 0.032 |
| C14:1n-9 | 14:1(5Z) | 5Z-tetradecenoic acid | LMFA01030248 | Physeteric acid | 6.99 | H | 1.5 | 0.022 |
| C14:1n-8 | 14:1(6Z) | 6Z-tetradecenoic acid | LMFA01031183 | 6Z-tetradecenoic acid | 6.92 | H | 84 | 1.2 |
| C14:1n-8_(E) | 14:1(6E) | 6E-tetradecenoic acid | Not found in LIPID MAPS. | = | 7.11 | H | 2.6 | 0.038 |
| C15:1n-5 | 15:1(10Z) | 10Z-pentadecenoic acid | LMFA01030259 | 10Z-pentadecenoic acid | 7.49 | H | 0.096 | 0.0013 |
| C15:1n-3 | 15:1(12Z) | 12Z-pentadecenoic acid | Not found in LIPID MAPS. | = | 7.5 | H | 0.38 | 0.0050 |
| C15:1n-4 | 15:1(11Z) | 11Z-pentadecenoic acid | Not found in LIPID MAPS. | = | 7.49 | H | 2.1 | 0.028 |
| C15:1n-6 | 15:1(9Z) | 9Z-pentadecenoic acid | LMFA01030866 | 15:1(9Z) | 7.49 | H | 0.53 | 0.0070 |
| C15:1n-9_Branched | 15:1(6)_Br |  |  |  | 7.49 | H | 13.7 | 0.18 |
| C15:1n-7 | 15:1(8Z) | 8Z-pentadecenoic acid | Not found in LIPID MAPS. | = | 7.49 | H | 2 | 0.026 |
| C15:1n-8 | 15:1(7Z) | 7Z-pentadecenoic acid | LMFA01030855 | 15:1(7Z) | 7.56 | H | 1.8 | 0.024 |
| C15:1n-9 | 15:1(6Z) | 6Z-pentadecenoic acid | LMFA01031184 | 6Z-pentadecenoic acid | 7.62 | H | 73.8 | 0.97 |
| C15:1n-4_(E) | 15:1(11E) | 11E-pentadecenoic acid | Not found in<br>LIPID MAPS. | = | 7.66 | H | 0.4 | 0.0053 |
| C15:1n-6_(E) | 15:1(9E) | 9E-pentadecenoic acid | Not found in<br>LIPID MAPS. | = | 7.7 | H | 0.32 | 0.0042 |
| C15:1n-9_(E) | 15:1(6E) | 6E-pentadecenoic acid | Not found in<br>LIPID MAPS. | = | 7.81 | H | 1.6 | 0.021 |
| C15:1n-10 | 15:1(5Z) | 5Z-pentadecenoic acid | LMFA01031197 | 5Z-pentadecenoic acid | 7.7 | H | 3.2 | 0.042 |
| C16:1n-9_Branched | 16:1(7)_Br |  | = | = | 8.11 | H | 0.09 | 0.015 |
| C16:1n-5 | 16:1(11Z) | 11Z-hexadecenoic acid | LMFA01030262 | cis-Palmitvaccenic acid | 8.19 | H | 3.1 | 0.50 |
| C16:1n-7 | 16:1(9Z) | 9Z-hexadecenoic acid | LMFA01030056 | cis-9-palmitoleic acid | 8.19 | H | 6.9 | 1.1 |
| C16:1n-3 | 16:1(13Z) | 13Z-hexadecenoic acid | LMFA01030264 | 13Z-hexadecenoic acid | 8.19 | H | 1 | 0.16 |
| C16:1n-6 | 16:1(10Z) | 10Z-hexadecenoic acid | LMFA01030058 | cis-10-palmitoleic acid | 8.16 | H | 0.11 | 0.018 |
| C16:1n-8 | 16:1(8Z) | 8Z-hexadecenoic acid | LMFA01031081 | 8Z-Hexadecenoic acid | 8.19 | H | 4.8 | 0.77 |
| C16:1n-4 | 16:1(12Z) | 12Z-hexadecenoic acid | Not found in<br>LIPID MAPS. | = | 8.19 | H | 0.37 | 0.060 |
| C16:1n-10_Branched | 16:1(6)_Br |  |  |  | 8.19 | H | 19.6 | 3.2 |
| C16:1n-9 | 16:1(7Z) | 7Z-hexadecenoic acid | LMFA01030766 |  | 8.2 | H | 1.4 | 0.23 |
| C16:1n-3_(E) | 16:1(13E) | 13E-hexadecenoic acid | Not found in<br>LIPID MAPS. | = | 8.33 | H | 1 | 0.16 |
| C16:1n-4_(E) | 16:1(12E) | 12E-hexadecenoic acid | Not found in<br>LIPID MAPS. | = | 8.32 | H | 0.37 | 0.060 |
| C16:1n-10 | 16:1(6Z) | 6Z-hexadecenoic acid | LMFA01030267 | Sapienic acid | 8.3 | H | 60.4 | 9.7 |
| C16:1n-5_(E) | 16:1(11E) | 11E-hexadecenoic acid | LMFA01030261 | Lycopodic acid | 8.32 | H | 3.1 | 0.50 |
| C16:1n-7_(E) | 16:1(9E) | 9E-hexadecenoic acid | LMFA01030057 | trans-9-palmitoleic acid | 8.34 | H | 6.9 | 1.1 |
| C16:1n-11 | 16:1(5Z) | 5Z-hexadecenoic acid | LMFA01030854 | 16:1(5Z) | 8.33 | H | 0.054 | 0.0087 |
| C16:1n-10_(E) | 16:1(6E) | 6E-hexadecenoic acid | LMFA01031082 | 6E-Hexadecenoic acid | 8.48 | H | 2.0 | 0.32 |
| C17:1n-11_Branched | 17:1(6)_Br |  | = | = | 8.77 | H | 21.4 | 0.52 |
| C17:1n-6 | 17:1(11Z) | 11Z-heptadecenoic acid | LMFA01030856 | 17:1(11Z) | 8.77 | H | 4.9 | 0.12 |
| C17:1n-8 | 17:1(9Z) | 9Z-heptadecenoic acid | LMFA01030060 | Margaroleic acid | 8.77 | H | 17 | 0.41 |
| C17:1n-9 | 17:1(8Z) | 8Z-heptadecenoic acid | LMFA01030288 | Civetic acid | 8.81 | H | 21 | 0.51 |
| C17:1n-4 | 17:1(13Z) | 13Z-heptadecenoic acid | Not found in<br>LIPID MAPS. | = | 8.81 | H | 0.67 | 0.016 |
| C17:1n-10 | 17:1(7Z) | 7Z-heptadecenoic acid | LMFA01030286 | 7Z-heptadecenoic acid | 8.86 | H | 3.7 | 0.089 |
| C17:1n-11 | 17:1(6Z) | 6Z-heptadecenoic acid | LMFA01031186 | 6Z-heptadecenoic acid | 8.96 | H | 29.2 | 0.70 |
| C17:1n-11_(E) | 17:1(6E) | 6E-heptadecenoic acid | Not found in<br>LIPID MAPS. | = | 9.14 | H | 2.2 | 0.053 |
| C18:1n-10_Branched | 18:1(8)_Br |  | = | = | 9.37 | H | 2.8 | 0.28 |
| C18:1n-7 | 18:1(11Z) | 11Z-octadecenoic acid | LMFA01030076 | cis-vaccenic acid | 9.44 | H | 20 | 2.0 |
| C18:1n-9_Branched | 18:1(9)_Br |  | = | = | 9.34 | H | 0.13 | 0.013 |
| C18:1n-12_Branched | 18:1(6)_Br |  | = | = | 9.44 | H | 1.4 | 0.14 |
| C18:1n-5 | 18:1(13Z) | 13Z-octadecenoic acid | LMFA01030290 | 13Z-octadecenoic acid | 9.43 | H | 0.66 | 0.066 |
| C18:1n-3 | 18:1(15Z) | 15Z-octadecenoic acid | LMFA01030291 | 15Z-octadecenoic acid | 9.42 | H | 0.12 | 0.012 |
| C18:1n-8 | 18:1(10Z) | 10Z-octadecenoic acid | LMFA01030074 | cis-10-oleic acid | 9.44 | H | 1 | 0.10 |
| C18:1n-9 | 18:1(9Z) | 9Z-octadecenoic acid | LMFA01030002 | Oleic acid | 9.44 | H | 32.4 | 3.2 |
| C18:1n-10 | 18:1(8Z) | 8Z-octadecenoic acid | LMFA01030070 | cis-8-oleic acid | 9.44 | H | 32.7 | 3.3 |
| C18:1n-11 | 18:1(7Z) | 7Z-octadecenoic acid | LMFA01030068 | 7Z-octadecenoic acid | 9.49 | H | 0.35 | 0.035 |
| C18:1n-13_Branched | 18:1(5)_Br |  | = | = | 9.44 | H | 0.23 | 0.023 |
| C18:1n-7_(E) | 18:1(11E) | 11E-octadecenoic acid | LMFA01030077 | trans-vaccenic acid | 9.59 | H | 20 | 2.0 |
| C18:1n-12 | 18:1(6Z) | 6Z-octadecenoic acid | LMFA01030066 | Petroselinic acid | 9.59 | H | 7.0 | 0.70 |
| C18:1n-13 | 18:1(5Z) | 5Z-octadecenoic acid | LMFA01030296 | 5Z-octadecenoic acid | 9.68 | H | 0.23 | 0.023 |
| C18:1n-10_(E) | 18:1(8E) | 8E-octadecenoic acid | LMFA01030071 | trans-8-elaidic acid | 9.68 | H | 2.8 | 0.28 |
| C18:1n-12_(E) | 18:1(6E) | 6E-octadecenoic acid | LMFA01030067 | Petroselaidic acid | 9.76 | H | 1.4 | 0.14 |
| C19:1n-11_Branched | 19:1(8)_Br |  | = | = | 9.91 | H | 6.7 | 0.018 |
| C19:1n-10_Branched | 19:1(9)_Br |  | = | = | 9.94 | H | 1.4 | 0.0038 |
| C19:1n-6 | 19:1(13Z) | 13Z-nonadecenoic acid | LMFA01030908 | 19:1(13Z) | 9.99 | H | 7.4 | 0.020 |
| C19:1n-8 | 19:1(11Z) | 11Z-nonadecenoic acid | LMFA01030887 | 19:1(11Z) | 10.04 | H | 25 | 0.068 |
| C19:1n-9 | 19:1(10Z) | 10Z-nonadecenoic acid | LMFA01030362 | 10Z-nonadecenoic acid | 10.04 | H | 8.4 | 0.023 |
| C19:1n-7 | 19:1(12Z) | 12Z-nonadecenoic acid | LMFA01030857 | 19:1(12Z) | 10 | H | 0.36 | 0.001 |
| C19:1n-13_Branched | 19:1(6)_Br |  | = | = | 9.99 | H | 1 | 0.0027 |
| C19:1n-10 | 19:1(9Z) | 9Z-nonadecenoic acid | LMFA01030886 | 19:1(9Z) | 10.04 | H | 1.4 | 0.0038 |
| C19:1n-11 | 19:1(8Z) | 8Z-nonadecenoic acid | Not found in<br>LIPID MAPS. | = | 10.04 | H | 6.7 | 0.018 |
| C19:1n-13 | 19:1(6Z) | 6Z-nonadecenoic acid | LMFA01030885 | 19:1(6Z) | 10.21 | H | 1 | 0.0027 |
| C20:1n-11_Branched | 20:1(9)_Br |  | = | = | 10.55 | H | 7.1 | 0.052 |
| C20:1n-10_Branched | 20:1(10)_Br |  | = | = | 10.52 | H | 4.7 | 0.035 |
| C20:1n-7 | 20:1(13Z) | 13Z-eicosenoic acid | LMFA01030367 | Paullinic acid | 10.58 | H | 42 | 0.31 |
| C20:1n-12_Branched | 20:1(8)_Br |  | = | = | 10.58 | H | 4.9 | 0.036 |
| C20:1n-8 | 20:1(12Z) | 12Z-eicosenoic acid | Not found in<br>LIPID MAPS. | = | 10.58 | H | 3.8 | 0.028 |
| C20:1n-9 | 20:1(11Z) | 11Z-eicosenoic acid | LMFA01030085 | cis-gondoic acid | 10.58 | H | 12 | 0.088 |
| C20:1n-5 | 20:1(15Z) | 15Z-eicosenoic acid | LMFA01030369 | 15Z-eicosenoic acid | 10.58 | H | 6 | 0.044 |
| C20:1n-3 | 20:1(17Z) | 17Z-eicosenoic acid | Not found in<br>LIPID MAPS. | = | 10.6 | H | 0.31 | 0.0023 |
| C20:1n-10 | 20:1(10Z) | 10Z-eicosenoic acid | LMFA01031093 | 10Z-Eicosenoic acid | 10.59 | H | 4.7 | 0.035 |
| C20:1n-11 | 20:1(9Z) | 9Z-eicosenoic acid | LMFA01030084 | cis-Gadoleic acid | 10.64 | H | 7.1 | 0.052 |
| C20:1n-12 | 20:1(8Z) | 8Z-eicosenoic acid | LMFA01030372 | 8Z-eicosenoic acid | 10.68 | H | 4.9 | 0.036 |
| C20:1n-14_Branched | 20:1(6)_Br |  |  |  | 10.68 | H | 0.52 | 0.0038 |
| C20:1n-13 | 20:1(7Z) | 7Z-eicosenoic acid | LMFA01030858 | 20:1(7Z) | 10.74 | H | 2.4 | 0.018 |
| C20:1n-14 | 20:1(6Z) | 6Z-eicosenoic acid | LMFA01031098 | 6Z-Eicosenoic acid | 10.8 | H | 0.52 | 0.0038 |
| C20:1n-9 (E) | 20:1(11E) | 11E-eicosenoic acid | LMFA01030086 | trans-gondoic acid | 10.77 | H | 12 | 0.088 |
| C21:1n-11_Branched | 21:1(10)_Br |  |  |  | 11.02 | H | 18 | 0.0097 |
| C21:1n-6 | 21:1(15Z) | 15Z-heneicosenoic acid | Not found in LIPID MAPS. |  | 11.15 | H | 26 | 0.014 |
| C21:1n-8 | 21:1(13Z) | 13Z-heneicosenoic acid | Not found in LIPID MAPS. |  | 11.15 | H | 32 | 0.017 |
| C21:1n-9 | 21:1(12Z) | 12Z-heneicosenoic acid | LMFA01030398 | 12Z-heneicosenoic acid | 11.15 | H | 7.5 | 0.004 |
| C21:1n-14_Branched | 21:1(7)_Br |  |  |  | 11.14 | H | 6.1 | 0.0033 |
| C21:1n-7 | 21:1(14Z) | 14Z-heneicosenoic acid | Not found in LIPID MAPS. |  | 11.15 | H | 2 | 0.0011 |
| C21:1n-15_Branched | 21:1(6)_Br |  |  |  | 11.18 | H | 3.9 | 0.0021 |
| C21:1n-11 | 21:1(10Z) | 10Z-heneicosenoic acid | Not found in LIPID MAPS. |  | 11.18 | H | 18 | 0.0097 |
| C22:1n-9_Branched | 22:1(13)_Br |  |  |  | 11.59 | H | 0.37 | 0.0011 |
| C22:1n-11_Branched | 22:1(11)_Br |  |  |  | 11.66 | H | 1.4 | 0.0043 |
| C22:1n-12_Branched | 22:1(10)_Br |  |  |  | 11.67 | H | 2.1 | 0.0064 |
| C22:1n-14_Branched | 22:1(8)_Br |  |  |  | 11.7 | H | 0.67 | 0.0020 |
| C22:1n-7 | 22:1(15Z) | 15Z-docosenoic acid | LMFA01030402 | 15Z-docosenoic acid | 11.69 | H | 47 | 0.14 |
| C22:1n-9 | 22:1(13Z) | 13Z-docosenoic acid | LMFA01030089 | cis-erucic acid | 11.69 | H | 11.5 | 0.035 |
| C22:1n-10_Branched | 22:1(12)_Br |  |  |  | 11.59 | H | 6.2 | 0.019 |
| C22:1n-10 | 22:1(12Z) | 12Z-docosenoic acid | Not found in LIPID MAPS. |  | 11.69 | H | 8.8 | 0.027 |
| C22:1n-14 | 22:1(8Z) | 8Z-docosenoic acid | Not found in LIPID MAPS. |  | 11.82 | H | 0.67 | 0.002 |
| C22:1n-5 | 22:1(17Z) | 17Z-docosenoic acid | LMFA04000085 | 17Z-Docosenoic acid | 11.7 | H | 12.4 | 0.038 |
| C22:1n-3 | 22:1(19Z) | 19Z-docosenoic acid | LMFA01030403 | 19Z-docosenoic acid | 11.75 | H | 0.78 | 0.0024 |
| C22:1n-12 | 22:1(10Z) | 10Z-docosenoic acid | Not found in LIPID MAPS. |  | 11.74 | H | 0.66 | 0.002 |
| C22:1n-15_Branched | 22:1(7)_Br |  |  |  | 11.74 | H | 1.17 | 0.0036 |
| C22:1n-13 | 22:1(9Z) | 9Z-docosenoic acid | LMFA01030911 | 22:1(9Z) | 11.77 | H | 2.4 | 0.0073 |
| C22:1n-16_Branched | 22:1(6)_Br |  |  |  | 11.82 | H | 2.9 | 0.0088 |
| C22:1n-15 | 22:1(7Z) | 7Z-docosenoic acid | LMFA01030910 | 22:1(7Z) | 11.87 | H | 0.6 | 0.0018 |
| C22:1n-9 (E) | 22:1(13E) | 13E-docosenoic acid | LMFA01030090 | trans-brassidic acid | 11.83 | H | 0.7 | 0.0021 |
| C23:1n-11_Branched | 23:1(12)_Br |  |  |  | 12.08 | H | 14 | 0.0045 |
| C23:1n-8 | 23:1(15Z) | 15Z-tricosenoic acid | Not found in LIPID MAPS. |  | 12.08 | H | 7.1 | 0.0023 |
| C23:1n-6 | 23:1(17Z) | 17Z-tricosenoic acid | LMFA01030415 | 17Z-tricosenoic acid | 12.23 | H | 41 | 0.013 |
| C23:1n-9 | 23:1(14Z) | 14Z-tricosenoic acid | LMFA01030413 | 14Z-tricosenoic acid | 12.23 | H | 9.4 | 0.0030 |
| C23:1n-7 | 23:1(16Z) | 16Z-tricosenoic acid | LMFA01030414 | 16Z-tricosenoic acid | 12.23 | H | 12 | 0.0039 |
| C23:1n-14 | 23:1(9Z) | 9Z-tricosenoic acid | LMFA01031074 | 9Z-Tricosenoic acid | 12.3 | H | 16 | 0.0052 |
| C24:1n-10_Branched | 24:1(14)_Br |  |  |  | 12.62 | H | 17 | 0.049 |
| C24:1n-11_Branched | 24:1(13)_Br |  |  |  | 12.64 | H | 1.2 | 0.0035 |
| C24:1n-12_Branched | 24:1(12)_Br |  |  |  | 12.64 | H | 1.5 | 0.0043 |
| C24:1n-14_Branched | 24:1(10)_Br |  |  |  | 12.73 | H | 1.1 | 0.0032 |
| C24:1n-7 | 24:1(17Z) | 17Z-tetracosenoic acid | LMFA01031084 | 17Z-Tetracosenoic acid | 12.73 | H | 47 | 0.14 |
| C24:1n-9 | 24:1(15Z) | 15Z-tetracosenoic acid | LMFA01030092 | Nervonic acid | 12.73 | H | 7.4 | 0.021 |
| C24:1n-10 | 24:1(14Z) | 14Z-tetracosenoic acid | Not found in LIPID MAPS. |  | 12.73 | H | 17 | 0.049 |
| C24:1n-15_Branched | 24:1(9)_Br |  |  |  | 12.73 | H | 0.84 | 0.0024 |
| C24:1n-16_Branched | 24:1(8)_Br |  |  |  | 12.75 | H | 1.7 | 0.0049 |
| C24:1n-5 | 24:1(19Z) | 19Z-tetracosenoic acid | LMFA01031022 | 19Z-Tetracosenoic acid | 12.76 | H | 13 | 0.038 |
| C24:1n-15 | 24:1(9Z) | 9Z-tetracosenoic acid | Not found in LIPID MAPS. |  | 12.84 | H | 0.84 | 0.0024 |
| C25:1n-11_Branched | 25:1(14)_Br |  |  |  | 13.09 | H | 31 | 0.01 |
| C25:1n-6 | 25:1(19Z) | 19Z-pentacosenoic acid | LMFA01030421 | 19Z-pentacosenoic acid | 13.24 | H | 32 | 0.011 |
| C25:1n-8 | 25:1(17Z) | 17Z-pentacosenoic acid | LMFA01030419 | 17Z-pentacosenoic acid | 13.24 | H | 20 | 0.0066 |
| C25:1n-9 | 25:1(16Z) | 16Z-pentacosenoic acid | LMFA01030418 | 16Z-pentacosenoic acid | 13.24 | H | 6.4 | 0.0021 |
| C25:1n-16_Branched | 25:1(9)_Br |  |  |  | 13.24 | H | 10 | 0.0033 |
| C26:1n-10_Branched | 26:1(16)_Br |  |  |  | 13.59 | H | 20 | 0.033 |
| C26:1n-12_Branched | 26:1(14)_Br |  |  |  | 13.59 | H | 3.8 | 0.0063 |
| C26:1n-15_Branched | 26:1(11)_Br |  |  |  | 13.62 | H | 2.3 | 0.0038 |
| C26:1n-7 | 26:1(19Z) | 19Z-hexacosenoic acid | LMFA01030874 | 26:1(19Z)) | 13.71 | H | 40 | 0.066 |
| C26:1n-9 | 26:1(17Z) | 17Z-hexacosenoic acid | LMFA01030095 | Ximenic acid | 13.71 | H | 6.8 | 0.011 |
| C26:1n-16_Branched | 26:1(10)_Br |  | = | = | 13.71 | H | 1.2 | 0.002 |
| C26:1n-10 | 26:1(16Z) | 16Z-hexacosenoic acid | Not found in LIPID MAPS. | = | 13.71 | H | 20 | 0.033 |
| C26:1n-5 | 26:1(21Z) | 21Z-hexacosenoic acid | LMFA01031077 | 21Z-Hexacosenoic acid | 13.73 | H | 16 | 0.027 |
| C26:1n-3 | 26:1(23Z) | 23Z-hexacosenoic acid | Not found in LIPID MAPS. | = | 13.78 | H | 1.1 | 0.0018 |
| C27:1n-11_Branched | 27:1(16)_Br |  | = | = | 14.02 | H | 34 | <0.01 |
| C27:1n-6 | 27:1(21Z) | 21Z-heptacosenoic acid | Not found in LIPID MAPS. | = | 14.19 | H | 66 | <0.01 |
| C28:1n-10_Branched | 28:1(18)_Br |  | = | = | 14.48 | H | 12 | 0.016 |
| C28:1n-12_Branched | 28:1(16)_Br |  | = | = | 14.52 | H | 2.4 | 0.0031 |
| C28:1n-7 | 28:1(21Z) | 21Z-octacosenoic acid | LMFA01030431 | 21Z-octacosenoic acid | 14.61 | H | 57 | 0.074 |
| C28:1n-9 | 28:1(19Z) | 19Z-octacosenoic acid | LMFA01030096 | 19Z-octacosenoic acid | 14.61 | H | 2.2 | 0.0029 |
| C28:1n-10 | 28:1(18Z) | 18Z-octacosenoic acid | Not found in LIPID MAPS. | = | 14.61 | H | 12 | 0.016 |
| C28:1n-5 | 28:1(23Z) | 23Z-octacosenoic acid | LMFA01030432 | 23Z-octacosenoic acid | 14.62 | H | 18 | 0.023 |
| C29:1n-6 | 29:1(23Z) | 23Z-nonacosenoic acid | Not found in LIPID MAPS. | = | 15.02 | H | 66 | 0.0084 |
| C29:1n-8 | 29:1(21Z) | 21Z-nonacosenoic acid | Not found in LIPID MAPS. | = | 15.02 | H | 34 | 0.0043 |
| C30:1n-10_Branched | 30:1(20Z)_Br |  | = | = | 15.33 | H | 5 | 0.0093 |
| C30:1n-7 | 30:1(23Z) | 23Z-triacontenoic acid | LMFA01030875 | 30:1(23Z)) | 15.42 | H | 77 | 0.14 |
| C30:1n-9 | 30:1(21Z) | 21Z-triacontenoic acid | LMFA01030097 | Lumequeic acid | 15.42 | H | 2.6 | 0.0048 |
| C30:1n-10 | 30:1(20Z) | 20Z-triacontenoic acid | Not found in LIPID MAPS. | = | 15.42 | H | 5 | 0.0093 |
| C30:1n-5 | 30:1(25Z) | 25Z-triacontenoic acid | Not found in LIPID MAPS. | = | 15.44 | H | 13 | 0.024 |
| C16:2n-2,10 | 16:2(6,14) | 6,14-hexadecadienoic acid | Not found in LIPID MAPS. | = | 7.44 | H | 21 | 0.039 |
| C16:2n-3,10 | 16:2(6,13) | 6,13-hexadecadienoic acid | Not found in LIPID MAPS. | Menzeleic acid | 7.44 | H | 47 | 0.087 |
| C16:2n-4,8 | 16:2(8,12) | 8,12-hexadecadienoic acid | Not found in LIPID MAPS. | = | 7.43 | L |  |  |
| C16:2n-4,10 | 16:2(6,12) | 6,12-hexadecadienoic acid | Not found in LIPID MAPS. | = | 7.44 | L |  |  |
| C16:2n-5,8 | 16:2(8,11) | 8,11-hexadecadienoic acid | Not found in LIPID MAPS. | = | 7.42 | H | 2.8 | 0.0052 |
| C16:2n-6,10 | 16:2(6,10) | 6,10-hexadecadienoic acid | Not found in LIPID MAPS. | = | 7.44 | L |  |  |
| C17:2n-6,9 | 17:2(8,11) | 8Z,11Z-heptadecadienoic acid | LMFA01031037 | Norlinoleic acid | 8.05 | H |  | <0.05 |
| C17:2n-8,11 | 17:2(6,9) | 6Z,9Z-heptadecadienoic acid | Not found in LIPID MAPS. | = | 8.18 | L |  | <0.05 |
| C18:2n-3,9 | 18:2(9,15) | 9,15-octadecadienoic acid | Not found in LIPID MAPS. | = | 8.63 | L |  |  |
| C18:2n-3,10 | 18:2(8,15) | 8,15-octadecadienoic acid | Not found in LIPID MAPS. | Foigeic acid | 8.63 | H | 5.1 | 0.089 |
| C18:2n-4,9 | 18:2(9,14) | 9,14-octadecadienoic acid | Not found in LIPID MAPS. | = | 8.62 | L |  |  |
| C18:2n-4,10 | 18:2(8,14) | 8,14-octadecadienoic acid | Not found in LIPID MAPS. | = | 8.62 | L |  |  |
| C18:2n-6,10 | 18:2(8,12) | 8,12-octadecadienoic acid | Not found in LIPID MAPS. | = | 8.62 | L |  |  |
| C18:2n-3,6 | 18:2(12,15) | 12,15-octadecadienoic acid | LMFA01030302 | 12Z,15Z-octadecadienoic acid | 8.65 | L |  |  |
| C18:2n-6,9 | 18:2(9,12) | 9Z,12Z-octadecadienoic acid | LMFA01030120 | Linoleic acid | 8.63 | H | 37 | 0.64 |
| C18:2n-7,10 | 18:2(8,11) | 8Z,11Z-octadecadienoic acid | LMFA01030116 | 8Z,11Z-octadecadienoic acid | 8.67 | H | 15 | 0.26 |
| C18:2n-2,10 | 18:2(8,16) | 8,16-octadecadienoic acid | Not found in LIPID MAPS. | = | 8.74 | L |  |  |
| C18:2n-2,12 | 18:2(6,16) | 6,16-octadecadienoic acid | Not found in LIPID MAPS. | = | 8.73 | L |  |  |
| C18:2n-3,10_(E) | 18:2(8,15) | 8,15-octadecadienoic acid | Not found in LIPID MAPS. | = | 8.74 | L |  |  |
| C18:2n-3,12 | 18:2(6,15) | 6,15-octadecadienoic acid | Not found in LIPID MAPS. | = | 8.81 | L |  |  |
| C18:2n-7,9 | 18:2(9,11) | 9,11-octadecadienoic acid | LMFA01030117 | Ricinenic acid | 8.85 | H |  | <0.01 |
| C18:2n-6,8 | 18:2(10,12) | 10,12-octadecadienoic acid | LMFA01030124 | 10Z,12Z-octadecadienoic acid | 8.83 | H |  | <0.01 |
| C18:2n-9,12 | 18:2(6,9) | 6Z,9Z-octadecadienoic acid | LMFA01030332 | 6Z,9Z-octadecadienoic acid | 8.83 | H | 1.2 | 0.021 |
| C18:2n-10,13 | 18:2(5,8) | 5Z,8Z-octadecadienoic acid | LMFA01030323 | Sebaleic acid | 8.93 | H | 17 | 0.30 |
| C19:2n-7,10 | 19:2(9,12) | 9Z,12Z-nonadecadienoic acid | Not found in LIPID MAPS. | = | 9.3 | H | 28 | 0.014 |
| C19:2n-8,11 | 19:2(8,11) | 8Z,11Z-nonadecadienoic acid | Not found in LIPID MAPS. | = | 9.3 | H | 45 | 0.023 |
| C19:2n-11,14 | 19:2(5,8) | 5Z,8Z-nonadecadienoic acid | Not found in LIPID MAPS. | = | 9.57 | H | 27 | 0.014 |
| C20:2n-6,9 | 20:2(11,14) | 11Z,14Z-eicosadienoic acid | LMFA01031043 | Dihomolinsoleic acid | 9.81 | H | 7.8 | 0.014 |
| C20:2n-7,10 | 20:2(10,13) | 10Z,13Z-eicosadienoic acid | Not found in LIPID MAPS. | = | 9.81 | H | 18 | 0.033 |
| C20:2n-9,12 | 20:2(8,11) | 8Z,11Z-eicosadienoic acid | LMFA01030377 | 8Z,11Z-eicosadienoic acid | 9.9 | H | 18 | 0.033 |
| C20:2n-10,13 | 20:2(7,10) | 7Z,10Z-eicosadienoic acid | Not found in LIPID MAPS. | = | 9.96 | H | 53 | 0.096 |
| C20:2n-12,15 | 20:2(5,8) | 5Z,8Z-eicosadienoic acid | Not found in LIPID MAPS. | = | 10.05 | H | 3.2 | 0.0058 |
| C22:2n-7,10 | 22:2(12,15) | 12Z,15Z-docosadienoic acid | Not found in LIPID MAPS. | — | 10.91 | H | 100 | 0.048 |
| C24:2n-7,10 | 24:2(14,17) | 14Z,17Z-tetracosadienoic acid | Not found in LIPID MAPS. | — | 12 | H | 100 | 0.018 |
| C26:2n-7,10 | 26:2(16,19) | 16Z,19Z-hexacosadienoic acid | Not found in LIPID MAPS. | — | 12.98 | H | 100 | <0.01 |
| C30:2n-7,10 | 30:2(20,23) | 20Z,23Z-triacontadienoic acid | Not found in LIPID MAPS. | — | 14.76 | H | 100 | 0.0088 |
| C18:3n-3,6,9 | 18:3(9,12,15) | 9Z,12Z,15Z-octadecatrienoic acid | LMFA01030152 | alpha-Linolenic acid | 7.95 | H | 15 | 0.0097 |
| C18:3n-6,9,12 | 18:3(6,9,12) | 6Z,9Z,12Z-octadecatrienoic acid | LMFA01030141 | gamma-Linolenic acid | 8.11 | H | 11 | 0.0071 |
| C18:3n-3,10,13 | 18:3(5,8,15) | 5,8,15-octadecatrienoic acid | Not found in LIPID MAPS. | *Dinheic acid* | 8.11 | H | 24 | 0.016 |
| C18:3n-7,10,13 | 18:3(5,8,11) | 5Z,8Z,11Z-octadecatrienoic acid | Not found in LIPID MAPS. | — | 8.19 | H | 50 | 0.033 |
| C20:3n-6,9,12 | 20:3(8,11,14) | 8Z,11Z,14Z-eicosatrienoic acid | LMFA01030387 | — | 9.15 | H | 25.5 | 0.037 |
| C20:3n-7,10,13 | 20:3(7,10,13) | 7Z,10Z,13Z-eicosatrienoic acid | LMFA01030384 | 7Z,10Z,13Z-eicosatrienoic acid | 9.19 | H | 18 | 0.026 |
| C20:3n-9,12,15 | 20:3(5,8,11) | 5Z,8Z,11Z-eicosatrienoic acid | LMFA01030381 | Mead acid | 9.42 | H | 56.4 | 0.081 |
| C22:3n-10,13,16 | 22:3(6,9,12) | 6Z,9Z,12Z-docosatrienoic acid | Not found in LIPID MAPS. | — | 10.49 | H | 100 | 0.030 |
| C20:4n-6,9,12,15 | 20:4(5,8,11,14) | 5Z,8Z,11Z,14Z-eicosatetraenoic acid | LMFA01030001 | Arachidonic acid | 8.68 | H | 100 | 0.039 |

### 3.6 Fatty acids in Fetal Bovine Serum (FBS)

Fetal bovine serum analyzed here was heat inactivated prior to lipid extraction. Lipids from 0.1 mL fetal bovine serum were extracted and hydrolyzed as per the procedures described in the methods section of this publication. Fatty acids were derivatized with AMPP and analyzed according to the methods described within this work. Twelve fatty acid species were discovered in fetal bovine serum that have to the best of our knowledge not been reported previously (including discoveries reported in human plasma or vernix caseosa herein). Additionally, 42 fatty acids that were discovered for the first time in either human plasma or vernix caseosa within this work, were also identified in fetal bovine serum.

**Figure S29:**
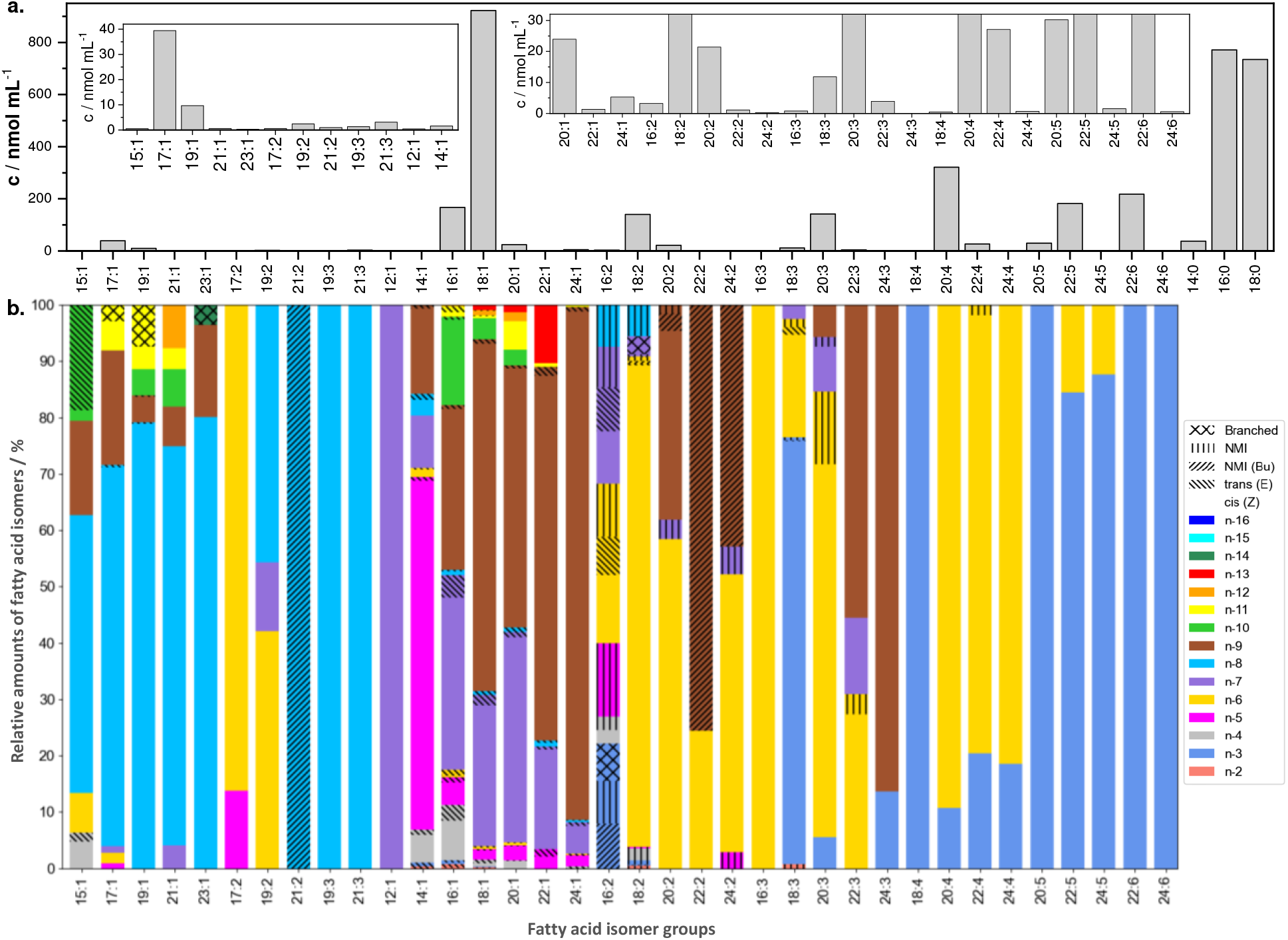
Fatty acids in fetal bovine serum. a. Estimation of absolute quantities of non-isomeric fatty acids by direct infusion mass spectrometry. b. Relative amounts of fatty acid isomers by LC-OzID-MS and LC-OzID-MS/MS.

**Table S7:**
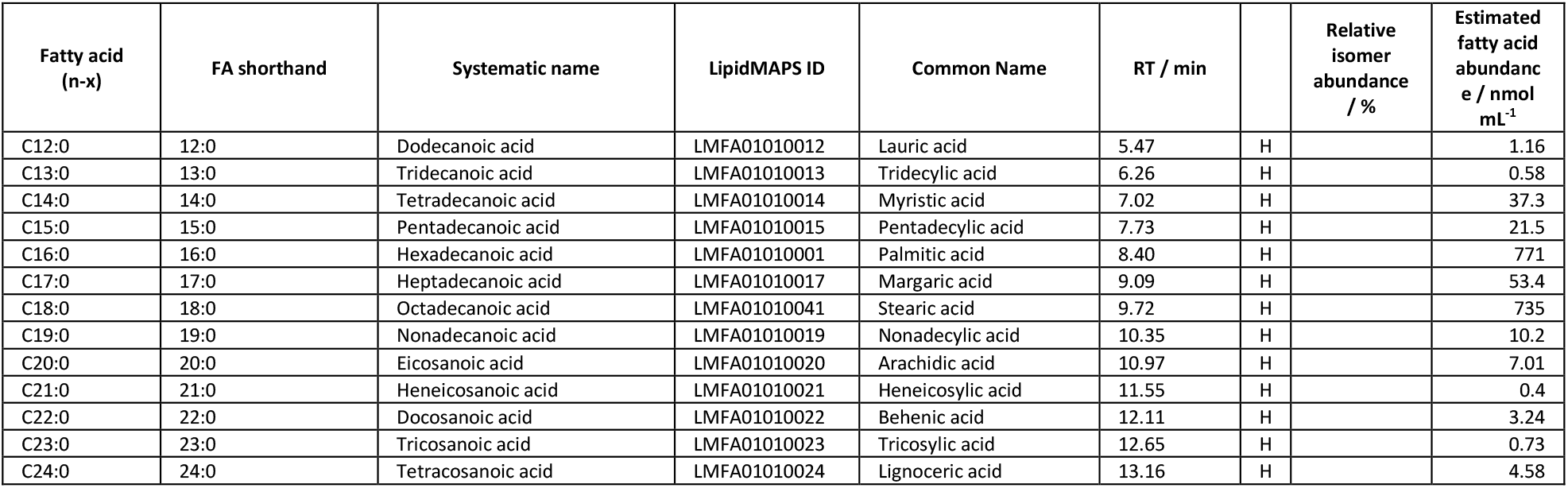
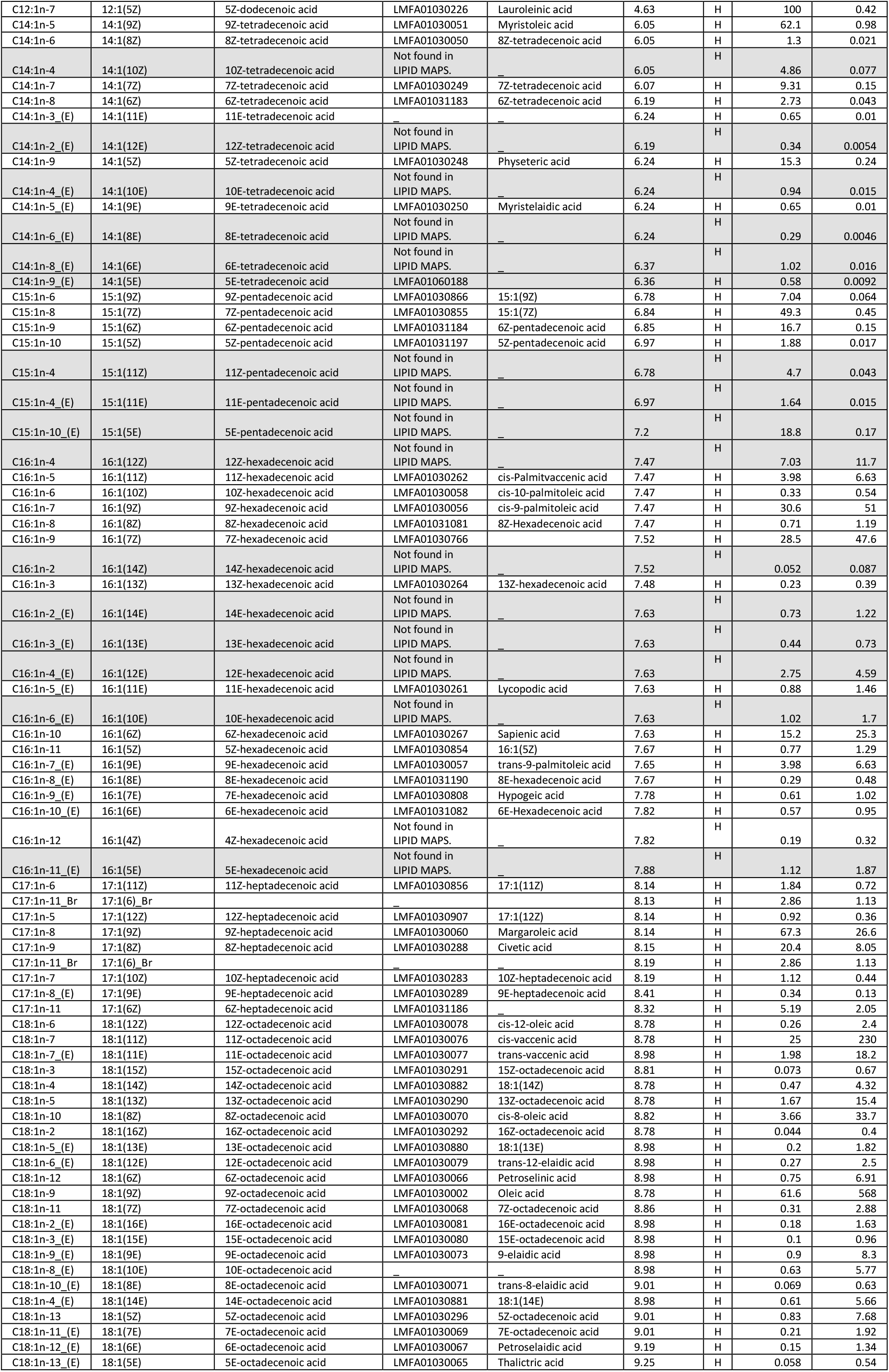
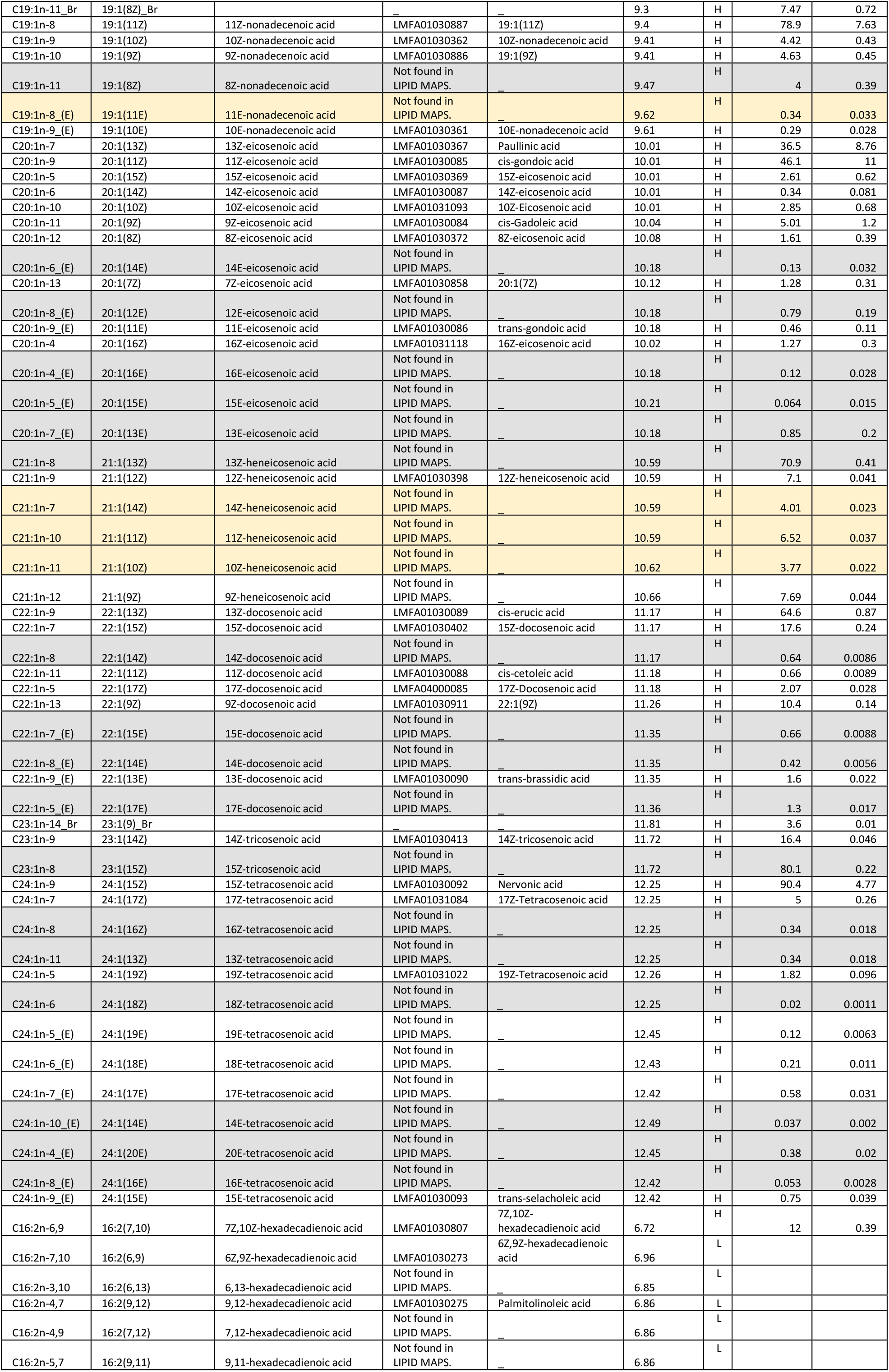
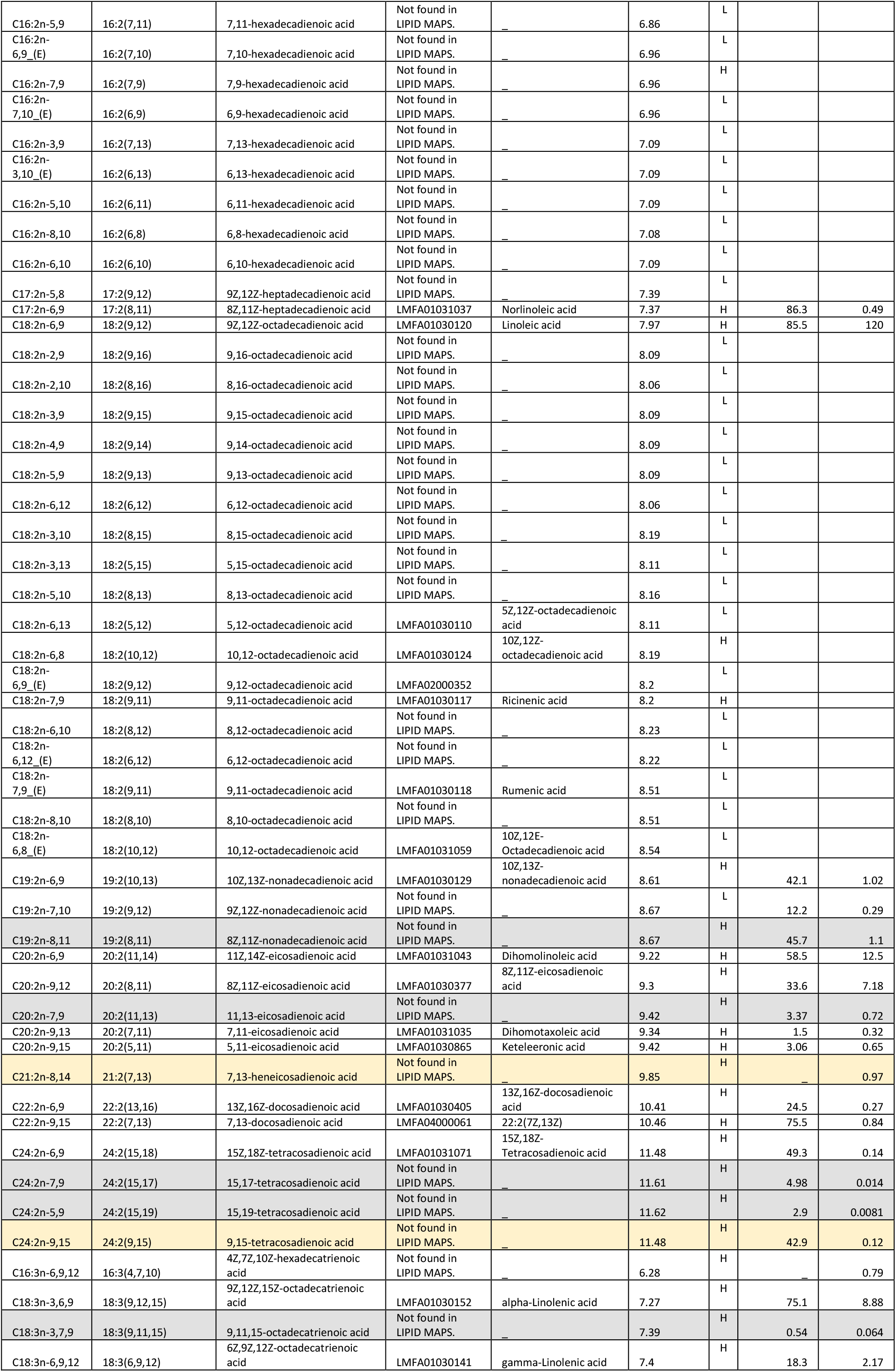
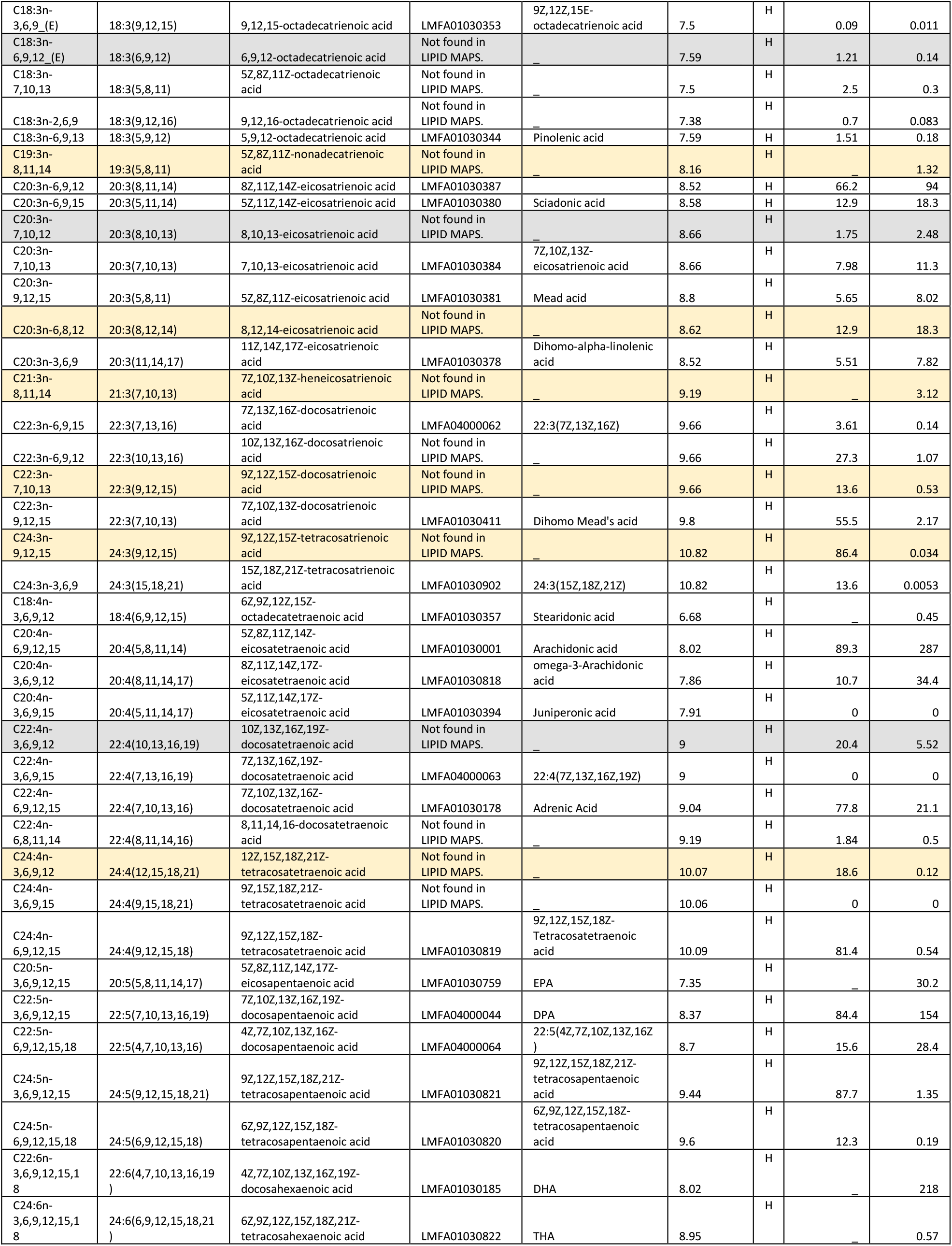
Overview of fatty acids that are identified here in fetal bovine serum. Fatty acid isomers that, to the best of our knowledge, have not been reported before, are highlighted with a yellow background, while those that we also discovered in pooled human plasma or vernix caseosa are highlighted with a grey background.

### 3.7 Fatty acids in cancer cell extracts

**Figure S30:**
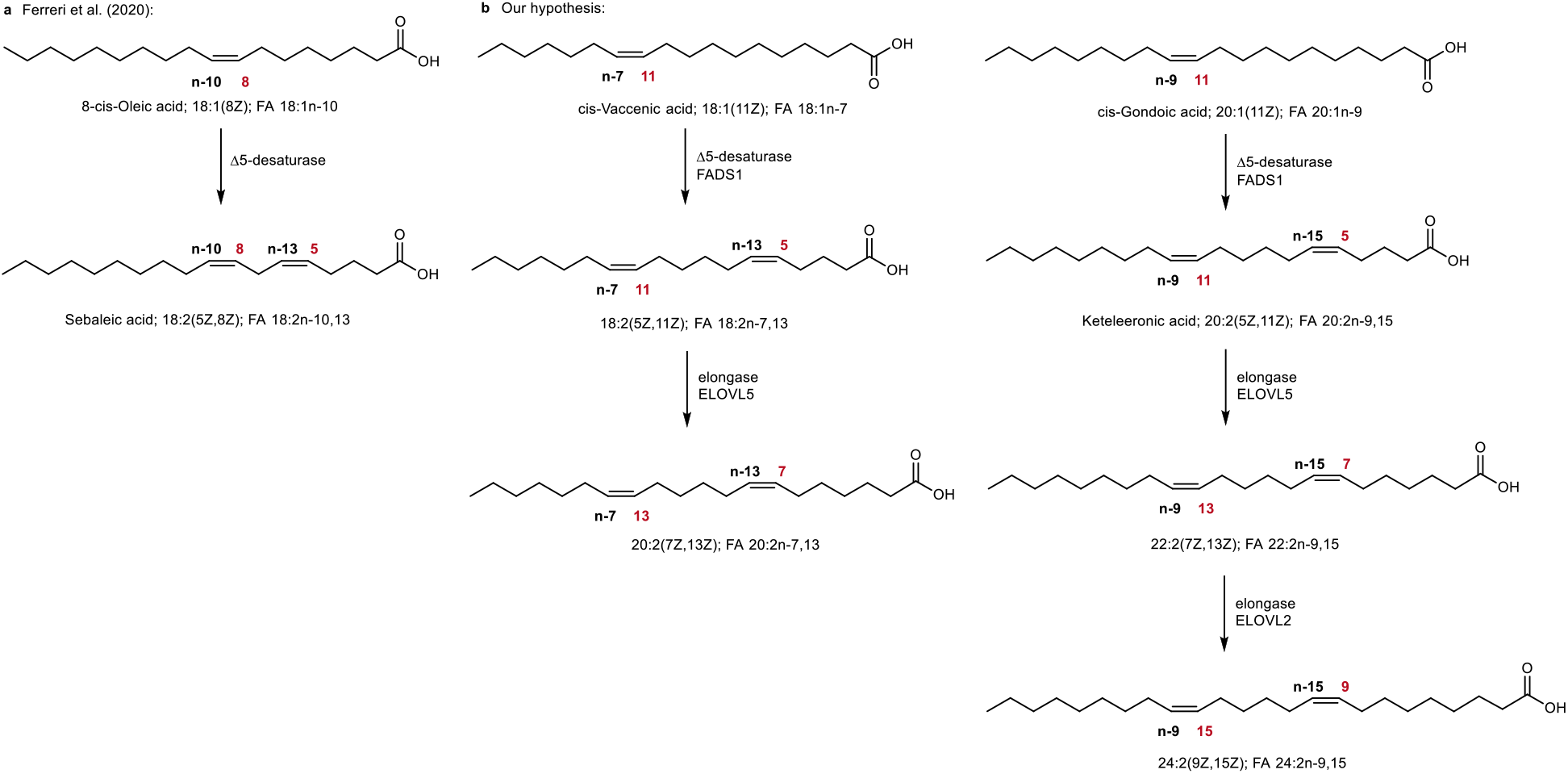
**a**. Proposed biosynthetic pathway to sebaleic acid by Ferreri et al.^40^ **b**. Our proposed pathways to butylene interrupted species.

#### 3.7.1 MCF7 Breast Cancer - Human breast adenocarcinoma, metastatic

The MCF7 cell line, named after the Michigan Cancer Foundation, consists of epithelial cells from a metastatic adenocarcinoma (breast tissue) of a 69-year-old female patient of Caucasian ethnic background.

**Figure S31:**
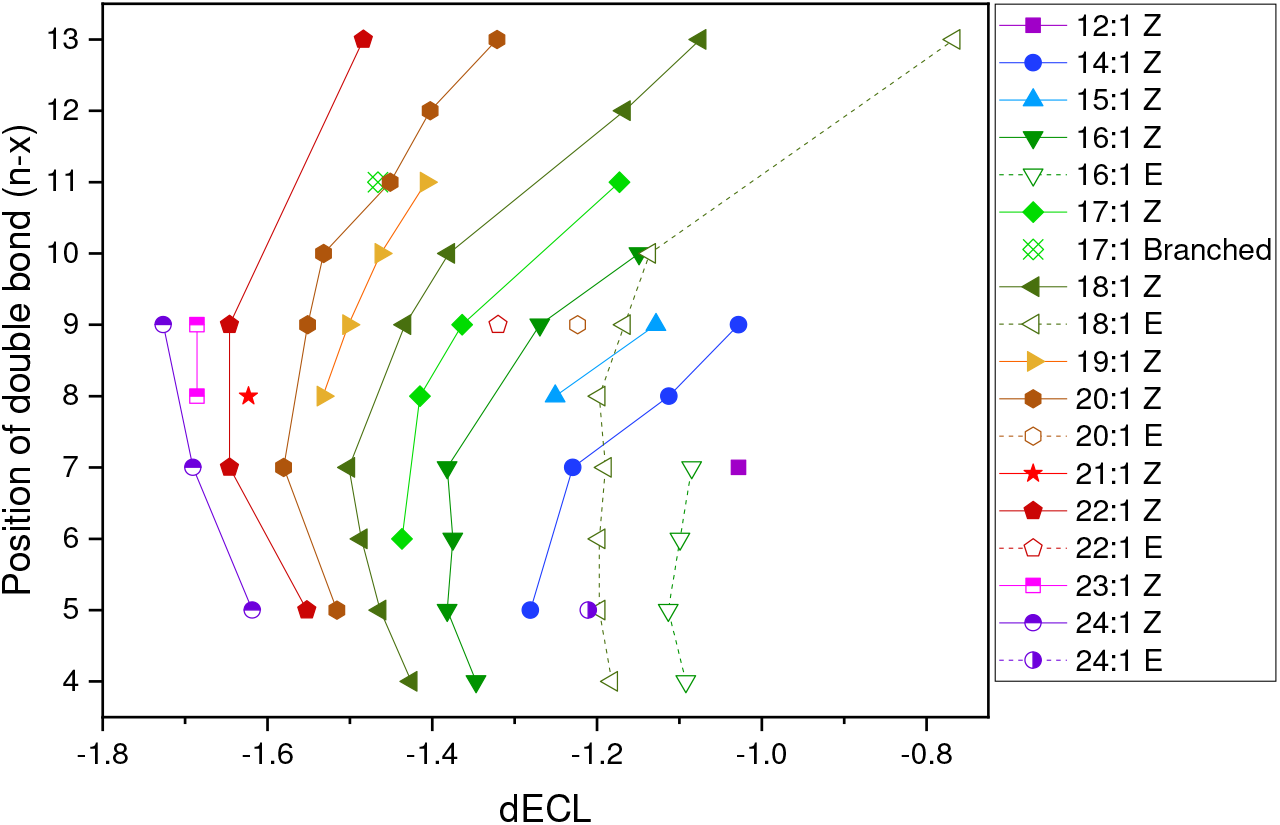
Analysis of differential equivalent chain lengths (dECL) of monounsaturated fatty acids in MCF7 cells. ECL and dECL values were determined with the equation ECL= 0.04115 * (RT)^2^ + 0.79914 * RT + 6.50734, based on the retention times of saturated fatty acids in the same sample.

**Figure S32:**
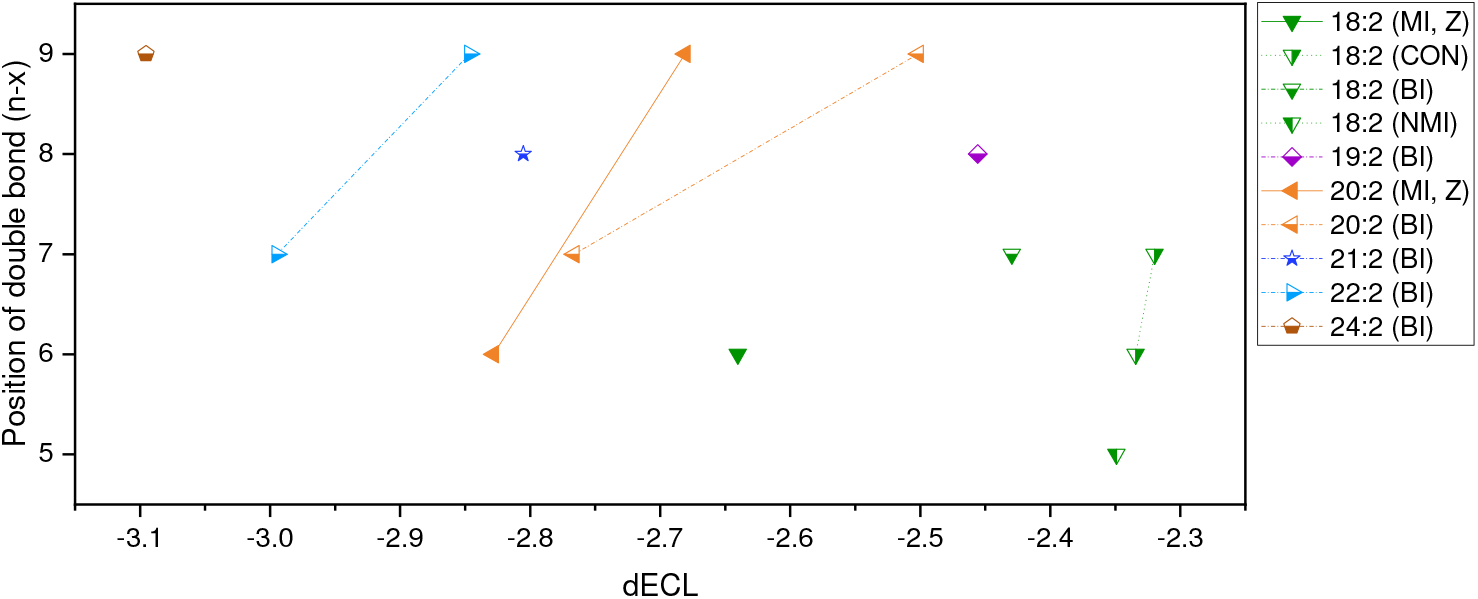
Analysis of differential equivalent chain lengths (dECL) of bisunsaturated fatty acids in MCF7 cells. Non-methylene interrupted fatty acids are assigned as conjugated (CON), butylene-interrupted (BI) or other non-methylene interrupted fatty acids (NMI). Double bond configurations cannot be established unequivocally for all fatty acid species. ECL and dECL values were determined with the equation ECL= 0.04115 * (RT)^2^ + 0.79914 * RT + 6.50734, based on the retention times of saturated fatty acids in the same sample.

**Figure S33:**
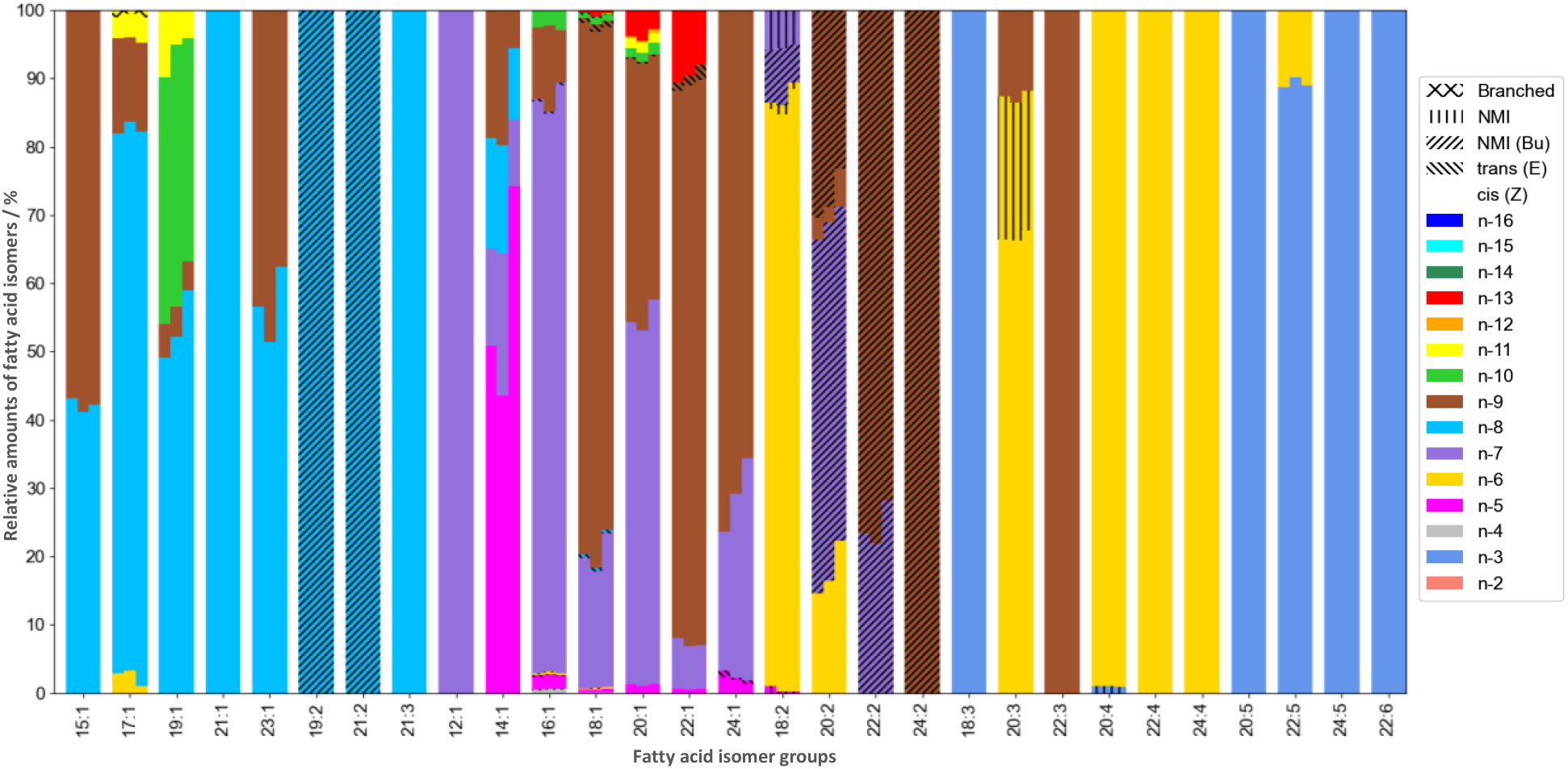
Relative amounts of fatty acid isomers identified in lipid extracts of MCF7 cancer cells (total fatty acid content of hydrolyzed lipids). Shown are the relative amounts of three replicates (three instances of MCF7 cells undergoing cell culture). Note that the deviation between the replicates is the largest for tetradecenoic acids. For example, the third replicate shows a significantly larger relative amount of 9Z-tetradecenoic acid (n-5) compared to the other replicates. Yet, relative amounts of other fatty acid isomers are not deviating substantially between the replicates.

**Table S8:** Overview of fatty acids that are identified here in lipid extracts of MCF7 cells. Column seven shows mean values of relative isomer abundances and their associated standard deviations of three replicates of cell cultures of MCF7 cells. Fatty isomers that, to the best of our knowledge, have not been reported before, are highlighted with a yellow background, while those that we also discovered in pooled human plasma or vernix caseosa are highlighted with a grey background. Fatty acid abundance is reported here in nmol per million cells (nmol M-Cells^-1^) as an estimate of the fatty acid content of each isomer in the cell culture.

| Fatty acid (n-x) | FA shorthand | Systematic name | LipidMAPS ID | Common Name | RT / min | Relative isomer abundance / % | COV (rel. isomer abundance) | Estimated fatty acid abundance / nmol M-Cells<sup>-1</sup> |
| --- | --- | --- | --- | --- | --- | --- | --- | --- |
| C12:0 | 12:0 | Dodecanoic acid | LMFA01010012 | Lauric acid | 5.38 | n.a. | n.a. | 1.0 ± 0.7 |
| C13:0 | 13:0 | Tridecanoic acid | LMFA01010013 | Tridecylic acid | 6.17 | n.a. | n.a. | 0.3 ± 0.3 |
| C14:0 | 14:0 | Tetradecanoic acid | LMFA01010014 | Myristic acid | 6.91 | n.a. | n.a. | 5 ± 1 |
| C15:0 | 15:0 | Pentadecanoic acid | LMFA01010015 | Pentadecylic acid | 7.63 | n.a. | n.a. | 14 ± 5 |
| C16:0 | 16:0 | Hexadecanoic acid | LMFA01010001 | Palmitic acid | 8.31 | n.a. | n.a. | 32 ± 7 |
| C17:0 | 17:0 | Heptadecanoic acid | LMFA01010017 | Margaric acid | 8.97 | n.a. | n.a. | 1.7 ± 0.3 |
| C18:0 | 18:0 | Octadecanoic acid | LMFA01010041 | Stearic acid | 9.61 | n.a. | n.a. | 27 ± 6 |
| C19:0 | 19:0 | Nonadecanoic acid | LMFA01010019 | Nonadecylic acid | 10.26 | n.a. | n.a. | 0.4 ± 0.3 |
| C20:0 | 20:0 | Eicosanoic acid | LMFA01010020 | Arachidic acid | 10.84 | n.a. | n.a. | 2.2 ± 0.7 |
| C21:0 | 21:0 | Heneicosanoic acid | LMFA01010021 | Heneicosylic acid | 11.42 | n.a. | n.a. | 0.02 ± 0.02 |
| C22:0 | 22:0 | Docosanoic acid | LMFA01010022 | Behenic acid | 12.00 | n.a. | n.a. | 3 ± 1 |
| C23:0 | 23:0 | Tricosanoic acid | LMFA01010023 | Tricosylic acid | 12.55 | n.a. | n.a. | 0.001 ± 0.002 |
| C24:0 | 24:0 | Tetracosanoic acid | LMFA01010024 | Lignoceric acid | 13.07 | n.a. | n.a. | 3 ± 1 |
| C12:1n-7 | 12:1(5Z) | 5Z-dodecenoic acid | LMFA01030226 | Lauroleinic acid | 4.56 |  |  | 0.045 ± 0.007 |
| C14:1n-5 | 14:1(9Z) | 9Z-tetradecenoic acid | LMFA01030051 | Myristoleic acid | 5.96 | 60 ± 20 | 0.4 | 0.13 ± 0.09 |
| C14:1n-7 | 14:1(7Z) | 7Z-tetradecenoic acid | LMFA01030249 | 7Z-tetradecenoic acid | 5.99 | 15 ± 6 | 0.4 | 0.03 ± 0.006 |
| C14:1n-8 | 14:1(6Z) | 6Z-tetradecenoic acid | LMFA01031183 | 6Z-tetradecenoic acid | 6.07 | 14 ± 3 | 0.2 | 0.03 ± 0.004 |
| C14:1n-9 | 14:1(5Z) | 5Z-tetradecenoic acid | LMFA01030248 | Physeteric acid | 6.13 | 15 ± 8 | 0.5 | 0.028 ± 0.009 |
| C15:1n-8 | 15:1(7Z) | 7Z-pentadecenoic acid | LMFA01030855 | 15:1(7Z) | 6.73 | 42 ± 1 | 0.02 | 0.012 ± 0.003 |
| C15:1n-9 | 15:1(6Z) | 6Z-pentadecenoic acid | LMFA01031184 | 6Z-pentadecenoic acid | 6.82 | 58 ± 1 | 0.02 | 0.016 ± 0.003 |
| C16:1n-4 | 16:1(12Z) | 12Z-hexadecenoic acid | Not found in LIPID MAPS. |  | 7.37 | 0.41 ± 0.03 | 0.07 | 0.07 ± 0.03 |
| C16:1n-5 | 16:1(11Z) | 11Z-hexadecenoic acid | LMFA01030262 | cis-Palmitvaccenic acid | 7.37 | 1.9 ± 0.1 | 0.05 | 0.3 ± 0.1 |
| C16:1n-6 | 16:1(10Z) | 10Z-hexadecenoic acid | LMFA01030058 | cis-10-palmitoleic acid | 7.37 | 0.26 ± 0.04 | 0.2 | 0.05 ± 0.02 |
| C16:1n-7 | 16:1(9Z) | 9Z-hexadecenoic acid | LMFA01030056 | cis-9-palmitoleic acid | 7.37 | 84 ± 2 | 0.02 | 14 ± 5 |
| C16:1n-9 | 16:1(7Z) | 7Z-hexadecenoic acid | LMFA01030766 |  | 7.4 | 10 ± 2 | 0.2 | 1.7 ± 0.5 |
| C16:1n-4_(E) | 16:1(12E) | 12E-hexadecenoic acid | Not found in LIPID MAPS. |  | 7.52 | 0.14 ± 0.01 | 0.07 | 0.024 ± 0.006 |
| C16:1n-5_(E) | 16:1(11E) | 11E-hexadecenoic acid | LMFA01030261 | Lycopodic acid | 7.52 | 0.12 ± 0.02 | 0.2 | 0.02 ± 0.004 |
| C16:1n-6_(E) | 16:1(10E) | 10E-hexadecenoic acid | Not found in LIPID MAPS. |  | 7.53 | 0.08 ± 0.01 | 0.1 | 0.013 ± 0.003 |
| C16:1n-10 | 16:1(6Z) | 6Z-hexadecenoic acid | LMFA01030267 | Sapienic acid | 7.52 | 2.5 ± 0.3 | 0.1 | 0.4 ± 0.2 |
| C16:1n-7_(E) | 16:1(9E) | 9E-hexadecenoic acid | LMFA01030057 | trans-9-palmitoleic acid | 7.54 | 0.32 ± 0.07 | 0.2 | 0.06 ± 0.02 |
| C17:1n-6 | 17:1(11Z) | 11Z-heptadecenoic acid | LMFA01030856 | 17:1(11Z) | 8.03 | 2 ± 1 | 0.4 | 0.02 ± 0.01 |
| C17:1n-8 | 17:1(9Z) | 9Z-heptadecenoic acid | LMFA01030060 | Margaroic acid | 8.03 | 80 ± 1 | 0.01 | 0.8 ± 0.2 |
| C17:1n-9 | 17:1(8Z) | 8Z-heptadecenoic acid | LMFA01030288 | Civetic acid | 8.04 | 13.1 ± 0.8 | 0.06 | 0.13 ± 0.02 |
| C17:1n-11_Br | 17:1(6Z)_Br |  |  |  | 8.02 | 0.9 ± 0.3 | 0.3 | 0.009 ± 0.003 |
| C17:1n-11 | 17:1(6Z) | 6Z-heptadecenoic acid | LMFA01031186 | 6Z-heptadecenoic acid | 8.19 | 3.4 ± 0.4 | 0.1 | 0.04 ± 0.01 |
| C18:1n-6 | 18:1(12Z) | 12Z-octadecenoic acid | LMFA01030078 | cis-12-oleic acid | 8.69 | 0.12 ± 0.04 | 0.3 | 0.06 ± 0.03 |
| C18:1n-7 | 18:1(11Z) | 11Z-octadecenoic acid | LMFA01030076 | cis-vaccenic acid | 8.69 | 20 ± 3 | 0.2 | 9 ± 2 |
| C18:1n-7_(E) | 18:1(11E) | 11E-octadecenoic acid | LMFA01030077 | trans-vaccenic acid | 8.86 | 0.24 ± 0.02 | 0.08 | 0.12 ± 0.02 |
| C18:1n-4 | 18:1(14Z) | 14Z-octadecenoic acid | LMFA01030882 | 18:1(14Z) | 8.69 | 0.06 ± 0.008 | 0.1 | 0.029 ± 0.008 |
| C18:1n-5 | 18:1(13Z) | 13Z-octadecenoic acid | LMFA01030290 | 13Z-octadecenoic acid | 8.69 | 0.44 ± 0.07 | 0.2 | 0.21 ± 0.06 |
| C18:1n-10 | 18:1(8Z) | 8Z-octadecenoic acid | LMFA01030070 | cis-8-oleic acid | 8.7 | 0.8 ± 0.2 | 0.3 | 0.4 ± 0.2 |
| C18:1n-5_(E) | 18:1(13E) | 13E-octadecenoic acid | LMFA01030880 | 18:1(13E) | 8.86 | 0.026 ± 0.0006 | 0.02 | 0.013 ± 0.002 |
| C18:1n-6_(E) | 18:1(12E) | 12E-octadecenoic acid | LMFA01030079 | trans-12-elaidic acid | 8.86 | 0.039 ± 0.0008 | 0.02 | 0.019 ± 0.004 |
| C18:1n-12 | 18:1(6Z) | 6Z-octadecenoic acid | LMFA01030066 | Petroselinic acid | 8.86 | 0.039 ± 0.008 | 0.2 | 0.019 ± 0.006 |
| C18:1n-9 | 18:1(9Z) | 9Z-octadecenoic acid | LMFA01030002 | Oleic acid | 8.69 | 77 ± 3 | 0.04 | 37 ± 7 |
| C18:1n-9_(E) | 18:1(9E) | 9E-octadecenoic acid | LMFA01030073 | 9-elaidic acid | 8.86 | 0.9 ± 0.2 | 0.2 | 0.5 ± 0.2 |
| C18:1n-8_(E) | 18:1(10E) | 10E-octadecenoic acid | LMFA01030075 | 10E-octadecenoic acid | 8.86 | 0.29 ± 0.03 | 0.1 | 0.14 ± 0.04 |
| C18:1n-10_(E) | 18:1(8E) | 8E-octadecenoic acid | LMFA01030071 | trans-8-elaidic acid | 8.86 | 0.23 ± 0.02 | 0.09 | 0.11 ± 0.03 |
| C18:1n-4_(E) | 18:1(14E) | 14E-octadecenoic acid | LMFA01030881 | 18:1(14E) | 8.86 | 0.044 ± 0.002 | 0.05 | 0.021 ± 0.005 |
| C18:1n-13 | 18:1(5Z) | 5Z-octadecenoic acid | LMFA01030296 | 5Z-octadecenoic acid | 8.92 | 0.5 ± 0.3 | 0.6 | 0.2 ± 0.2 |
| C18:1n-13_(E) | 18:1(5E) | 5E-octadecenoic acid | LMFA01030065 | Thalicttric acid | 9.12 | 0.07 ± 0.02 | 0.3 | 0.03 ± 0.02 |
| C19:1n-9 | 19:1(10Z) | 10Z-nonadecenoic acid | LMFA01030362 | 10Z-nonadecenoic acid | 9.28 | 4.5 ± 0.3 | 0.07 | 0.003 ± 0.004 |
| C19:1n-8 | 19:1(11Z) | 11Z-nonadecenoic acid | LMFA01030887 | 19:1(11Z) | 9.28 | 53 ± 5 | 0.09 | 0.04 ± 0.06 |
| C19:1n-10 | 19:1(9Z) | 9Z-nonadecenoic acid | LMFA01030886 | 19:1(9Z) | 9.28 | 36 ± 3 | 0.08 | 0.02 ± 0.03 |
| C19:1n-11 | 19:1(8Z) | 8Z-nonadecenoic acid | Not found in LIPID MAPS. |  | 9.29 | 6 ± 3 | 0.5 | 0.003 ± 0.004 |
| C20:1n-7 | 20:1(13Z) | 13Z-eicosenoic acid | LMFA01030367 | Paullinic acid | 9.88 | 54 ± 2 | 0.04 | 0.9 ± 0.2 |
| C20:1n-9 | 20:1(11Z) | 11Z-eicosenoic acid | LMFA01030085 | cis-gondoic acid | 9.88 | 38 ± 2 | 0.05 | 0.6 ± 0.1 |
| C20:1n-5 | 20:1(15Z) | 15Z-eicosenoic acid | LMFA01030369 | 15Z-eicosenoic acid | 9.88 | 1.2 ± 0.2 | 0.2 | 0.018 ± 0.001 |
| C20:1n-10 | 20:1(10Z) | 10Z-eicosenoic acid | LMFA01031093 | 10Z-Eicosenoic acid | 9.88 | 1.5 ± 0.3 | 0.2 | 0.024 ± 0.006 |
| C20:1n-11 | 20:1(9Z) | 9Z-eicosenoic acid | LMFA01030084 | cis-Gadoleic acid | 9.91 | 1.46 ± 0.04 | 0.03 | 0.023 ± 0.004 |
| C20:1n-12 | 20:1(8Z) | 8Z-eicosenoic acid | LMFA01030372 | 8Z-eicosenoic acid | 10.03 | 0.4 ± 0.1 | 0.3 | 0.006 ± 0.002 |
| C20:1n-13 | 20:1(7Z) | 7Z-eicosenoic acid | LMFA01030858 | 20:1(7Z) | 10.04 | 3.7 ± 0.9 | 0.3 | 0.06 ± 0.02 |
| C20:1n-9_(E) | 20:1(11E) | 11E-eicosenoic acid | LMFA01030086 | trans-gondoic acid | 10.09 | 0.27 ± 0.01 | 0.04 | 0.0044 ± 0.0009 |
| C21:1n-8 | 21:1(13Z) | 13Z-heneicosenoic acid | Not found in LIPID MAPS. |  | 10.46 | 100 ± 0 | 0.0 | 0.00066 ± 0.001 |
| C22:1n-9 | 22:1(13Z) | 13Z-docosenoic acid | LMFA01030089 | cis-erucic acid | 11.04 | 82 ± 1 | 0.01 | 1.0 ± 0.3 |
| C22:1n-7 | 22:1(15Z) | 15Z-docosenoic acid | LMFA01030402 | 15Z-docosenoic acid | 11.04 | 6.8 ± 0.6 | 0.09 | 0.08 ± 0.02 |
| C22:1n-5 | 22:1(17Z) | 17Z-docosenoic acid | LMFA04000085 | 17Z-Docosenoic acid | 11.05 | 0.57 ± 0.09 | 0.2 | 0.006 ± 0.001 |
| C22:1n-13 | 22:1(9Z) | 9Z-docosenoic acid | LMFA01030911 | 22:1(9Z) | 11.13 | 9 ± 1 | 0.1 | 0.11 ± 0.03 |
| C22:1n-9_(E) | 22:1(13E) | 13E-docosenoic acid | LMFA01030090 | trans-brassicidic acid | 11.23 | 1.6 ± 0.5 | 0.3 | 0.019 ± 0.009 |
| C23:1n-9 | 23:1(14Z) | 14Z-tricosenoic acid | LMFA01030413 | 14Z-tricosenoic acid | 11.6 | 43 ± 6 | 0.1 | 0.005 ± 0.009 |
| C23:1n-8 | 23:1(15Z) | 15Z-tricosenoic acid | Not found in LIPID MAPS. | — | 11.6 | 57 ± 6 | 0.1 | 0.005 ± 0.009 |
| C24:1n-9 | 24:1(15Z) | 15Z-tetracosenoic acid | LMFA01030092 | Nervonic acid | 12.15 | 71 ± 5 | 0.07 | 0.31 ± 0.05 |
| C24:1n-7 | 24:1(17Z) | 17Z-tetracosenoic acid | LMFA01031084 | 17Z-Tetracosenoic acid | 12.15 | 27 ± 6 | 0.2 | 0.12 ± 0.06 |
| C24:1n-5 | 24:1(19Z) | 19Z-tetracosenoic acid | LMFA01031022 | 19Z-Tetracosenoic acid | 12.15 | 1.8 ± 0.4 | 0.2 | 0.0076 ± 0.0007 |
| C24:1n-5_(E) | 24:1(19E) | 19E-tetracosenoic acid | Not found in LIPID MAPS. | — | 12.29 | 0.7 ± 0.5 | 0.8 | 0.003 ± 0.001 |
| C18:2n-6,9 | 18:2(9,12) | 9Z,12Z-octadecadienoic acid | LMFA01030120 | Linoleic acid | 7.86 | 86 ± 2 | 0.02 | 2.1 ± 0.6 |
| C18:2n-7,13 | 18:2(5,11) | 5,11-octadecadienoic acid | LMFA01030322 | Ephedrenic acid | 8.02 | 7 ± 1 | 0.1 | 0.17 ± 0.06 |
| C18:2n-5,9 | 18:2(9,13) | 9,13-octadecadienoic acid | Not found in LIPID MAPS. | — | 8 | 0.4 ± 0.4 | 0.9 | 0.008 ± 0.006 |
| C18:2n-6,8 | 18:2(10,12) | 10,12-octadecadienoic acid | LMFA01030124 | 10Z,12Z-octadecadienoic acid | 8.08 | 1.0 ± 0.2 | 0.2 | 0.03 ± 0.01 |
| C18:2n-7,9 | 18:2(9,11) | 9,11-octadecadienoic acid | LMFA01030117 | Ricinenic acid | 8.08 | 5.5 ± 0.4 | 0.07 | 0.13 ± 0.04 |
| C19:2n-8,14 | 19:2(5,11) | 5,11-nonadecadienoic acid | Not found in LIPID MAPS. | — | 8.67 | — | — | 0.07 ± 0.04 |
| C20:2n-6,9 | 20:2(11,14) | 11Z,14Z-eicosadienoic acid | LMFA01031043 | Dihomolinoleic acid | 9.1 | 18 ± 4 | 0.2 | 0.09 ± 0.04 |
| C20:2n-7,13 | 20:2(7,13) | 7,13-eicosadienoic acid | Not found in LIPID MAPS. | — | 9.14 | 51 ± 2 | 0.04 | 0.25 ± 0.05 |
| C20:2n-9,12 | 20:2(8,11) | 8Z,11Z-eicosadienoic acid | LMFA01030377 | 8Z,11Z-eicosadienoic acid | 9.19 | 4 ± 2 | 0.5 | 0.02 ± 0.01 |
| C20:2n-9,15 | 20:2(5,11) | 5Z,11Z-eicosadienoic acid | LMFA01030865 | Keteleeronic acid | 9.29 | 28 ± 4 | 0.2 | 0.13 ± 0.02 |
| C21:2n-8,14 | 21:2(7,13) | 7,13-heneicosadienoic acid | Not found in LIPID MAPS. | — | 9.73 | — | — | 0.05 ± 0.03 |
| C22:2n-7,13 | 22:2(9,15) | 9,15-docosadienoic acid | Not found in LIPID MAPS. | — | 10.24 | 24 ± 3 | 0.1 | 0.05 ± 0.02 |
| C22:2n-9,15 | 22:2(7,13) | 7,13-docosadienoic acid | LMFA04000061 | 22:2(7Z,13Z) | 10.33 | 76 ± 3 | 0.04 | 0.16 ± 0.09 |
| C24:2n-9,15 | 24:2(9,15) | 9,15-tetracosadienoic acid | Not found in LIPID MAPS. | — | 11.36 | — | — | 0.09 ± 0.05 |
| C18:3n-3,6,9 | 18:3(9,12,15) | 9Z,12Z,15Z-octadecatrienoic acid | LMFA01030152 | alpha-Linolenic acid | 7.16 | — | — | 0.13 ± 0.06 |
| C20:3n-6,9,12 | 20:3(8,11,14) | 8Z,11Z,14Z-eicosatrienoic acid | LMFA01030387 | Bishomo-gamma-linolenic acid | 8.41 | 66.8 ± 0.8 | 0.01 | 1.0 ± 0.3 |
| C20:3n-6,9,15 | 20:3(5,11,14) | 5Z,11Z,14Z-eicosatrienoic acid | LMFA01030380 | Sciadonic acid | 8.51 | 20.6 ± 0.4 | 0.02 | 0.3 ± 0.07 |
| C20:3n-9,12,15 | 20:3(5,8,11) | 5Z,8Z,11Z-eicosatrienoic acid | LMFA01030381 | Mead acid | 8.7 | 12.6 ± 0.9 | 0.07 | 0.19 ± 0.06 |
| C21:3n-8,11,14 | 21:3(7,10,13) | 7Z,10Z,13Z-heneicosatrienoic acid | Not found in LIPID MAPS. | — | 9.06 | — | — | 0.03 ± 0.03 |
| C22:3n-9,12,15 | 22:3(7,10,13) | 7Z,10Z,13Z-docosatrienoic acid | LMFA01030411 | Dihomo Mead's acid | 9.68 | — | — | 0.14 ± 0.04 |
| C20:4n-6,9,12,15 | 20:4(5,8,11,14) | 5Z,8Z,11Z,14Z-eicosatetraenoic acid | LMFA01030001 | Arachidonic acid | 7.93 | 99.1 ± 0.02 | 0.0002 | 6 ± 1 |
| C20:4n-3,6,9,15 | 20:4(5,11,14,17) | 5Z,11Z,14Z,17Z-eicosatetraenoic acid | LMFA01030394 | Juniperonic acid | 7.81 | 0.93 ± 0.02 | 0.02 | 0.06 ± 0.01 |
| C22:4n-6,9,12,15 | 22:4(7,10,13,16) | 7Z,10Z,13Z,16Z-docosatetraenoic acid | LMFA01030178 | Adrenic Acid | 8.93 | — | — | 0.3 ± 0.2 |
| C24:4n-6,9,12,15 | 24:4(9,12,15,18) | 9Z,12Z,15Z,18Z-tetracosatetraenoic acid | LMFA01030819 | 9Z,12Z,15Z,18Z-Tetracosatetraenoic acid | 9.97 | — | — | 0.05 ± 0.02 |
| C20:5n-3,6,9,12,15 | 20:5(5,8,11,14,17) | 5Z,8Z,11Z,14Z,17Z-eicosapentaenoic acid | LMFA01030759 | EPA | 7.25 | — | — | 1.1 ± 0.3 |
| C22:5n-3,6,9,12,15 | 22:5(7,10,13,16,19) | 7Z,10Z,13Z,16Z,19Z-docosapentaenoic acid | LMFA04000044 | DPA | 8.27 | 89.3 ± 0.8 | 0.009 | 1.6 ± 0.4 |
| C22:5n-6,9,12,15,18 | 22:5(4,7,10,13,16) | 4Z,7Z,10Z,13Z,16Z-docosapentaenoic acid | LMFA04000064 | 22:5(4Z,7Z,10Z,13Z,16Z) | 8.6 | 10.7 ± 0.8 | 0.08 | 0.19 ± 0.06 |
| C24:5n-3,6,9,12,15 | 24:5(9,12,15,18,21) | 9Z,12Z,15Z,18Z,21Z-tetracosapentaenoic acid | LMFA01030821 | 9Z,12Z,15Z,18Z,21Z-tetracosapentaenoic acid | 9.33 | — | — | 1.4 ± 0.3 |
| C22:6n-3,6,9,12,15,18 | 22:6(4,7,10,13,16,19) | 4Z,7Z,10Z,13Z,16Z,19Z-docosahexaenoic acid | LMFA01030185 | DHA | 7.93 | — | — | 2.8 ± 0.6 |

#### 3.7.2 LNCaP Lymph node carcinoma of the Prostate - Human Prostatic Cancer Cells

LNCaP cells are derived from the left supraclavicular lymph node (metastasis of prostate cancer) of a 50-year-old male patient.

Cell culture, lipid extraction, hydrolysis and fixed charge derivatization (AMPP) was described previously.^41^ The derivatized fatty acids in Methanol were kept at -18°C prior to analysis with the methods described in this study. For direct infusion ESI-MS measurement, 10 μL were injected, while 2.5 μL were injected for LC-OzID-MS analyses (both untargeted and targeted analysis).

**Figure S34:**
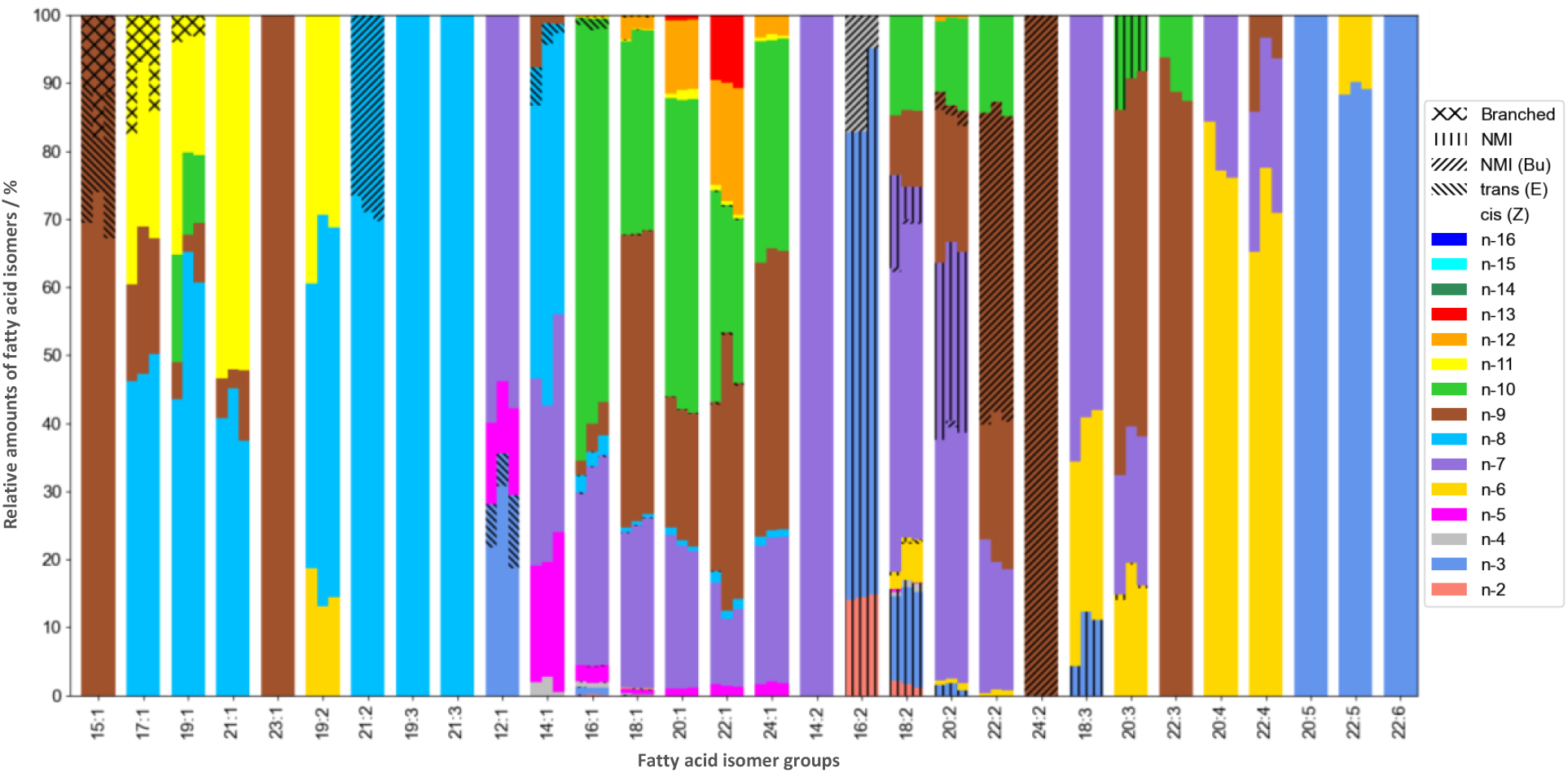
Relative amounts of fatty acid isomers identified in lipid extracts of LNCaP cancer cells (total fatty acid content of hydrolyzed lipids). Shown are the relative amounts of three replicates (three instances of LNCaP cells undergoing cell culture).

**Figure S35:**
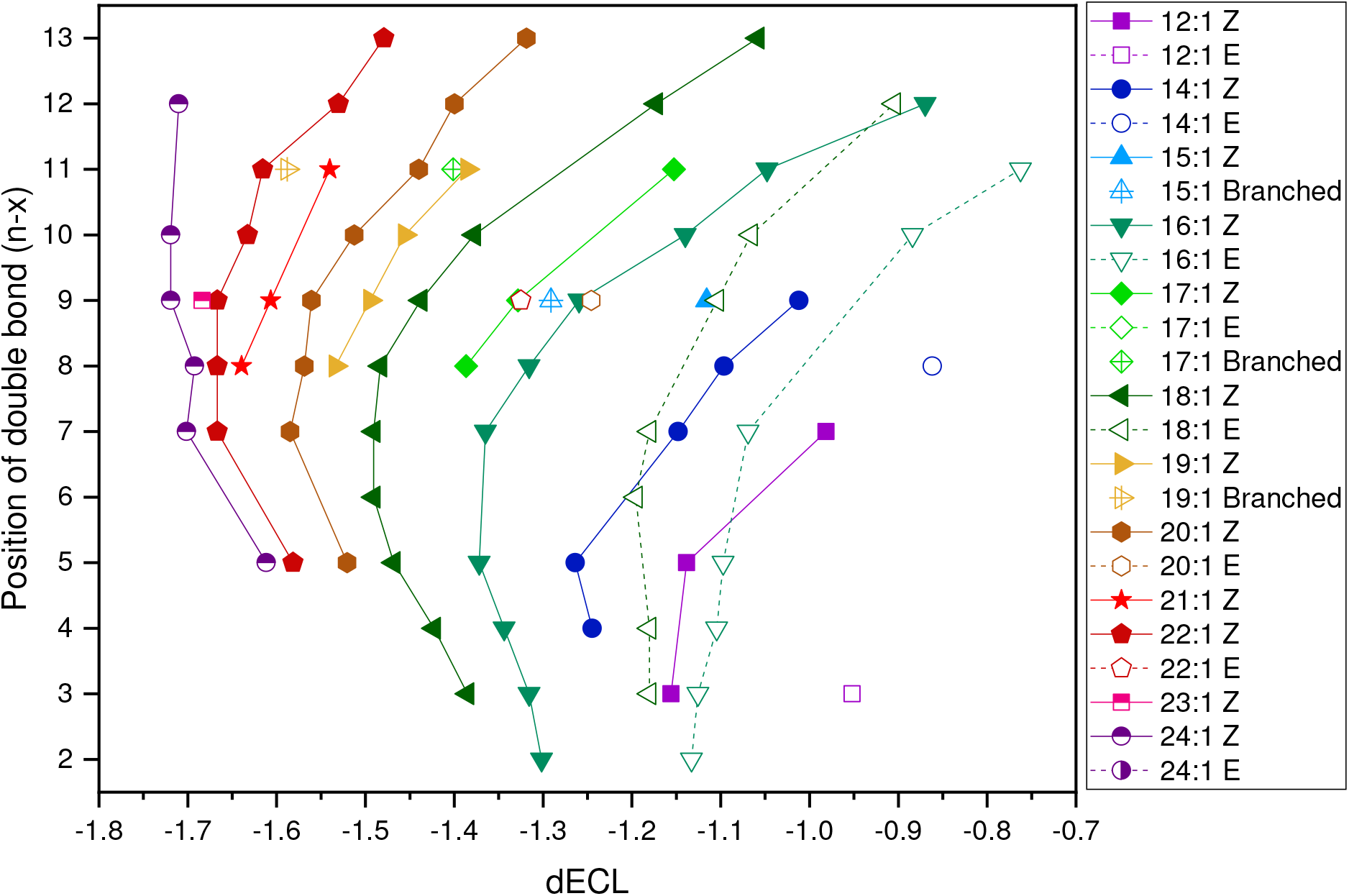
Analysis of differential equivalent chain lengths (dECL) for monounsaturated fatty acids in LNCaP cell extracts. For determination of ECL and dECL values, the equation (ECL = 0.04095 * (RT)^2 + 0.79599 * RT + 6.53741) was determined from retention times of saturated fatty acids in the same sample.

**Figure S36:**
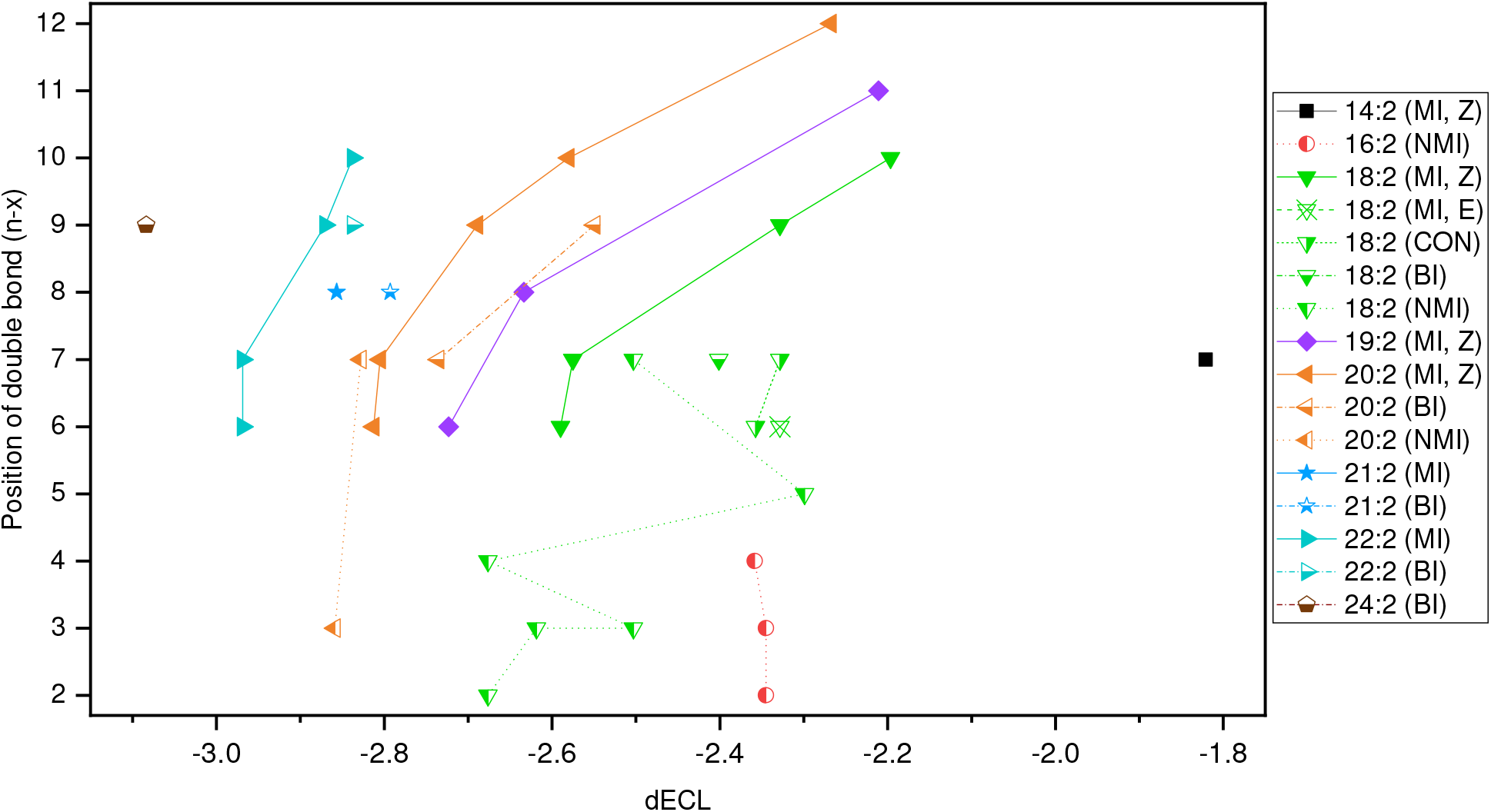
Analysis of differential equivalent chain lengths (dECL) for bisunsaturated fatty acids in LNCaP cell extracts. Non-methylene interrupted fatty acids are assigned as conjugated (CON), butylene-interrupted (BI) or other non-methylene interrupted fatty acids (NMI). Double bond configurations cannot be established unequivocally for all fatty acid species. For determination of ECL and dECL values, the equation (ECL = 0.04095 * (RT)^2 + 0.79599 * RT + 6.53741) was determined from retention times of saturated fatty acids in the same sample.

**Table S9:**
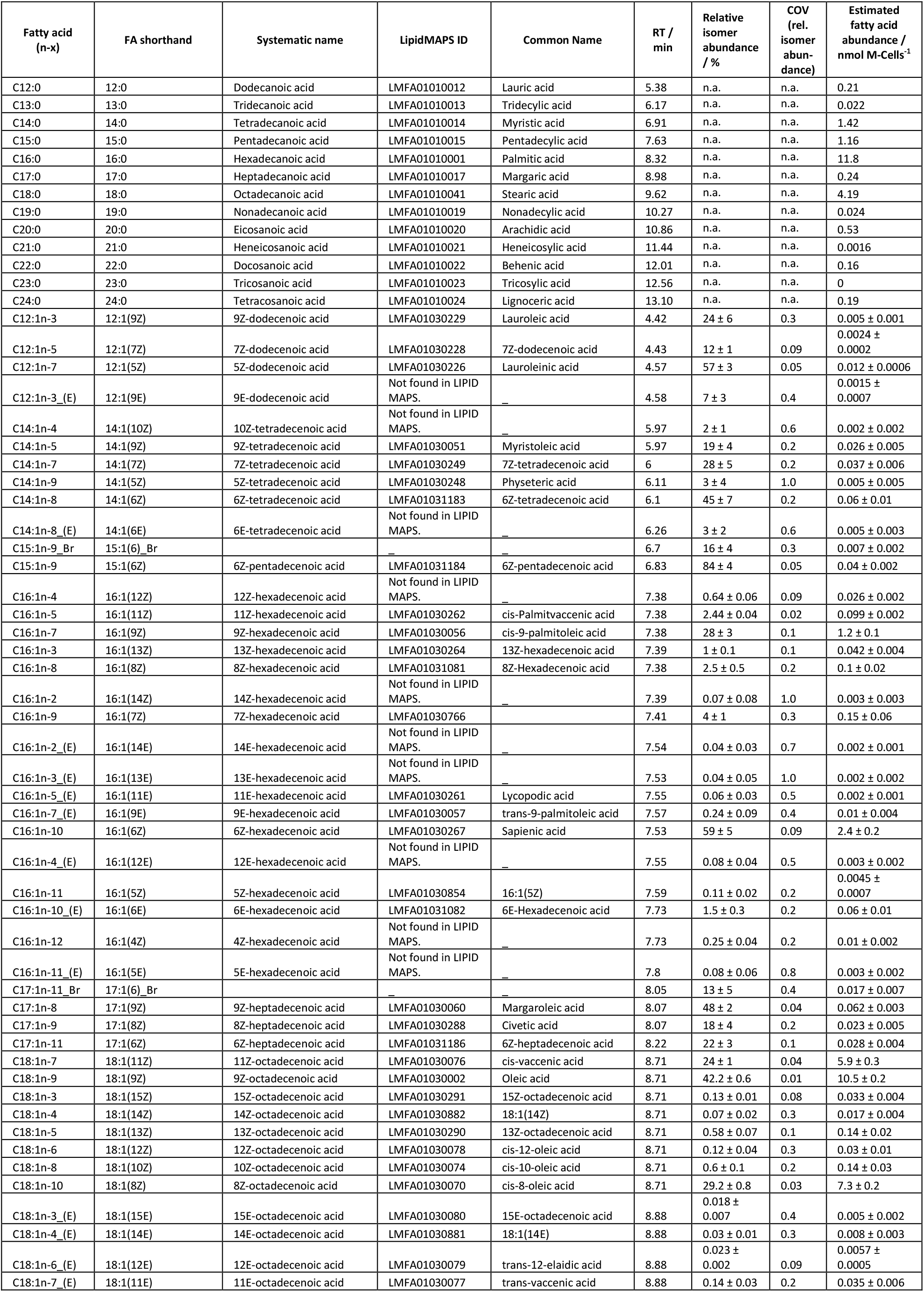
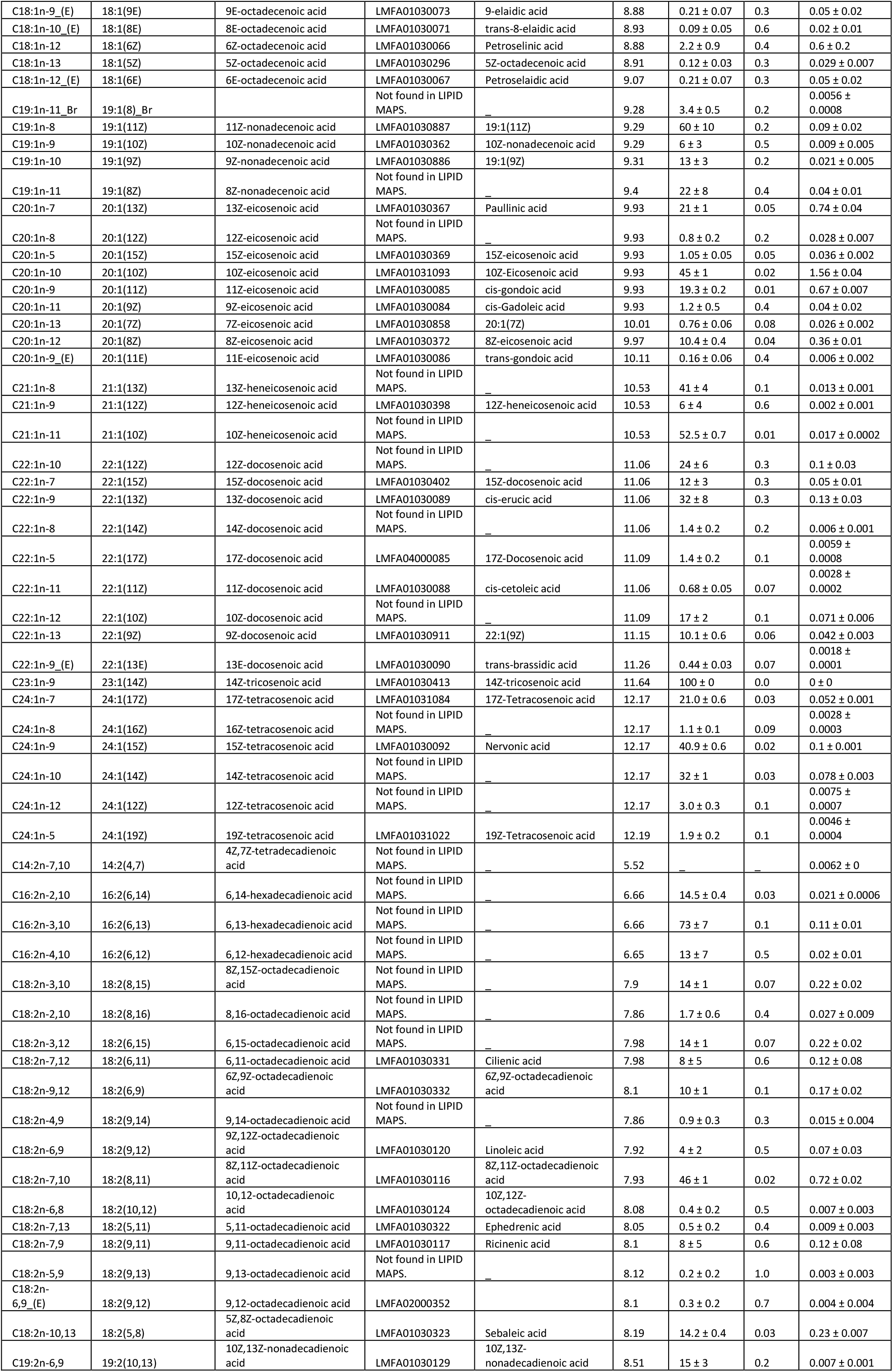
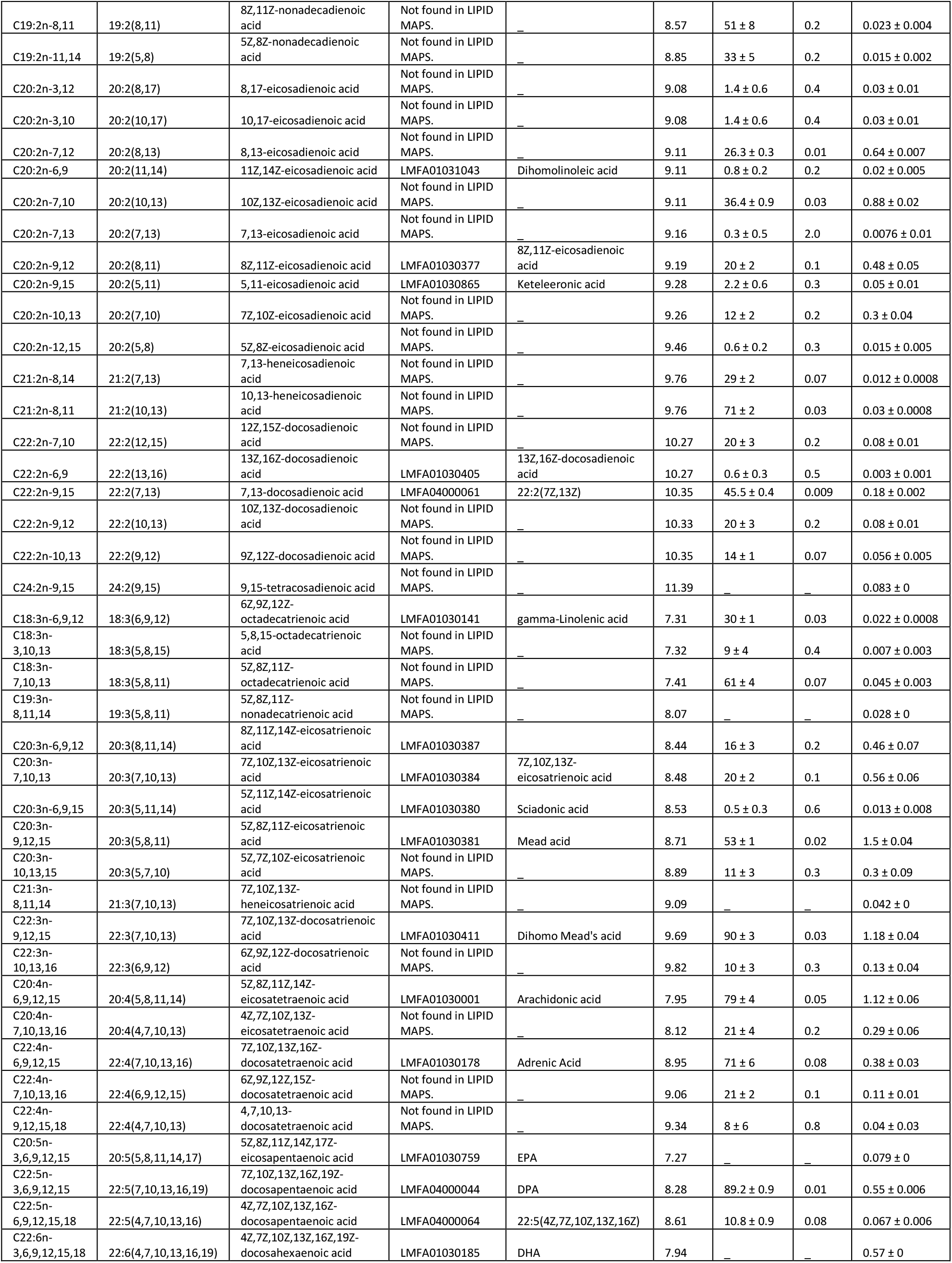
Overview of fatty acids that are identified here in lipid extracts of LNCaP cells. Column seven shows mean values of relative isomer abundances and their associated standard deviations of three replicates of cell cultures of LNCaP cells. Fatty acid abundance is reported here in nmol per million cells (nmol M-Cells^-1^) as an estimate of the fatty acid content of each isomer in the cell culture.

#### 3.7.3 LNCaP_SCD-1i Cancer of the Prostate - Human Prostatic Cancer Cells, SCD-1 inhibited

Cell culture, lipid extraction, hydrolysis and fixed charge derivatization (AMPP) was described previously.^41^ The derivatized fatty acids in Methanol were kept at -18°C prior to analysis with the methods described in this study. For direct infusion ESI-MS measurement, 10 μL were injected, while 2.5 μL were injected for LC-OzID-MS analyses (both untargeted and targeted analysis).

**Figure S37:**
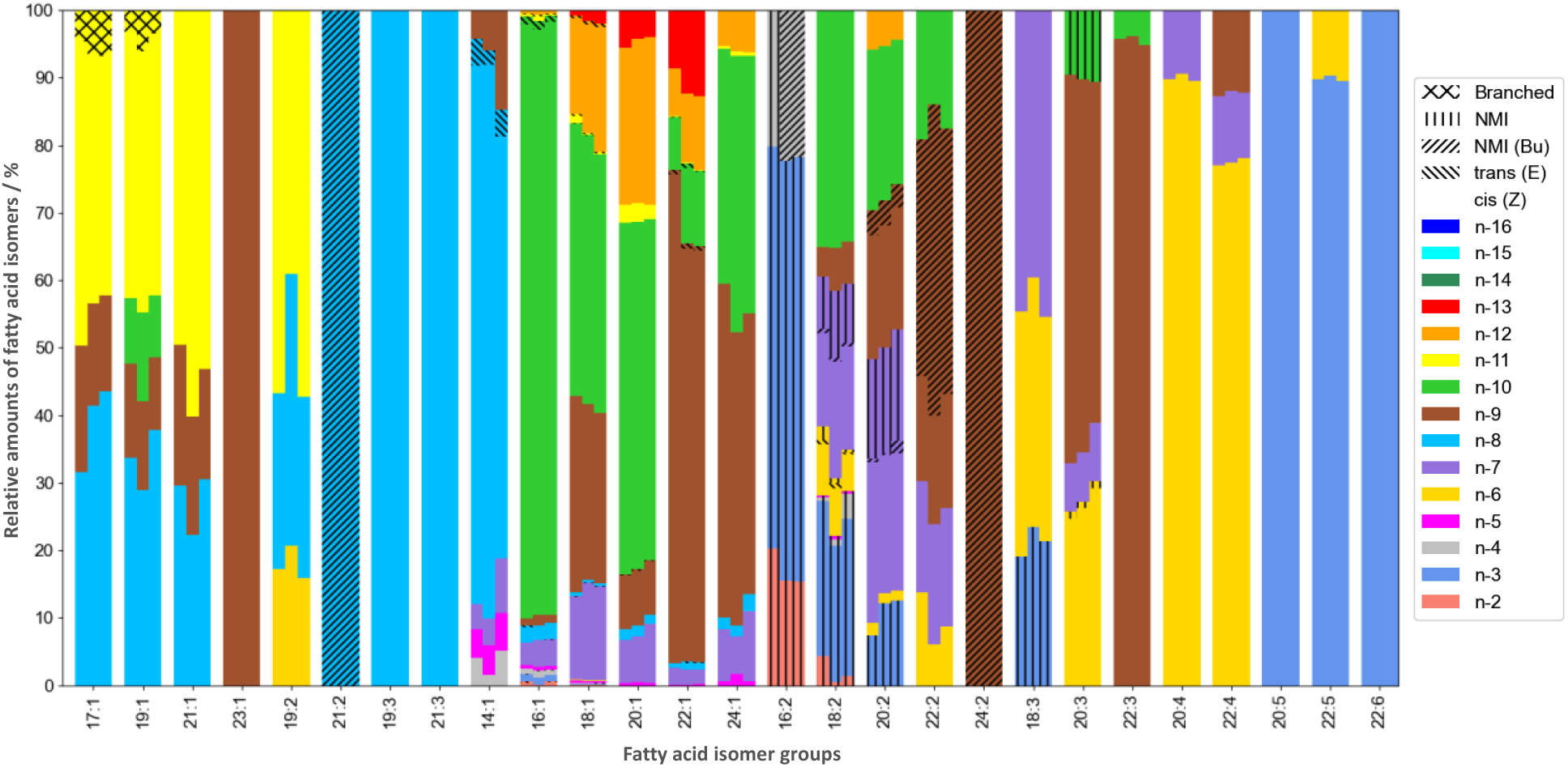
Relative amounts of fatty acid isomers identified in lipid extracts of LNCaP_SCD-1i cancer cells (total fatty acid content of hydrolyzed lipids). Shown are the relative amounts of three replicates (three instances of SCD-1 inhibited LNCaP cells undergoing cell culture).

**Table S10:**
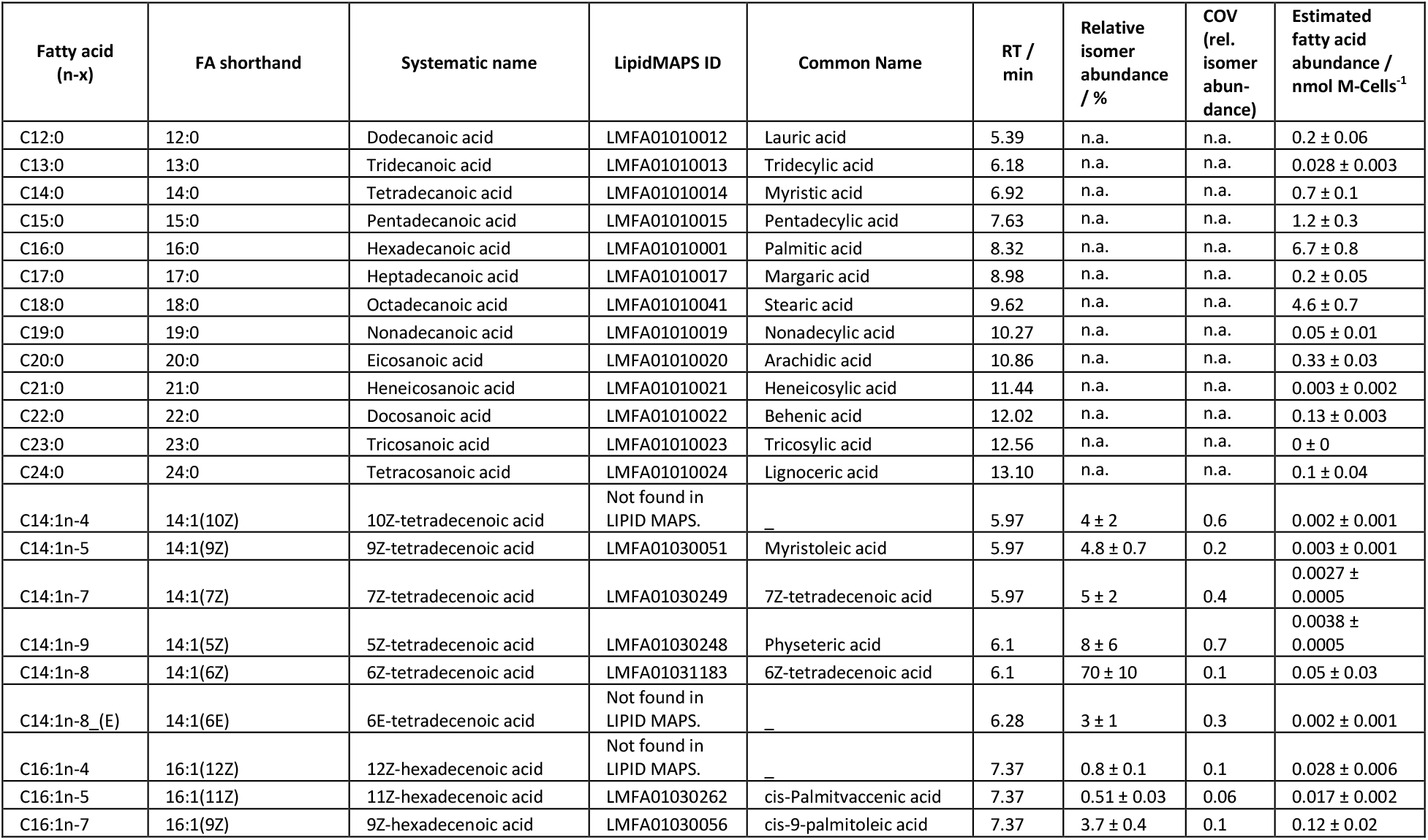
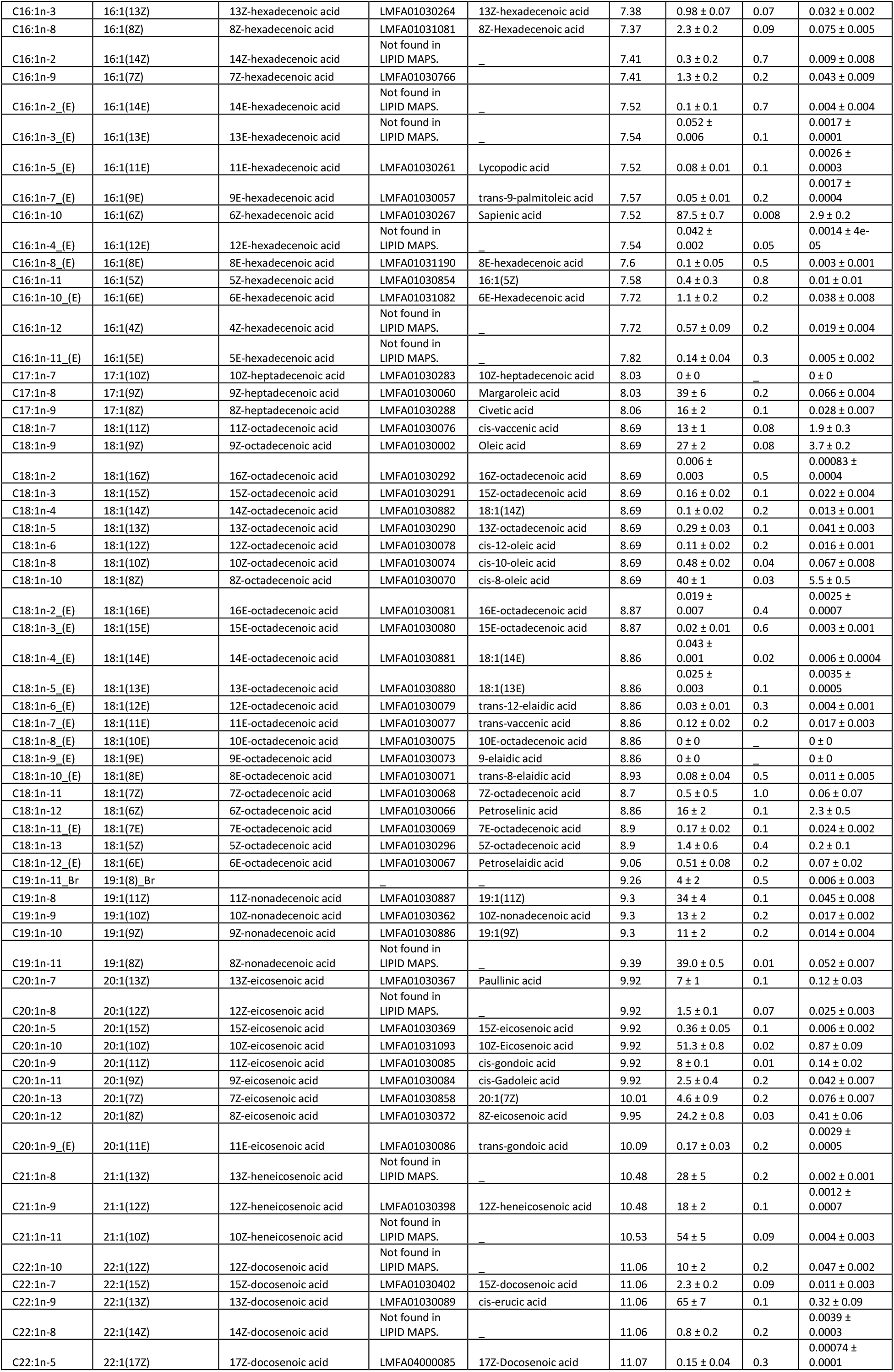
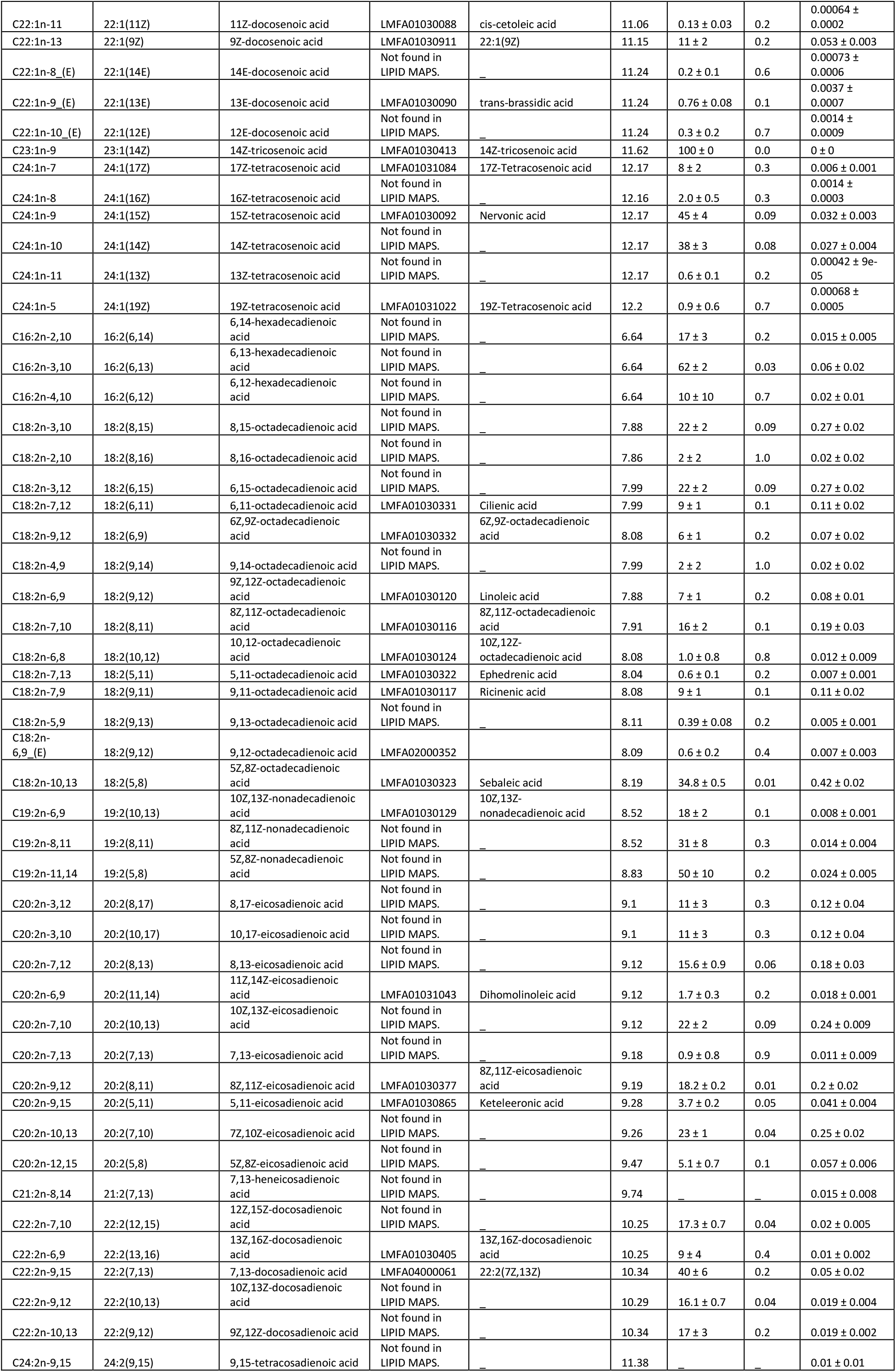
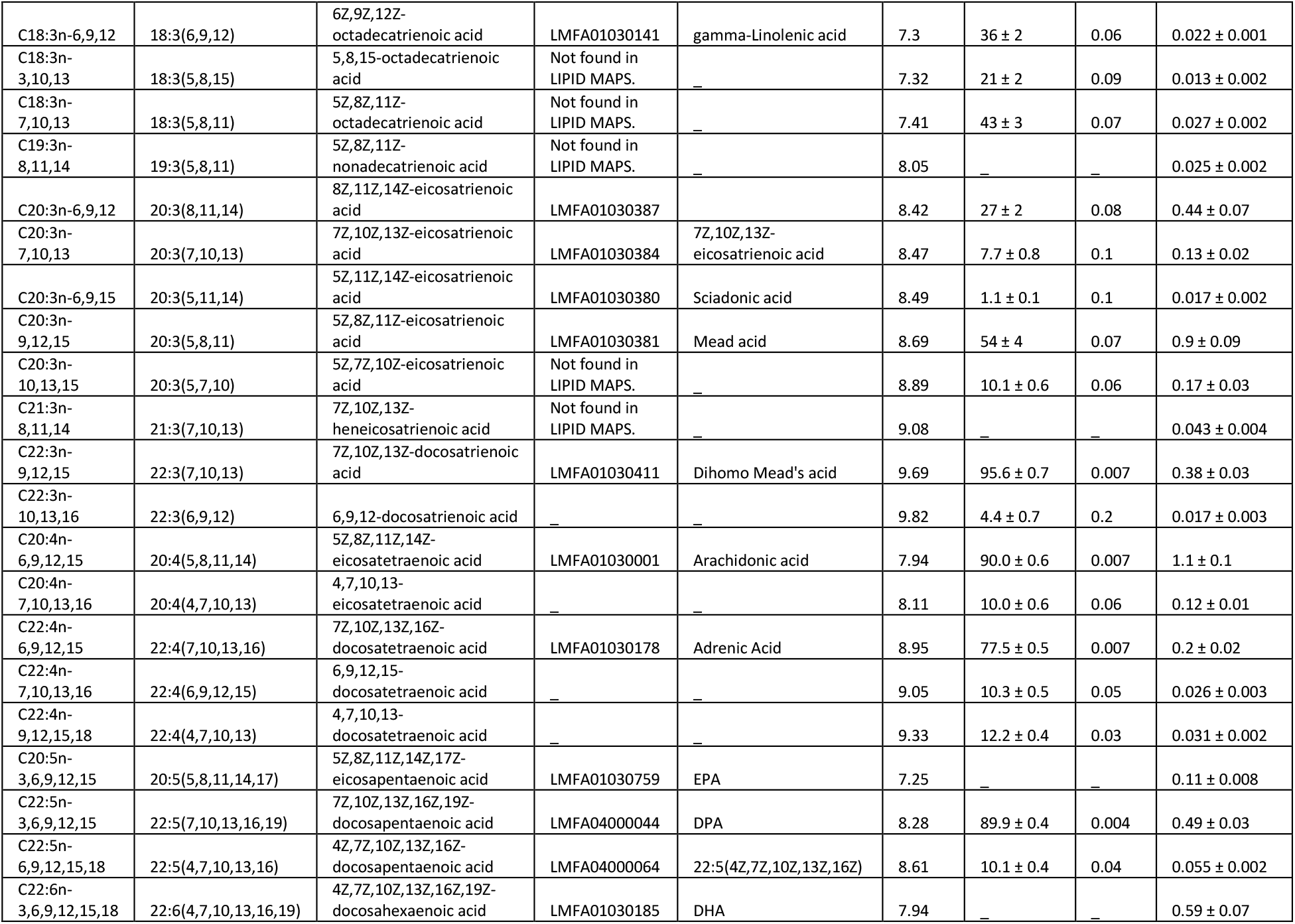
Overview of fatty acids that are identified here in lipid extracts of LNCaP*_*SCD-1i cells. Column seven shows mean values of relative isomer abundances and their associated standard deviations of three replicates of cell cultures of LNCaP*_*SCD-1i cells. Fatty acid abundance is reported here in nmol per million cells (nmol M-Cells^-1^) as an estimate of the fatty acid content of each isomer in the cell culture.

| Fatty acid (n-x) | FA shorthand | Systematic name | LipidMAPS ID | Common Name | RT / min | Relative isomer abundance / % | COV (rel. isomer abundance) | Estimated fatty acid abundance / nmol M-Cells<sup>-1</sup> |
| --- | --- | --- | --- | --- | --- | --- | --- | --- |
| C12:0 | 12:0 | Dodecanoic acid | LMFA01010012 | Lauric acid | 5.39 | n.a. | n.a. | 0.2 ± 0.06 |
| C13:0 | 13:0 | Tridecanoic acid | LMFA01010013 | Tridecylic acid | 6.18 | n.a. | n.a. | 0.028 ± 0.003 |
| C14:0 | 14:0 | Tetradecanoic acid | LMFA01010014 | Myristic acid | 6.92 | n.a. | n.a. | 0.7 ± 0.1 |
| C15:0 | 15:0 | Pentadecanoic acid | LMFA01010015 | Pentadecylic acid | 7.63 | n.a. | n.a. | 1.2 ± 0.3 |
| C16:0 | 16:0 | Hexadecanoic acid | LMFA01010001 | Palmitic acid | 8.32 | n.a. | n.a. | 6.7 ± 0.8 |
| C17:0 | 17:0 | Heptadecanoic acid | LMFA01010017 | Margaric acid | 8.98 | n.a. | n.a. | 0.2 ± 0.05 |
| C18:0 | 18:0 | Octadecanoic acid | LMFA01010041 | Stearic acid | 9.62 | n.a. | n.a. | 4.6 ± 0.7 |
| C19:0 | 19:0 | Nonadecanoic acid | LMFA01010019 | Nonadecylic acid | 10.27 | n.a. | n.a. | 0.05 ± 0.01 |
| C20:0 | 20:0 | Eicosanoic acid | LMFA01010020 | Arachidic acid | 10.86 | n.a. | n.a. | 0.33 ± 0.03 |
| C21:0 | 21:0 | Heneicosanoic acid | LMFA01010021 | Heneicosylic acid | 11.44 | n.a. | n.a. | 0.003 ± 0.002 |
| C22:0 | 22:0 | Docosanoic acid | LMFA01010022 | Behenic acid | 12.02 | n.a. | n.a. | 0.13 ± 0.003 |
| C23:0 | 23:0 | Tricosanoic acid | LMFA01010023 | Tricosylic acid | 12.56 | n.a. | n.a. | 0 ± 0 |
| C24:0 | 24:0 | Tetracosanoic acid | LMFA01010024 | Lignoceric acid | 13.10 | n.a. | n.a. | 0.1 ± 0.04 |
| C14:1n-4 | 14:1(10Z) | 10Z-tetradecenoic acid | Not found in LIPID MAPS. |  | 5.97 | 4 ± 2 | 0.6 | 0.002 ± 0.001 |
| C14:1n-5 | 14:1(9Z) | 9Z-tetradecenoic acid | LMFA01030051 | Myristoleic acid | 5.97 | 4.8 ± 0.7 | 0.2 | 0.003 ± 0.001 |
| C14:1n-7 | 14:1(7Z) | 7Z-tetradecenoic acid | LMFA01030249 | 7Z-tetradecenoic acid | 5.97 | 5 ± 2 | 0.4 | 0.0027 ± 0.0005 |
| C14:1n-9 | 14:1(5Z) | 5Z-tetradecenoic acid | LMFA01030248 | Phytseteric acid | 6.1 | 8 ± 6 | 0.7 | 0.0038 ± 0.0005 |
| C14:1n-8 | 14:1(6Z) | 6Z-tetradecenoic acid | LMFA01031183 | 6Z-tetradecenoic acid | 6.1 | 70 ± 10 | 0.1 | 0.05 ± 0.03 |
| C14:1n-8 (E) | 14:1(6E) | 6E-tetradecenoic acid | Not found in LIPID MAPS. |  | 6.28 | 3 ± 1 | 0.3 | 0.002 ± 0.001 |
| C16:1n-4 | 16:1(12Z) | 12Z-hexadecenoic acid | Not found in LIPID MAPS. |  | 7.37 | 0.8 ± 0.1 | 0.1 | 0.028 ± 0.006 |
| C16:1n-5 | 16:1(11Z) | 11Z-hexadecenoic acid | LMFA01030262 | cis-Palmitvaccenic acid | 7.37 | 0.51 ± 0.03 | 0.06 | 0.017 ± 0.002 |
| C16:1n-7 | 16:1(9Z) | 9Z-hexadecenoic acid | LMFA01030056 | cis-9-palmitoleic acid | 7.37 | 3.7 ± 0.4 | 0.1 | 0.12 ± 0.02 |
| C16:1n-3 | 16:1(13Z) | 13Z-hexadecenoic acid | LMFA01030264 | 13Z-hexadecenoic acid | 7.38 | 0.98 ± 0.07 | 0.07 | 0.032 ± 0.002 |
| C16:1n-8 | 16:1(8Z) | 8Z-hexadecenoic acid | LMFA01031081 | 8Z-Hexadecenoic acid | 7.37 | 2.3 ± 0.2 | 0.09 | 0.075 ± 0.005 |
| C16:1n-2 | 16:1(14Z) | 14Z-hexadecenoic acid | Not found in<br>LIPID MAPS. | = | 7.41 | 0.3 ± 0.2 | 0.7 | 0.009 ± 0.008 |
| C16:1n-9 | 16:1(7Z) | 7Z-hexadecenoic acid | LMFA01030766 |  | 7.41 | 1.3 ± 0.2 | 0.2 | 0.043 ± 0.009 |
| C16:1n-2_(E) | 16:1(14E) | 14E-hexadecenoic acid | Not found in<br>LIPID MAPS. | = | 7.52 | 0.1 ± 0.1 | 0.7 | 0.004 ± 0.004 |
| C16:1n-3_(E) | 16:1(13E) | 13E-hexadecenoic acid | Not found in<br>LIPID MAPS. | = | 7.54 | 0.052 ±<br>0.006 | 0.1 | 0.0017 ±<br>0.0001 |
| C16:1n-5_(E) | 16:1(11E) | 11E-hexadecenoic acid | LMFA01030261 | Lycopodic acid | 7.52 | 0.08 ± 0.01 | 0.1 | 0.0026 ±<br>0.0003 |
| C16:1n-7_(E) | 16:1(9E) | 9E-hexadecenoic acid | LMFA01030057 | trans-9-palmitoleic acid | 7.57 | 0.05 ± 0.01 | 0.2 | 0.0017 ±<br>0.0004 |
| C16:1n-10 | 16:1(6Z) | 6Z-hexadecenoic acid | LMFA01030267 | Sapientic acid | 7.52 | 87.5 ± 0.7 | 0.008 | 2.9 ± 0.2 |
| C16:1n-4_(E) | 16:1(12E) | 12E-hexadecenoic acid | Not found in<br>LIPID MAPS. | = | 7.54 | 0.042 ±<br>0.002 | 0.05 | 0.0014 ± 4e-05 |
| C16:1n-8_(E) | 16:1(8E) | 8E-hexadecenoic acid | LMFA01031190 | 8E-hexadecenoic acid | 7.6 | 0.1 ± 0.05 | 0.5 | 0.003 ± 0.001 |
| C16:1n-11 | 16:1(5Z) | 5Z-hexadecenoic acid | LMFA01030854 | 16:1(5Z) | 7.58 | 0.4 ± 0.3 | 0.8 | 0.01 ± 0.01 |
| C16:1n-10_(E) | 16:1(6E) | 6E-hexadecenoic acid | LMFA01031082 | 6E-Hexadecenoic acid | 7.72 | 1.1 ± 0.2 | 0.2 | 0.038 ± 0.008 |
| C16:1n-12 | 16:1(4Z) | 4Z-hexadecenoic acid | Not found in<br>LIPID MAPS. | = | 7.72 | 0.57 ± 0.09 | 0.2 | 0.019 ± 0.004 |
| C16:1n-11_(E) | 16:1(5E) | 5E-hexadecenoic acid | Not found in<br>LIPID MAPS. | = | 7.82 | 0.14 ± 0.04 | 0.3 | 0.005 ± 0.002 |
| C17:1n-7 | 17:1(10Z) | 10Z-heptadecenoic acid | LMFA01030283 | 10Z-heptadecenoic acid | 8.03 | 0 ± 0 | = | 0 ± 0 |
| C17:1n-8 | 17:1(9Z) | 9Z-heptadecenoic acid | LMFA01030060 | Margaroleic acid | 8.03 | 39 ± 6 | 0.2 | 0.066 ± 0.004 |
| C17:1n-9 | 17:1(8Z) | 8Z-heptadecenoic acid | LMFA01030288 | Civetic acid | 8.06 | 16 ± 2 | 0.1 | 0.028 ± 0.007 |
| C18:1n-7 | 18:1(11Z) | 11Z-octadecenoic acid | LMFA01030076 | cis-vaccenic acid | 8.69 | 13 ± 1 | 0.08 | 1.9 ± 0.3 |
| C18:1n-9 | 18:1(9Z) | 9Z-octadecenoic acid | LMFA01030002 | Oleic acid | 8.69 | 27 ± 2 | 0.08 | 3.7 ± 0.2 |
| C18:1n-2 | 18:1(16Z) | 16Z-octadecenoic acid | LMFA01030292 | 16Z-octadecenoic acid | 8.69 | 0.006 ±<br>0.003 | 0.5 | 0.00083 ±<br>0.0004 |
| C18:1n-3 | 18:1(15Z) | 15Z-octadecenoic acid | LMFA01030291 | 15Z-octadecenoic acid | 8.69 | 0.16 ± 0.02 | 0.1 | 0.022 ± 0.004 |
| C18:1n-4 | 18:1(14Z) | 14Z-octadecenoic acid | LMFA01030882 | 18:1(14Z) | 8.69 | 0.1 ± 0.02 | 0.2 | 0.013 ± 0.001 |
| C18:1n-5 | 18:1(13Z) | 13Z-octadecenoic acid | LMFA01030290 | 13Z-octadecenoic acid | 8.69 | 0.29 ± 0.03 | 0.1 | 0.041 ± 0.003 |
| C18:1n-6 | 18:1(12Z) | 12Z-octadecenoic acid | LMFA01030078 | cis-12-oleic acid | 8.69 | 0.11 ± 0.02 | 0.2 | 0.016 ± 0.001 |
| C18:1n-8 | 18:1(10Z) | 10Z-octadecenoic acid | LMFA01030074 | cis-10-oleic acid | 8.69 | 0.48 ± 0.02 | 0.04 | 0.067 ± 0.008 |
| C18:1n-10 | 18:1(8Z) | 8Z-octadecenoic acid | LMFA01030070 | cis-8-oleic acid | 8.69 | 40 ± 1 | 0.03 | 5.5 ± 0.5 |
| C18:1n-2_(E) | 18:1(16E) | 16E-octadecenoic acid | LMFA01030081 | 16E-octadecenoic acid | 8.87 | 0.019 ±<br>0.007 | 0.4 | 0.0025 ±<br>0.0007 |
| C18:1n-3_(E) | 18:1(15E) | 15E-octadecenoic acid | LMFA01030080 | 15E-octadecenoic acid | 8.87 | 0.02 ± 0.01 | 0.6 | 0.003 ± 0.001 |
| C18:1n-4_(E) | 18:1(14E) | 14E-octadecenoic acid | LMFA01030881 | 18:1(14E) | 8.86 | 0.043 ±<br>0.001 | 0.02 | 0.006 ± 0.0004 |
| C18:1n-5_(E) | 18:1(13E) | 13E-octadecenoic acid | LMFA01030880 | 18:1(13E) | 8.86 | 0.025 ±<br>0.003 | 0.1 | 0.0035 ±<br>0.0005 |
| C18:1n-6_(E) | 18:1(12E) | 12E-octadecenoic acid | LMFA01030079 | trans-12-elaidic acid | 8.86 | 0.03 ± 0.01 | 0.3 | 0.004 ± 0.001 |
| C18:1n-7_(E) | 18:1(11E) | 11E-octadecenoic acid | LMFA01030077 | trans-vaccenic acid | 8.86 | 0.12 ± 0.02 | 0.2 | 0.017 ± 0.003 |
| C18:1n-8_(E) | 18:1(10E) | 10E-octadecenoic acid | LMFA01030075 | 10E-octadecenoic acid | 8.86 | 0 ± 0 | = | 0 ± 0 |
| C18:1n-9_(E) | 18:1(9E) | 9E-octadecenoic acid | LMFA01030073 | 9-elaidic acid | 8.86 | 0 ± 0 | = | 0 ± 0 |
| C18:1n-10_(E) | 18:1(8E) | 8E-octadecenoic acid | LMFA01030071 | trans-8-elaidic acid | 8.93 | 0.08 ± 0.04 | 0.5 | 0.011 ± 0.005 |
| C18:1n-11 | 18:1(7Z) | 7Z-octadecenoic acid | LMFA01030068 | 7Z-octadecenoic acid | 8.7 | 0.5 ± 0.5 | 1.0 | 0.06 ± 0.07 |
| C18:1n-12 | 18:1(6Z) | 6Z-octadecenoic acid | LMFA01030066 | Petroselinic acid | 8.86 | 16 ± 2 | 0.1 | 2.3 ± 0.5 |
| C18:1n-11_(E) | 18:1(7E) | 7E-octadecenoic acid | LMFA01030069 | 7E-octadecenoic acid | 8.9 | 0.17 ± 0.02 | 0.1 | 0.024 ± 0.002 |
| C18:1n-13 | 18:1(5Z) | 5Z-octadecenoic acid | LMFA01030296 | 5Z-octadecenoic acid | 8.9 | 1.4 ± 0.6 | 0.4 | 0.2 ± 0.1 |
| C18:1n-12_(E) | 18:1(6E) | 6E-octadecenoic acid | LMFA01030067 | Petroselaidic acid | 9.06 | 0.51 ± 0.08 | 0.2 | 0.07 ± 0.02 |
| C19:1n-11_Br | 19:1(8)_Br |  | = | = | 9.26 | 4 ± 2 | 0.5 | 0.006 ± 0.003 |
| C19:1n-8 | 19:1(11Z) | 11Z-nonadecenoic acid | LMFA01030887 | 19:1(11Z) | 9.3 | 34 ± 4 | 0.1 | 0.045 ± 0.008 |
| C19:1n-9 | 19:1(10Z) | 10Z-nonadecenoic acid | LMFA01030362 | 10Z-nonadecenoic acid | 9.3 | 13 ± 2 | 0.2 | 0.017 ± 0.002 |
| C19:1n-10 | 19:1(9Z) | 9Z-nonadecenoic acid | LMFA01030886 | 19:1(9Z) | 9.3 | 11 ± 2 | 0.2 | 0.014 ± 0.004 |
| C19:1n-11 | 19:1(8Z) | 8Z-nonadecenoic acid | Not found in<br>LIPID MAPS. |  | 9.39 | 39.0 ± 0.5 | 0.01 | 0.052 ± 0.007 |
| C20:1n-7 | 20:1(13Z) | 13Z-eicosenoic acid | LMFA01030367 | Paullinic acid | 9.92 | 7 ± 1 | 0.1 | 0.12 ± 0.03 |
| C20:1n-8 | 20:1(12Z) | 12Z-eicosenoic acid | Not found in<br>LIPID MAPS. | = | 9.92 | 1.5 ± 0.1 | 0.07 | 0.025 ± 0.003 |
| C20:1n-5 | 20:1(15Z) | 15Z-eicosenoic acid | LMFA01030369 | 15Z-eicosenoic acid | 9.92 | 0.36 ± 0.05 | 0.1 | 0.006 ± 0.002 |
| C20:1n-10 | 20:1(10Z) | 10Z-eicosenoic acid | LMFA01031093 | 10Z-Eicosenoic acid | 9.92 | 51.3 ± 0.8 | 0.02 | 0.87 ± 0.09 |
| C20:1n-9 | 20:1(11Z) | 11Z-eicosenoic acid | LMFA01030085 | cis-gondoic acid | 9.92 | 8 ± 0.1 | 0.01 | 0.14 ± 0.02 |
| C20:1n-11 | 20:1(9Z) | 9Z-eicosenoic acid | LMFA01030084 | cis-Gadoleic acid | 9.92 | 2.5 ± 0.4 | 0.2 | 0.042 ± 0.007 |
| C20:1n-13 | 20:1(7Z) | 7Z-eicosenoic acid | LMFA01030858 | 20:1(7Z) | 10.01 | 4.6 ± 0.9 | 0.2 | 0.076 ± 0.007 |
| C20:1n-12 | 20:1(8Z) | 8Z-eicosenoic acid | LMFA01030372 | 8Z-eicosenoic acid | 9.95 | 24.2 ± 0.8 | 0.03 | 0.41 ± 0.06 |
| C20:1n-9_(E) | 20:1(11E) | 11E-eicosenoic acid | LMFA01030086 | trans-gondoic acid | 10.09 | 0.17 ± 0.03 | 0.2 | 0.0029 ±<br>0.0005 |
| C21:1n-8 | 21:1(13Z) | 13Z-heneicosenoic acid | Not found in<br>LIPID MAPS. | = | 10.48 | 28 ± 5 | 0.2 | 0.002 ± 0.001 |
| C21:1n-9 | 21:1(12Z) | 12Z-heneicosenoic acid | LMFA01030398 | 12Z-heneicosenoic acid | 10.48 | 18 ± 2 | 0.1 | 0.0012 ±<br>0.0007 |
| C21:1n-11 | 21:1(10Z) | 10Z-heneicosenoic acid | Not found in<br>LIPID MAPS. | = | 10.53 | 54 ± 5 | 0.09 | 0.004 ± 0.003 |
| C22:1n-10 | 22:1(12Z) | 12Z-docosenoic acid | Not found in<br>LIPID MAPS. | = | 11.06 | 10 ± 2 | 0.2 | 0.047 ± 0.002 |
| C22:1n-7 | 22:1(15Z) | 15Z-docosenoic acid | LMFA01030402 | 15Z-docosenoic acid | 11.06 | 2.3 ± 0.2 | 0.09 | 0.011 ± 0.003 |
| C22:1n-9 | 22:1(13Z) | 13Z-docosenoic acid | LMFA01030089 | cis-erucic acid | 11.06 | 65 ± 7 | 0.1 | 0.32 ± 0.09 |
| C22:1n-8 | 22:1(14Z) | 14Z-docosenoic acid | Not found in<br>LIPID MAPS. | = | 11.06 | 0.8 ± 0.2 | 0.2 | 0.0039 ±<br>0.0003 |
| C22:1n-5 | 22:1(17Z) | 17Z-docosenoic acid | LMFA04000085 | 17Z-Docosenoic acid | 11.07 | 0.15 ± 0.04 | 0.3 | 0.00074 ±<br>0.0001 |
| C22:1n-11 | 22:1(11Z) | 11Z-docosenoic acid | LMFA01030088 | cis-cetoleic acid | 11.06 | 0.13 ± 0.03 | 0.2 | 0.00064 ± 0.0002 |
| C22:1n-13 | 22:1(9Z) | 9Z-docosenoic acid | LMFA01030911 | 22:1(9Z) | 11.15 | 11 ± 2 | 0.2 | 0.053 ± 0.003 |
| C22:1n-8_(E) | 22:1(14E) | 14E-docosenoic acid | Not found in LIPID MAPS. | = | 11.24 | 0.2 ± 0.1 | 0.6 | 0.00073 ± 0.0006 |
| C22:1n-9_(E) | 22:1(13E) | 13E-docosenoic acid | LMFA01030090 | trans-brassicidic acid | 11.24 | 0.76 ± 0.08 | 0.1 | 0.0037 ± 0.0007 |
| C22:1n-10_(E) | 22:1(12E) | 12E-docosenoic acid | Not found in LIPID MAPS. | = | 11.24 | 0.3 ± 0.2 | 0.7 | 0.0014 ± 0.0009 |
| C23:1n-9 | 23:1(14Z) | 14Z-tricosenoic acid | LMFA01030413 | 14Z-tricosenoic acid | 11.62 | 100 ± 0 | 0.0 | 0 ± 0 |
| C24:1n-7 | 24:1(17Z) | 17Z-tetracosenoic acid | LMFA01031084 | 17Z-Tetracosenoic acid | 12.17 | 8 ± 2 | 0.3 | 0.006 ± 0.001 |
| C24:1n-8 | 24:1(16Z) | 16Z-tetracosenoic acid | Not found in LIPID MAPS. | = | 12.16 | 2.0 ± 0.5 | 0.3 | 0.0014 ± 0.0003 |
| C24:1n-9 | 24:1(15Z) | 15Z-tetracosenoic acid | LMFA01030092 | Nervonic acid | 12.17 | 45 ± 4 | 0.09 | 0.032 ± 0.003 |
| C24:1n-10 | 24:1(14Z) | 14Z-tetracosenoic acid | Not found in LIPID MAPS. | = | 12.17 | 38 ± 3 | 0.08 | 0.027 ± 0.004 |
| C24:1n-11 | 24:1(13Z) | 13Z-tetracosenoic acid | Not found in LIPID MAPS. | = | 12.17 | 0.6 ± 0.1 | 0.2 | 0.00042 ± 9e-05 |
| C24:1n-5 | 24:1(19Z) | 19Z-tetracosenoic acid | LMFA01031022 | 19Z-Tetracosenoic acid | 12.2 | 0.9 ± 0.6 | 0.7 | 0.00068 ± 0.0005 |
| C16:2n-2,10 | 16:2(6,14) | 6,14-hexadecadienoic acid | Not found in LIPID MAPS. | = | 6.64 | 17 ± 3 | 0.2 | 0.015 ± 0.005 |
| C16:2n-3,10 | 16:2(6,13) | 6,13-hexadecadienoic acid | Not found in LIPID MAPS. | = | 6.64 | 62 ± 2 | 0.03 | 0.06 ± 0.02 |
| C16:2n-4,10 | 16:2(6,12) | 6,12-hexadecadienoic acid | Not found in LIPID MAPS. | = | 6.64 | 10 ± 10 | 0.7 | 0.02 ± 0.01 |
| C18:2n-3,10 | 18:2(8,15) | 8,15-octadecadienoic acid | Not found in LIPID MAPS. | = | 7.88 | 22 ± 2 | 0.09 | 0.27 ± 0.02 |
| C18:2n-2,10 | 18:2(8,16) | 8,16-octadecadienoic acid | Not found in LIPID MAPS. | = | 7.86 | 2 ± 2 | 1.0 | 0.02 ± 0.02 |
| C18:2n-3,12 | 18:2(6,15) | 6,15-octadecadienoic acid | Not found in LIPID MAPS. | = | 7.99 | 22 ± 2 | 0.09 | 0.27 ± 0.02 |
| C18:2n-7,12 | 18:2(6,11) | 6,11-octadecadienoic acid | LMFA01030331 | Cilienic acid | 7.99 | 9 ± 1 | 0.1 | 0.11 ± 0.02 |
| C18:2n-9,12 | 18:2(6,9) | 6Z,9Z-octadecadienoic acid | LMFA01030332 | 6Z,9Z-octadecadienoic acid | 8.08 | 6 ± 1 | 0.2 | 0.07 ± 0.02 |
| C18:2n-4,9 | 18:2(9,14) | 9,14-octadecadienoic acid | Not found in LIPID MAPS. | = | 7.99 | 2 ± 2 | 1.0 | 0.02 ± 0.02 |
| C18:2n-6,9 | 18:2(9,12) | 9Z,12Z-octadecadienoic acid | LMFA01030120 | Linoleic acid | 7.88 | 7 ± 1 | 0.2 | 0.08 ± 0.01 |
| C18:2n-7,10 | 18:2(8,11) | 8Z,11Z-octadecadienoic acid | LMFA01030116 | 8Z,11Z-octadecadienoic acid | 7.91 | 16 ± 2 | 0.1 | 0.19 ± 0.03 |
| C18:2n-6,8 | 18:2(10,12) | 10,12-octadecadienoic acid | LMFA01030124 | 10Z,12Z-octadecadienoic acid | 8.08 | 1.0 ± 0.8 | 0.8 | 0.012 ± 0.009 |
| C18:2n-7,13 | 18:2(5,11) | 5,11-octadecadienoic acid | LMFA01030322 | Ephedrenic acid | 8.04 | 0.6 ± 0.1 | 0.2 | 0.007 ± 0.001 |
| C18:2n-7,9 | 18:2(9,11) | 9,11-octadecadienoic acid | LMFA01030117 | Ricinenic acid | 8.08 | 9 ± 1 | 0.1 | 0.11 ± 0.02 |
| C18:2n-5,9 | 18:2(9,13) | 9,13-octadecadienoic acid | Not found in LIPID MAPS. | = | 8.11 | 0.39 ± 0.08 | 0.2 | 0.005 ± 0.001 |
| C18:2n-6,9_(E) | 18:2(9,12) | 9,12-octadecadienoic acid | LMFA02000352 |  | 8.09 | 0.6 ± 0.2 | 0.4 | 0.007 ± 0.003 |
| C18:2n-10,13 | 18:2(5,8) | 5Z,8Z-octadecadienoic acid | LMFA01030323 | Sebaleic acid | 8.19 | 34.8 ± 0.5 | 0.01 | 0.42 ± 0.02 |
| C19:2n-6,9 | 19:2(10,13) | 10Z,13Z-nonadecadienoic acid | LMFA01030129 | 10Z,13Z-nonadecadienoic acid | 8.52 | 18 ± 2 | 0.1 | 0.008 ± 0.001 |
| C19:2n-8,11 | 19:2(8,11) | 8Z,11Z-nonadecadienoic acid | Not found in LIPID MAPS. | = | 8.52 | 31 ± 8 | 0.3 | 0.014 ± 0.004 |
| C19:2n-11,14 | 19:2(5,8) | 5Z,8Z-nonadecadienoic acid | Not found in LIPID MAPS. | = | 8.83 | 50 ± 10 | 0.2 | 0.024 ± 0.005 |
| C20:2n-3,12 | 20:2(8,17) | 8,17-eicosadienoic acid | Not found in LIPID MAPS. | = | 9.1 | 11 ± 3 | 0.3 | 0.12 ± 0.04 |
| C20:2n-3,10 | 20:2(10,17) | 10,17-eicosadienoic acid | Not found in LIPID MAPS. | = | 9.1 | 11 ± 3 | 0.3 | 0.12 ± 0.04 |
| C20:2n-7,12 | 20:2(8,13) | 8,13-eicosadienoic acid | Not found in LIPID MAPS. | = | 9.12 | 15.6 ± 0.9 | 0.06 | 0.18 ± 0.03 |
| C20:2n-6,9 | 20:2(11,14) | 11Z,14Z-eicosadienoic acid | LMFA01031043 | Dihomolinoleic acid | 9.12 | 1.7 ± 0.3 | 0.2 | 0.018 ± 0.001 |
| C20:2n-7,10 | 20:2(10,13) | 10Z,13Z-eicosadienoic acid | Not found in LIPID MAPS. | = | 9.12 | 22 ± 2 | 0.09 | 0.24 ± 0.009 |
| C20:2n-7,13 | 20:2(7,13) | 7,13-eicosadienoic acid | Not found in LIPID MAPS. | = | 9.18 | 0.9 ± 0.8 | 0.9 | 0.011 ± 0.009 |
| C20:2n-9,12 | 20:2(8,11) | 8Z,11Z-eicosadienoic acid | LMFA01030377 | 8Z,11Z-eicosadienoic acid | 9.19 | 18.2 ± 0.2 | 0.01 | 0.2 ± 0.02 |
| C20:2n-9,15 | 20:2(5,11) | 5,11-eicosadienoic acid | LMFA01030865 | Keteleeronic acid | 9.28 | 3.7 ± 0.2 | 0.05 | 0.041 ± 0.004 |
| C20:2n-10,13 | 20:2(7,10) | 7Z,10Z-eicosadienoic acid | Not found in LIPID MAPS. | = | 9.26 | 23 ± 1 | 0.04 | 0.25 ± 0.02 |
| C20:2n-12,15 | 20:2(5,8) | 5Z,8Z-eicosadienoic acid | Not found in LIPID MAPS. | = | 9.47 | 5.1 ± 0.7 | 0.1 | 0.057 ± 0.006 |
| C21:2n-8,14 | 21:2(7,13) | 7,13-heneicosadienoic acid | Not found in LIPID MAPS. | = | 9.74 | = | = | 0.015 ± 0.008 |
| C22:2n-7,10 | 22:2(12,15) | 12Z,15Z-docosadienoic acid | Not found in LIPID MAPS. | = | 10.25 | 17.3 ± 0.7 | 0.04 | 0.02 ± 0.005 |
| C22:2n-6,9 | 22:2(13,16) | 13Z,16Z-docosadienoic acid | LMFA01030405 | 13Z,16Z-docosadienoic acid | 10.25 | 9 ± 4 | 0.4 | 0.01 ± 0.002 |
| C22:2n-9,15 | 22:2(7,13) | 7,13-docosadienoic acid | LMFA04000061 | 22:2(7Z,13Z) | 10.34 | 40 ± 6 | 0.2 | 0.05 ± 0.02 |
| C22:2n-9,12 | 22:2(10,13) | 10Z,13Z-docosadienoic acid | Not found in LIPID MAPS. | = | 10.29 | 16.1 ± 0.7 | 0.04 | 0.019 ± 0.004 |
| C22:2n-10,13 | 22:2(9,12) | 9Z,12Z-docosadienoic acid | Not found in LIPID MAPS. | = | 10.34 | 17 ± 3 | 0.2 | 0.019 ± 0.002 |
| C24:2n-9,15 | 24:2(9,15) | 9,15-tetracosadienoic acid | Not found in LIPID MAPS. | = | 11.38 | = | = | 0.01 ± 0.01 |
| C18:3n-6,9,12 | 18:3(6,9,12) | 6Z,9Z,12Z-octadecatrienoic acid | LMFA01030141 | gamma-Linolenic acid | 7.3 | 36 ± 2 | 0.06 | 0.022 ± 0.001 |
| C18:3n-3,10,13 | 18:3(5,8,15) | 5,8,15-octadecatrienoic acid | Not found in LIPID MAPS. | — | 7.32 | 21 ± 2 | 0.09 | 0.013 ± 0.002 |
| C18:3n-7,10,13 | 18:3(5,8,11) | 5Z,8Z,11Z-octadecatrienoic acid | Not found in LIPID MAPS. | — | 7.41 | 43 ± 3 | 0.07 | 0.027 ± 0.002 |
| C19:3n-8,11,14 | 19:3(5,8,11) | 5Z,8Z,11Z-nonadecatrienoic acid | Not found in LIPID MAPS. | — | 8.05 | — | — | 0.025 ± 0.002 |
| C20:3n-6,9,12 | 20:3(8,11,14) | 8Z,11Z,14Z-eicosatrienoic acid | LMFA01030387 | — | 8.42 | 27 ± 2 | 0.08 | 0.44 ± 0.07 |
| C20:3n-7,10,13 | 20:3(7,10,13) | 7Z,10Z,13Z-eicosatrienoic acid | LMFA01030384 | 7Z,10Z,13Z-eicosatrienoic acid | 8.47 | 7.7 ± 0.8 | 0.1 | 0.13 ± 0.02 |
| C20:3n-6,9,15 | 20:3(5,11,14) | 5Z,11Z,14Z-eicosatrienoic acid | LMFA01030380 | Sciadonic acid | 8.49 | 1.1 ± 0.1 | 0.1 | 0.017 ± 0.002 |
| C20:3n-9,12,15 | 20:3(5,8,11) | 5Z,8Z,11Z-eicosatrienoic acid | LMFA01030381 | Mead acid | 8.69 | 54 ± 4 | 0.07 | 0.9 ± 0.09 |
| C20:3n-10,13,15 | 20:3(5,7,10) | 5Z,7Z,10Z-eicosatrienoic acid | Not found in LIPID MAPS. | — | 8.89 | 10.1 ± 0.6 | 0.06 | 0.17 ± 0.03 |
| C21:3n-8,11,14 | 21:3(7,10,13) | 7Z,10Z,13Z-heneicosatrienoic acid | Not found in LIPID MAPS. | — | 9.08 | — | — | 0.043 ± 0.004 |
| C22:3n-9,12,15 | 22:3(7,10,13) | 7Z,10Z,13Z-docosatrienoic acid | LMFA01030411 | Dihomo Mead's acid | 9.69 | 95.6 ± 0.7 | 0.007 | 0.38 ± 0.03 |
| C22:3n-10,13,16 | 22:3(6,9,12) | 6,9,12-docosatrienoic acid | — | — | 9.82 | 4.4 ± 0.7 | 0.2 | 0.017 ± 0.003 |
| C20:4n-6,9,12,15 | 20:4(5,8,11,14) | 5Z,8Z,11Z,14Z-eicosatetraenoic acid | LMFA01030001 | Arachidonic acid | 7.94 | 90.0 ± 0.6 | 0.007 | 1.1 ± 0.1 |
| C20:4n-7,10,13,16 | 20:4(4,7,10,13) | 4,7,10,13-eicosatetraenoic acid | — | — | 8.11 | 10.0 ± 0.6 | 0.06 | 0.12 ± 0.01 |
| C22:4n-6,9,12,15 | 22:4(7,10,13,16) | 7Z,10Z,13Z,16Z-docosatetraenoic acid | LMFA01030178 | Adrenic Acid | 8.95 | 77.5 ± 0.5 | 0.007 | 0.2 ± 0.02 |
| C22:4n-7,10,13,16 | 22:4(6,9,12,15) | 6,9,12,15-docosatetraenoic acid | — | — | 9.05 | 10.3 ± 0.5 | 0.05 | 0.026 ± 0.003 |
| C22:4n-9,12,15,18 | 22:4(4,7,10,13) | 4,7,10,13-docosatetraenoic acid | — | — | 9.33 | 12.2 ± 0.4 | 0.03 | 0.031 ± 0.002 |
| C20:5n-3,6,9,12,15 | 20:5(5,8,11,14,17) | 5Z,8Z,11Z,14Z,17Z-eicosapentaenoic acid | LMFA01030759 | EPA | 7.25 | — | — | 0.11 ± 0.008 |
| C22:5n-3,6,9,12,15 | 22:5(7,10,13,16,19) | 7Z,10Z,13Z,16Z,19Z-docosapentaenoic acid | LMFA04000044 | DPA | 8.28 | 89.9 ± 0.4 | 0.004 | 0.49 ± 0.03 |
| C22:5n-6,9,12,15,18 | 22:5(4,7,10,13,16) | 4Z,7Z,10Z,13Z,16Z-docosapentaenoic acid | LMFA04000064 | 22:5(4Z,7Z,10Z,13Z,16Z) | 8.61 | 10.1 ± 0.4 | 0.04 | 0.055 ± 0.002 |
| C22:6n-3,6,9,12,15,18 | 22:6(4,7,10,13,16,19) | 4Z,7Z,10Z,13Z,16Z,19Z-docosahexaenoic acid | LMFA01030185 | DHA | 7.94 | — | — | 0.59 ± 0.07 |

#### 3.7.4 Comparison of relative abundance of fatty acid isomers between cancer cell lines and associated P values

**Figure S38:**
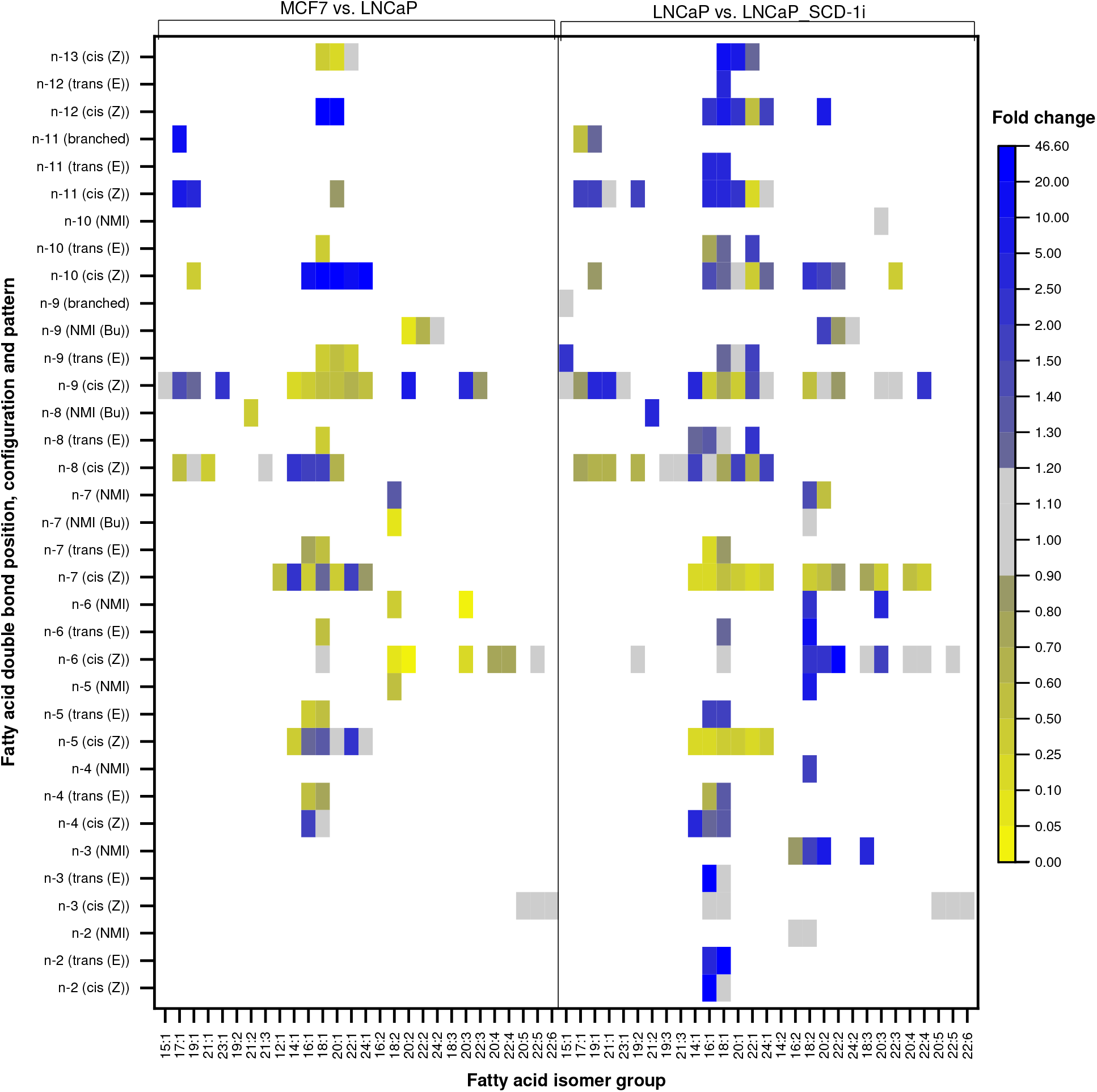
Mean fold change of three biological replicates (n=3) between relative isomer abundances of fatty acid isomers in cancer cell lines MCF7 and LNCaP (left side of the heatmap) as well as between LNCaP and LNCaP_SCD-1i (right side of the heatmap). White colour represents that the respective fatty acid isomer was not detected in either some or all of the six samples. The colour scale represents the mean of the fold changes of relative isomer abundance, with a change of less than ± 10% of the value represented by the grey colour and yellow colours indicating a decrease from MCF7 to LNCaP or LNCaP to LNCaP_SCD-1i. Conversely, blue colours indicate an increase in relative abundance.

**Figure S39:**
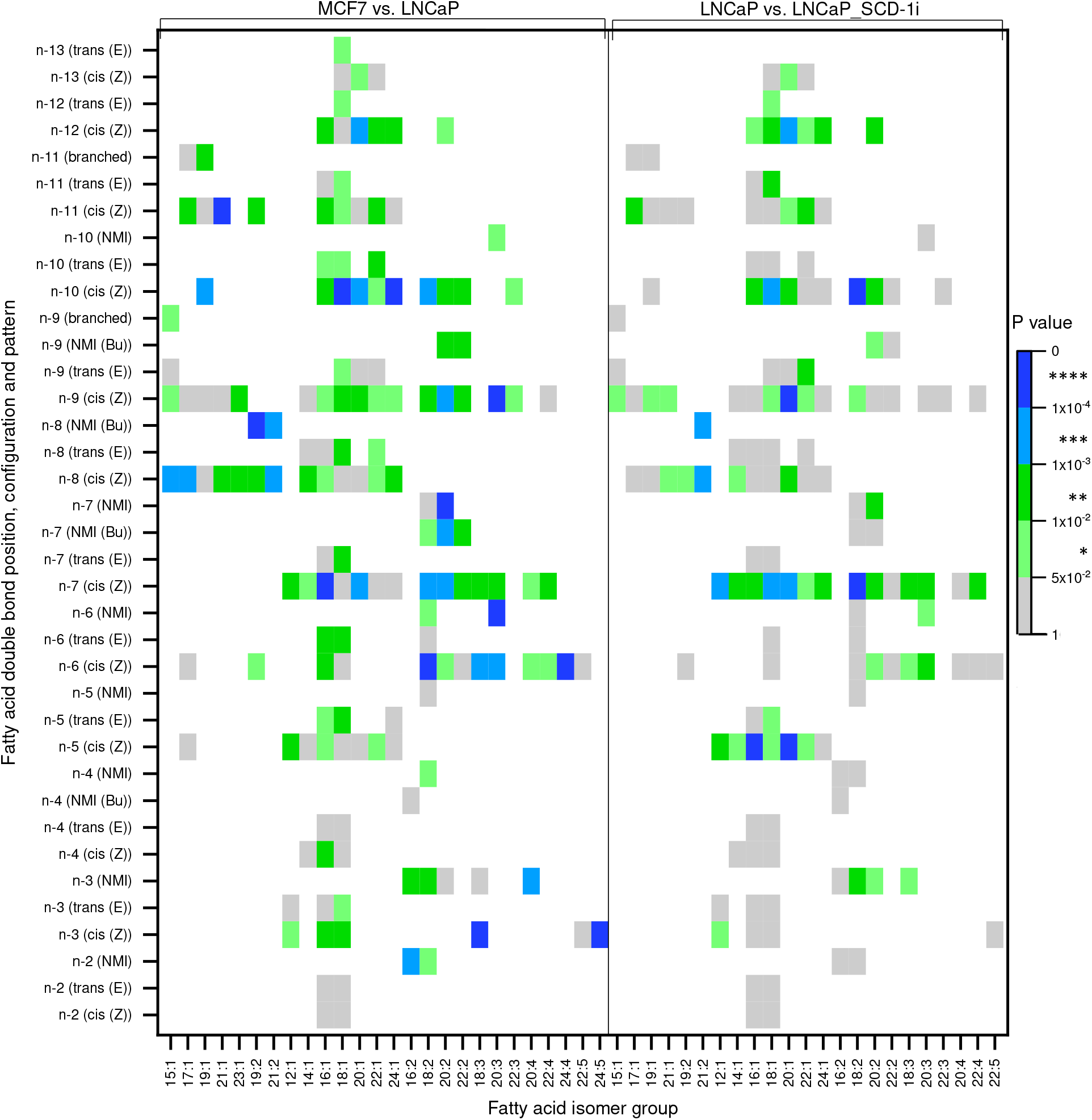
P values of fatty acid isomers according to a two-sided Welsh’s t-test applied each to the three replicate values (n=3) of the cancer cell lines MCF7 and LNCaP (left side of the heatmap) and LNCaP and LNCaP_SCD-1i (right side of the heatmap). White colour represents either that the respective fatty acid isomer was not detected in any of the six samples or that only one isomer within the fatty acid isomer group was detected, such as C22:6n-3. The colour scale represents statistical significance of the rejection of the null hypothesis that the respective cell lines exhibit no different relative abundances of fatty acid isomers. P values were calculated with a custom python script employing the python package scipy.

**Table S11:** Fold changes in relative abundance of selected fatty acid isomers between cell lines MCF7 and LNCaP as well as between LNCaP and LNCaP_SCD-1*i*.

| Correlated desaturase<br>(Proposed) | FA isomer desaturation product | MCF7 to LNCaP |  | LNCaP to LNCaP_SCD-1*i* |  |
| --- | --- | --- | --- | --- | --- |
|  |  | Fold change | P value | Fold change | P value |
| SCD-1 | FA 14:1*n*-5cis | 0.36 | 0.061 | 0.21 | 0.018 |
| SCD-1 | FA 16:1*n*-7cis | 0.34 | 0.00004 | 0.13 | 0.0043 |
| SCD-1 | FA 18:1*n*-9cis | 0.56 | 0.0018 | 0.77 | 0.014 |
| SCD-1 | FA 17:1*n*-8cis | 0.56 | 0.00019 | 0.79 | 0.11 |
| SCD-1 | FA 19:1*n*-10cis | 0.35 | 0.0006 | 0.87 | 0.41 |
| FADS2 | FA 14:1*n*-8cis | 2.2 | 0.0088 | 1.6 | 0.01 |
| FADS2 | FA 16:1*n*-10cis | 18.9 | 0.0022 | 1.5 | 0.008 |
| FADS2 | FA 17:1*n*-11cis | 6.2 | 0.0065 | 1.8 | 0.0087 |
| FADS2 | FA 18:1*n*-12cis | 28.3 | 0.051 | 7.6 | 0.0035 |
| FADS1 | FA 20:3*n*-6,9,15 | 0.023 | 0.0000008 | 2.7 | 0.048 |
| FADS1 | FA 20:2*n*-9,15 | 0.079 | 0.0061 | 1.8 | 0.041 |
| FADS1 | FA 18:2*n*-7,13 | 0.073 | 0.014 | 1.2 | 0.51 |
| FADS1 | FA 18:2*n*-10,13 | n.a. | n.a. | 2.4 | 0.0000008 |
| FADS1 | FA 18:1*n*-13 | 0.3 | 0.14 | 11.6 | 0.064 |

#### 3.7.5 Comparison of relative abundance of eicosa-docosa- and tetracosadienoic acids

**Figure S40:**
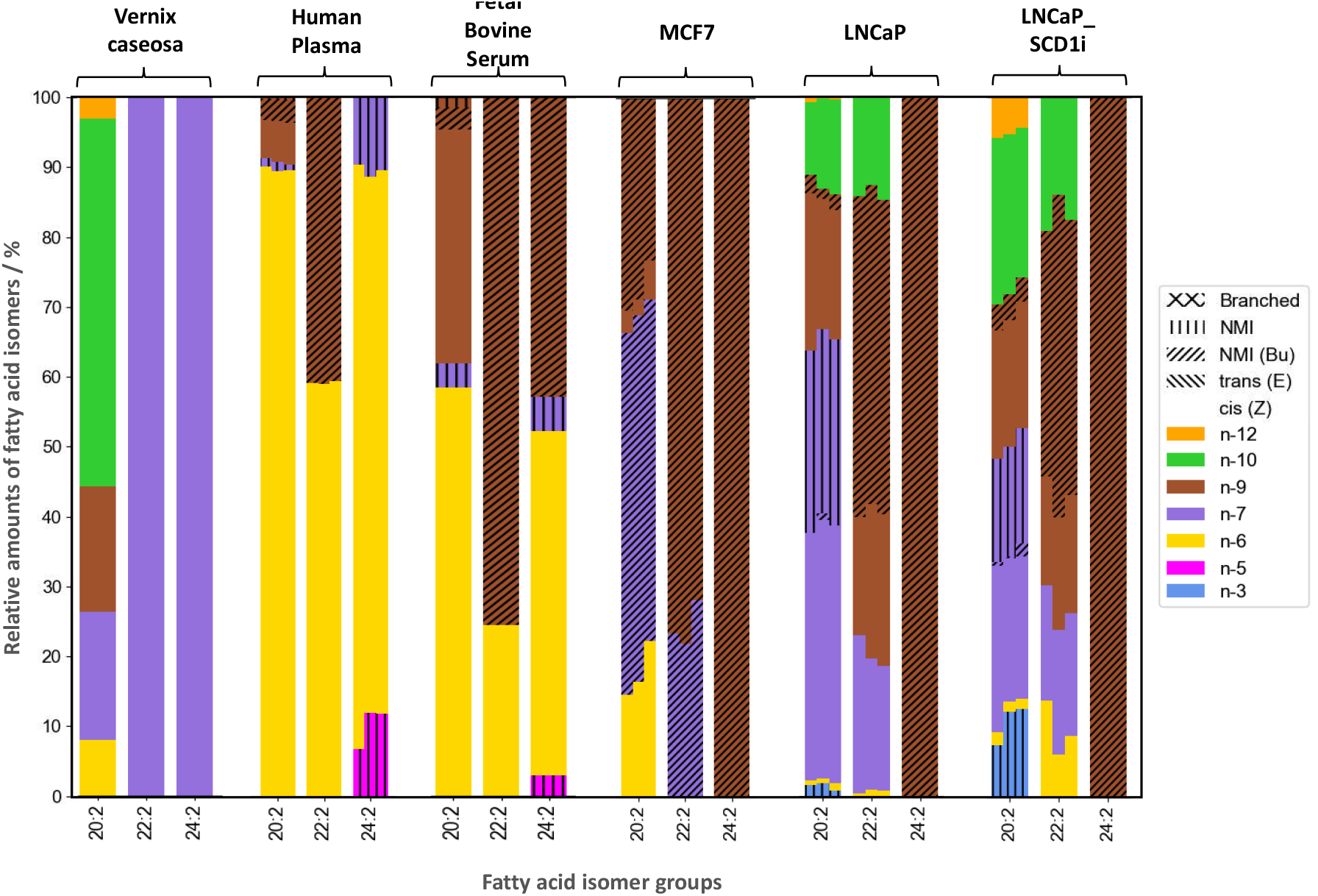
Relative abundance of selected fatty acid isomers as observed in the biological contexts studied herein. Not only changes in relative abundance of fatty acid isomers between the samples but importantly the presence of distinct desaturation patterns characterize each aspect of the human or bovine lipidome. This comparative visualization of data that is already shown above in the respective sections supports the claim that analysis of full fatty acid profiles with the OzFAD workflow has the potential to reveal previously unknown molecular markers for unique metabolism including disease specific aberrations.

